# Rapid adaptation and extinction across climates in synchronized outdoor evolution experiments of *Arabidopsis thaliana*

**DOI:** 10.1101/2025.05.28.654549

**Authors:** Xing Wu, Tatiana Bellagio, Yunru Peng, Lucas Czech, Meixi Lin, Patricia Lang, Ruth Epstein, Mohamed Abdelaziz, Jake Alexander, Mireille Caton-Darby, Carlos Alonso-Blanco, Heidi Lie Andersen, Modesto Berbel, Joy Bergelson, Liana Burghardt, Carolin Delker, Panayiotis G. Dimitrakopoulos, Kathleen Donohue, Walter Durka, Gema Escribano-Avila, Steven J. Franks, Felix B. Fritschi, Alexandros Galanidis, Alfredo Garcia-Fernández, Ana García-Muñoz, Elena Hamann, Martijn Herber, Allison Hutt, José M. Iriondo, Thomas E. Juenger, Stephen Keller, Karin Koehl, Arthur Korte, Pamela Korte, Alexander Kuschera, Carlos Lara-Romero, Laura Leventhal, Daniel Maag, Arnald Marcer, Martí March-Salas, Juliette de Meaux, Belén Méndez-Vigo, Javier Morente-López, Timothy C. Morton, Zuzana Münzbergova, Anne Muola, Meelis Pärtel, F. Xavier Picó, Brandie Quarles-Chidyagwai, Marcel Quint, Niklas Reichelt, Agnieszka Rudak, Johanna Schmitt, Merav Seifan, Basten L. Snoek, Remco Stam, John R. Stinchcombe, Marc Stift, Mark A. Taylor, Peter Tiffin, Irène Till-Bottraud, Anna Traveset, Jean-Gabriel Valay, Martijn van Zanten, Vigdis Vandvik, Cyrille Violle, Maciej Wódkiewicz, Detlef Weigel, Oliver Bossdorf, Robert Colautti, François Vasseur, J.F. Scheepens, Moises Exposito-Alonso

**Author notes:** Co-first authors (Xing Wu, Tatiana Bellagio, Yunru Peng, Lucas Czech, Meixi Lin). Genomics of rapid Evolution to Novel Environment network consortium. See Appendix with GrENE-net participant author affiliations.

## Abstract

Climate change is threatening species with extinction, and rapid evolutionary adaptation may be their only option for population rescue over short ecological timescales. However, direct observations of rapid genetic adaptation and population dynamics across climates are rare across species. To fill this gap, we conducted a replicated, globally synchronized evolution experiment with the plant *Arabidopsis thaliana* for 5 years in over 30 outdoor experimental gardens with distinct climates across Europe, the Levant, and North America. We performed whole-genome sequencing on ∼70,000 surviving reproductive individuals and directly observed rapid and repeatable adaptation across climates. Allele frequency changes over time were parallel in experimental evolution replicates within the same climates, while they diverged across contrasting climates—with some allele frequency shifts best explained by strong selection between −46% to +60%. Screening the genome for signals of rapid climate adaptation identified a polygenic architecture with both known and novel adaptive genetic variants connected to important ecological phenotypes including environmental stress responses, *CAM5* and *HEAT SHOCK FACTORs,* and germination and spring flowering timing*, CYTOCHROME P450s* and *TSF*. We found evolutionary adaptation trends were often predictable, but variable across environments. In warm climates, high evolutionary predictability was associated with population survival up to 5 years, while erratic trends were an early warning for population extinction. Together, these results show rapid climate adaptation may be possible, but understanding its limits across species will be key for biodiversity forecasting.

---

Rapid evolutionary adaptation at ecological time scales in the wild has been documented across eukaryotes, from field mustard (*1*), barley (*2*), and Darwin’s finches (*3*), to fruit flies (*4*), stick insects (*5*), and sticklebacks (*6*). Despite evidence of rapid adaptation in natural environments, it is still unknown the extent to which rapid adaptation can rescue vulnerable populations from climate-change-driven population and species extinctions (*7*, *8*), but we still do not fully understand the tempo, dynamics, and predictability of rapid evolutionary adaptation in complex climates and organisms. The gold standard to experimentally study the dynamics of evolution has been represented by microbial long-term laboratory experiments combined with genome re-sequencing (*9*, *10*)—where evolutionary adaptation typically occurs via *de novo* mutations over thousands of generations—and by animal and plant field experiments as in fruit fly orchards (*11*, *12*) or domesticated plant trials (*2*)—where generation times are longer, population sizes are smaller, and rapid adaptation often occurs via selection on standing pre-existing genetic variation (*13*). However, to address major unanswered questions on rapid evolution to climate and population responses we are still lacking large-scale animal or plant experimental evolution replicated across continents over years.

Common garden and reciprocal transplant experiments, pioneered with plants (*14*), are powerful methods for comparing fitness across multiple genotypes of a species from different locales in the same environment. These approaches have been key tools to reveal pervasive within-species standing variation and past climate adaptation across macro-organisms (*15–17*). However, knowledge about within-species variation only teaches us about the species potential for adaptation, but is not a direct observation of adaptation dynamics or population rescue occurring in time. Here, we study the process of rapid evolution over 5 years by melding common garden experiments in multiple climates and genome re-sequencing in the annual plant *Arabidopsis thaliana*. We find evidence of rapid evolutionary responses to climate, identify novel genes, and predict short-term evolutionary trends. However, evolutionary adaptation was not possible across all environments, highlighting the importance of understanding not only the potential but also the limits of evolution under the exceptional pressures raised by climate change.

## A synchronized evolution experiment in *Arabidopsis thaliana* across climates

To understand the genetic basis of rapid evolution across a wide range of climates, we conducted a multi-year and multi-location evolution experiment in outdoor gardens, using the hermaphroditic highly-selfing annual plant *Arabidopsis thaliana* as a model species. We established the GrENE-net consortium (www.GrENE-net.org) to implement a simplified and standardized protocol for evolution experiments (see protocol, **Text S1**). Experiments were coordinated across 43 locations across Europe, the Levant, and US (**Fig. 1, Fig. S1, Table S1**), and began in the fall of 2017. Experimental sites spanned contrasting climates—from urban European environments to the likely edge of the species’ niche, the Negev desert. At each location, 12 independent replicate trays of plants were established and maintained for up to five years (*A. thaliana* typically undergoes one generation per year with a spring flowering, or two generations in some climates with spring and fall flowering). Experimental trays were filled with homogenized soil, tagged with temperature and humidity sensors, placed outdoors to grow with minimal human intervention (i.e. no watering, fertilization, or shelter), and sown with ∼15,000 seeds of the same founder genotypes: 231 *A. thaliana* accessions. These accessions were selected to represent the entire native geographic range of *A. thaliana* and were mixed at roughly equal proportions and validated by whole-genome sequencing (**Fig. 1A, Table S2**, see sequence validation in **Text S4**, **S5**, **Dataset S4, S5**, **Fig. S12, S13**). The population in each tray was allowed to reproduce naturally, allowing for the study of demography and evolution across generations and climates. In predominantly self-fertilizing populations, adaptation is expected to advance through the differential fitness of inbred lineages, whereas any outcrossing that does occur generates recombinant genotypes on which selection can act at a finer genomic scale. We note that we detected signs of outcrossing in ∼6-16% of samples (see **Text S10, Fig. S14**) in line with observations from natural populations (*18*). Out of 43 initial sites, seven sites failed due to experimental logistics, and six sites established populations that all died off within a few months, possibly due to extreme climatic conditions. Plants from the remaining 30 sites successfully reproduced for at least one generation and yielded high-quality genomic and demographic data (**Fig. S5**).

**Fig. 1.**
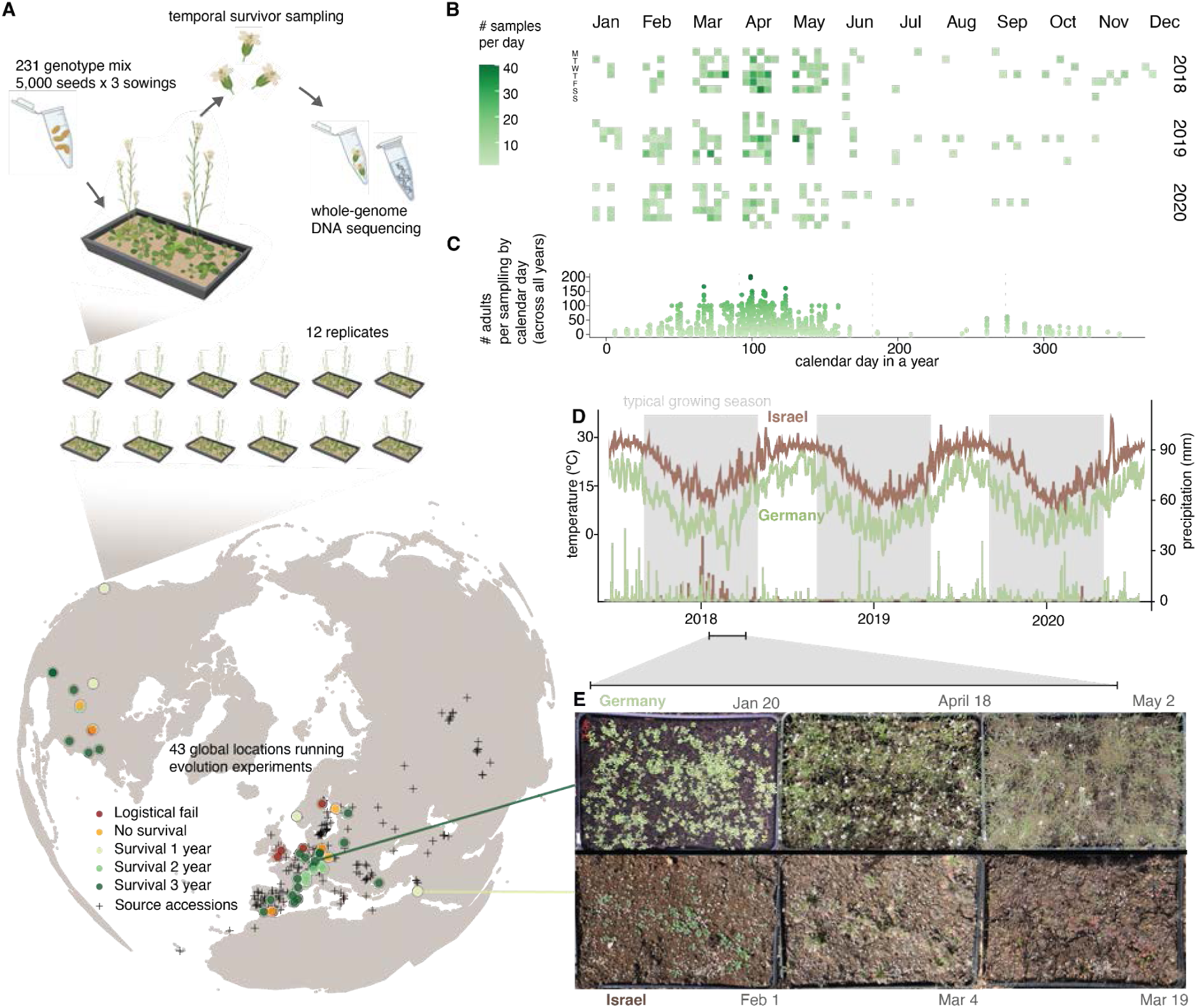
GrENE-net’s globally-distributed evolution experiment of *A. thaliana*. (**A**) GrENE-net experimental design with 231 *A. thaliana* accessions mixed in tubes of ∼5,000 seeds. Each experimental tray was sown with three tubes and seeds were spread every two weeks throughout fall 2017 to ensure establishment. Each site started 12 trays as independent experimental replicates. The map shows 43 gardens (sites) where participants started the experiment; colors indicate experiment outcomes, with 30 sites successfully completing at least one generation and producing genomic data. (**B**) Calendar of time-series collections of flower tissues used for genomic sequencing for the first three years. (**C**) Density of samples collected along the calendar year, combining data from all three years. (**D**) Daily temperatures curves and precipitation bars over the first three years of the experiment in two example locations: humid continental (Würzburg, Germany, site #46, green) and arid desert (Sde Boker in Negev desert, Israel, site #26, brown). (**E**) Example photographs of the experimental populations in Germany and Israel during spring of the first growing season.

Here, we present genomic data from the first three years of GrENE-net (2017-2020) along with complete census and environmental data (2017-2022) (Phase I data release, www.GrENE-net.org/data and **Supplemental Materials**). This includes daily climate data (**Dataset S2**), biweekly per-tray photographs from the growing seasons (**Dataset S3**), 1,141 demographic measurements (**Fig. S5**, **Dataset S1**, **Table S3**) and 74,491 tissue samples of reproductive plants collected and sequenced in 2,415 pools (**Fig. S4**, **Fig. S2, Table S5**) (*19*). Each genomic DNA library was generated from 1 to ∼200 pooled flowers collected from a single tray of surviving plants at standardized sampling times, providing a snapshot of the genetic makeup of the reproductive population (**Fig. 1A-C, Text S1, S2**). For each sample, we reconstructed allele frequencies for ca. 3.2 million single nucleotide polymorphisms (SNPs) with estimated ∼0.7% error rate. To achieve this, we combined allele count information from ∼10X coverage of pool-sequencing per experimental population sample and sampling error reduction using linkage in the 231 founder accessions from high-coverage sequencing (**Text S5**, **Fig. S11, S12, S13**, **Dataset S6, S7**), following established methods from the evolve & resequence literature (*20*, *21*). The composition of 231 founder accessions in the starting seed mix was also reconstructed by high coverage sequencing of the seed mix, confirming a very even representation of ≈0.05% per accession (i.e. 1/231) with 0.1% error rate (**Text S5**, **Fig. S12**). By merging data from multiple samplings of the same tray and growing season, we generated genome-wide allele frequencies and accession relative frequencies of 738 replicated populations across 30 locations, spanning up to 12 replicate populations per location and up to three sequenced years (i.e. equivalent to ≈3–6 generations in addition to starting generation). Combining these paired environmental and demographic metadata with evolutionary trajectories, we then studied the patterns and genomic architecture of rapid evolution across climates.

## Evolution is rapid across climates in GrENE-net

To directly study rapid adaptation from standing genetic variation, we analyzed genome-wide allele frequency changes and population differentiation across all generations and experimental gardens. We reason that allele frequency trends that are significantly synchronized (increasing or decreasing) across independent population replicates within one garden (i.e. repeatable adaptation) or different gardens of similar climate (i.e. parallel adaptation) must reflect the action of natural selection.

We first measured the degree of population differentiation by comparing shifts in allele frequencies between starting founder and evolved populations across time and space using *F*_ST_ (*22*). We observed that the median *F_ST_* across all experimental gardens with respect to the founder population increased with each generation, indicating the gradual differentiation from the founder population over time (across all samples: *F_ST_* _y1 median [IQR]_ = 0.002 [0.001-0.006], *n* = 319, *F_ST_* _y2 median [IQR]_ = 0.017 [0.010-0.036], *n* = 217, *F_ST_* _y3 median [IQR]_ = 0.024 [0.010-0.055], *n* = 182, **Fig. S19**A). In addition, *F_ST_* divergence was significantly larger between-gardens compared to within-gardens (Mann-Whitney U Test *P* = 2×10^−90^, *n* = 50,403 pairwise combinations, **Fig. S19**B). To better understand population divergence across environments, we then decomposed allele frequency changes of evolved populations across all gardens using a principal component analysis and diffusion maps (**Fig. 2A, Fig. S20 S21**). The major axes of allele frequency change separate evolved populations according to the climate of the experimental gardens they were planted in, where experiments in similar climates led to similar evolutionary trajectories and vice versa (Pearson’s correlation PC1-annual mean temperature [BIO1], *r* = 0.436, *P* = 2.348×10^−10^, *n* = 193; Pearson’s correlation PC2-precipitation of wettest month [BIO13], *r* = 0.248, *P* = 4.918×10^−4^, *n* = 193).

**Fig. 2.**
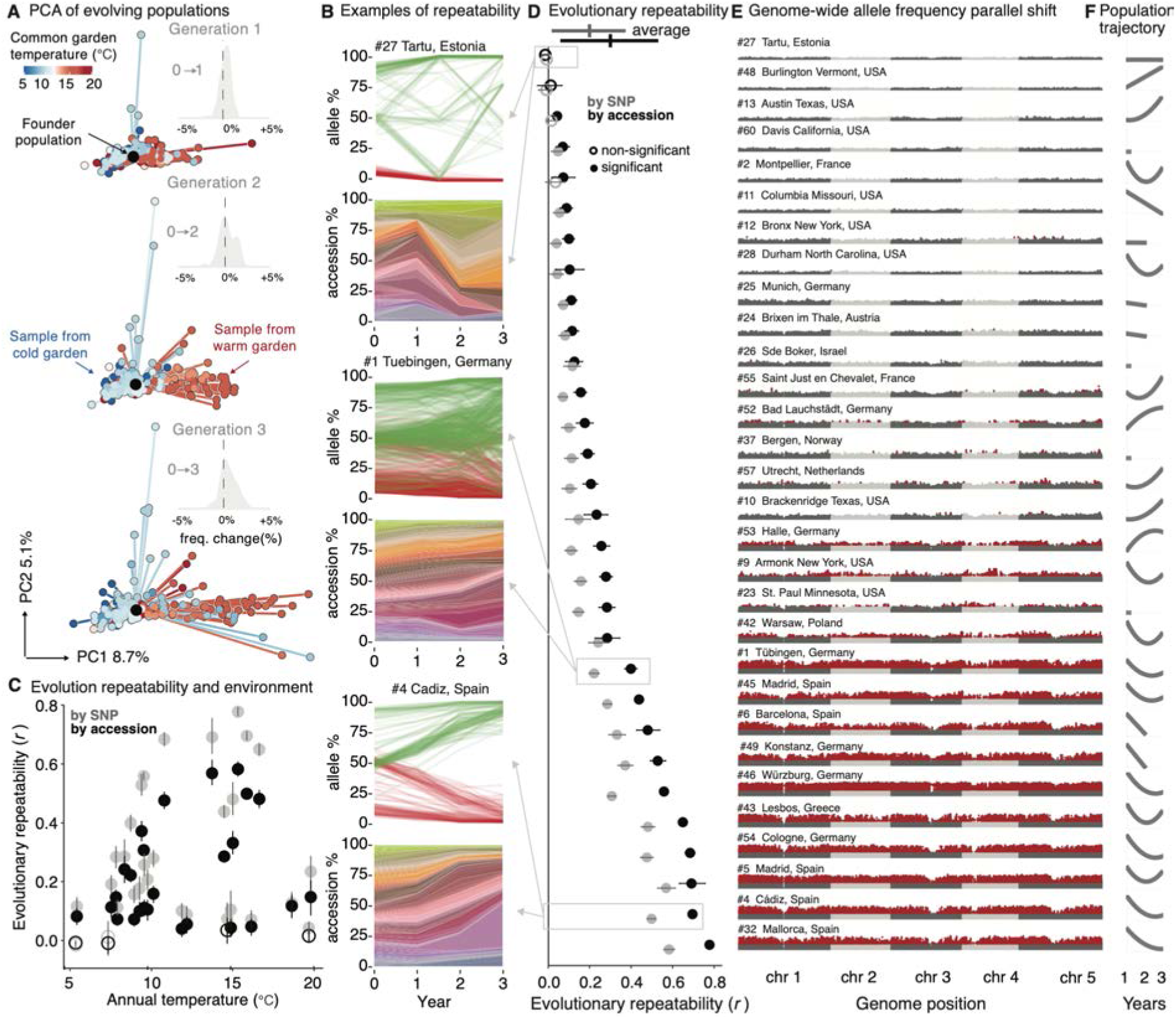
Genomic evolution in GrENE-net is rapid and parallel (**A**) Principal Component Analysis (PCA) of allele frequencies and samples over three generations with up to 12 replicates per location (n=738). The genome-sequenced founder population, common to all experiments, was projected into the PCA space (black). Insets show the distribution of genome-wide allele frequency changes between generations. (**B**) Example of three sites with low, intermediate, and high evolutionary repeatability displayed at the allele and accession level. At the allele level, the 100 fastest increasing or decreasing allele frequencies over time are plotted for illustration. At the accession level, all 231 accessions are displayed using a Muller plot with accessions sorted based on the temperature of origin from colder (green-yellow) to warmer (purple-blue). (**C-D**) Evolutionary repeatability measured at the allele or accession level as an average correlation of change in frequency from the founder frequency to first generation is displayed against the (**C**) garden annual temperature or (**D**) as a vertical rank. (**E**) Manhattan plots of Genome-Wide Likelihood Ratio Tests (LRTs) of alleles changing in frequency across 12 replicates within a site in the first generation (red indicates alleles with significant natural selection under Bonferroni correction). (**F**) Population trajectories of each location estimated across all years and replicates displaying the fitting of a polynomial regression (for expanded visualization **Fig. S8**).

To test whether the observed magnitude of evolution was significantly larger than expected from neutral genetic drift, we compared the observed variance of allele frequency changes (*Var(Δp)=(p_1_-p_0_)^2^*) with neutral evolution expectations from theory and simulations. Our rationale is that genetic drift naturally creates shifts in allele frequencies from just stochastic sampling (measured genome-wide as *Var(Δp)*). Then, if we find larger observed shifts than expectations of shifts from several stochastic demographic processes, we will need to invoke other evolutionary forces. First, comparing observed frequency variance in experimental populations with the classic Wright-Fisher population expectation (*Var_WF_=Var(Δp)_WF_=p_0_(1-p_0_)/2N*), we found on average 3-fold larger shifts across samples of different population sizes and starting allele frequency classes (*Var_observed_/Var_WF_ =* 2.99 [CI_95%_=2.94–3.04], Mann-Whitney U Test *P*<2×10^−16^; assuming *N* as the sample size sequenced, which yields a conservative test, **Fig**. **S16**, **S17**, **Text S7**). Larger allele frequency deviations would be expected if WF assumptions are violated, such as due to lack of complete outcrossing or equal reproduction contributions common in naturally evolving *A. thaliana* populations. We then conducted another set of non-WF neutral expectation, by simulating random accession sorting (i.e. no outcrossing) and unequal reproduction (i.e. uniform or Poisson distributed seed set) (see **Text S7**). We still found significantly larger observed frequency changes than expected under non-WF sorting dynamics, especially in experimental replicates of larger population sizes (**Fig. S17**), with increasing deviations of observed evolution from neutral expectations over generations (t_0_→t_1_:*Var_observed_/Var_neutral-sorting_* = 4.155 [95% CI: 2.576-5.734], t_0_→t_2_:*Var_observed_/Var_neutral-sorting_* = 9.131 [95% CI: 7.167-11.095], t_0_→t_3_:*Var_observed_/Var_neutral-sorting_* = 8.398 [95% CI: 6.584-10.211], **Fig. 2A, Fig. S15, S16, S17, Text S7, S8**).

Having established that allele frequency changes significantly depart from several neutral stochastic expectations, we hypothesize that environment-driven natural selection may have created larger, deterministic allele trajectories. Such natural selection should create repeatable allele frequency shifts in multiple replicates in the same garden (i.e. correlation of [*p_1_-p_0_*] between replicate *i* and *j*, see **Text S9**). We found significant evolutionary repeatability in 24 out of 30 gardens, as shown by high rank correlations of genome-wide allele frequency trends (**Fig. 2B-D, Fig. S24, S25, S25, S27, S28**, mean[sd] *r_ρ snp_* = 0.293 [0.237], e.g. highest repeatability site #32 Mallorca, Spain, *r_ρ snp_* = 0.778 [95% CI: 0.762–0.794]). Repeatable trends across replicates can be further used to map genomic regions with more predictable frequency. Using a likelihood ratio test (LRT-1, **Text S11**) (*23*), we found that such signals were widespread along the genome, likely due to many alleles being linked to causal loci and thus experiencing indirect selection or “genetic draft” (*24*) (**Fig. 2E**). Overlaying the selection signals onto 16,917 linkage disequilibrium (LD) blocks along the *A. thaliana* genome estimated in the founder population (see **Methods** and **Text S12**) identified 377 LD blocks that showed repeatable signals within gardens and that overlapped in 10 or more gardens (overlap higher than expected by permutation test, *P* = 10^−6^, **Fig. S31**), suggesting that adaptation through standing variation may be rapid and highly polygenic.

An additional way to evaluate selection on standing variation is by analyses of accession sorting. We reconstructed the relative abundance of the 231 founder *A. thaliana* accessions over time using allele frequencies from pool-sequencing and the genomes of the founders. We note that although we detect some outcrossing (**Text S10**, **Fig. S14**), ignoring outcrossing allows us to implement this intuitive analysis of accession frequency evolution. Muller plots reveal patterns akin to strain evolution in microbial studies (**Fig. 2B, Fig. S22, S23**) where multiple adaptive variants are competing to rise in frequency (*10*). These dynamics are expected since the starting population was rich in standing genetic variation. Following the previous rationale that deterministic trends of accession relative frequency must be at least partially owed to differences in fitness of accessions, and thus natural selection, we also found high rank correlation of accessions relative frequency showcasing repeatable trends within garden environments (**Fig. 2B,C**, mean[sd] *r_ρ accession_* = 0.194[0.179]). Another approach to the repeatability in frequency shifts used in evolve and resequence experiments has aimed to quantify the heritability of frequency changes (*25*). Having accession relative frequency in multiple replicates within a garden, heritability of frequency changes can be simply estimated using random effect regression (*H^2^ =* 12.9–79.6%*, n=*30 gardens), and indeed strongly correlates with repeatability (correlation *H^2^*–repeatability, *r*=0.93, *P*=5.3×10^−14^, see **Text S9**).

Not only did similar accessions rise in frequency in replicates within an environmental garden, but they also prospered in parallel across gardens of similar climates. For example, we see strongly parallel changes in three cold locations in Germany (mean *r*_cold_= 0.451 [95% CI= 0.459-0.443]), and three warm locations in south Spain (*r*_warm_ = 0.453 [95% CI=0.437–0.468], **Fig. S29**), indicating similar relative fitness ranks of accessions are maintained in similar environments even in geographically distinct locations (*26*, *27*). Correlations among replicates within gardens were naturally higher than correlations between gardens of similar climates (*r_cold within_ =* 0.548 [0.534–0.562] *, r_warm within_ =* 0.699 [0.682–0.716], **Fig. S29**), which may be attributed to technical factors in the experimental design (e.g. correlated experimenter temporal sampling, or dispersal among trays, although our dispersal estimates indicate <1% of seeds in a tray could be migrants, **Fig. S3**). Alternatively, these results may indicate that the environmental selection pressures are complex and unique within each garden, even if we classify several gardens as belonging to similar climates.

We conclude that patterns from genomic time series support non-neutral, natural-selection-driven evolutionary dynamics, presumably involved in rapid adaptation. Under such rapid adaptation we may expect populations that are initially maladapted would decline and then rebound as adaptive genotypes rise in frequency (*28*). By tracking population sizes through annual census (**Text S1**), eight out of 30 experimental gardens showed average significant signs of population recovery across replicates in the third generation, with U-shaped trajectories reminiscent of evolutionary rescue (**Fig. 2F, Fig. S7, S8, S20, Text S8**). Together, the significant allele frequency shifts and the U-shape population size trajectories support the notion that adaptive evolutionary rescue occurred across climates.

## Rapid evolution follows the pattern of past local adaptation

The strong evidence of rapid adaptation in our experiment is likely attributable to the fact that we drew lines from natural populations, which presumably were locally adapted to their different native conditions. We next wanted to determine if our observed rapid adaptation mimics past local adaptation to climatic conditions. Previous studies have found local adaptation in *A. thaliana* (*26*, *27*, *29–32*) and many other species (*15*, *16*). Here, we used information on each accession’s climate of origin (**Fig. 1A**) and change in frequency in the experiment (**Fig. 3A**) to determine if genotypes from matching climatic origins increased in frequency. We focused on the first generation sequencing since sampling reproductive adults is a proxy of relative fitness per accession (since *p_t+1_ / p_t_ = w / ŵ*). We indeed found a strong negative correlation between accession’s relative frequency in one generation and increasing climate distance squared across all gardens (*r_ρ_ =* −0.25*, P<* 2×10^−16^, *n* = 169,115) (**Fig. 3A**, **Fig. S32, S34, S35, S42**, see **Text S13**). To formally quantify climate-driven natural selection, we used a Gaussian stabilizing natural selection model (*17*) extended to accession frequency measurements in experimental evolution: *log(p_t+1_ / p_t_)* = *log(W_max_ / ŵ) - V_s_^−1^ (z_origin_ - z_garden_)^2^*. Here, *W_max_* denotes the accession-specific maximum fitness at the origin environment; *V_s_ ^−1^*denotes the accession-specific strength of natural selection measuring the rate of fitness decay; *ŵ* denotes garden-specific average fitness; *z_garden_* denotes the garden environment and *z_origin_* denotes the accession-specific optimal environment (assumed to be the climate of accession origin described by a chosen environmental variable. See details in **Methods**). With this framework, we quantified the strength of climatic local adaptation for each of 19 temperature and precipitation climate variables (BIOCLIM variables calculated from ERA5-land database, **Fig. S34, S40**, (*33*)). We found evidence of climate adaptation from both temperature and precipitation variables, with the strongest local adaptation signal being annual mean temperature (BIO1) (*R^2^* = 0.337, *P* < 2.2×10^−16^, *n* = 71,976, **Fig. 3A, B, Fig. S33**).

**Fig. 3.**
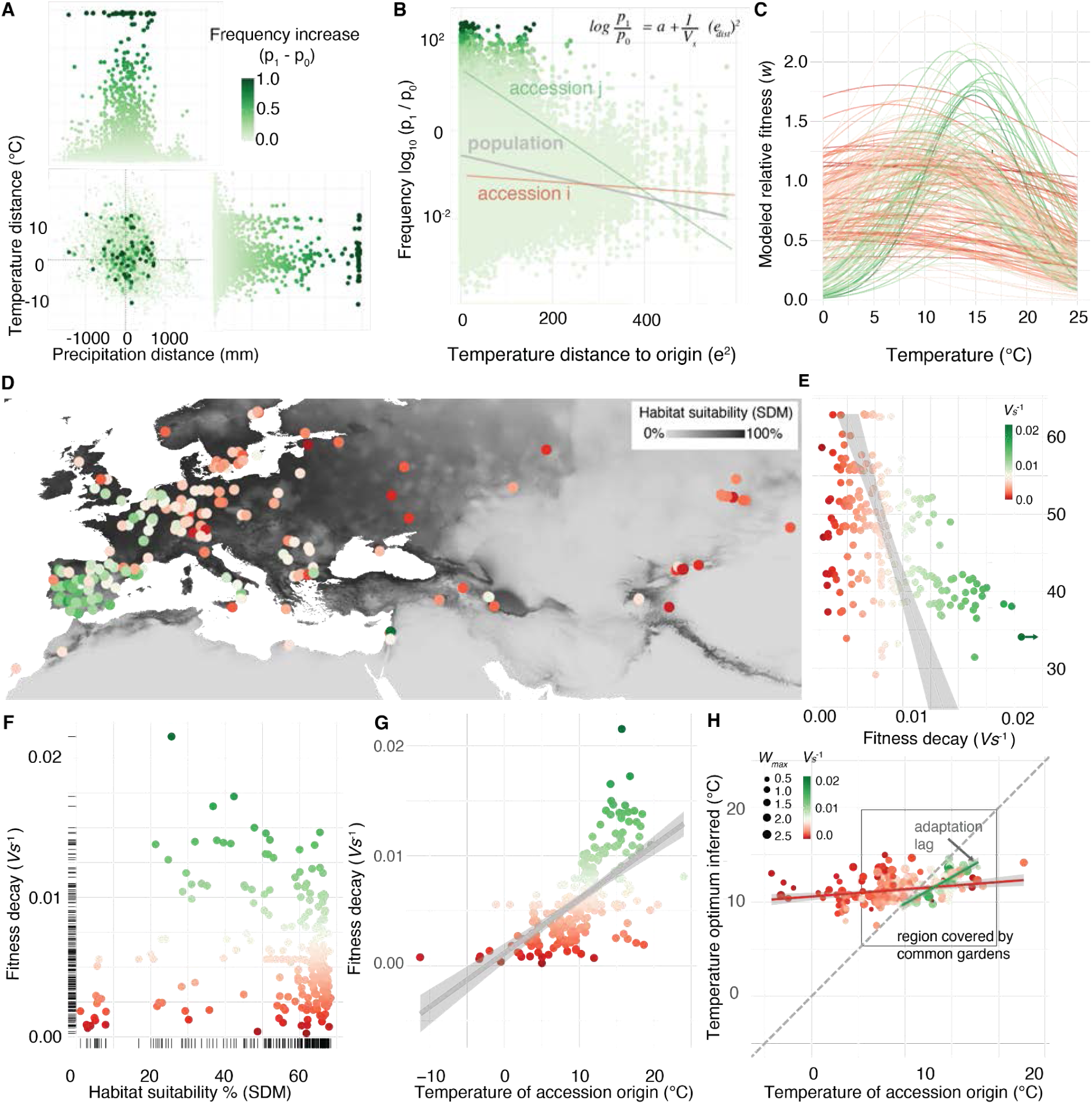
Rapid evolution follows local adaptation. (**A**) Accession relative frequency change (*p_1_-p_0_*) over climatic distance across all three years (*n* = 75,075 garden-accession origin transplant combinations) showing that planted accessions at sites with annual temperature and precipitation most similar to their home environment typically increase in frequency more than those transplanted to climatically distant environments. (**B**) Transformation of data in (A) to display *log (p_1_/p_0_)* and squared temperature distances to fit a model of stabilizing local adaptation. Grey line regression represents the average fitness decline of the GrENE-net accessions with climate distance transplant (i.e. the stabilizing selection parameter *V_s_^−1^*) while accessions *i* and *j* are examples of accession-specific *V_s_^−1^* slopes. (**C**) Idealized stabilizing selection curves for all 231 accessions based on fitting (B) equation of *V_s_ and W_max_*. (**D**) Per-accession local adaptation parameter *V_s_^−1^*visualized in a map of the accessions’ geographic collection of origin colored by habitat suitability and (**E**) across a latitudinal gradient of accession’s location origin. (**F**) Relationship between the strength of the per-accession local adaptation parameter and habitat suitability of the accessions’ locations of origin and (**G**) the accessions’ temperatures of origin. (**H**) Annual temperature averages at accessions’ origins against temperatures of gardens weighted by the accessions’ frequency, as a proxy of temperature optimum. The gray lines represent regression lines, and the shaded areas indicate their confidence intervals.

To understand possible differences between “accession niches”, we expanded the Gaussian framework to be accession-specific (**Fig. 3B, C, Fig. S39**). This revealed a trade-off between maximum fitness and rate of fitness decay across environments (**Fig. 3H, Fig. S38**) (*17*), whereby accessions’ maximum fitness correlated with more rapid fitness decay when planted in a garden with a different temperature profile (*r_ρ Wmax_* _and_ *_Vs-1_* = 0.344, *P* = 8.086×10^−8^, *n* = 231, **Fig. 3B, Fig. S41**). These results are reminiscent of the ecological trade-off observed between specialists and generalists (*27*, *34*). Consequently, we found that “generalist” accessions with wider niches (low *V_s_^−1^*) are originally from regions of lower habitat suitability at the species distribution range, and typically from colder regions (**Fig. 3E, F, Fig. S10** see **Methods**) (*17*), while “specialist” accessions with narrower but higher fitness curves appear to come from central-to-warmer native environments (*r _Vs - temp_* = 0.634, *P=* 2.2×10^−16^, *n*=231, **Fig. 3G**) (*35*). For those accessions with narrower niches (*1/V_s_ >0.15*), where the local adaptation signal is strongest, we found a notable adaptation lag (*17*, *36*), whereby accession’s realized optimum estimated from the experimental gardens was on average colder than the current climate at their geographic origin, on par with the magnitude of ∼1.5°C climate change to date (*37*) (Estimated optimum - origin temperature = −1.87°C [IQR = −0.796 – −2.84°C], **Fig. 3H**).

Because natural selection ultimately operates on phenotypes, we sought to identify the phenotypic basis of rapid adaptation across gardens. Our field observations revealed that spring flowering rapidly synchronized with the expected growth season along a latitudinal temperature gradient within three years (*r_y1_ =* 0.347, *P=* 2.89×^−31^*, n=*1060*; r_y2_ =* 0.526*, P=* 9.14×^−60^*, n=*822*; r_y2_ =* 0.554*, n =* 535*, P =* 2.67×^−44^, **Fig. S6**). Specifically, flowering periods extended to July in high latitudes and started as early as February in low latitudes (**Fig. S4**). To extend our phenotypic evolution study to phenotypes that are difficult to measure in the field, we used a curated and imputed database of traits of known heritability across *A. thaliana* accessions (*38*) with GrENE-net founder accessions (*n=*213). By correlating *ex situ* phenotype with the accession’s relative frequency change at each garden (**Text S15**), we found evidence that a number of traits likely diverged across environments (**Fig. S44**). For instance, as proof of concept, using the highly heritable flowering time measured in growth chambers (*h^2^_kinship_ =* 0.93 [95%CI 0.898-0.975]), we found accessions with known late flowering times showed a weak but significant correlation in the coldest locations (*r _freq - ft._ =* 0.074*, P =* 4.6×10 ^−6^, site #27 Tartu, Estonia, ∼5℃ mean annual temperature), while in warm locations the correlation was strong and reversed, with early-flowering accessions becoming more common (*r_freq - ft._ =* −0.25*, P =* 1.1×10^−117^, site #5 Madrid, Spain, ∼14℃ mean annual temperature). Similarly, strong seed dormancy of accessions (i.e., days of seed dry storage required to reach 50% germination, DSDS50, *h^2^_kinship_ =* 0.987 [95%CI 0.934-0.998]) correlated with accession relative frequency increase in warm, low precipitation environments (<100 mm summer rain [BIO18], Madrid (Spain), and Lesbos (Greece), *r =* 0.24–0.25*, P <* 10^−104^), where strong dormancy prevents summer germination and increases bet-hedging (*39–41*). This follows an expected growth season gradient from late flowering and high autumn germination in high latitudes to early flowering and high seed dormancy in low latitudes (*41*, *42*). Beyond phenology, we found a suite of other traits associated with climatic evolution, such as increased leaf area in cold environments (e.g. <10℃, Warsaw, Poland, *r_leaf area_* = 0.115, *P =* 2.09×10^−7^), or decreased leaf stomatal density in summer dry environment (<20 mm summer precipitation, Lesbos, Grece, *r_stomata_*= −0.07, *P =* 1.8×10^−5^) (see different phenotypes and environments: **Fig. S43 S44**, **Table S7**). Together, this supports the hypothesis that rapid adaptation trends across local environments are also driven by phenotypic evolutionary divergence.

## Mapping the genetic basis of climate adaptation

To map the genetic basis of climate adaptation, we scanned the genome for highly divergent allele frequencies across gardens. We used experimental evolution Genome-Environment Associations (eGEA) to identify SNP frequency changes across experimental gardens associated with environmental selective forces as reflected in BIOCLIM variables (see **Methods**). We used three modeling frameworks: a Latent Factor Mixed Model (LFMM) to account for population structure (*43*), a binomial GLM to account for variable population sizes, and Kendall-τ ranked correlations (see **Methods)** to detect nonlinear associations in combination with an LD block partition and *P*-value pooling with WZA (**Fig. S46, S47, S48**) (*44*). After false discovery rate (FDR) correction, we identified 44 significant blocks associated with multiple climate variables (**Fig. 4A Dataset S9**). Several blocks included genes known to affect growth, flowering and dormancy while other blocks included genes that are likely involved in environmental stress responses (**Text S17**, **Table S9**, **S10**). We then compared our experimental evolution eGEA with classic population GEA (or climate GWA). The classic GEA approach uses genomic sequencing of natural populations and directly associates genetic variants with the climate of collection of accessions to find enrichments, which requires careful population structure correction as population history correlates both with geographic and genetic patterns. This classic GEA approach has been used previously to map climate adaptation in *A. thaliana* populations (*30*). Instead, our eGEA uses the fact that standing genetic variants start at equal frequency across all experiments, and climate-driven natural selection will increase or decrease their frequencies over time. We surprisingly found little overlap in the top significant genomic LD blocks (0–3 overlapping FDR significant blocks across 19 BIOCLIM variables (**Fig. S49**, **Table S16**. See classic GEA in **Text S16** and interpretations of partial overlap). Regardless, this novel experimental evolution eGEA confirmed well-known loci or revealed novel genes important for climate adaptation.

**Fig. 4.**
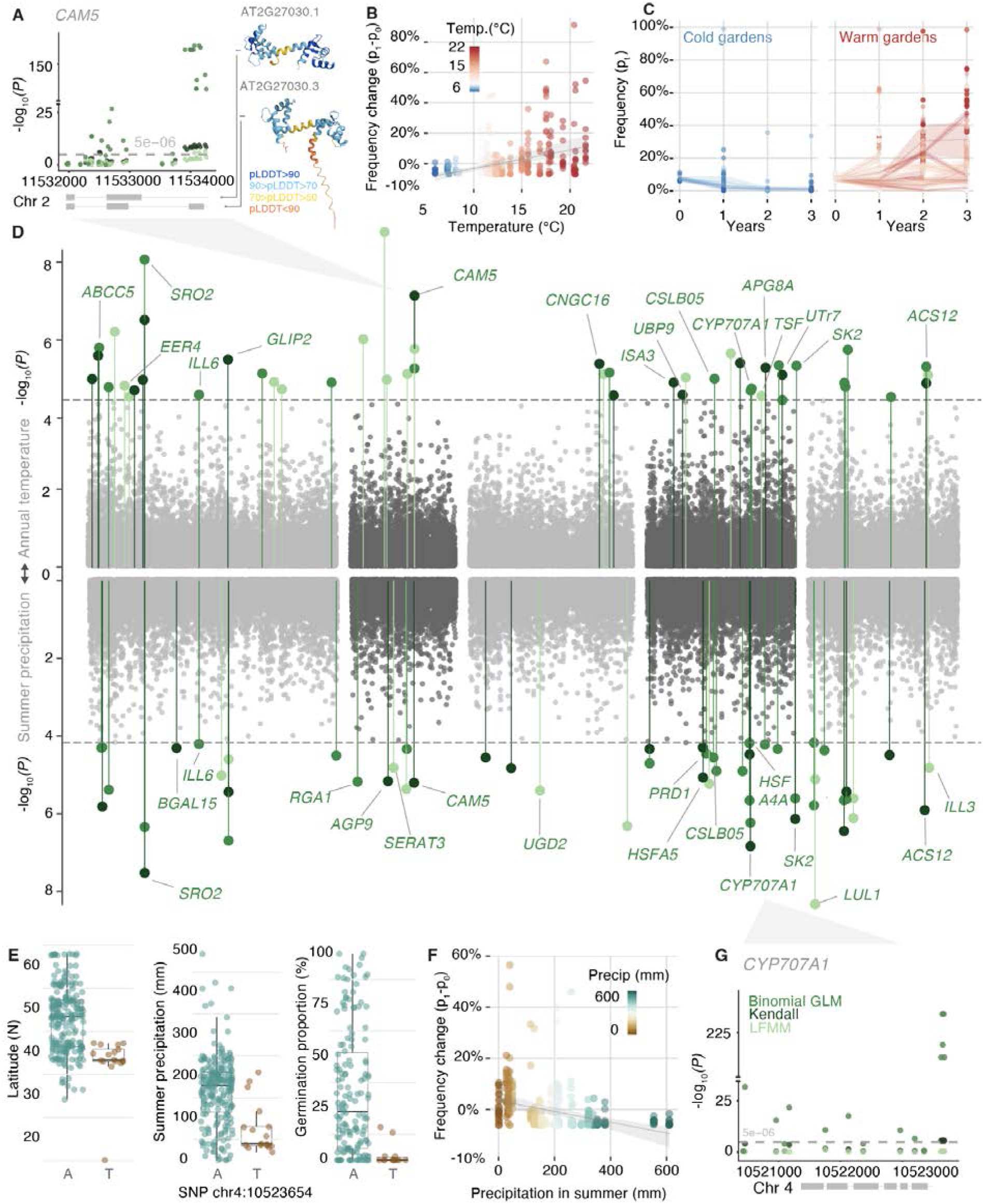
Rapid adaptation signals along the *A. thaliana* genome Experimental-evolution Genome-Environment Associations (eGEA) of rapid allele frequency trajectories with temperature (**A-D**) and precipitation in summer (**D-G**) using three statistical approaches: Latent Factor Mixed Model (LFMM), quasi-binomial Generalized Linear Mixed model (GLM), and Kendall correlation. (**A**) Zoom into the temperature Manhattan plot with *CAM5* SNP associations, reporting *P-values* obtained from the three models before inflation correction with WZA. The protein structures (AlphaFold computed) of two alternative splicing isoforms of the *CAM5* gene are depicted: isoform 1 (AT2G27030.1) and isoform 3 (AT2G27030.3). Grey boxes along the genome (x-axis) indicate two gene models of the TAIR reference genome which are present in published transcriptome data (*54*). (**B**) Example of divergent allele frequency trajectories of the *CAM5* top allele (chr4:11533937) across experimental locations along a temperature gradient. (**C**) Frequency trajectories of top *CAM5* allele over years separating experimental gardens in high (>10°C) and low (<10°C) mean annual temperature. (**D**) Manhattan plot of eGEA association of mean annual temperature (up) and summer precipitation (down) combining results from the three applied statistical approaches with haplotype block *P-value* pooling with WZA. Five *A. thaliana* chromosomes indicated in grey and black. (**E**) Relation of the top *CYP707A1* SNP allele identified in precipitation eGEA and boxplots of allele distribution relative to accession origin latitude (left) and precipitation (mid), and the expected effect of reduced germination (right). (**F**) Relation between changes in *CYP707A1* alleles and precipitation in summer. (**G**) CYP707A1 gene model and zoom into top SNP associations, reporting *P-values* obtained from the three models before inflation correction with WZA, the grey boxes along the genome (x-axis) indicate the gene model of the TAIR reference genome.

A gene well-known to be involved in spring flowering we identified in our eGEAs is the “florigen”-encoding gene *TWIN SISTER OF FT* (*TSF,* AT4G20370) (*45–47*) (LFMM-WZA block significance *P =* 3.9×10^−5^; LFMM of lead SNP, *P =* 3.63×10^−7^, Kendall *P =* 7.2×10^−10^, binomial GLM *P =* 3.55×10^−42^) (**Fig. S55, S56**, **Text S17**). We observed SNPs significantly shifting frequency across the experimental temperature gradient in the same genomic region detected in an earlier local adaptation study (*48*). *TSF* alleles were previously associated with flowering time variation within the Iberian Peninsula (Spain and Portugal) both in natural populations and common gardens (*49*, *50*). In our study, we also found that the accessions carrying the top *TSF* alleles had a significantly earlier flowering times (Wilcoxon tests, *P* < 0.05, *n* = 220, **Fig. S57**, **Table S18**).

For an annual plant, both onset of flowering and the timing of germination determine adaptation to seasonal climates, and vary strongly across the *A. thaliana* range (*41*). It is thus no surprise that we also found strong genotype-environment associations for dormancy-related genes such as *CYTOCHROME P450 (CYP707A1)* (**Fig. 4D-F**), a gene encoding an ABA-catabolic enzyme highly expressed during germination (*51*, *52*). This gene acts to promote germination by reducing ABA accumulation (*52*). The alternate *CYP707A1* allele became enriched in dry study sites (<80 mm summer precipitation) (**Fig. 4F-G**). In nature, this allele is mainly detected at lower latitudes with extremely low germination rates in the laboratory (mean germination % difference = −30%, Wilcoxon test, *n*=220, *P =* 2.648×10^−6^, **Fig. 4E**). This suggests that there has been rapid adaptation through changes in seed dormancy timing in our experimental evolution plots.

Our eGEA also identified genes that have not been previously implicated in climate adaptation. We identified a strong significant association with variation in *CALMODULIN 5* (*CAM5*) in all three eGEA methods (LFMM-WZA *P* = 2×10^−6^, Kendall’s τ WZA *P* = 1×10^−7^, binomial GLM WZA *P* = 8×10^−6^, **Fig. 4B**) (see **Text S17)**. Calmodulins bind stress-triggered calcium to modulate signaling in the context of environmental stress or pathogen responses. *CAM5* expression has been shown to be triggered by high temperature exposure in laboratory conditions (*53*). We found that the top associated SNPs are located in the intron before a third exon that is alternative spliced (**Fig. 4A**). Accessions from warm environments tend to have increased expression of a *CAM5* isoform that includes the alternative third exon downstream of the intron with the top SNP (*54*) (*r =* 0.124*, P =* 0.004*, n =* 521, **Fig. S53**). In concordance with this prediction, we find that frequencies of alternative alleles in the second *CAM5* intron increase in frequency over time in warm gardens (change rate: +1%/year, *P* = 3.58×10^−15^), and decrease in cold gardens (change rate: −1.6%/year*, P =* 1.6×10^−18^, **Fig. 4B, C**). Taking gardens in both extremes of the temperature gradient, either cold or warm, we estimated selection coefficients on *CAM5* in Cadiz (Spain) to be *s =* 57% (95% CI *s* = 49% – 66%, *p_year3_ = +*46%) and in Brixen im Thale (Austria), *s* = −47% (95% CI = −56 – −38%, *p_year2_ =* 0.2%, **Fig S52**, see selection estimation in **Text S18**). Other eGEA hits with links to stress responses include genes encoding heat shock transcription factors *(HSF4A4, HSFA5)* or an aquaporin-like protein (**Fig. 4D, Text S17**). The magnitude of environment-driven natural selection we inferred on *CAM5* was highly significant but hardly unique, with abundant polygenic signals detected along the genome (**Fig. 4D**). These findings are on par with our observed genome-wide patterns of large selection coefficients and rapid evolutionary responses, akin to those seen in fruit flies or stick insects adapting to seasonal environments (*4*, *55*).

## The direction of rapid evolution across climates is predictable

There is an urgent need to predict potential (mal)adaptation of species to future climates, both for species of conservation focus as well as domesticated species (*56*). We thus asked whether the observed changes in allele frequency (*p_0_→p_1_*) across experimental evolution gardens could have been predicted from knowledge of the genetic basis of local adaptation of the species. We reason that the climatic factors that drive differences in survival in experimental gardens likely also occur in natural populations, so we can use local adaptation signals of natural populations as a predictive signal (*57*). In agreement with this rationale, we found that alleles of warm origins showed upward frequency trajectories (*Δ p/(1-p)*/°C) in warm experimental gardens and downward trajectories in cold sites (**Fig. 5A**, *R^2^* = 0.242, *P* < 2.2×10^−100^, see **Methods, Fig. S45**). Likewise, experimental gardens in similar climates showed concordant changes in allele frequencies (e.g. Madrid vs Barcelona [Spain], *r* = 0.657, *P <* 2.2×10^−100^, and Cádiz [Spain] vs Lesbos [Greece] *r* = 0.733, *P <* 2.2×10^−100^, **Fig. 5B**), whereas gardens of contrasting climates showed opposite trajectories (**Fig. 5B** Konstanz [Germany] vs Madrid [Spain], *r* = −0.393, *P <* 2.2×10^−100^, and Warsaw [Poland] vs Madrid [Spain], *r* = −0.262 *P* = 3.8×10^−165^, *n* = 16,757 LD blocks) (see systematic analyses of antagonistic pleiotropy in **Text S19** and **Table S11**). Such a strong signal is likely driven by a combination of high polygenicity of adaptation, antagonistic pleiotropy, and genome-wide linkage disequilibrium (*27*, *58*). Using this signal, we fitted so-called “genomic offset” models that assign genotypes’ or populations’ fitness scores based on the allelic associations with climate (*59–61*) (GO_score_ = *(1/n) ∑ (|p_adapt.,i_ - X_accession,i_|),* see **Methods**). Using leave-one-out (LOO) cross-validation, we aimed to predict evolutionary trajectories for each garden (*p_0_→p_1_*) using the other gardens as genomic offset training data (**Fig. 5C, Fig. S62**). This showed substantial rank predictive accuracy across all gardens (*r_ρ_* = 0.263 [IQR 0.160–0.368], *P* = 5.752×10^−7^, *n* = 325; *r^2^* range = 0–10.9%) beyond what climate distance alone could predict (*r_ρ_* = 0.181 [IQR 0.020–0.332], *n* = 325; *r^2^* range = 0–10.1%), which is in agreement with previous findings in common gardens of *A. thaliana*, steppe grasses, and poplar tree provenances (*59*, *62*). A similar genomic offset model that additionally incorporated the Gaussian stabilizing local adaptation parameters, *V_s_^−1^ W_max_*, has a similar or higher cross-validation predictability (*r_ρ_* = 0.415 [IQR 0.311–0.544], *P* < 2.2×10^−16^, *n* = 325, *r^2^* range = 0–29.3%, **Fig. S63, S64**, see **Methods**).

**Fig. 5.**
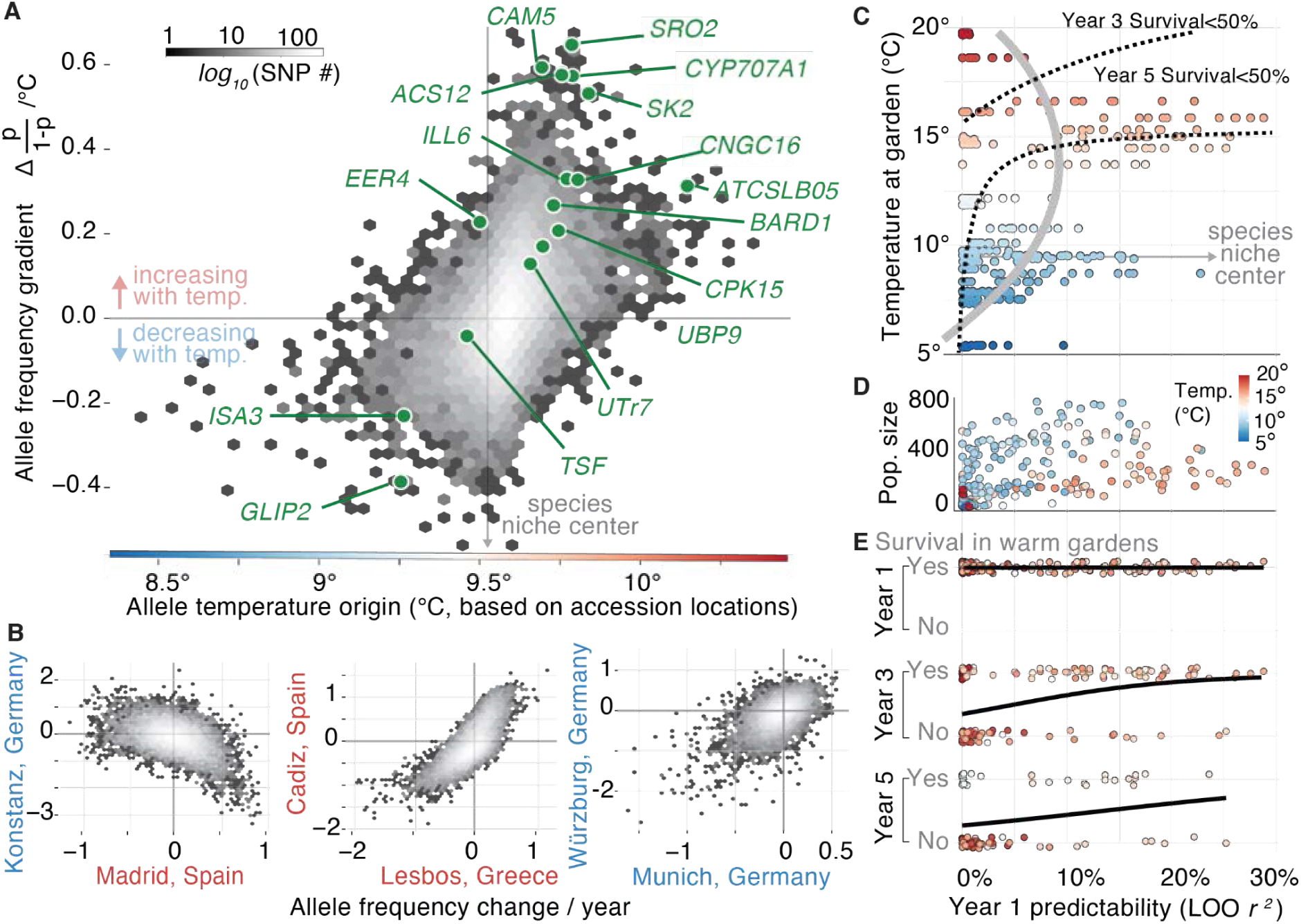
Predictability of genome-wide evolution and population survival across environments (**A**) Allele frequency changes with temperature (logistic parameter *β* = *Δp/(1-p)*/°C) and its relation to allele’s temperature origin based on the average annual temperature of the *A. thaliana* accessions carrying such alleles. Logistic regression *β* was calculated per allele and averaged within each LD block (n = 16,656). Top gene associations (Fig. 4) are highlighted in green. (**B**) Example of allele frequency trajectory over time fitting a logistic regression (*Δp/(1-p)*/year), comparing several warm (>10 °C, red) and cold (<10°C, blue) experimental gardens (**Fig. S69**). (**C**) Leave-one-out (LOO) predictability of year 1 evolutionary trends (*log(p_1_/p_0_)*) per replicate (*n* = 325) based on new genomic offset and stabilizing selection across gardens of different temperatures (see other metrics **Fig. S69**). Grey line indicates the fitted second term polynomial between predictability and temperature. Dotted lines indicate isolines of population survival from fitted logistic regressions in (E). Species niche center represents the average temperature of origin across all founder accessions (9.6°C). (**D**) Relationships between LOO predictability (year 1) and population size over time (summed total number of individuals sampled year 1-3). (**E**) Logistic regressions of LOO predictability of evolutionary trends of population replicates and survival in the 1^st^, 3^rd^, and 5^th^ years.

We ultimately predict early signs of rapid evolution to be informative about long-term population survival in changing climates. So far, evolutionary studies in the wild have typically been limited by either the breadth of climate gradients studied or by the studies’ short duration (*63*). To address this gap, we leveraged the geographic span of our experimental evolution plots and the census monitoring for up to five years. First, we tested whether predictability of early rapid evolution trends from genomic offset varies across climates. Correlating environmental data with predictability metrics, we found evolutionary predictability increased with annual temperature (14°C–17°C) but had a significant concave drop at high (>18°C) annual temperatures (regression’s quadratic coefficient = −0.001, *P* = 2.24×10^−9^, **Fig. 5D**, **Fig. 2C**). In contrast, predictability remained lower in cold and wet environments, where mortality was rare and natural selection was presumably low (**Fig. 5C**). Second, we used logistic regression-based methods to test whether evolutionary predictability was associated with survival or extinction of experimental population replicates. We found climate, evolutionary predictability, as well as their interaction to be generally significant (**5E**; logistic regression 3^nd^ year survival: *P_pred._* = 0.02, *P_temp._ =* 0.02, interaction *P_pred._* _×_ *_temp_* = 0.03; 5^th^ year survival: *P_pred._*= 0.01, *P_temp._ =* 0.02, *P_pred._* _×_ *_temp_* = 0.02; **Fig. S70**). The significant interaction indicated that evolutionary predictability correlated with increased survival especially in the warmest climates: for instance, the likelihood of population survival at ∼15°C annual temperature is over 50% only when initial evolutionary predictability is *r^2^*>15% (see isolines, **Fig. 5C**). This reminds us of eco-evolutionary tipping points that have been long theorized in population genetic literature (*64*), where in extreme environments natural selection increases mortality and overpowers the efficiency of evolutionary adaptation, leading to erratic evolutionary trends. Given natural populations of short-lived plants and animals show evidence of evolutionary and demographic responses in sub-decadal scales (*4*, *55*, *65*, *66*), our results should be helpful to downscale predictions to conditions with limited genetic diversity or less extreme climate gradients. In the future it will be key to better understand eco-evolutionary tipping points across species that may help us anticipate when species’ evolutionary responses may succeed or fail under climate change (*67*).

## Data & code availability

Supplemental Tables and Datasets are available in the online version of the paper and in www.GrENE-net.org/data and Github: https://github.com/moiexpositoalonsolab/grene. Founder genomes are available at http://1001genomes.org/data/GMI-MPI/releases/v3.1/. Sequencing Illumina reads for GrENE-net experimental evolution plots are deposited at NCBI with accession number https://www.ncbi.nlm.nih.gov/sra/PRJNA1256468. Processed frequency files are available at www.GrENE-net.org/data. Scripts to reproduce analyses and figures are available at: https://github.com/moiexpositoalonsolab/grenephase1-paper. Both intermediate data and scripts are available also at Zenodo with doi: … Software to analyze Pool-seq data: g_r_enepipe and g_r_enedalf are available on Github: github.com/moiexpositoalonsolab/grenepipe, github.com/lczech/grenedalf, github.com/moiexpositoalonsolab/hapfire. The 1001G seed collection can be obtained from the Arabidopsis Biological Resource Center (ABRC) under accession <u>CS78942</u>.

## Author contribution

M.E-A, J.F.S, F.V. conceptualized the experiment and co-led the coordination of GrENE-net. M.E-A led and coordinated this first GrENE-net paper, data production, and release including sample curation, genomic sequencing, and data analyses. O.B. R.C. and D.W. provided initial support and resources for starting this initiative at the University of Tübingen and Max Planck Institute for Biology Tübingen. All GrENE-net participants contributed to experimental design and running the experiment. M.E-A, J.F.S coordinated sample handling. Sample processing and genomic sequencing was conducted in M.E-A laboratory. Y.P. performed DNA sample processing and sequencing. T.B. and M.L. performed experimental data curations and climate data retrieval. L.C., X.W., T.B., M.L., M.E-A. conducted software development. X.W., T.B., M.L., L.C., M.E-A. performed genomic analyses. B.Q. and K.D. performed the dispersal experiment and data analyses. Data interpretation was conducted by all coauthors. The manuscript was drafted by M.E-A, X.W., T.B. and edited, commented, discussed, and improved by all coauthors.

## Acknowledgements

We are thankful for feedback from and discussions on genomic and rapid adaptation with Dmitri Petrov, Seth Rudman, Molly Schumer, and Ben Good, for discussions of intrinsically disordered domains with Alex Holehouse, and for discussions on initial project design with Magnus Nordborg. We appreciate John Kelly for his help with running LRT-1 analysis, and Tom Booker for discussions of WZA code. We thank the members of the MOILAB for feedback on the research and manuscript. We are grateful to all colleagues in the Arabidopsis field that have maintained a vibrant seed collection and shared resources.

## Affiliations of GrENE-net.org consortia authors

*By alphabetical order*

Abdelaziz, Mohamed (1), Alonso-Blanco, Carlos (2), Andersen, Heidi Lie (3), Berbel, Modesto (1), Bergelson, Joy (4), Burghardt, Liana (5), Caton-Darby, Mireille (6), Delker, Carolin (7), Dimitrakopoulos, Panayiotis G. (8), Donohue, Kathleen (9), Durka, Walter (10), Escribano-Avila, Gema (11), Franks, Steven J. (12), Fritschi, Felix B. (13), Galanidis, Alexandros (8), Garcia-Fernández, Alfredo (14), García-Muñoz, Ana (14, 1), Hamann, Elena (12, 15), Hutt, Allison (16), Iriondo, José M. (14), Juenger, Thomas E. (16), Keller, Steve (17), Koehl, Karin (18), Korte, Pamela (19), Kutschera, Alexander (20), Lara-Romero, Carlos (14), Leventhal, Laura (6, 21, 22, 23), Maag, Daniel (19), March-Salas, Martí (14, 24), Marcer, Arnald (25), Méndez-Vigo, Belén (2), Morente-López, Javier (14, 24), Morton, Timothy C. (26), Pärtel, Meelis (27), Picó, F. Xavier (28), Quarles-Chidyagwai, Brandie (9, 29), Quint, Marcel (30), Reichelt, Niklas (19), Rudak, Agnieszka (31), Seifan, Merav (32), Snoek, Basten L. (33), Stam, Remco (20, 34), Stift, Marc (35), Stinchcombe, John R. (36), Taylor, Mark A. (6), Tiffin, Peter (37), Till-Bottraud, Irène (38), Traveset, Anna (39), Valay, Jean-Gabriel (40), Van Zanten, Martijn (41), Vandvik, Vigdis (42), Violle, Cyrille (43), Westphal, Laura (18), Wódkiewicz, Maciej (31), Zerning, Dirk (18)

(1) BioChange, Department of Genetics, University of Granada, 18071 Granada, Spain
(2) Plant Molecular Genetics Department, Centro Nacional de Biotecnología (CNB-CSIC), 28049 Madrid, Spain
(3) Department of Natural History, University of Bergen, Bergen, Norway
(4) Center for Genomics and Systems Biology, New York University, New York, NY, USA
(5) Plant Science Department, The Pennsylvania State University, University Park, PA, USA
(6) Department of Evolution and Ecology, University of California Davis, Davis, CA, USA
(7) Crop Physiology, Institute of Agricultural and Nutritional Sciences, Martin Luther University Halle-Wittenberg, Germany
(8) Biodiversity Conservation Laboratory, Department of Environment, University of the Aegean, 81100 Mytilene, Lesbos, Greece
(9) Department of Biology, Duke University, Durham, NC, USA
(10) Department of Community Ecology, Helmholtz Centre for Environmental Research-UFZ, 06120 Halle, Germany
(11) Proyecto Mejora Conocimiento THIC, Grupo Tragsa – SEPI, Madrid, Spain
(12) Department of Biological Sciences and the Louis Calder Center, Fordham University, Bronx, NY, USA
(13) Division of Plant Science & Technology, University of Missouri, Columbia, MO, USA
(14) Global Change Research Institute, Rey Juan Carlos University, Móstoles, Madrid, Spain
(15) Biological Sciences, Fordham University, Bronx, NY, USA
(16) Department of Integrative Biology, University of Texas at Austin, Austin, TX, USA
(17) Department of Plant Biology, University of Vermont, Burlington, VT, USA
(18) Max Planck Institute of Molecular Plant Physiology, Potsdam-Golm, Germany
(19) Department of Pharmaceutical Biology, Julius-von-Sachs-Institute of Biosciences, Julius-Maximilians-Universität Würzburg, Würzburg, Germany
(20) Chair of Phytopathology, Technical University of Munich, Freising, Germany
(21) Department of Plant Biology, Carnegie Institution for Science, Stanford, CA, USA
(22) Department of Biology, Stanford University, Stanford, CA, USA
(23) Department of Integrative Biology, University of California Berkeley, Berkeley, CA, USA
(24) Plant Evolutionary Ecology, Faculty of Biological Sciences, Goethe University Frankfurt, Frankfurt am Main, Germany
(25) CREAF, Bellaterra (Cerdanyola del Vallès), Catalonia, Spain
(26) Department of Ecology and Evolution, University of Chicago, Chicago, IL, USA
(27) Institute of Ecology and Earth Sciences, University of Tartu, Tartu, Estonia
(28) Departamento de Ecología y Evolución, Estación Biológica de Doñana (EBD-CSIC), Sevilla 41092, Spain
(29) Department of Plant Biology, University of California Davis, Davis, CA, USA
(30) Institute of Agricultural and Nutritional Sciences, Martin Luther University Halle-Wittenberg, Halle (Saale), and German Centre for Integrative Biodiversity Research (iDiv), Germany
(31) Faculty of Biology, University of Warsaw, Warsaw, Poland
(32) Mitrani Department of Desert Ecology, Swiss Institute for Dryland Environmental & Energy Research, Ben-Gurion University of the Negev, Israel
(33) Theoretical Biology & Bioinformatics, Institute of Biodynamics and Biocomplexity, Utrecht University, Utrecht, Netherlands
(34) Institute of Phytopathology, Christian-Albrechts University, Kiel, Germany
(35) Ecology, University of Konstanz, 78457 Konstanz, Germany & Fraunhofer-Institut für Molekularbiologie und Angewandte Oekologie IME, 52074 Aachen, Germany
(36) Ecology & Evolutionary Biology, University of Toronto, Toronto, Canada
(37) Department of Plant and Microbial Biology, University of Minnesota, St Paul, MN, USA
(38) Université Clermont Auvergne, CNRS, GEOLAB, 63000 Clermont-Ferrand, France
(39) Global Change Research Group, Mediterranean Institute of Advanced Studies, Esporles, Mallorca, Balearic Islands, Spain
(40) Lautaret Garden, Université Grenoble Alpes, CNRS, Grenoble, France
(41) Plant Stress Resilience, Institute of Environmental Biology, Utrecht University, Utrecht, Netherlands
(42) Department of Biological Sciences, University of Bergen, Bergen, Norway
(43) CEFE, Univ Montpellier, CNRS, EPHE, IRD, France

## Methods Summary

### The GrENE-net consortium experiment protocol summary

The GrENE-net project follows an Evolve and Resequence (E&R) approach (*68*) and a common garden approach (*15*) to understand rapid plant evolution in different natural environments. Prior to the experiment, we assembled a GrENE-net founder population (∼5000 seeds) consisting of a roughly equal-mix of 231 natural accessions of *Arabidopsis thaliana* from Europe, Asia, and Africa. The same founder population was planted in 45 realistic outdoor environmental gardens using soil with homogenized texture and nutrient composition. In each garden, the founder population was sown in 12 independent plastic trays as replicates, each with dimensions of 40 x 60 cm. In collaboration with GrENE-net participants, we synchronized the launch of the experiment in the fall of 2017, with its first generation in the spring of 2018, and the following generations occurred relying on the population’s own capacity to reproduce offspring. GrENE-net participants sampled flower heads in each replicated plot three to four times during the reproductive season. In each sampling, GrENE-net participants pooled the flower samples from one tray into an Eppendorf tube and stored them in −80 ℃ freezer. In addition, GrENE-net participants recorded climate data with the provided environmental sensors, and recorded population size and census data by taking high-resolution photos of each tray throughout the experiment. The flower samples from each experimental garden were shipped to GrENE-net coordinators for DNA extraction and sequencing (See detailed protocol sent to participants in **Text S1**).

### DNA extraction, library preparation, and whole genome sequencing

We performed whole genome sequencing on eight tubes of seeds from the founder population (∼2,500 seeds, estimated based on weights), and 2,415 tubes of pooled flower samples from GrENE-net experimental gardens. The founder seed samples were homogenized twice using the Quickprep adapter in a FastPrep-24 machine (MP Biomedicals, Irvine, CA, USA) with 6 m/s for 40 seconds each time. Homogenate was incubated at 65℃ for 10 minutes and DNA was extracted following the standard Qiagen DNeasy Plant Mini protocol described previously (*19*). The frozen field flower samples were homogenized using a TissueLyser II (Qiagen, Hilden, Germany) at 22 s^−1^ for 35 seconds until the flowers appeared as greenish white powders. DNA was extracted from the pulverized flower samples using a custom protocol (*20*) adapted from the widely used 2x CTAB protocol (*69*).

The library preparation and sequencing steps for the seed samples followed the protocol described in (*20*). The flower DNA extracts were processed into Illumina sequencing libraries using a modified Nextera protocol (*70*). Up to 96 libraries were prepared in one batch with combinatorial indexing and pooled into one multiplexed library. In total, 26 multiplexed 96-well plates with a total of 2,415 sequencing libraries were generated from the GrENE-net flower samples collected from 2018 to 2020. Each multiplexed library plate was sequenced on one Illumina HiSeq 3000 2×150 lane, and several plates were sequenced twice. Because GrENE-net samples were precious, we used sample randomization to avoid batch sequencing effects, where temporal samples from all replicate plots within one site were never included in the same 96 well plate to avoid complete losses of evolutionary timelines in one environmental location.

### GrENE-net founder reference panel construction

Of the 231 GrENE-net founder accessions, 225 accessions were from the Arabidopsis 1001G project (*35*), three accessions were from the Arabidopsis 1001G pilot sequencing (*71*), and two accessions were collected from Israel to include more relict populations, and one accession was from the Arabidopsis RegMap project (*72*). We downloaded the paired-end sequence files of three accessions from the 1001G pilot project. In addition, we extracted genomic DNA from two Israel accessions, and performed 150 bp paired-end Illumina whole-genome DNA sequencing. We processed five sequence files following a plant variant calling pipeline (*73*), and used the Arabidopsis 1001G VCF file as the reference panel for variant calling. We then merged the Arabidopsis 1001G VCF, Arabidopsis RegMap VCF and the newly generated five-sample VCF using BCFtools (*74*). We then used Beagle v5.2 to impute the missing genotypes (*75*) and phase SNPs (*76*) in the merged VCF. After imputation, we only kept bi-allelic SNPs whose minor allele count was above 7 and heterozygosity less than 1%. Lastly, we subsetted the merged VCF to 231 GrENE-net founders, yielding a final GrENE-net founder reference panel with 231 Arabidopsis accessions and 3,234,480 high-confidence SNPs.

### Allele frequency inference and founder accession reconstruction

We used the g_r_enepipe workflow (*20*) for processing pooled sequencing reads for all 2,415 GrENE-net sequencing libraries, as well as the 8 seed samples of the founder population. In g_r_enepipe, we used Trimmomatic (*77*) for adapter and quality trimming, BWA-mem (*78*) for aligning trimmed reads to Arabidopsis reference genome TAIR 10.1 (*79*, *80*), and Picard (https://broadinstitute.github.io/picard) and samtools (*81*) to remove PCR duplicates and sort and index BAM files. To be able to quickly manipulate allele frequency over two thousand genome sequencing samples, we developed a command line C++ tool g_r_enedalf, akin to bcftools for Pool-seq (*22*) that corrects for finite population sampling when computing statistics.

When information about founders of experimental populations is known, whole-genome linkage information has been shown to help infer allele frequency, making the effective sequencing coverage much larger than the true observed genome-wide coverage (*21*). We developed a Python-based program, hapFIRE (github.com/moiexpositoalonsolab/HapFIRE), for inferring SNP and founder accession frequencies from BAM files. One major assumption of HapFIRE is that the reference panel includes all haplotypes and no new haplotypes emerged in the pool sequencing data. In other words, we assume no recombination of reference haplotypes, nor invasion by local *Arabidopsis*. Violation of this assumption will have a bigger impact on frequency estimation of the founders than SNPs. In brief, hapFIRE takes a known phased reference panel of genomic variants and identifies genome-wide independent haplotype blocks. In each haplotype block, hapFIRE relies on harp (*82*) to estimate frequencies for all unique haplotypes based on read alignments. hapFIRE then uses the haplotype frequency to calculate the frequencies of SNPs and individuals in the reference panel (see detailed description about hapFIRE in **Text S4**). The underlying concept of hapFIRE is similar to HAF-pipe (*21*), but utilizes haplotype blocks instead of simple fixed-sized windows, in order to increase accuracy of the allele frequency estimation. Once re-inferred, allele frequencies can be analyzed without much concern for sequencing depth, but finite sample size still must be taken into account. Hence, g_r_enedalf also allows reading of hapFIRE/HAF-pipe frequency tables.

### Assembling accession and allele frequencies

Of the 2,415 genomic libraries, 2,169 passed quality tests. These libraries, which included temporal samples within one generation and site, were merged into a single population genomic composition average for one experimental garden and tray replicate within the same year. To do this merge, we calculated the weighted average of per-sample frequencies for the sample tray in the same generation, weighted by the number of flowers collected from the tray in the generation. The resulting merged frequencies have 738 entries representing the average frequency of the evolved founder population in a given generation, experimental garden and replicates. The SNP and individual frequencies of the founder population were calculated by taking the average of eight founder seed samples.

### Assemble environmental data

We obtained ERA5-Land climatic data from the Copernicus Climate Change Service (https://cds.climate.copernicus.eu) for two distinct datasets: GrENE-net founders and accessions in the Arabidopsis 1001G project (1995–2005) and experimental gardens (2018–2022), based on their geographic coordinates. We transformed this data into BIOCLIM variables (*83*) typically available via WorldClim v2 (*33*) using dismo R package (https://rdrr.io/cran/dismo/). We also used an in-house Python script to process the iButton temperature sensor results for GrENE-net gardens (See details in **Text S1**).

### Analyses of allele and accession frequency changes in GrENE-net

We identified several lines of evidence of rapid evolution in GrENE-net at both genetic and individual levels. In order to alleviate the redundancy due to high linkage disequilibrium (LD) in *Arabidopsis thaliana*, we performed LD-pruning on the GrENE-net reference panel using PLINK v1.9 (*84*) with option: −-indep-pairwise 1000 100 0.1, and then we calculated the per-generation frequency change, Δ*p_t→t+1_*, for 13,985 LD-pruned SNP sites. We then applied two dimensionality reduction techniques, PCA (prcomp function in R) and diffusion maps (*85*) on the LD-pruned Δ*p_t→t+1_* to visualize the differentiation of the founder population in different gardens per generation. To do this, we constructed the PCA and diffusion map for the third generation, and then projected the first and the second generation data to the third generation axes.

We next examined the distribution of average frequency change per evolved population, for both LD-pruned genome-wide alleles and founder individuals, and compared them to the simulated null distribution under random drift assumption, to inform the dynamics of natural selection and drift during GrENE-net. To simulate the null distribution of allele frequency change under random drift, we simulated 500 populations by random uniform sampling of founder individuals for three consecutive generations under the assumption of random drift, and using the real GrENE-net population size and reference panel. For each population, individuals in the current generation were randomly sampled, with replacement, from the previous generation population. For example, the first generation individuals were randomly and proportionally sampled from the founder population. The second generation individuals were then randomly and proportionally sampled from the first generation pool based on the proportion of each founder individual, and so on and so forth. The mean and variance of allele frequency change for each simulated population, for both LD-pruned SNPs and founder accessions, were then estimated for each generation.

Leveraging 12 independent replicates in each experimental garden, we calculated the evolutionary repeatability for each garden as a proxy for experienced strength of natural selection. The evolutionary repeatability is defined as the average of pairwise Pearson correlations of frequency changes, Δ*p_t→t+1_,* among all replicate combinations within one experimental garden. We first calculated the pairwise Pearson correlation coefficients between any two replicates in the same experimental garden based on LD-pruned genome-wide SNP frequency and founder individual frequency change, and then we used a Bootstrap resampling approach (*n*=10000) to estimate the average and 95% confidence interval repeatability.

### Models of allele frequency over time

To test whether alleles significantly changed in frequency coordinatedly across replicates, we used a likelihood ratio test, LRT-1 (*86*), to identify parallel-shifted genomic variants in response to selection across 12 replicates, accounting for sampling noise. This test systematically outperformed others in a recent benchmark (*87*). We used hapFM (*88*) to partition the Arabidopsis reference panel into haplotype blocks, and then assigned parallel-shift blocks if there was at least one SNP in the block that was significant (after Bonferroni correction of *P*-values) in the LRT-1 test for each experimental garden. We next summarized the overlapping parallel-shift blocks across all experimental gardens, and used a permutation test to determine the null distribution of overlapping blocks. The significant threshold is defined at 99% quantile of the null distribution. We then extracted the genes under significant parallel-shift blocks and performed gene ontology enrichment analysis using PANTHER v19.0 (*89*).

An alternative approach to LRT-1 is to use the standard GWA framework using the change in frequency of accessions (a_t -_ a_t-1_) as a phenotype. Using GEMMA software (*90*) we identified genomic regions associated with adaptation, analyzing changes in accession frequencies from the founder to the final generation across various common gardens. Rather than traditional phenotypes, these frequency changes served as proxies for relative fitness. We employed GEMMA’s linear mixed model approach, focusing on SNPs with a minor allele frequency MAF >5%, (all results are in **Dataset S11**).

Finally, with longer time-series (e.g. 4 timepoints including founders), we can fit Generalized Linear Models (GLM) using Binomial or quasi-Binomial variances (*12*): *k*/*n ∼ a + β t +ε*. This allows us to incorporate the fact that frequency over time *p* needs to be modeled including the limited number of flowers, *n*, we sample and thus the number of observed alternative allele *k = p× n*. The resulting model provides a slope *β,* of Odds ratio change over time, *β= p/(1-p)/year*. This repeatability of frequency change over years provides an even higher confidence that alleles are under positive or negative natural selection, so we used them to test for conditional neutrality and antagonistic pleiotropy in a pairwise manner across sites (**Fig. 5**) (*91*).

### Local climate adaptation and Gaussian stabilizing selection model

In GrENE-net, we observed accession relative frequency and climatic distance between founder accessions’ origin of collection and experiment gardens, whereby accessions transplanted proximally to the garden increased the most in frequency, resembling local adaptation.

To formally model the strength of local adaptation we extended the Gaussian phenotypic stabilizing selection framework from evolutionary biology and quantitative genetics (*92*). Under this framework, an individual (accession) *i* growing in environment *j*, has a fitness value, *w_ij_*, which decays following a Gaussian distribution as the individual is further away from an optimum:

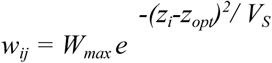

Where *W_max_*is the maximum fitness at the selection optimum, *z_opt_*, and phenotype of the *i* accession, *z_i_,* results in a decrease in fitness proportional to the variance or width of the Gaussian curve, dictated by *V_S_*, also thought of as the strength of natural selection parameter.

Log-transformations help reveal that the log fitness of an accession decays linearly with squared environmental distance with *V_S_ ^−1^* as the slope of fitness declines.

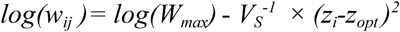

This framework has also been utilized in the context of local adaptation and reciprocal common garden studies to model fitness with transplant distance (*93*). This expansion of stabilizing natural selection to local adaptation assumes different environments having different local optima, *z_j garden_*, and natural accessions have evolved to their environment of origin optima*, z_i origin_*. A descriptor of common garden environments and collections of origin, such as a climate variable like temperature can then be used in the same framework for garden *j* and accession *i* pairs:

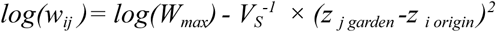

In GrENE-net, we did not directly observe a fitness value but rather relative frequencies after a period of time. However, systematic changes in relative frequency over one generation (*t→t+1; or t=0→t=1*) in an accession *i* in a location *j* ought to be due to differences in relative fitness: *p_i t+1_ = (w_ij_ / ŵ_j_) p_i t_*; where *ŵ* represents the mean fitness of a population (this expectation follows from a Wright Fisher population (discrete generations, infinite population size, etc.). By dividing the aforementioned local adaptation selection function by the mean fitness of a population *ŵ_j_* we can get:

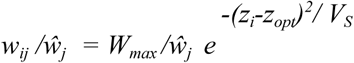

and thus:

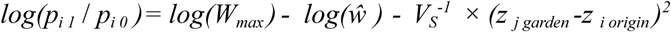

This final equation now links the frequency ratio of founder accessions in two consecutive generations in a specific environment with its maximum fitness, average fitness of the population in the environment, and the climatic distance between the growing and optimal environments.

This first model can be fitted to the GrENE-net data, as response variable (*y= p_i 1_*/ *p_i 0_*) and predictor climate distance variable (x= *z _j garden_ - z _i origin_*) are known. The parameters can be estimated using an Ordinary Least Squares (OLS) regression where the intercept is: *a=log(W_max_) - log(ŵ)*; and the slope is: *b= V_S_ ^−1^*.

This model can further be extended to be more realistic and account for accession-specific differences in fitness decline over transplant distance or maximum fitness. To do this, we reformulate the equation as a linear mixed model (LMM) with random intercepts and random slopes using MCMCglmm package (*94*) in R. Specifically, we idealize the fitness slope to have a accession-specific deviation, *β = V_S i_^−1^,* the maximum fitness achieved to have a accession-specific deviation, *α_i_* = *log(W_max i_)*, and the average fitness of a population to have a location or garden specific deviation, *γ_j_=log(ŵ _j_)*. Of course, every observation *k* will have its specific residual deviation too, *ε_ijk_.* We then fit a MCMCglmm as:

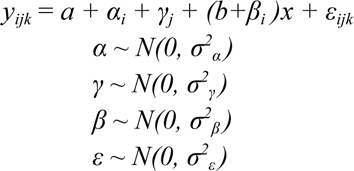

In MCMCglmm, we used 30000 iterations for MCMC chains and 5000 iterations for burn-in.

### Experimental evolution Genome-Environmental Associations (eGEA) and classic GEA/climate GWAs

We conducted Genome-Environmental Association (GEA) analyses to explore the relationship between changes in SNP frequencies and climatic selective forces across experimental gardens, aiming to identify genomic regions and putative genes responsible for climate adaptation. We assessed allele frequency shifts at two key points: after the first generation (Δpt→t+1) and at the final generation (Δpt→t terminal) of each common garden experiment.

We correlated these allele frequency changes (Δp) with the 19 BIOCLIM variables at each common garden. The BIOCLIM variables were computed from the ERA5-Land dataset downloaded from the Copernicus Climate Change Service (https://cds.climate.copernicus.eu). We used 2018 data for the first generation analysis and data spanning 2018-2020 for the terminal generation analysis.

Our analysis employed three established GEA models to identify adaptive genomic regions across environmental gradients in natural populations (*59*, *95–99*):

1. Latent Factor Mixed Model (LFMM) (*43*): This model accounts for population structure by estimating unknown confounding factors (the latent factors). We conducted a Principal Component Analysis (PCA) on the allele frequency matrix, and based on the scree plot and the elbow rule, we used 17 latent factors for the first generation analysis and 24 for the terminal generation analysis.
2. Binomial Regression: This model evaluates each SNP for the probability of observing the alternative allele relative to an environmental variable while accounting for sample size (population size). The model’s formula is log(p_i_ / 1 - p_i_) = *B*_0_ + B_1_X + E, where p represents the alternative allele’s probability, *B*_0_ is the intercept, B_1_ shows the environmental impact, and Xi is the environmental variable. This model was implemented using the Python library Statsmodels.
3. Kendall-τ Rank Correlation: This method assesses nonlinear associations by correlating the changes in allele frequencies of each SNP with each BIOCLIM variable. This model was implemented using the Scipy Python library.

From the initial dataset we filtered SNPs to ensure robust statistical analysis to a minimum allele pseudo-count threshold of 5%. We narrowed down the dataset from 3.2 million SNPs in the founder population to 1,054,574 SNPs in the first generation dataset, and further to 1,048,635 SNPs in the terminal generation dataset. To leverage the increased linkage disequilibrium expected around sites subject to local adaptation and enhance power, we used a window-based method and partitioned SNPs into 16k unique haplotype blocks (*88*) to then transform SNP-based p-values from our three models into haplotype-based p-values with WZA (*44*, *95*).

#### Climate GWAs

We used the climatic origin of each accession as phenotypes, defined using each of the 10 BIOCLIM variables calculated from ERA5-land data. We filtered SNPs with a minor allele frequency (MAF) less than 0.05 and used the same window-based approach, WZA, (*44*) used in the GEA models for direct comparability.

### Relationship between allele’ climatic origin and rapid trajectories in experimental evolution

To estimate the climatic origin of each haplotype block, we used the mean annual temperature (BIO1) of the sites of origin for each *Arabidopsis* accession (*e*) and then multiplied the genotype matrix (G), where rows represent accessions and columns represent SNPs, by the centered temperature vector (*e*). This dot product, G*^T^e* yielded a vector representing the association of each SNP with temperature. To obtain the average climatic origin of each haplotype, we normalized this vector by the number of accessions G*^T^e /n* and then averaged alleles within a block to finally estimate the average climate where a haplotype is geographically found.

To estimate the haplotype frequency change in response to temperature changes, we used the binomial regression coefficient β estimated from the third generation data (see **GEA methods**). β relates the changes in allele frequencies with the temperature gradient (BIO1) across experimental sites (β = Δp/(1-p)/°C). Finally, we averaged β within each haplotype block to obtain the estimated haplotype-level change.

For the regression analysis between haplotype climatic origin and their frequency change across the climatic gradient in the third generation of the GrENE-net experiment we used 16,665 independent haplotype blocks. This subset was derived from the initial 16,917 founder haplotype blocks after filtering for a minor allele pseudo-count of 5% in the third generation.

For the correlation analysis between haplotype trajectories at contrasting and similar experimental climates, we used 16,757 independent haplotype blocks. This subset was derived from the initial 16,917 founder haplotype blocks after filtering for a minor allele frequency (MAF) threshold of 0.05.

### Predictability under leave-one-out cross-validation

We employed four approaches to predict the evolutionary trajectories of founder accession frequencies in GrENE-net across three generations and 30 sites. To account for differences in output across these approaches, we used both Spearman’s correlation and variance explained (R²) to evaluate their performance.

#### First Approach: Climate Distance-based Prediction

In this approach, we used the climate distance between the founder origin and the experimental garden. For each garden, we calculated the climate distance of annual mean temperature (BIO1) between the garden and each founder’s origin. These distances were then quantified against the observed founder frequencies in each replicate population.

#### Second Approach: Gaussian Stabilizing Selection Model

We applied a Gaussian stabilizing selection model using annual mean temperature to predict founder trajectories. A leave-one-out cross-validation method was employed, where data from 29 gardens were used to train the model, and the remaining garden was used for testing. This process was repeated for all 30 gardens. In each training-testing split, we estimated accession-level parameters (W_max and V_s) using data from the 29 training gardens. The evolutionary trajectory of accession relative frequency in the testing site was then predicted using Equation (4).

#### Third Approach: Generalized Linear Model (GLM) Based on GEA

We applied a binomial Generalized Linear Model (GLM) based on the learned relationship between allele frequency changes and environmental variables from Genome-Environment Association (GEA) models. Using a leave-one-out cross-validation approach across the 30 experimental gardens, we trained the GLM on allele frequency changes observed in 29 gardens across the mean annual temperature gradient (BIO1), while the remaining site was reserved for testing. The GLM predicted the probability of detecting the alternative allele at the test site. We calculated a genomic offset for each accession at the test site, analogous to the Risk of Non-Adaptiveness (RONA) concept (Rellstab et al., 2016) but adapted for accessions. This was expressed as: *RONA_accession_ = (1/n) ∑ (|p_adapt.,i_ - X_accession,i_|)* where *p_adapt.,i_*is the predicted optimal frequency of the alternative allele at the *i* SNP at the given environment based on the evolved GrENE-net populations, and *X_accession,i_*is standardized accession genotype, *X*∈ *(0,0.5,1)*, at the *i* SNP. We also tested the effect of the number of SNPs included in the model and its effect on prediction accuracy, and found that increasing the number of SNPs included in the calculation of genomic offset enhances the Spearman correlation with actual relative fitness of accessions (**Fig. S63**).

For genomic offset we also could calculate the distance of the founder population based on starting frequency of all alleles (*p_0,i_*) to the predicted ideal local adaptation frequency at a given environment (*GO_founder_=(1/n) ∑ (|p_adapt,i_ - p_0_*_,*i*_*|)*). This would ideally provide a metric of overall founder population maladaptation to the transplanted environment.

#### Fourth Approach: Replicate-Based Predictions as the Positive Control

In the final approach, we leveraged replicate populations within the same experimental garden. Each garden contained up to 12 replicated populations. To predict the founder accession frequencies in one population, we used the mean founder accession relative frequency values from the other replicates within that garden. This process was iteratively applied across all replicate populations in each garden.

## SUPPLEMENTAL TEXT

### Text S1 Detailed GrENE-net protocol provided to participants

GrENE-net participants will receive identical mixtures of thousands of seeds generated from three different greenhouse experiments to dilute maternal effects. Prior to the distribution of seeds among the participants, the coordinators have prepared a master seed mix by pooling accurately weighed amounts of seeds from the parental genotypes. Because it was not feasible to count the exact number of seeds for all accessions, seed number per accession has been estimated by weighing seeds of three accessions with contrasting seed size, from which we took the average. In total, we estimated that the seed mix contains *ca*. 3.5 million seeds from the 231 accessions (see **Table S2**). Sowing, which will be at a density of *ca*. 2 seeds / cm^2^, will take place in Autumn 2017. For up to 5 years, participants will sample pools of flower heads in each replicated plot during the reproductive season. These flower head samples will be processed centrally by the coordinators and used for pool-sequencing. In parallel, **the actual genetic composition of the seed mix** sent to participants (mean and variation), as well as **the effective population size (*N*e)** will be characterized directly by sequencing of repeated subsamples of the master mix (identical to the subsamples used for sowing in a plot).

The **pool-sequencing approach (pool-seq) is a cost-effective method to track allelic frequency changes** in populations evolving under different environments (*100–103*). When the parent lines are genetically known, as it is in our case, methods also exist to estimate haplotype (*i.e.* accessions) abundances from pooled reads (*104*). The main sources of error in pool-seq are related to low sample size (number of individuals pooled) and low genome coverage (*100*). However, previous studies indicate that when over 100 individuals are pooled, the accuracy of allele frequency estimation is equal or higher than single-individual sequencing (*100*) and the unequal representation of DNA per individual to the pool is negligible (*100*, *105*, *106*). These technical issues have been considered when defining the sample sizes and sequencing coverage in the GrENE-net protocol.

### 1.2 Experimental materials

#### Provided

- **36 Eppendorf tubes**, each containing *ca*. 0.1 g of **seeds** (approximately 5000 seeds from the 231 natural accessions in total) will be provided to each GrENE-net participant to perform three sowings in each of the 12 plots. Depending on the location and climatic conditions, participants will propose to the coordinators the expected optimal timing for sowing (ideally a relatively cold but not freezing, rainy period between September and November). Then the first 12 tubes will be sown at the beginning of that month (one in each plot), the second 12 tubes in the middle, and the last 12 at the end of the month. This procedure will likely increase the chances of seed germination and seedling establishment. On average, there will be over 20 seeds per accession per plot after the three sowings being performed. The tube bottom will be already punctured and sealed with parafilm and Scotch tape. When participants remove these covers, they will be able to spread seeds by shaking the tube, like a salt shaker (see photo). 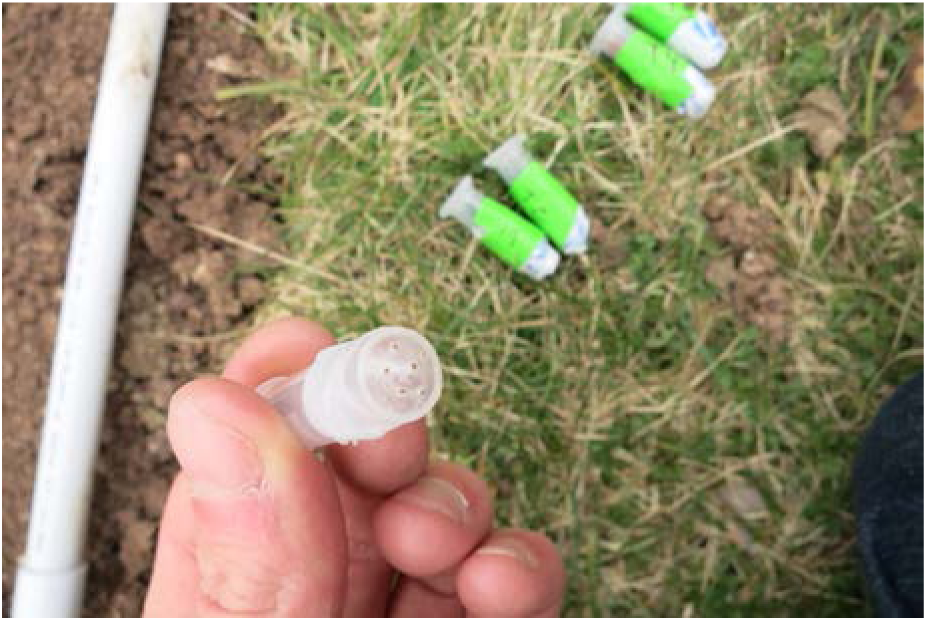
- **12 plastic trays of 60 cm width × 40 cm length × 7 cm depth with draining holes** will be provided for the 12 plots. 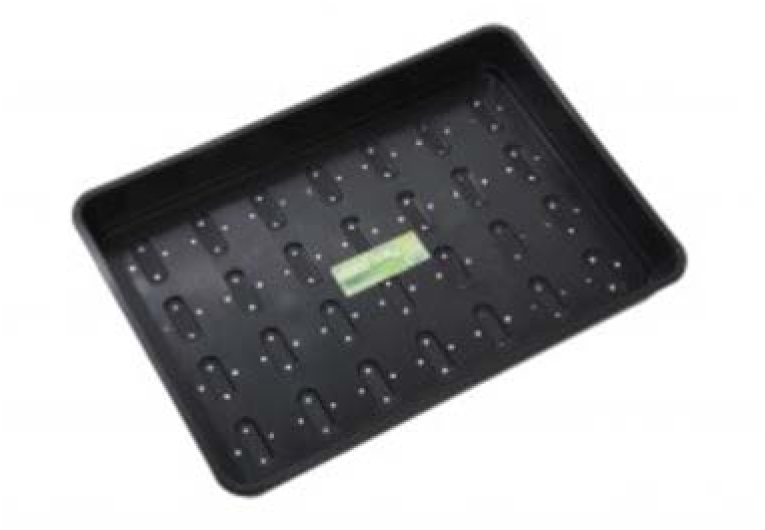
- **3** activated **iButtons**: 2 temperature loggers with plastic holder in a plastic bag and 1 temperature plus air humidity logger attached to a 20 cm wooden construction (see photo). All three iButtons have already been activated for the participants. 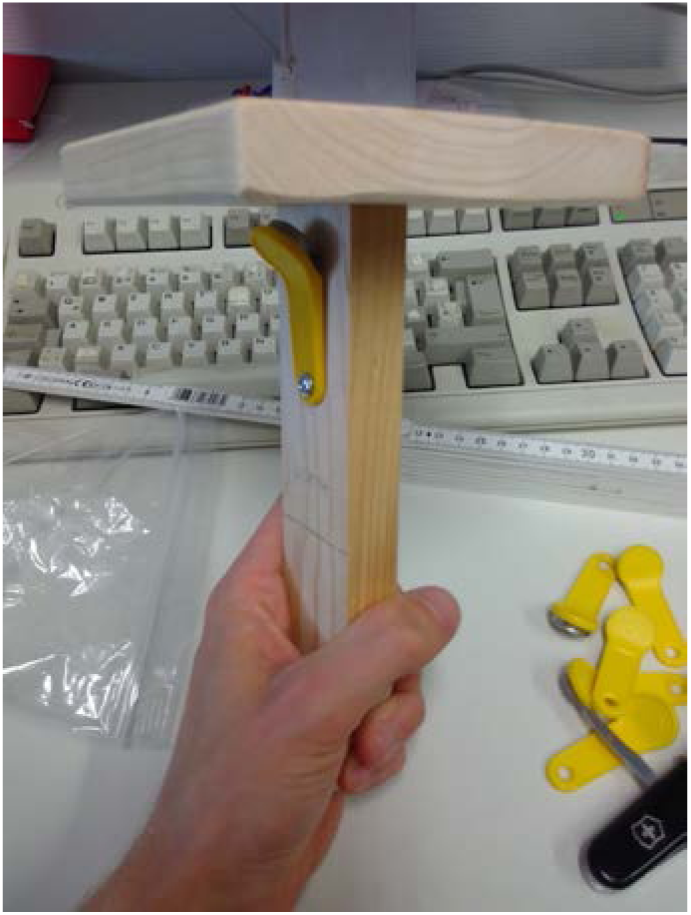
- **60 Eppendorf 2 mL** tubes to sample flowers from the 12 different plots at different times.
- Several **falcon tubes** for soil sampling (for nutrient analysis as well as seed bank analysis).

#### Required

- We will **use commercial pot soil rather than natural soil** to (1) avoid contamination from native *Arabidopsis thaliana* seeds, (2) reduce soil variation among sites since the focus is on adaptation to climatic conditions, and (3) reduce weed pressure from other species. A total of 200 Liters will be required (16L per tray). We ask participants to acquire directly <u>Brill-substrate Propagation Substrate PRO start</u> (see specifications below) or to find another 100% peat-based soil. Ideally, 500g NPK /m^3^ soil is preferred, but up to 1000g/m^3^ would also be accepted, and fine to medium-fine structure. The use of 100% peat soil assures fairly standard structural and nutrient properties irrespective of the supplier. It turned out impractical to distribute soil from a single company to all sites, which is why we ask participants to organize soil for themselves. In case you are unsure whether a particular soil would be fine, please contact us. 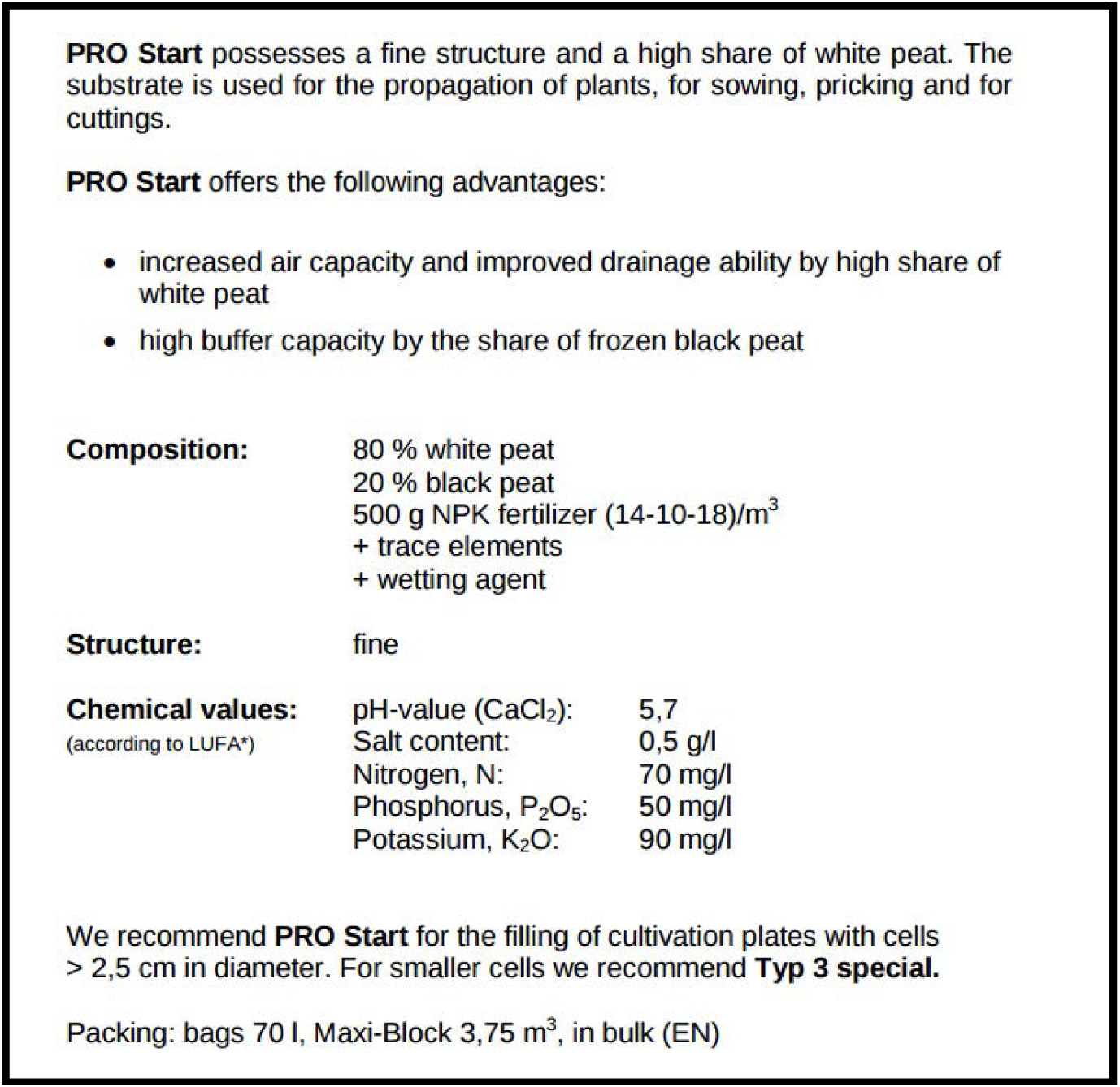
- A high-resolution **digital camera** (>12 megapixels). A digital SLR camera capable of capturing ‘RAW’ images is ideal.
- Materials to facilitate random flower head sampling (see cartoon):

- When there are more than 100 flowering individuals in a plot: **Two measuring sticks**. The first stick can be placed in one side of the plot. The second can be placed perpendicularly and be moved 10 times (every 8 cm). Then the experimenter can sample 10 times along the second stick (every 8 cm) the closest plant. This procedure can be used to sample flower heads in a grid fashion 100 times.
- When there are less than 100 flowering individuals in a plot: A 1 m bright-coloured **rope** can be used instead. By moving this rope along the plot, one can keep track of sampled individuals. 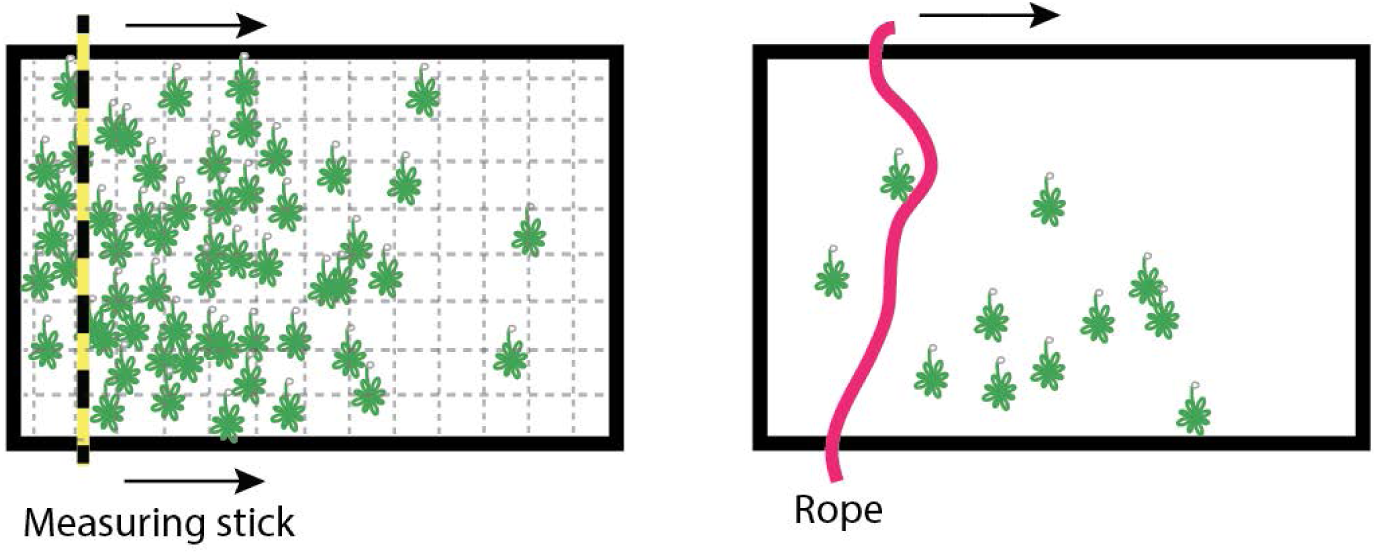

#### Optional

- iButton reader to read out iButton temperature and air humidity data and to reset iButton for further data logging (discuss this with coordinators).

### 1.3 Setting the experiment

For setting up the experimental plots:

1. **Choose a lawn, ruderal or open area** which can be used for at least three consecutive years and preferably that is not accessible to the public (e.g. an institution’s ground, for security reasons). If there is a risk of disturbance from animals, it would be ideal to fence the area. Make sure that local law does not conflict with any of the procedures of this experiment. And make sure that risk of contamination of *Arabidopsis thaliana* from native populations to experimental plots and *vice versa* is negligible.
2. For all locations, the **initial experimental setup** will take place between September and November 2017. All experiments will start the same year, but participants are asked to discuss the **optimal timing** of key stages and activities of the experiment (sowing, surveying, harvesting, etc.) with the coordinators.
3. Establish the **12 plots** using the plastic 60×40 cm trays.

- Choose 12 locations for the plots. These can be in a grid or loosely arranged within the area. Each plot must be spatially separated from any other plot by a minimum of half a meter and a maximum of ca. 5 meters.
- Remove all vegetation in each of the plots, dig out ca. 5-10 cm depth of soil and place the tray in the digged square. Then, fill the tray with compost soil (16 L per tray). Trays must be prepared in time before a rainy period to ensure good germination (see above the decision rules regarding the expected optimal timing for sowing depending on the location and participant expertise). The first sowing should take place on the same day of establishment of the plots.
- Bury the 2 iButtons with temperature logger in 2 separate plots which are well distributed among all plots (e.g. the third and sixth if the plots are arranged in a single line). Bury them at 5 cm depth in the middle of the plot (at least 10 cm away from tray edges). Make a note in which plots the loggers are buried. In addition, install the wooden construction (iButton with Air humidity logger) by sticking it at 10 cm depth at a random place located roughly centrally among all plots.
- Sketch the positions of the plots and note their ID (written and carved on the plastic trays) on a map (e.g. a local map or a Google earth orthophoto) and send it to the coordinators. Also note the day and exact time of the first sowing.

Below are some photos of an earlier pilot study using plots of 50 x 50 cm with natural soil and PVC tubes:

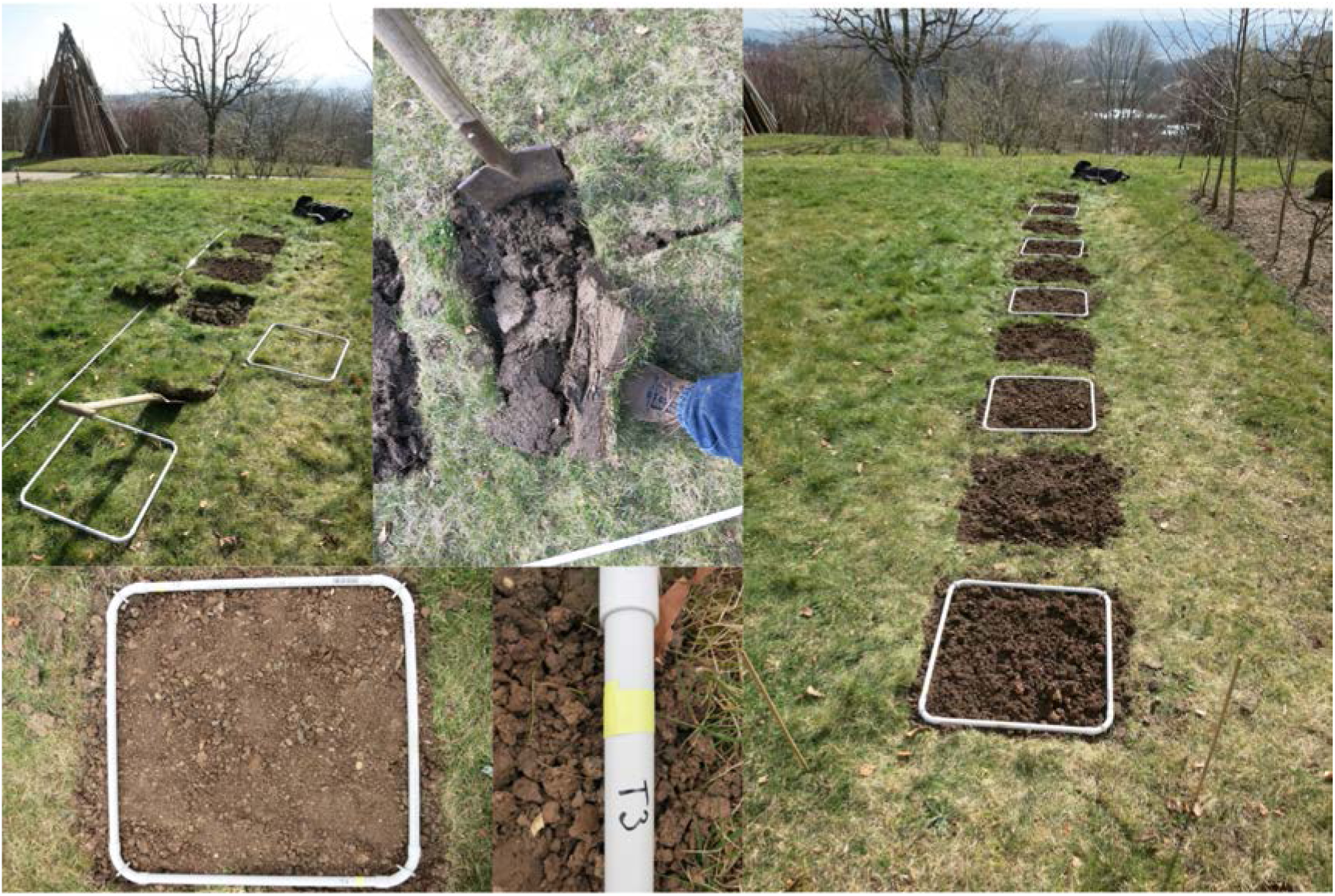

1. If deer or rabbit **herbivory** is likely, set up appropriate exclosures in the perimeter. Do not exclude insect herbivores with cloth mats.
2. Sample ca. **50 mL of the commercial soil** used for the experiment. Store these samples in a freezer (any temperature below zero) until shipment. Write “SOIL”, the site code and the date on the falcon tube (e.g. SOIL-32-20171024). Write this in the falcon tube, the paper notebook, and in the online spreadsheet (**Dataset S1**). This soil will be used for soil nutrient analysis.
3. **Sowing** will take place in each of the plots in Autumn 2017 during three separate sowing events with two weeks in between. Optimal timing for sowing has to be defined by the participant for his/her own site (fill the following table :https://docs.google.com/spreadsheets/d/16YBmFWdxQP8Fy1cFRjxz3X-BseZzAvOlFrVIVsu_bGM/edit?usp=sharing), and it will be discussed with coordinators prior to starting the experiment. Ideally, the optimal timing must be a relatively cold (but not freezing/snowing), humid period between September (for northern sites) and November (for southern sites). Importantly, there is **sowing only in the first year** not in consecutive years, unless complete failure occurs in the first year.
4. Register the exact location of the experimental garden in latitude and longitude <u>WGS84</u> **coordinates**.
5. Find the **closest weather station** and request temperature and precipitation data from the last 5 years and later for every year that the experiment runs.

➡ **General note for participants:** Participants are allowed to pursue parallel experiments that address specific questions of their interest as long as they do not interfere with the success of the main GrENE-net experiment. For instance, doing further measurements such as density of plants within the plots, or do independent sequencing of tissues, etc. Nonetheless, we recommend to discuss such additional activities with the coordinators before starting the experiment.

➡ **Note for experiments at extreme environments:** if the participant thinks his/her site might be too harsh for proper seed germination (*e.g.* very dry, desert environments), it is allowed to artificially water the plants during the sowing period. However, such artificial watering must be exceptional and restricted to extreme sites, as the goal of the experiment is to investigate adaptation to various climatic conditions with limited human interferences. Please discuss with coordinators the planned watering events.

➡ **Note regarding precautions and measures to prevent the spread of the species:** all participants must check local rules regarding the use of biological material in their location. We suggest to spread herbicide at the end of the experiment to avoid the spread of the species and autoclave the used soil when possible. Please read the ethics policy statement (grenenet.wordpress.com/policy-and-ethics-statement/).

### 1.3 Surveys of plants and census

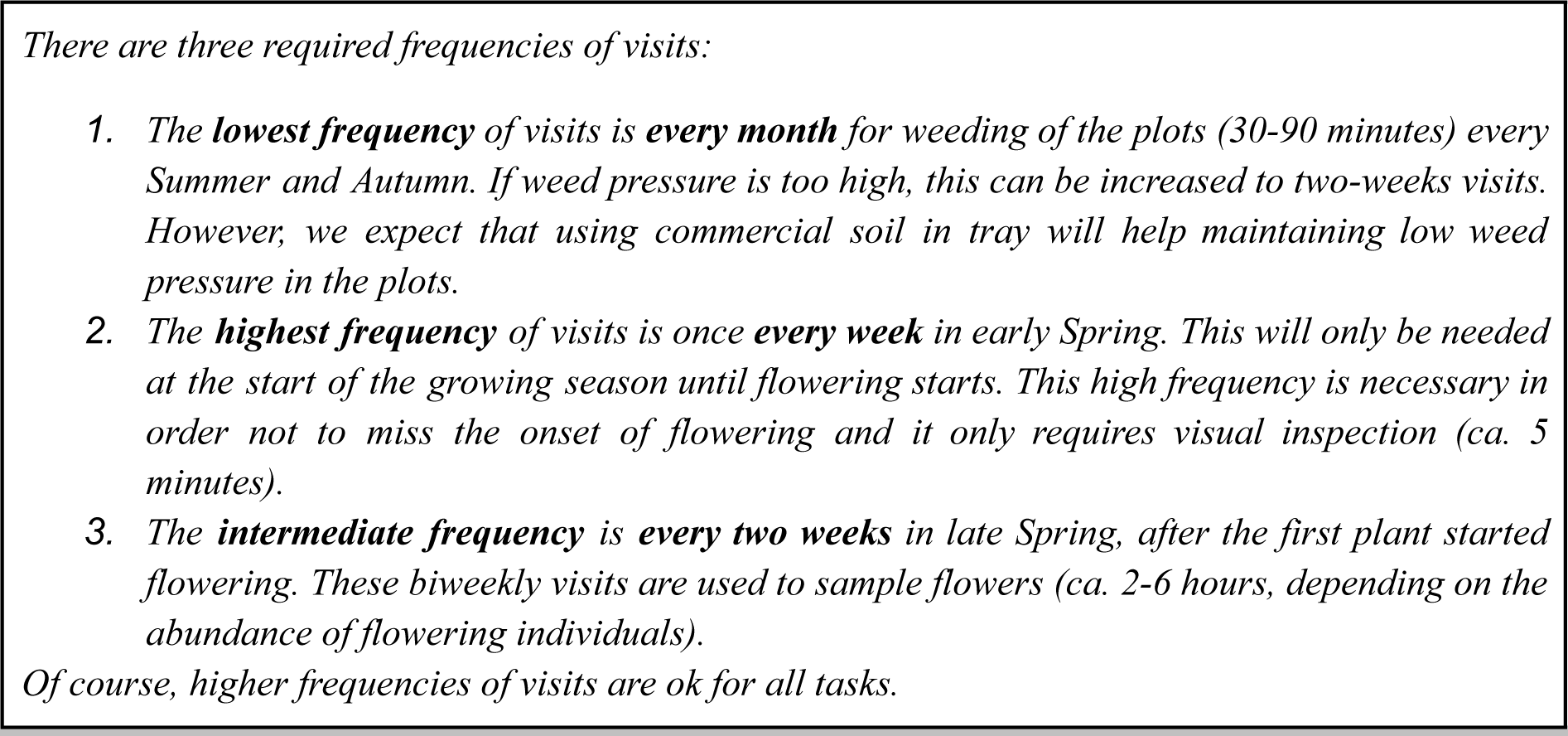

1. Keep a **notebook** to record observations and your activities throughout the experiment. It is notably important to record the timing of germination at the beginning of the experiment and every year (as good as the bi-monthly visits allow such an assessment).
2. Upon each visit to the plots, check if the plot IDs are still readable. If not, use an edding marker to improve readability. If you spot any flowering Arabidopsis individual out of the expected period, e.g. in Autumn, proceed to flower heads sampling right away as in Spring (see next section about material collection).
3. During each monthly visit, perform careful **weeding** of all non-*Arabidopsis thaliana* plants in the plots. Take pictures before and after weeding to estimate weed pressure in the plots. It is also recommended to include quantitative information about weed pressure in the notebook. **Important exception**: no weeding during vegetative growth nor flowering to not disturb Arabidopsis plants.
4. As soon as environmental conditions become more favourable for plant growth (after snow melt or after temperatures increase), **check for flowering** on a weekly basis.
5. When you see the **first flowering individual**:

a. In case there are many flowering individuals, start harvesting right away (see section IV below).
b. In case there are only few flowering individuals per plot (<<5% of all rosettes), start harvesting the following week.
c. After the first flower sampling effort, return every 2 weeks for flower sampling until flowering ceased or is negligible. In general, the flowering period ranges between 1-2 months (rarely 3 months) depending on the geographic location. We aim to sample a minimum of 3 times and a maximum of 5 times which include a sample during peak flowering (more samplings are, of course, possible but we might pool the flowers from two consecutive samplings).
d. During each flower sampling effort, take <u>photographs of all plots</u>. Make sure that the entire plot is visible and as large as possible in the picture, that all <u>4 sides are parallel in the frame</u> (zenithal imaging) and that the plot ID is readable.
6. After the first year, remove the three iButtons and read out the temperature and air humidity data to a laptop, reset the data loggers and replace the loggers in the same location. A detailed protocol for these steps will be provided later. If you do not have an iButton reading device, either we will provide iButton readers or iButtons should be replaced with new ones which we sent to you and the old iButtons should be sent back to us.

Population size will be estimated once per year when the first individual(s) started flowering. This timing is chosen to obtain measurements at a comparable timing among the sites. Although the timing will depend on the climate and will vary from year to year, we expect this to be done around February-March. Participants need to be attentive to not disturb emerging flowering buds during this estimate.

Participants will count the number of rosettes overlapping a string or measuring rule in a diagonal and an off-diagonal of the tray. In the example below, both the diagonal and the off-diagonal overlap with 2 rosettes. These two numbers will be written down in a paper notebook, and in the online spreadsheet, page census size (**Dataset S2**).

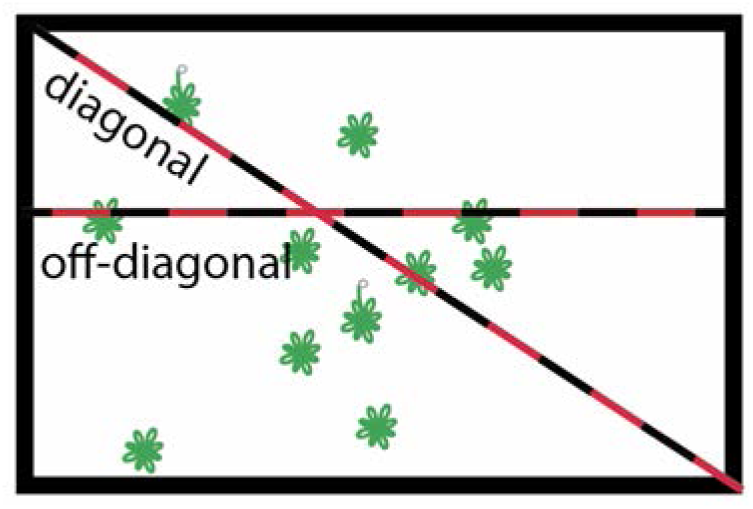

Although these estimates will provide valuable yet quick information of the density of plants, we can also use images of the trays to go back and potentially validate population density per tray.

### 1.4 Tissue collection

1. During each flower sampling event, if there are up to **100 flowering individuals and a minimum of 10**, pick **one open flower from each flowering individual** and add all flowers in one 2 mL Eppendorf tube per plot (see photo). Sampled flowers should be in anthesis, i.e. the white petals should be visible and the silique should not yet be visible. Except for very specific reasons that must be discussed with the coordinators, we do not aim to collect vegetative tissue but only flower heads. Mark each Eppendorf tube with the sample type, “FH” from flower heads, the site code, the plot ID, and the date of sampling (e.g. FH-54-1-20170524 indicates the flower heads collected in the site 54, the plot 1 and the date 2017 May 24). **Please also write the number of flower heads that have been collected.** All this information should be written not only on the tube, but also in a paper notebook, and in <u>the online spreadsheet</u>. 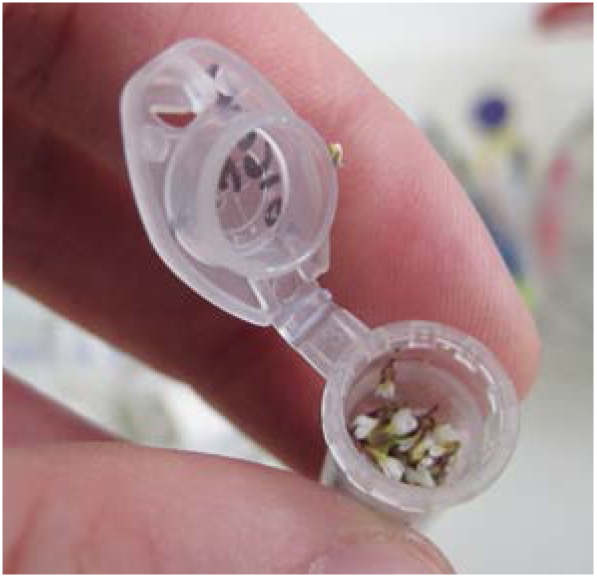
2. Exception: If a plant has only one flower, do not sample it, as it would affect its fitness too much. Note: We do not want sampling of flower heads to be proportional to the individual size, as this will entail a high complexity of the protocol (i.e. we aim for **only one flower head per individual** across the flowering season). Ideally one individual would be sampled only once, but for individuals that are very large and their flowering period spans more than two weeks, it might happen that they are sampled two times. In order to avoid this, it would be required to monitor each plant and add a mark when a plant has been sampled, but again the high complexity of this protocol would be unfeasible. Thus, it is **not necessary to mark plant individuals**.
3. In case there are more than 100 flowering individuals, collect evenly within the plot and stop collecting when you have sampled *ca*. 100 flower heads (max. 150). When using two measuring sticks or a rope (see “Required material” in Section I).
4. After sampling of flower heads, put the **Eppendorf tubes into a −80 °C freezer** for storage until shipment. If no −80 °C freezer is available, please keep the material at max 4 °C and send it as soon as possible to the coordinators (ideally during the same day).
5. **Every summer**, when plants have ripened, dispersed their seeds, and died, collect ca. **50 mL of soil per plot**. Each sample should combine 9 soil subsamples. Take one spoon of soil including the potentially seed-rich surface but simultaneously reaching ca. 2 cm deep, ca. 5 mL, from 9 locations in the tray with a minimum distance of 10 cm from the edge (see cartoon). Store these samples in a freezer (any temperature below zero) until shipment. Write “SB” (seed bank), the site code, plot ID and the date on the falcon tube (e.g. SB-32-1-20170524). Write this on the falcon tube, in a paper notebook, and in <u>the online Spreadsheet</u>. **Note:** there will be one sample for soil analysis before starting the experiment (at Setup), and then samples of seed bank per plot every summer. 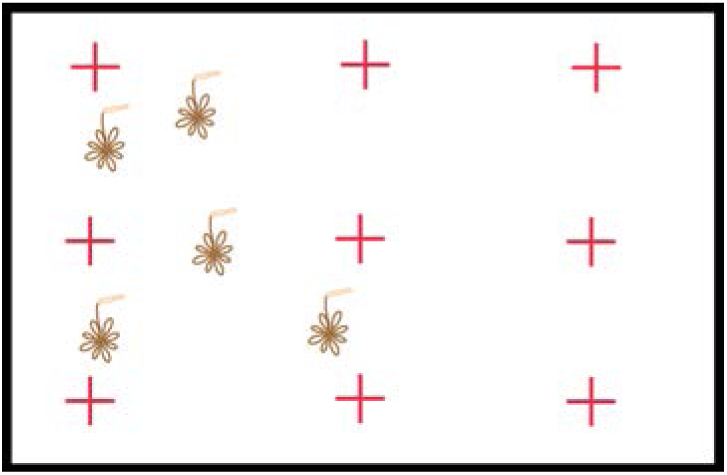

In summary, for every location, we expect the following material to be sent to the coordinators in summer:

- 1 Falcon tube with soil at the experiment setup (first year).
- 12 x 1 Falcon tube with soil for the seed bank analyses.
- 12 x 3-5 Eppendorf tubes with flower heads.

### 1.5 Shipment of samples

Every year after flower sampling and soil sampling, all material should be sent to the GrENE-net coordinators for analyses. Put all tubes for each single date together in a small plastic bag. Put all materials in a styropor box with dry ice. Close the styropor box properly wrapped. Communicate with the coordinators for the shipment. Participants from Europe sent the materials to Niek Scheepens who assembled them for a large shipment with dry ice to Moi Exposito-Alonso.

### 1.6 Policy on sample mishaps

As in any field experiment, unpredictable accidents might happen. These include pests of herbivores or pathogens, floods and droughts, human interventions, etc. In all these cases, please contact the GrENE-net coordinators to discuss potential solutions.

- Generally, climatic disasters are part of the aim of GrENE-net to characterize climate-driven natural selection. Only in cases in which these happen due to flawed experimental setup (e.g. experiment is done in a artificially compacted soil), we would try to amend it (e.g. designing a draining system).
- If there are clear signs of people interfering with the experiments, we can consider moving the experiment to a more suitable location not accessible to the public.
- If herbivores are mammals or big animals, setup an exclusion net. If they are insects or other small herbivores, contact the GrENE-net coordinators to discuss the application of a pesticide.
- Likewise for pathogen pests. Contact the GrENE-net coordinators to discuss the application of a pesticide. Note that we might want to collect samples before any spraying.

### 1.7 Example sampling timeline

This timeline is an example of likely timings in temperate areas of Europe. Precise timings will be discussed with each participant.

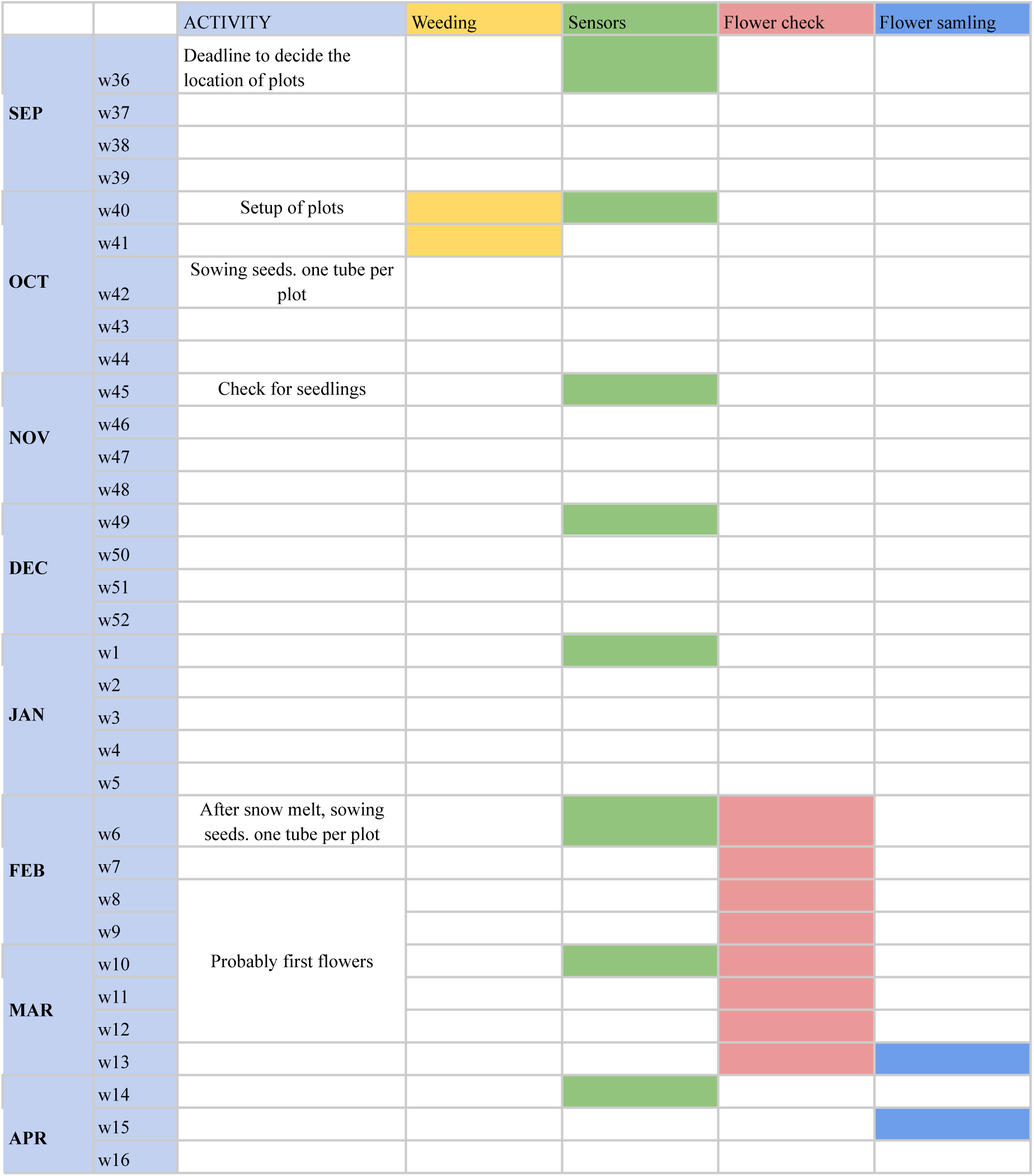
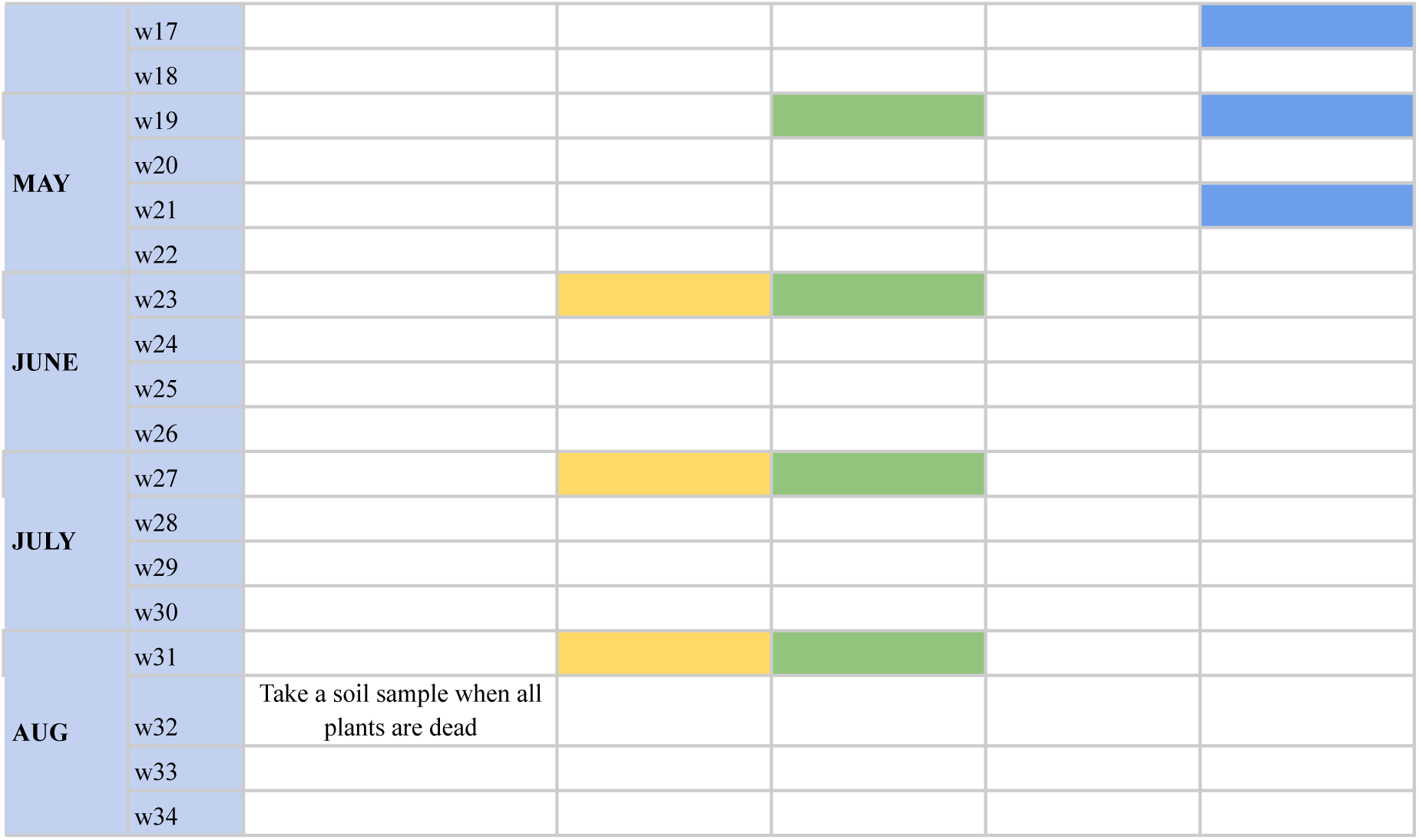

In fall 2017, packages containing materials and seeds of the cosmopolitan *Arabidopsis thaliana* were sent to experimental facilities across Eurasia and North America to set up a Globally Distributed Evolution Experiment, investigating the Genomics of rapid Evolution to Novel Environments (GrENE-net.org). The aim of this experiment was to expose a set of wild lines of *A. thaliana* to replicated semi-natural conditions and observe genetic evolution as it happens. The experiment has now been running for the minimum agreed term of two annual cycles. In this report, we show a detailed analysis of the overall success of the first phase, experimental setup and population survival in the 2017/18 and 2018/19 seasons, describe the results of a sequencing proof-of-concept and bioinformatic pipelines that will be used in this second laboratory phase of the project, and propose a future strategy for this unique collaborative experiment. In the years 2018 and 2019, the GrENE-net participants collected flower and soil samples, monitored individuals and the environment, and sent the materials to the GrENE-net coordinators. Fig. 1 summarizes the efforts so far (NB: subject to change upon receiving materials from a fraction of participants; send an email to if any irregularity is detected).

### Text S2 Sample and census data cleaning, curating, and modeling

We cleaned and homogenized participant-provided spreadsheets containing census measurements of overwintering populations (**Dataset S1**) by correcting formatting and typos, and cross-referenced participant diaries to verify and correct missing entries. The census data included total, diagonal, and off-diagonal plant counts (see **Text S1**). Using this transect census data, we estimated the total number of plants for all trays. Some participants additionally conducted an additional census counting all plants within a tray, which we used as validation and confirmed robust predictability *R^2^*=0.564. Some participants did not collect any census data. In addition, the number of flowers collected for sequencing represents additional census information of reproductive adults during the flowering season, which we use to our advantage to impute population sizes across sites.

The final dataset includes information on ‘site’, ‘plot’, ‘date’, ‘diagonalplantnumber’, ‘offdiagonalplantnumber’, ‘totalplantnumber’, ‘meanfruitsperplant’, ‘sdfruitsperplant’, ‘comments’, ‘flowering_plants’, ‘totalplantnumber_estimates’, and ‘totalplantnumber_complete’ (**Dataset S2**).

The steps to arrive at such a final dataset are as follows:

**A)** We processed spreadsheets detailing flower collection, corrected typos, and resolved inconsistencies by reviewing participant diaries. The resulting dataset includes columns like ‘number_flowers_collected’, ‘site’, ‘plot’, ‘date’, ‘sampleid’, ‘year’, ‘month’, ‘day’, ‘to_skip’, ‘source’, and ‘generation’ (**Dataset S3**). (Cleaning script <u>census_clean_step1.ipynb, samples_clean_step2.ipynb</u>).
**B)** Because the total number of flowers in a population tray sometimes exceeded 100 flowers, participants were instructed to limit it to 100 flowers. In those cases, this would not represent the total census. Some participants inadvertently collected more than 100 flowers, which we ended up using to our advantage to adjust the expected total number of flowering adults on sites where collections were capped at 100 by interpolating based on longitude, latitude, and altitude from the sites that did not follow the limit (Cleaning script Solving_cap_flowers_col_step3.ipynb)
**C)** To fill in the blanks of sites without total census sizes, we finally used the flower count data, we estimated population sizes with predictors including ‘generation’, ‘longitude’, ‘latitude’, ‘altitude’, ‘max_flower’, and ‘flowerscollected_corrected’. We trained a RandomForestRegressor on these variables, resulting in an *R^2^=*0.87 when compared to sites with measured population sizes. The comprehensive final dataset contains columns: ‘site’, ‘plot’, ‘generation’, ‘max_flower’, ‘min_flower’, ‘flowerscollected_corrected’, ‘totalplantnumber_complete’, ‘longitude’, ‘latitude’, ‘altitude’, and ‘predicted’ (**Dataset S4**). (Cleaning script <u>estimate_pop_size_step4.ipynb</u>)

### Text S3. g_r_enepipe and g_r_enedalf: Bioinformatics tools for large-scale Pool-seq analyses

The unprecedented size of the GrENE-net experiment necessitated novel software development, as existing tools did not scale to the amount of data produced. In particular, we needed (a) an automated workflow for processing pool-sequenced samples, and (b) scalable software to conduct population genetic analyses on Pool-seq data.

First, we developed g_r_enepipe (*20*), a variant and frequency calling pipeline. The main workflow of g_r_enepipe can be used both for individuals and for pooled populations. Starting from raw sequencing reads, the initial steps are identical in both use cases: read trimming and mapping to a reference genome, as well as a plethora of quality control tools. Then, when using g_r_enepipe for samples of individuals, variant calling is conducted; the overall process for this follows the GATK Best Practices (*107*). When using g_r_enepipe for pooled samples, we also offer to run a HAF-pipe module to compute allele frequencies if individual whole-genome data is available of founders mixed prior to Pool-seq samples (*21*, *82*) (we updated HAF-pipe for a Python-based and upgraded hapFIRE, **Text S3**). In all steps, users can choose from a variety of established tools, and optional downstream tools are offered, for instance for ancient DNA, or for annotation of variants. The backend of g_r_enepipe is the Snakemake workflow management system (*108*, *109*), which takes care of background work necessary to run an analysis on thousands of samples. This includes installing all needed dependencies (i.e., the bioinformatics tools that are run within the workflow), the file bookkeeping, job submission to computer clusters, and more.

Second, we developed g_r_enedalf (*22*), a population genomics C++ tool for fast pool-sequencing data handling and population genomic statistics. The Pool-seq approach introduces two nested levels of sampling noise (*100*): First, individuals are sampled from a population into a pool, and second, reads are sampled from the pool when sequencing. When computing estimators of typical population genetic statistics, the biases introduced by this noise need to be accounted for. Existing tools for this such as PoPoolation (*110*, *111*) offer these corrections for computing nucleotide diversity (Theta Pi, Theta Watterson, Tajima’s D) as well as population differentiation (*F*_ST_). In g_r_enedalf, we have re-derived the statistics estimators, and found several shortcomings of the existing estimator and implementation of Tajima’s D in PoPoolation, as well as fixed a remaining bias of the Pool-seq estimator of *F*_ST_ in PoPoolation2. Furthermore, existing tools did by far not scale to our dataset sizes in GrENE-net, and were lacking needed functionality. The implementation of g_r_enedalf in C++ is several orders of magnitude (×100 times) faster than existing tools, and has multi-threading support for large datasets. It furthermore offers considerably more options, such as different input file formats, region and numerical filters, masking options, different windowing modes, different strategies for computing per-window averages, and more. In the future, we also plan to extend the functionality to other typical estimators such as the F- and G-statistics and genome-wide associations with metadata.

Both tools were instrumental for the basic processing of the data in this study. We used g_r_enepipe for trimming the reads and mapping the samples to the *Arabidopsis* reference genome, and subsequently used g_r_enedalf to extract allele frequencies from the mapped data, as well as compute diversity and differentiation as explained in the main text (e.g. **Fig. S19**).

### Text S4. Detailed description of HapFIRE for constructing accession and allele frequencies

A popular approach to characterize allele frequencies in Pool-seq Evolve & Resequence approaches where genomic data of the founder accessions or strains is known is a haplotype-based frequency reconstruction using HAF-pipe & HARP (**Text S3**). The previous software has several shortcomings that we solved with HapFIRE (https://github.com/moiexpositoalonsolab/HapFIRE). The main goal is the same: estimates SNP and founder frequencies from pool sequencing data given a reference panel of the founders, with the assumption of no recombination among reference haplotypes. Violation of this assumption will have a stronger impact on frequency estimation of founder individuals than SNPs. HapFIRE takes a phased VCF file as the reference panel, from which it estimates the frequencies of SNP and founder individuals based on aligned pool sequencing reads.

#### Improved allele frequency estimates from Pool-seq

HapFIRE first uses the phased VCF file to calculate the independent linkage disequilibrium blocks based on *r^2^*. Within each independent block, hapFIRE enumerates all unique haplotypes and relies on HARP (*82*) to calculate the maximum likelihood estimate of haplotype frequencies from the observed pooled read samples. The SNP frequencies are calculated with the following formula:

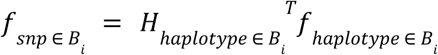

Where 𝑓_𝑠𝑛𝑝 ∈ *B*𝑖_ is the frequency of 𝑚 SNPs in the independent block *B*𝑖, 𝐻_ℎ𝑎𝑝𝑙𝑜𝑡𝑦𝑝𝑒 ∈ *B*_𝑖 is the indicator matrix {0, 1} of 𝑛 unique haplotypes in block *B*_𝑖_. 𝐻_ℎ𝑎𝑝𝑙𝑜𝑡𝑦𝑝𝑒 ∈ *B*_𝑖 has a dimension of 𝑛 × 𝑚. Each row of 𝐻_ℎ𝑎𝑝𝑙𝑜𝑡𝑦𝑝𝑒 ∈ *B*_𝑖 represents a unique haplotype which is a vector of zeros and ones, and each item in the vector represents the presence or absence of the SNP in the haplotype. 𝑓_ℎ𝑎𝑝𝑙𝑜𝑡𝑦𝑝𝑒 ∈ *B*_𝑖 is the frequency of unique haplotypes estimated by harp. The frequency of SNPs in the block is the product of matrix multiplication between 𝐻_ℎ𝑎𝑝𝑙𝑜𝑡𝑦𝑝𝑒 ∈ *B*_𝑖^𝑇^ and 𝑓_ℎ𝑎𝑝𝑙𝑜𝑡𝑦𝑝𝑒 ∈ *B*_𝑖.

#### Reconstruction of relative proportion of founder genotypes

In cases where Evolve & Resequence experiments are conducted starting from known strains or accessions, it may be helpful to reconstruct the theoretical relative proportion of founders (this may be most useful in species that are partially or predominantly selfing). The founder individual frequencies are calculated by solving the following equation:

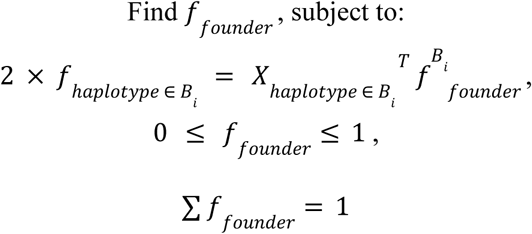

where 𝑓*B*𝑖_𝑓𝑜𝑢𝑛𝑑𝑒𝑟_ is the frequency of 𝑝 founder individuals based on the information of block *B*_𝑖_, 𝑋_ℎ𝑎𝑝𝑙𝑜𝑡𝑦𝑝𝑒 ∈ *B*_𝑖 is the dosage matrix {0,1,2}with a dimension of 𝑝 × 𝑛 recording the dosage of unique haplotypes each founder contains. Each row of 𝑋_ℎ𝑎𝑝𝑙𝑜𝑡𝑦𝑝𝑒 ∈ *B*_𝑖 represents a founder individual and each column represents a unique haplotype. 𝑋_𝑎𝑏_ represents the dosage of haplotype 𝑏 in founder 𝑎. The row sum of 𝑋_ℎ𝑎𝑝𝑙𝑜𝑡𝑦𝑝𝑒 ∈ *B*_𝑖 equals to the ploidy of the organism, which is 2 in the case of *Arabidopsis thaliana*. HapFIRE uses the CVXPY package (https://www.cvxpy.org/) to solve the optimization problem. Each block will output an estimate of founder individual frequencies, 𝑓_*B*_𝑖_𝑓𝑜𝑢𝑛𝑑𝑒𝑟_. In our benchmarking, we noticed that the accuracy of 𝑓^*B*^_𝑖𝑓𝑜𝑢𝑛𝑑𝑒𝑟_ positively correlated with the number of unique haplotypes in block *B*_𝑖_, and will reach a plateau when the ratio between unique haplotypes and founder individuals is above 0.9. This suggests that the model cannot accurately estimate the founder frequencies when only a few unique haplotypes are present in the block, or regions where many founder individuals share the same haplotypes. To address this problem, HapFIRE only uses founder frequencies estimations from blocks with high numbers of unique haplotypes (> 90% quantile in number of unique haplotypes), and averages the estimates across these blocks to result in the final founder individual frequency estimation, which we show is highly accurate with simulations (**Fig. S13**).

### Text S5. Accuracy of the allele and accession frequency reconstruction from pool sequencing for founder and evolved populations

For each founder and evolved pool-sequenced population, we reconstructed ∼3.2 million SNP allele frequencies and the relative frequencies of 231 founder accessions. Allele and accession frequency reconstructions were conducted using hapFIRE, which leverages known relationships among accessions, haplotypes, and SNPs (see Methods and Text S4).

We assessed hapFIRE’s accuracy and benchmarked it against the existing tool HAF-pipe through extensive in silico simulations. We created pooled samples containing between 2 and 150 accessions and simulated sequencing depths of 1× and 10×. Performance was evaluated using predictive R² and root-mean-squared error (RMSE).

In allele-frequency reconstruction, hapFIRE consistently outperformed HAF-pipe across all pooling and sequencing-depth scenarios. For example, with 150 accessions pooled at 10× coverage, hapFIRE achieved an average RMSE of 0.00799 (IQR: 0.00783–0.00823) and predictive R² of 0.9994 (IQR: 0.9991–0.9996). Under the same conditions, HAF-pipe yielded an RMSE of 0.0184 (IQR: 0.0181–0.0187) and R² of 0.9930 (IQR: 0.9928–0.9931) (**Fig. S13**). Increasing sequencing depth from 1× to 10× provided only marginal gains in accuracy, in line with Tilk et al (*21*).

In the accession frequency reconstruction, we only evaluated the performance of hapFIRE because HAF-pipe does not support accession frequency reconstruction. The accuracy of accession frequency reconstruction decreased as the number of pooled accessions increased. The highest accuracy was achieve with two accessions at 10x sequencing depth, yielding R2 of 0.965 (IQR: 0.956-0.993) and normalized RMSE (RMSE scaled by standard deviation of the group truth) of 0.179 (IQR: 0.0923-0.231). In the simulation with 50 pooled accessions with 10x sequencing depth, hapFIRE maintained robust performance with R2 of 0.828 (IQR: 0.810-0.849) and normalized RMSE of 0.610 (IQR: 0.604-0.616), suggesting on average predictions deviate from true values by 0.6 standard deviation of the true frequencies.

The starting frequency of founder accessions was inferred from the sequencing of founder seeds sent to GrENE-net participants. We took eight samples of seeds and individually sequenced each sample. We then used hapFIRE to infer the founder accession frequencies from the sequencing data. We found the average Pearson’s correlations among eight samples to be 0.523 (IQR: 0.427 - 0.627). The final starting frequency for each founder accession was the average value across eight samples (Dataset S5). We then calculated two metrics to see the deviation of the reconstructed accession frequencies from the expected uniform value of 1/231 for each accession (Rooted mean squared error: 0.0014, Mean absolute error: 0.0009).

### Text S6. Quantifying seed dispersal between independent trays in GrENE-net

In site 28, a fallow old-field site on Duke Forest property, Durham, NC (36.009169, −79.018746), 12 GrENE-net trays were established and placed ∼2.35-m apart. To investigate seed dispersal in these experimental plots, we added empty trays containing the same soil as the source populations (Sunshine Mix #1, SunGro Horticulture), placed at three distances, 0.29-m 0.02 (close trays), 0.97-m 0.04 (middle trays), and 1.67-m 0.08 (far trays), from the experimental source populations to catch dispersed seeds (Fig. 1). Empirical evidence suggests that wind-dispersed plants are characterized by mostly short dispersal events so this short distance should adequately capture natural dispersal dynamics and provide sufficient sample sizes for the comparisons across distance (*112*). There were 4 blocks in the field, and each block consisted of 3 source trays, 4 close trays, 2 middle trays, and 1 far tray, as shown in the figure below.

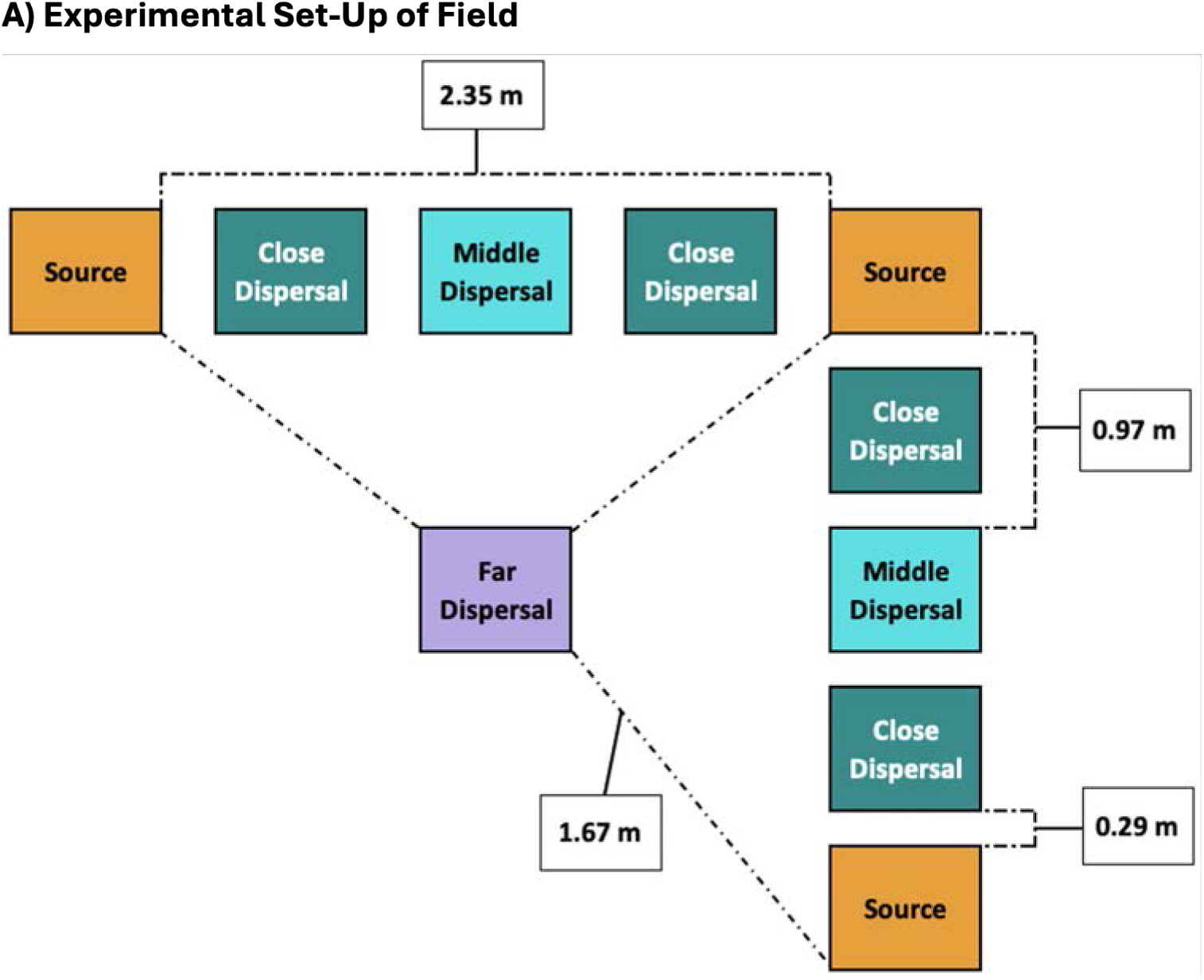

After the first dispersal season, the dispersal trays were brought into the greenhouse and grown under optimal germination conditions to recover plants from the seeds that were dispersed into them. Given the large number of trays and limited space in the greenhouse, the dispersal trays were assessed in batches (Batch 1 = blocks 1 and 2; Batch 2 = blocks 2 and 4). First, to account for any movement of seeds during transportation and to try to maximize *A. thaliana* germination, we transplanted the soil from different levels from the field dispersal trays into new trays. we took two samples from the top layer of soil (samples T1 and T2) and one sample from the bottom layer of soil (sample B). Therefore, for every field dispersal tray, there were three greenhouse dispersal trays. Next, to break any dormancy, we moved the greenhouse dispersal trays into a dark cold room set at 4°C for stratification. After 2 or 6 days in stratification for batch 1 and 2, respectively, we moved the trays to the greenhouse in warm light conditions (16 h light at 22°C and 8 h dark at 16°C) and monitored them for germination. At the 4-leaf stage, a sample of individuals were transplanted into plug trays, so they would be grown at a constant density. Due to large sample sizes and COVID shutdowns, we was not able to transplant all individuals, so some individuals remained in their greenhouse dispersal trays.

To estimate how many seeds dispersed across distance (as continuous), directly from the data, we took into account the proportion of the total area at each distance from the source trays that was covered by each dispersal tray. We also assumed that 100% of the seedlings observed in the “close” dispersal trays came from the nearest source tray, and that the proportion of seeds that arrived at the “middle” and “far” dispersal trays from each of the 2 or 3 nearest source trays, respectively, depended on the distance between each of those sources and the dispersal tray. For each pairing of dispersal tray and corresponding source tray(s), estimated dispersal at the distance of that dispersal tray (from the source(s)) was calculated by dividing the number of *A. thaliana* in the dispersal tray, that likely came from that source tray, by the proportion of the total area covered by that dispersal tray at that distance (**Fig. S3**).

### Text S7. Expectation of allele frequency changes under neutral evolution or accession sorting

The neutral evolution expectation from a classic Wright-Fisher (WF) population (i.e. a population of diploid individuals under random mating, discrete generations, linkage equilibrium, constant size, no selection, no mutation, and no migration) is a well-known (*113*) magnitude of allele squared frequency changes (i.e. Var(Δp)) between two generations (t_0_→t_1_) depends on the starting frequency and the population size: *p_0_(1-p_0_)/2N*.

Because it is typically expected that a mixture of *Arabidopsis* accessions would behave as a sampling of inbred lines–sometimes called accession sorting (but see **Text S10**)–that may differ from the theoretical WF haploid model, we also conducted simulations where for any *N* flower sample of GRENE-net (333 data points of site by replicate in the first year with at least 3 flowers) we sampled *N* out of 231 accessions with replacement. The sampling of 231 accessions was either conducted uniformly (i.e. equal probability of all accessions) or following a Poisson (λ=7; following an average from seed counts in similar common gardens. Analyses are very insensitive to either uniform or Poisson samplings of several lambdas). Then for each GrENE-net data point we produced a neutral expectation *E[ (p_t_ - p_t-1_)^2^]*. Because we know the expected variance in allele frequency depends strongly on starting frequency and population size, we calculate this expectation for ±0.5% bins and ±5 flower bins. We used the 13,985 LD-pruned SNPs.

By comparing to the founder population, the mean of the genome-wide average frequence change in neutral-simulated populations in all three generations were not significantly different from zero (generation 1: mean E*(p_1_ - p_0_)* = 0.001, SD[E*(p_1_ - p_0_)*] = 0.008, one-sample t-test: adjusted *P* = 0.07707, N = 500; generation 2: mean E*(p_2_ - p_0_)* = 0.001, SD[E*(p_2_ - p_0_)*] = 0.016, one-sample t-test: adjusted *P* = 0.135, N = 500; generation 3: mean E*(p_3_ - p_0_)* = 0.001, SD[E*(p_3_ - p_0_)*] = 0.021, one-sample t-test:adjusted *P* = 0.180, Fig S14). Similar to the mean of genome wide average frequency change, we also calculated the variance of genome-wide allele frequency change over three generations in our neutral-simulated populations (mean Var*(p_1_ - p_0_)* = 0.015, SD[Var*(p_1_ - p_0_)*] = 0.035; mean Var*(p_2_ - p_0_)* = 0.021, SD[Var*(p_2_ - p_0_)*] = 0.043; mean Var*(p_3_ - p_0_)* = 0.025, SD[Var*(p_3_ - p_0_)*] = 0.049). We then compared these two neutral evolutionary expectations, WF and non-WF, with the GrENE-net dataset calculating the average *(p_t_ - p_0_)^2^* for the starting allele frequency classes and population sizes (**Fig. S15, S16, S17**).

### Text. S8. Evidence of natural selection and adaptation in GrENE-net

To test whether the observed magnitude of evolution is beyond neutral genetic drift, we compared the mean and the variance of changes in allele frequencies with neutral simulations and theoretical expectations using populations that survived all three generations (N=142). Genome-wide frequency changes were not significantly departed from expectations from founder to first survivor generations (Two sample t-test, t_0_→t_1_, *P=*0.223), but became significant in the second and third generations (Two sample t-test t_0_→t_2_, *P=*0.001, t_0_→t_3_, *P=*9.518×10^−5^, **Fig. 2A, Fig. S14**). We then evaluated the observed variance of allele frequency changes and compared it to the neutral simulation. We observed magnitude of evolution, measured as variance in allele frequency change, was significantly beyond neutral genetic drift expectations (t_0_→t_1_:Var_observed_/Var_neutral_ = 4.155 [95% CI: 2.576-5.734], t_0_→t_2_:Var_observed_/Var_neutral_= 9.131 [95%CI: 7.167-11.095], t_0_→t_3_:Var_observed_/Var_neutral_= 8.398 [95%CI: 6.584-10.211]; two sample Kolmogorov-Smirnov test, t_0_→t_1_, *P=*2.794×10^−16^, t_0_→t_2_, *P<*2.2×10^−16^, t_0_→t_3_, *P<2.2*×10^−16^). Both the mean and variance of the allele frequency change showed significant deviation from the neutral expectation, therefore, suggesting natural selection and adaptation processes happened in GrENE-net.

In addition to allele frequency changes departing from neutral simulation, we also observed adaptive evolutionary rescue in GrENE-net. Eight out of 30 experimental gardens showed significant signs of population recovery in the third generation with U-shape trajectories indicating evolutionary rescue (**Fig. 2F**, **Fig. S9**, across sites: *N (t) ∼ a +bt+ct^2^*; *c*=37 *P*=3.03×10^−5^, *n*=670; example garden #4 Cadiz, Spain, *c*=130.71, *P*=0.0033, *n*=36, **Fig. S17** for all estimates). This also supports the notion that natural selection driven adaptation occurred in GrENE-net.

### Text S9. Explanation for repeatability vs parallelism

From population genetics, we expect the frequency of an allele to change directionally if the relative fitness of the individuals carrying the allele is higher than the population average: *p_t_= p_t-1_(w/w_avg_)*. Assuming complete selfing and using the reconstructed relative abundance of the 231 founder accessions, we can make the same argument about accession frequencies: an accession’s relative frequency will increase if its fitness is over the average (nb. *A. thaliana* generally self-fertilizes although we find evidence of outcrossing in ∼10% of samples from sequencing, see **Text S10**).

It could be that seed dispersion among replicates or other statistical processes leads to non-independence (e.g. experimenter, temporal sampling) or inflated cross-replicate correlations therefore weakening the argument of repeated evolution. To address this, we compared the correlation of accession frequency changes pairwise among two replicates within one site vs pairwise of one replicate from one site and another replicate from another site that has a similar climate (**Fig. S29**). We conducted this with an example set of three warm location site gardens and cold location site gardens, and found that indeed there is significant evolutionary parallelism in accession frequency changes in two gardens of similar climates. To assess whether this parallelism is significant, we also conducted two randomization approaches, each repeated one thousand times: (1) randomizing the entire dataset by re-sampling accession frequency across all sites, and (2) nested randomization re-sampling accession frequency within a site and replicating. We show that evolutionary parallelism in similar environmental sites is significant. We also find that evolutionary repeatability (i.e. within a site) is higher than parallelism. The increased correlation within-site, however, cannot be readily attributed to artifacts or lack of independence due to contamination, as the replicates within one site share more environmental pressures than two sites hundreds of kilometers apart that roughly have a similar annual temperature.

### Text S10. Approximating outcrossing rate from pool-sequenced flower samples

Collaborators reported the number of flowers in each sample while the frequency of all 231 accessions within each sample was estimated using HapFIRE. An outcrossing event was counted when the number of flowers exceeded the number of accessions present, assuming each flower is a separate accession if there was more than 1 flower. For an accession to be considered present in a sample, it had to have a frequency at or above 10%.We also implemented several measures to ensure proper outcrossing estimation: 1) samples lacking any of the 231 accessions above a 10% frequency were excluded, as they likely represented contamination and 2) samples with fewer than 10 flowers underwent manual verification. 3) To assess the impact of sample size, we analyzed how outcrossing rates varied with the number of flowers per sample across the three years of the experiment (see **Fig. S14**).

For samples containing a small number of flowers, sometimes even 1 flower (n=186 samples) or 2 flowers (n = 144), it was straightforward to detect outcrossing events as relative proportions followed Mendelian segregation ratios (e.g. see F1 hybrids, Inbred Line x F1, F1 x F1 cases in **Fig. S14A-C**).

To avoid overestimation of outcrossing events, we employed a conservative approach to detect outcrossing. We focused on samples containing a single flower (n=186), ensuring that the presence of multiple accessions definitively indicated an outcrossing event. To account for pre-experiment outcrossing, we only considered samples as outcrossed if they were F1 hybrids, characterized by two accessions present at approximately 50/50 frequency within a single flower sample (see **Fig. S14**). Using this stringent metric, we calculated a lower bound estimate for the outcrossing rate across all years to be 2.65%. Breaking this down by year, 2020 had a minimum outcrossing rate of 4.22%, 2019: 1.2%, and 2018: 2.7%.

The proportion of outcrossing events detected declined with an increase in the number of flowers per tube (**Fig. S14**). Since we applied a 10% cut-off to classify an accession as present, increasing the number of flowers diminishes our ability to detect outcrossing events. This is because the accession relative frequency in a recently outcrossed individual is typically around 50/50, or in some cases an even lower frequency, such as that of a segregating F2 individual. Therefore, once the flower count exceeds 3 or 4, the accession frequencies of an outcrossed individual drops below 10%, our detectable level (**Fig. S14A**). In addition, outcrossing events were detected in the first generation (2018), indicating that accessions had outcrossed during the seed bulking stage. The highest outcrossing rates, regardless of the number of flowers per sample, were observed in the third year (2020) of the experiment.

### Text S11. Per-garden genome-wide natural selection scans

To identify genomic region shifts across 30 experimental gardens we performed a SNP-based LRT-1 test for each garden. Next, we assigned significant SNPs (after Bonferroni correction) to haplotype blocks and conducted permutation tests to identify over-represented haplotype blocks across the 30 gardens. Finally, we extracted genes within the significant blocks and performed gene ontology (GO) enrichment analysis to characterize the biological functions of genes under selection.

Haplotype blocks were defined based on linkage disequilibrium (LD) patterns among the 231 GrENE-net founder accessions, using a genome-wide dataset of 3.2 million SNPs (*88*). In total, we identified 16,917 haplotype blocks, with an average block size of 6,963 base pairs. For each experimental garden, we projected the SNP-based LRT-1 results onto the haplotype blocks, assigning the lowest SNP *P*-value to its corresponding block. After completing this projection for all 30 gardens, we applied Bonferroni correction to the *P*-values and counted the number of gardens in which each block exhibited a parallel shift. This approach did not assume the same direction of shift across gardens, as *P*-values were calculated independently for each garden.

To establish a null distribution for the number of gardens in which a block significantly shifted, we performed 10,000 permutations. In each permutation, we randomized the *P*-values within each garden and identified the significant blocks. Using this approach, we determined a significance threshold of 10 gardens (**Fig. S30** permutation *P=*10^−6^). Based on this threshold, we identified 377 blocks significantly enriched across all gardens, encompassing 8,054 genes (**Table S11, S12**). GO enrichment analysis revealed 18 significantly enriched terms, spanning biological processes such as reproduction, development, and metabolism (**Table S13**). Notably, 22 haplotype blocks showed significant parallel shifts in at least 15 gardens. The genes within these blocks were significantly enriched for biological processes such as proline metabolism (**Table S11, S12**, **S13**). Proline has been well-documented in the literature for its critical role in plant responses to abiotic stress (*114*), suggesting that the proline metabolic pathway could be a promising target for engineering environmental resilience in plants.

### Text S12 LD patterns across generations

We estimated the LD blocks in the first, second, and third generations for experimental gardens with all three generations of data, and compared those with LD blocks calculated from the 231 founder population (**Fig. S17**). Compared to founder blocks (r2 = 0.5, median block length = 1.238 kb), 13 out of 19 experimental gardens had larger median values of LD blocks in the first generation (mean: 1.53 kb, IQR: 1.235 kb - 1.670 kb). In the second and third generations, we observed a further increase in the LD block sizes (r2=0.5) across GrENE-net gardens (mean second generation: 1.915 kb, IQR: 1.512 kb - 2.353 kb; mean third generation: 2.012 kb, IQR: 1.648 kb - 2.392 kb). In addition, we are sequencing individuals to estimate LD in evolved populations at high resolutions, and this will be for a future manuscript.

### Text S13. Supporting evidence for local adaptation

After detecting strong signals of natural selection in GrENE-net, evidenced by high repeatability within experimental gardens, we sought to understand the specific types of selection driving the evolution of GrENE-net populations. Among the three main types—directional selection, stabilizing selection, and disruptive selection—we found compelling evidence supporting the role of stabilizing selection in shaping these evolutionary dynamics.

Stabilizing selection was evident through patterns of local adaptation at the founder accession level. We recognize that we did not perform reciprocal transplants in GrENE-net, to test directly for local adaptation, but we used climate adaptation as a proxy of local adaptation. We evaluated local adaptation through our model, since the founder accessions achieved higher individual frequencies, indicating greater relative fitness, in experimental gardens with climates more similar to their regions of origin (**Fig. 3**). Local adaptation was observed across both geographic distances (**Fig. S32**) and climate distance (**Fig. 3A, Fig. S33**, **S34**), with individual accessions reaching higher relative frequencies when these distances approached zero.

To offer an alternative perspective, we identified a significant correlation between the climate of an accession’s origin and its inferred climate optimum, calculated from its relative success across experimental gardens (**Fig. 5D**, **Fig. S35**). Furthermore, we categorized GrENE-net gardens into high-repeatability (*r* > 0.2) and low-repeatability (*r* < 0.2) groups. In high-repeatability gardens, we found a significant relationship between the annual mean temperature of the experimental gardens and that of the best-performing founder accessions’ origins (**Fig. S36**, *R²* = 0.24, *P* = 1.75 × 10^−9^, *n* = 11). This suggests that in gardens with high repeatability, the most successful founder accessions originated from environments with similar annual mean temperatures, reinforcing the evidence for local adaptation in GrENE-net populations.

### Text S14 Species distribution model of Arabidopsis

Species Distribution Model or Environmental Niche Model fitted using MaxEnt and Worldclim v.2. Briefly, we used all *Arabidopsis thaliana* observations in GBIF (2022) (*115*), filtered them within main native Eurasian range (Longitude > 15W, Latitude >15N), and conducted a presence-only MaxEnt niche model with 19 Bioclim variables (scripts available at: https://github.com/MoisesExpositoAlonso/arabidopsisrange).

### Text S15. Phenotypic associations

We took a phenotypic selection approach using accession’s frequency as a proxy of fitness. Because the frequency of an accession, *log(p_1_/ p_0_)* can be reformulated into *log(w/ŵ)* (**Text S13**) we can use this for the classical Lande & Arnold approaches (*116*). With this, we can quantify correlations between accession frequencies and phenotypes from a curated and imputed dataset of accessions (*38*) (publicly available from arapheno.1001genomes.org, and (*117*)). Under the L&A approach, phenotypic selection is: *s = Cov (z, w)*. By conducting a simple Pearson correlation we have *r=s × Var(log(p_1_/p_0_)).* While we can transform correlation coefficients into phenotypic selection by scaling *Var(log(p_1_/p_0_))*, we report the correlation *r* for a simpler intuition (**Table S7**). Because our observed data is in fact change in frequency over time (*p_t_*) not simply relative fitness in one generation, we interpret correlations of phenotypes and frequency changes in later generations (e.g. three generations) as phenotypic evolutionary responses.

Focusing on life history and functional phenotypes (*38*), we found evolution of a suite of phenotypes (**Fig. S43, S44**). Because these shifts were across independent experimental population replicates within one site, and in parallel across sites of similar climates, we interpret this as a response to selection on populations. Further, we see that this phenotypic evolutionary response over time follows predictable climate gradients. For instance, the correlation coefficient between flowering time and accession relative frequency intensifies by Δ*r* = 0.1 for every one degree temperature change (*r _freq - ft._ ∼ a+ b temp.; b* = −0.1, *P =* 0.01*, R^2^ =* 0.31, see **Fig. S44**).

### Text S16. GEA overlap with climate GWAS

We compared our experimental GEA models with a classic climatic GWAS in *Arabidopsis* (*30*) by assessing the extent of overlap between haplotype blocks identified as significantly associated with climate in our experimental GEA models with a climate GWAS using GEMMA software (*118*) on the 231 founder accessions. For the latter, we used the climate of origin of each accession as phenotypes, defined using each of the 10 BIOCLIM variables calculated from ERA5-land data as outlined in the Methods section. We filtered SNPs with a minor allele frequency (MAF) less than 0.05 and used the same window-based approach, WZA, (*44*) used in the experimental GEA models for direct comparability. The extent of overlap in genomic regions and genes present within those blocks identified through the experimental GEA and GWAS methods is detailed in **Table S16** and **Fig. S49.**

Despite using similar linear statistical frameworks, we observed limited overlap between the genomic regions identified in our experimental GEA and those detected in the climate GWAS. We argue that these approaches are complementary rather than redundant, as they capture different aspects of adaptation, and while the exact causes of the discrepancy cannot be definitively determined, the methods and approaches differ in several things:

a. Historical vs. ongoing adaptation. Climate GWAS identify historical adaptation, where allele frequencies reflect past selection events in native environments. This approach assumes that alleles are fully locally adapted to their historical climates (*30*). Experimental GEA, in contrast, captures ongoing selection and adaptation processes by identifying allele frequency changes correlated with selective pressures in experimental gardens. Because populations are captured during ongoing adaptation, our approach may identify alleles most relevant to rapid responses to novel climates.
b. Alleles limited geographical distribution. In climate GWAS, alleles distribution is constrained by geographical barriers and limited gene flow between populations. As a result, some adaptive alleles may not be present where they would be most beneficial, leading to incomplete detection of adaptive loci. In our experimental GEA, we broke these limitations by planting a diverse population across different climates. This allowed all alleles to be tested in all climatic conditions, enabling the identification of novel genomic regions that may not be detected in climate GWAS.
c. Population structure as a confounding factor. Climate GWAS corrects for population structure to avoid spurious associations. However, this correction may remove adaptive alleles that are correlated with population structure (*119*). In our experimental GEA, we decoupled the climate of origin from the adaptive potential of alleles. While linkage between neutral and adaptive alleles due to population history remains a challenge, our approach reduces this confounding effect.
d. Pool sequencing vs. single genotype. Our experimental GEA used pooled sequencing of tens to hundreds of individuals per sample, providing robust estimates of allele frequencies and enhancing the detection of genetic-environmental associations. In contrast, climate GWAS relies on a single representative genotype per geographical location, typically the most prevalent or well-characterized accession.

Finally, GEAs conducted in natural populations or experimental settings are still considered a new approach, when compared to well established phenotypic GWAS. Nonetheless, simulations suggest that GEAs should be an effective tool for detecting locally adaptive loci (*120*, *121*) and recent empirical studies have begun validating GEA-identified genes, providing experimental evidence for their role in adaptation. For example, LSD1, a gene also detected in our analysis, was shown to influence flowering time and drought response in Arabidopsis under experimental conditions, further supporting its role in climate adaptation (*122*).

### Text S17. Genes associated with rapid adaptation

We identified a number of loci harboring natural variation whose evolutionary trajectories in GrENE-net experiments were correlated with climate. Below is a non-comprehensive list of top genes to showcase evidence of local adaptation, including *CAM5, TSF, Aquaporin-like* protein, and *CYP707A1* (For a comprehensive list of the top association peaks with temperature and precipitation see **Table S9, S10**).

#### CALMODULIN 5, CAM5, AT2G27030

*CAM5* (https://www.arabidopsis.org/locus?name=AT2G27030) encodes a Calmodulin gene out of 8 Calmodulin genes in *Arabidopsis*. *CAM5* was found to be the most reactive in expression under heat stress (*53*). Intriguingly, the top associations are found around an annotated third exon in transcript AT2G27030.3 which is spliced out of AT2G27030.1 based on TAIR (**Fig. 4, S50, S51**). To gather further evidence, we extracted normalized transcriptome data from the 1001 Transcriptomes & Methylomes (*54*) and showed that across accessions both transcripts exist and in fact the comparative level of expression of AT2G27030.1 across accessions decreases with the temperature at collection of origin of accessions whereas AT2G27030.3 increases (**Fig. S53**). Calmodulin protein structures are well studied in biochemistry and thus we wondered whether there were differences in protein predictions, especially in the amino acid sequence resulting from the additional third exon of AT2G27030.3. Comparing the isoforms 1 and 3 (https://alphafold.ebi.ac.uk/entry/Q682T9, https://alphafold.ebi.ac.uk/entry/F4IVN) (**Fig. S54**), we found a very low per-residue model confidence score (pLDDT<50), which made us hypothesize that a C-terminus intrinsically disordered domain (IDR) may be responsible for a change in function. To confirm the validity of the IDR prediction we used *Metapredict* v.3 (*123*), confirming the c-terminus of AT2G27030.3 is intrinsically disordered (**Fig. S54**). Several hypotheses emerge from the literature on IDR C-terminus of proteins, including the possibility of this with this IDR c-terminus altering the Ca^2+^ binding involved in signaling downstream (*124*) or the autoinhibition of *CAM5* effect by interference with a C-terminus (*125*).

#### TWIN SISTER OF FT, TSF, AT4G20370

*TSF* (https://www.arabidopsis.org/locus?name=AT4G20370) encodes a floral inducer that is a homolog of *FT* (AT1G65480). Plants overexpressing this gene flower earlier than Col-0. Loss-of-function mutations flower later in short days. TSF and FT play overlapping roles in promoting flowering, with FT as the dominant regulator. Together, they antagonize TFL1 in determining inflorescence meristem identity. *TSF* sequences show extensive variation among accessions and may contribute to quantitative variation in flowering time*. TSF* has a complex pattern of spatial expression; it is expressed mainly in phloem and expression is regulated by daylength and vernalization. Its homolog, *FT*, is well known to harbor natural variation including in the promoter region that changes expression, natural variation that may have been under natural selection and local adaptation (*126*, *127*). *TSF* is known to be a weaker but redundant form of *FT* and we thus expect to have similar phenotypic effects. In corroboration, we find a strong correlation between *FT* and *TSF* expression with flowering time measured in growth chamber conditions at 10℃ (Spearman’s *ρ_TSF_=* −0.35, *P=* 5.16×10^−9^*, n=*1220; *ρ_FT_ =* −0.64*, P=*2.09×10^−26^, *n=*1220). In agreement, the top alleles that were discovered to rapidly shift in frequency across the experimental environment showed an association with flowering time (**Fig. S56**, **S57, Table S18**). In previous studies *TSF* had been discovered in differentiation genomic scans (*48*) enriched in Iberian relict populations, in a peak that spans *TSF* and the neighboring *CHILLING SENSITIVE 4* (*CHS4*) or *LESION SIMULATING DISEASE 1* (*LSD1*), involved in pathogenesis and environmental stress. In agreement, we also found such a peak using a LMM GWA with climate of origin of the 231 founder accessions, although the peak is not the exact SNP as that identified in GrENE-net GEA (**Fig. S55**).

#### CRYPTOCHROME P940, CYP707A1, AT4G19230

*CYP707A1* (https://www.arabidopsis.org/locus?name=AT4G19230) encodes a protein with ABA 8’-hydroxylase activity, involved in ABA catabolism. Member of the *CYP707A* gene family. *CYP707A1* plays a role in determining the ABA levels in dry seeds and is involved in post germination growth (*51*). It has already been connected with natural variation in *Arabidopsis* (*52*). When overexpressed, *CYP707A1* leads to a decrease in ABA levels and a reduction in after-ripening period to break dormancy. We found accessions carrying the identified allele in experimental evolution, decrease the percentage of germination in laboratory conditions (**Fig. 4**).

#### Aquaporin-like protein, AT1G79780

AT1G79780 (https://www.arabidopsis.org/locus?name=AT1G79780) is a *CASP-LIKE PROTEIN 3A2, CASPL3A2*, uncharacterized protein family (UPF0497) for which we discovered natural alleles strongly correlating with precipitation of summer of experimental gardens (**Fig. S59 S60**).

### Text S18. Estimates of natural selection coefficient from allele frequency changes

To infer the magnitude of natural selection over alleles of interest, following a classic Wright-Fisher thinking, we used the knowledge of its allele frequency at a given site *p_i t_* across up to *i*=1…12 independent population replicates per year in *t*=1…3 time points. Because within each replicate *i* and time point *t* we know the number of reproductive adults sampled *n_i t_* and thus the number of adults with the alternative allele *k_i t_ = n_i t_ × p_i t_*, we can compute the probability of observing a vector *p_i t_* (i.e.) based on a Binomial likelihood that uses the number of collected samples and the frequency of the previous generation, typically expressed in log form: *log L(p_i t_ | n_it_, p_i t-1_)* = *Σ_i_ [ k_i_ × log(p) + (n_i_ - k_i_) × log(1 - p)]*.

We are interested in understanding whether an allele frequency change can be explained by neutral genetic drift or otherwise natural selection (whether it is causal or not, i.e. net natural selection can be driven by background selection or hitchhiking). Estimating selection genome-wide is challenging but because we are interested in focusing on a single top allele such as *CAM5* for which we have prior knowledge may be closely tagging a causal allele or may be one itself, we can simply conduct a grid search of selection coefficients *s_i_=* −1 – 1 by modifying the likelihood above *L(p_i t_ | n_it_, p_i t-1_, s_i_)*, using the expectation is that the true frequency at *p_t_ = [p_t-1_ × (1+s)] / [p_t-1_ × (1 + s) + (1 - p_t-1_)];* following the notation of relative fitness of the reference allele *A, w_A_* = 1, and alternative alleles *a*, *w_a_* = *1+s.* Scaling the calculated log likelihoods *L’_si_ = L_si_ / Σ_i_ L_si_*; we can plot the probability density of *s* given the data and generate an estimate of the mean (95% CI) of the selection coefficient for an allele on a given experimental environment between two generations (**Fig. S52**).

### Text S19. Conditional neutrality vs antagonistic pleiotropy tests

We found support of both antagonistic pleiotropy and conditional neutrality (**Fig. 5**). B performing enrichment tests, we found pleiotropy was significantly more likely to be observed than conditional neutrality (pleiotropy supported by 135/325 comparisons, Fisher’s exact test, odds ratio > 1.07, *P* < 0.039, conditional neutrality: 18/325 comparisons, Fisher’s exact test, odds ratio < 0.92, *P*<0.045). This comparison was conducted comparing LD block average frequency trajectories among experimental sites. We first identified haplotypes with significant frequency changes in each site by selecting the extreme 5% of *P*-values using a Binomial GLM with allele frequency as response and time as predictor. We then constructed contingency tables for each pair of sites, categorizing haplotypes based on whether they exhibited significant frequency changes in one or both sites. To assess non-independence, we performed Fisher’s exact test, calculating both the *P*-values and the odds ratio (OR) to determine support for pleiotropy (where haplotypes were more likely to change in both sites) versus conditional neutrality (where haplotypes changed only in one site). Second, for haplotypes showing significant frequency changes in both sites, we examined the direction of selection (increasing or decreasing frequency) to distinguish between antagonistic pleiotropy (selection in opposite directions) and synergistic pleiotropy (selection in the same direction). Fisher’s exact test was used to calculate the odds ratio (OR) and *P*-values for antagonistic vs synergistic pleiotropy.

## SUPPLEMENTAL DATASETS

Supplemental datasets are available in the online materials of this manuscript as well as in www.grene-net.org/data.

**Dataset S1. Samples and census records from participants**

Descriptions of a total of 2,410 flowers samples and 1,178 populations census records were collected and submitted by participants.

Dataset_S1_GrENE-net_records_2017-2021.xlsx

**Dataset S2. Daily climate loggers**

Dataset_S2_daily_climate_logger.csv

**Dataset S3. Biweekly photographic records during the growing seasons**

Dataset_S3_biweekly_photorecords_diary

**Dataset S4. Founder allele frequencies**

Dataset_S4_founder_allele_frequency.txt

**Dataset S5. Founder accession frequencies**

Dataset_S5_founder_accession_frequency.txt

**Dataset S6. Evolved populations’ allele frequencies**

Dataset_S6_evolved_allele_frequency.csv

**Dataset S7. Evolved populations’ accession frequencies**

Dataset_S7_evovled_accession_frequency.txt

**Dataset S8. Accessions relative frequency GWAs**

Raw results from GWAs on the relative frequencies of accession at each experimental garden Dataset_S8_accession_frequency_gwas.zip

**Dataset S9. GEA WZA results**

Raw results from GEAs + WZA models on the relationship between allele frequencies and environmental variables

Dataset_S9_wza_results.zip

## SUPPLEMENTAL TABLES

Supplemental tables are available in the online materials of this manuscript as well as in www.grene-net.org/data. Temporarily, all Supplemental tables are available via a Google Drive link:

**Table S1. GrENE-net sites information.**

Contact and location information about each experimental garden in the GrENE-net experiment.

Table_S1_site_information.xlsx

**Table S2. GrENE-net accession information.**

Founder accession information including country of origin, seed stock number, latitude and longitude, and relative proportion in the original seed mix.

Table_S2_founder_accession_information.xlsx

**Table S3. Census records curated**

Curated population’s census records from participant-submitted records (**Dataset S1**)

Table_S3_census_data_curated.csv

**Table S4. Population size estimations**

Population sizes measured by participants and estimated based on the number of flowers collected

Table_S4_population_size_estimations.csv

**Table S5. GrENE-net sample collection and sequencing coverage summary.**

Flower samples collected by GrENE-net participants including location, replicates, year, date, flowers inside tube, and sequencing information. Sample names are coded as a strings that contain key information following: [Moi Lab][Flower Heads][site number: 01…60][plot number:01…12][year][month][day]; for example: MLFH010120180409.

Table_S5_sample_collection_sequencing_library.xlsx

**Table S6. Summary of the sequenced GrENE-net samples.**

Summary of how many samples were collected per site per year, the total number of flowers collected.

Table_S6_summary_per_tray_generation.xlsx

**Table S7. Associations between accessions’ phenotypes and frequency change across sites**.

Correlation between accession frequencies and key phenotypes per experimental garden. Columns are coded by phenotype Pearson’s correlation estimate (r_<phenotype>) and *P-*value (r_p_<phenotype>).

Table_S7_accessions_phenotypes_n_frequency_change.tsv

**Table S8. Stabilizing selection parameters.**

Estimates of *W_max_*and *Vs* based on accession relative frequency change in the first generation and temperature distance to origin of accessions using a random effect model in MCMCglmm.

Table_S8_Wmax_Vs.csv

**Table S9. Top hits of genome-wide frequency associations with mean annual temperature.**

Genome-wide significant LD blocks across temperature GEA models (LFMM, Kendall, bGLM) and time samples (first generation, last generations). All genes within each LD block are reported including gene descriptions.

Table_S9_top_hits_GEA_matemperature.csv

**Table S10. Top hits of genome-wide frequency associations with precipitation.**

Genome-wide significant LD blocks across precipitation GEA models (LFMM, Kendall, bGLM) and time samples (first generation, last generations). All genes within each LD block are reported including gene descriptions.

Table_S10_top_hits_GEA_precipitation.csv

**Table S11. Comparisons of antagonistic pleiotropy or conditional neutrality.**

Pairwise comparisons of significant genomic LD blocks increasing or decreasing in frequency over time (bGLM) to test for evidence of natural selection trade-offs. We tested trade-offs and conditional neutrality using contingency tables and Fisher’s test (see Materials & Methods) and reported Odds ratios and *P-*values.

Table_S11_antagonistic_pleiotropy_conditional_neutrality_test.csv

**Table S12. Leave-one-out prediction accuracies.**

Leave-one-out cross-validation predictability of accession relative frequency based on four predictions (see **Text S12**). Pearson’s, Spearman’s correlations and *R^2^* are reported for every site and replicate.

Table_S12_LOO_accuracy.csv

**Table S13. LRT-1 significant parallel blocks across all experimental gardens.**

Table_S13_significant_parallel_blocks_aross_gardens.xlsx

The chromosomal positions of significantly over represented parallel-shifting LD blocks across all experimental gardens

**Table S14. Genes overlapping with LRT-1 significant parallel blocks.**

Table_S14_genes_overlapping_significant_parallel_blocks.xlsx

The Arabidopsis gene IDs of genes in the significantly over represented parallel-shifting LD blocks

**Table S15. Gene ontology enrichment results from genes identified by LRT-1.**

Table_S15_gene_ontology_enrichment_genes_overlapping_parallel_blocks.xlsx

The gene ontology enrichment analysis results from genes in the significantly over represented parallel-shifting LD blocks.

**Table S16. Top hits overlap between classic climate GWAS and GrENE-net’s GEA.**

Table summarizing overlap of LD blocks and the genes within those blocks, showing significant associations with different WorldClim BIOCLIM variables using GrENE-net’s GEA approach of allele frequency trajectories and the standard Genome-Wide Association using climate of origin of *Arabidopsis* accessions (see Methods).

Table_S16_hblocks_overlap_GWAS_GEA.xlsx

**Table S17. Coefficients for population dynamics analysis**

Table presenting the results of linear and quadratic regression analyses used to model population dynamics across 2 or 3 generations. Each row corresponds to a specific site and includes the intercept, slope (linear coefficient), and, where applicable, the quadratic coefficient along with their associated p-values. See Fig. 1F and Fig. S8 for visualizations of this data

Table_S17_population_dynamics.csv

**Table S18. Effect of GEA identified alleles near TSF in flowering time**

Effect of alleles identified by Genome-Environment Association (GEA) near the TSF gene on flowering time in *Arabidopsis*. The effect of each allele on flowering time was evaluated under two growth chamber conditions: 10°C (FT10) and 16°C (FT16). For each SNP, the table provides the effect size from LFMM (Latent Factor Mixed Models), correlation coefficients (Kendall, Pearson, and Spearman) of the temperature gradient and the change in allele frequency in the 3rd generation data of the expeirment with associated p-values, and the mean difference in flowering time between accession carrying the alternative and reference alleles. The significance of the mean difference was assessed using Wilcoxon tests.

Table_S18_tsfalleles_flowering_time.csv

## SUPPLEMENTAL FIGURES

### EXPERIMENT SETUP AND SUCCESS

**Figure S1.**
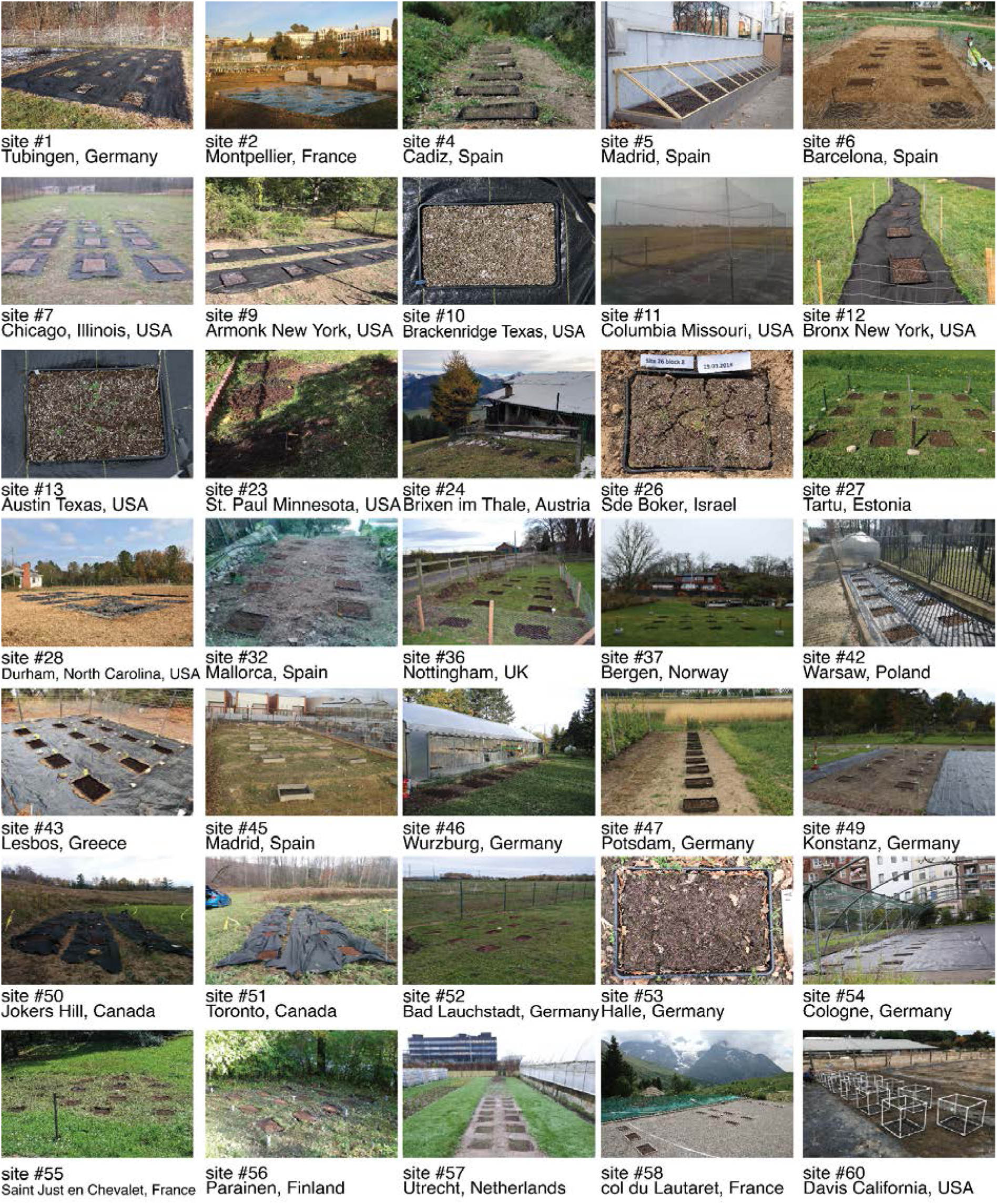
Experimental garden pictures of GrENE-net Overview of experimental setups across diverse GrENE-net gardens. In total, GrENE-net set up 43 experimental gardens globally, 35 out of 43 participating gardens took photographic records during the experiment.

**Figure S2.**
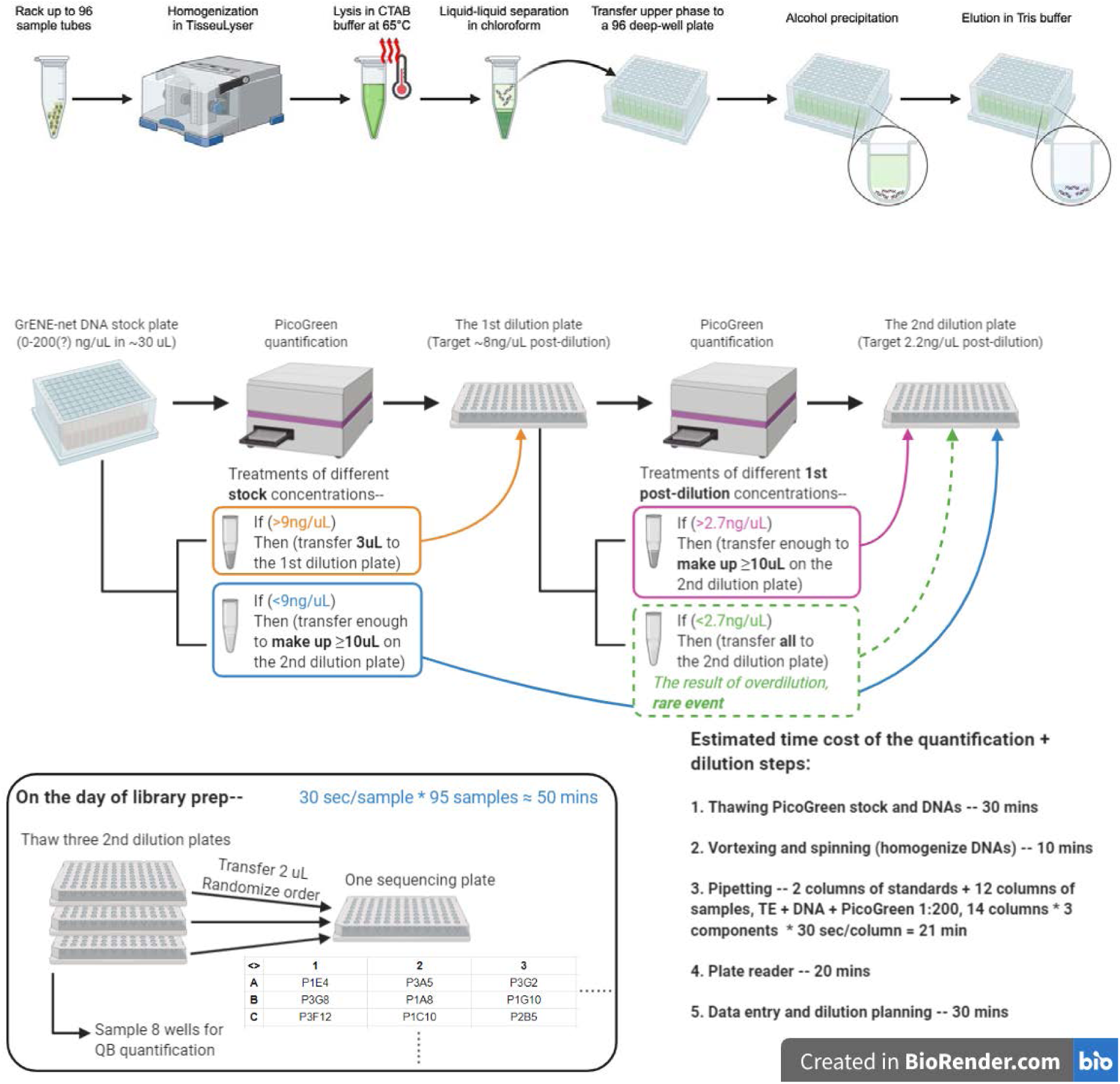
Preparation of GrENE-net DNA for sequencing. Workflow of sample processing starting with grinding flowers in 96 deep well plates for DNA extraction, followed by quantification for either further dilution or direct use. Tissue heterogeneity required an intermediate step in DNA normalization across 96 well samples. Next, we conducted tagmentation-based library preparation in 96 well plates.

**Figure S3.**
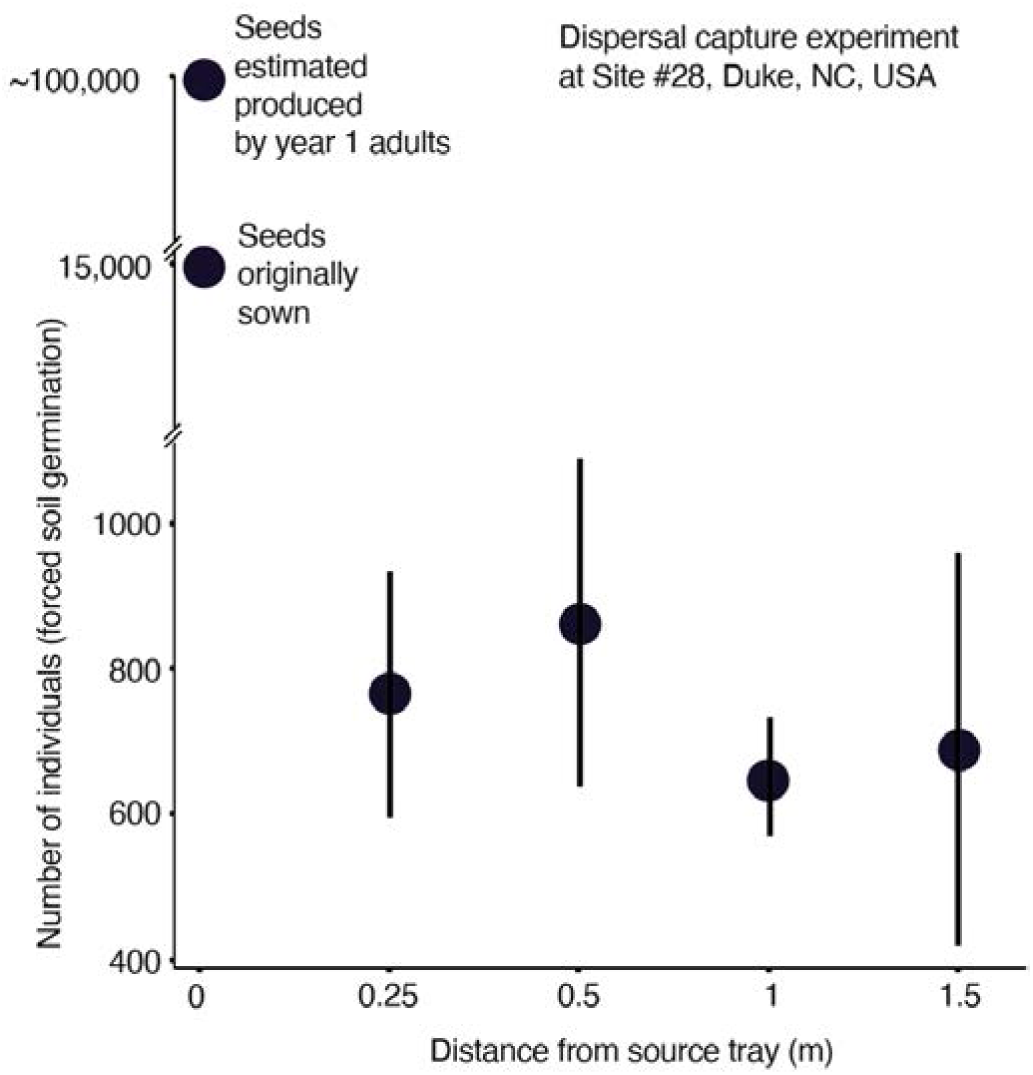
Mean estimated seed dispersal from source GrENE-net trays in site 28. Data from dispersal capturing experiment of site #28 (Duke, NC, USA), where empty soil trays were placed at different distances from GrENE-net (“source”) trays containing the seed mix and evolving naturally. The number of seeds dispersed into empty “capturing” trays were revealed by forced germination in the greenhouse (see **Text S6**) one year after original GrENE-net sowing. Distance from source tray is in 4 bins based on the distribution of distances of the dispersal trays. There were 11 capturing tray values per distance and the error bars are standard error. Source tray seeds are added for comparison (i.e. each GrENE-net tray was sown with ∼15,000 seeds). Estimated seed production from adult fecundity in the first year of reproduction (a season prior studying seed abundance in capture trays) was estimated based on the recorded number of adults that season and typical fruit and seed set of *A. thaliana* (**Text S6**).

### FLOWERING COLLECTIONS

**Figure S4.**
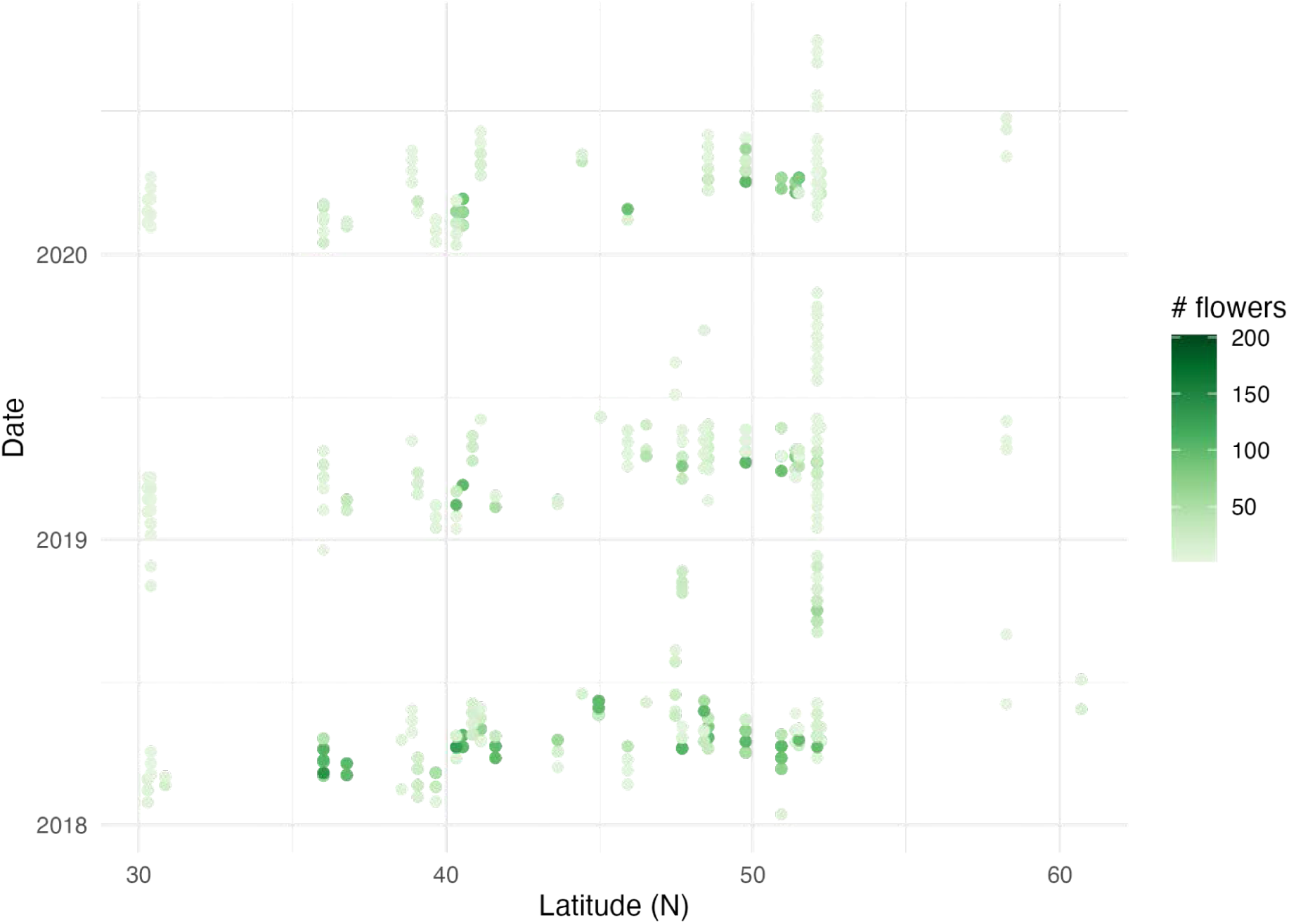
Flowering date collections across time and latitude Representation of all flower pool collections of Phase 1 GrENE-net that include the first three years (2018-2020). The number of pools sampled within a tube is represented by the green color gradient, and all samples across 30 sites are plotted across a latitudinal gradient.

**Figure S5.**
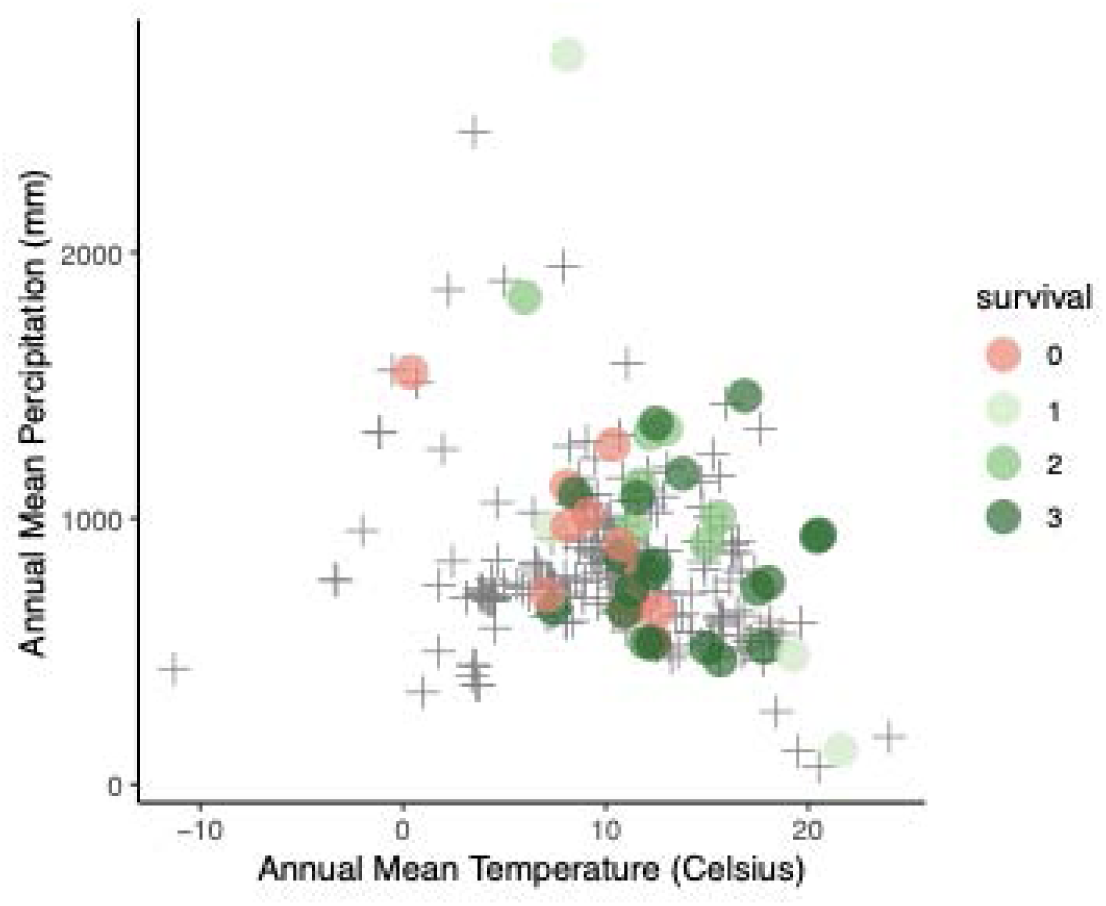
Environmental distribution of GrENE-net experimental gardens GrENE-net experiments, *n* = 43, organized by annual temperature and precipitation and colored by the number of years experimental populations survived. Crosses represent the 231 founder accessions’ climate of origin.

**Figure S6.**
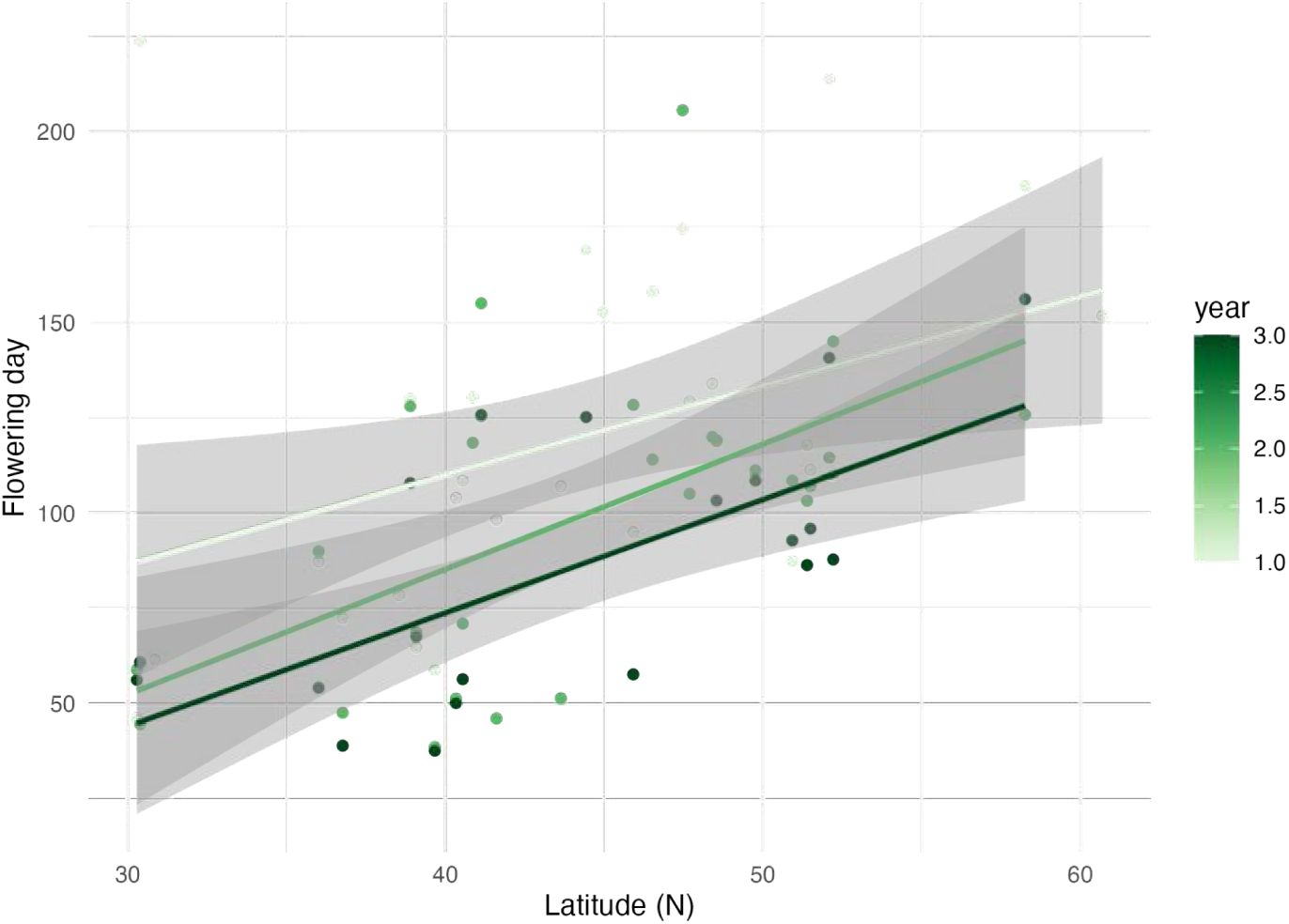
Weighted average of flowering time from GrENE-net The day of the year of peak sampled flowers is weighted by the proportion of flowers of the total collected per site within a season to estimate peak flowering time across years and sites. Regression of flowering time against latitude was conducted for each of the years.

### POPULATION SIZES & SURVIVAL

**Figure S7.**
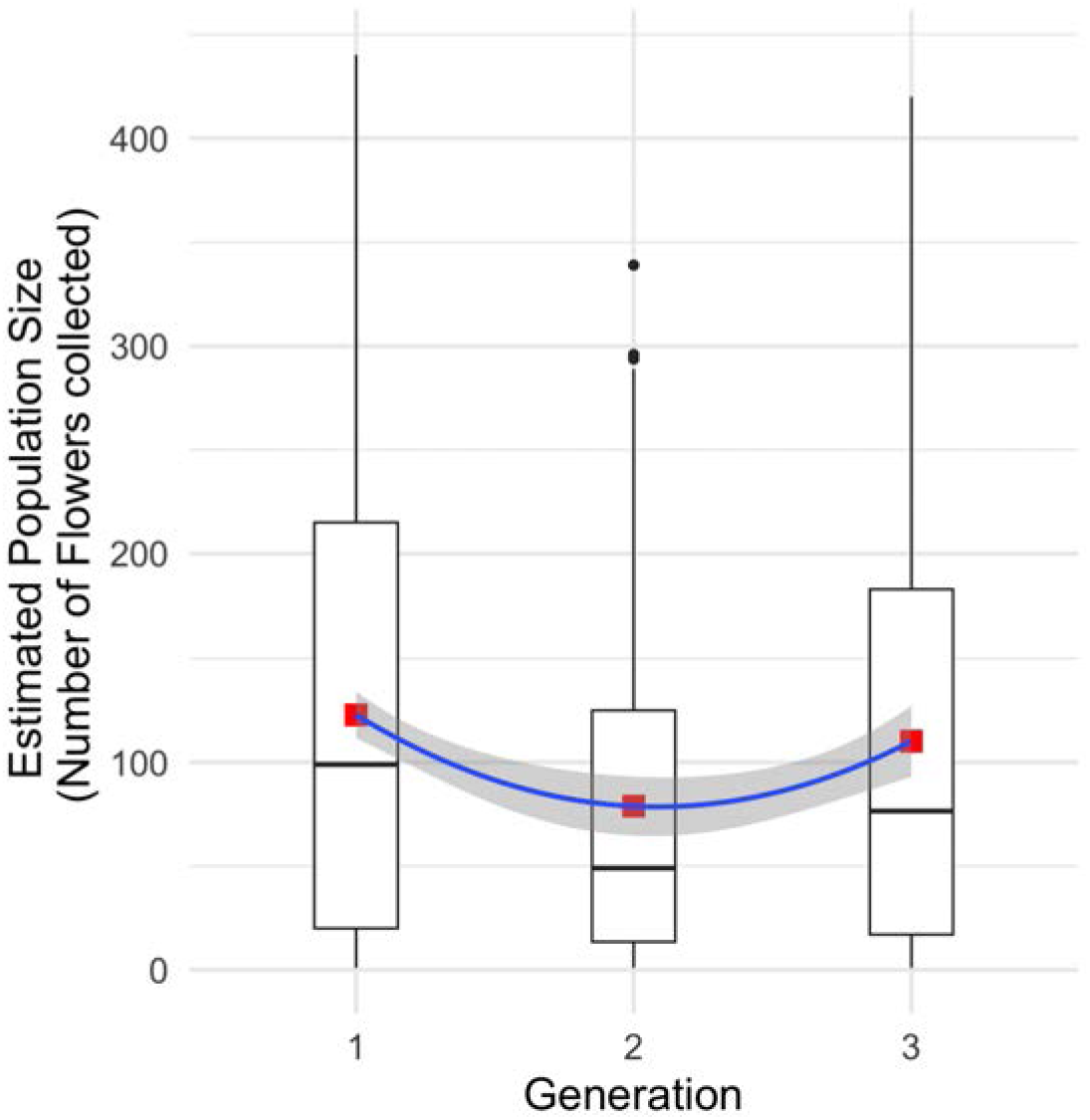
Boxplot of estimated census population size across generations. The red square in the box plot indicates the mean value of the distribution. The blue curve is the quadratic polynomial fit of the estimated population size over generations (populations ∼ a + b·time + c·time^2^). The gray shade indicates the standard error of the fit.

**Figure S8.**
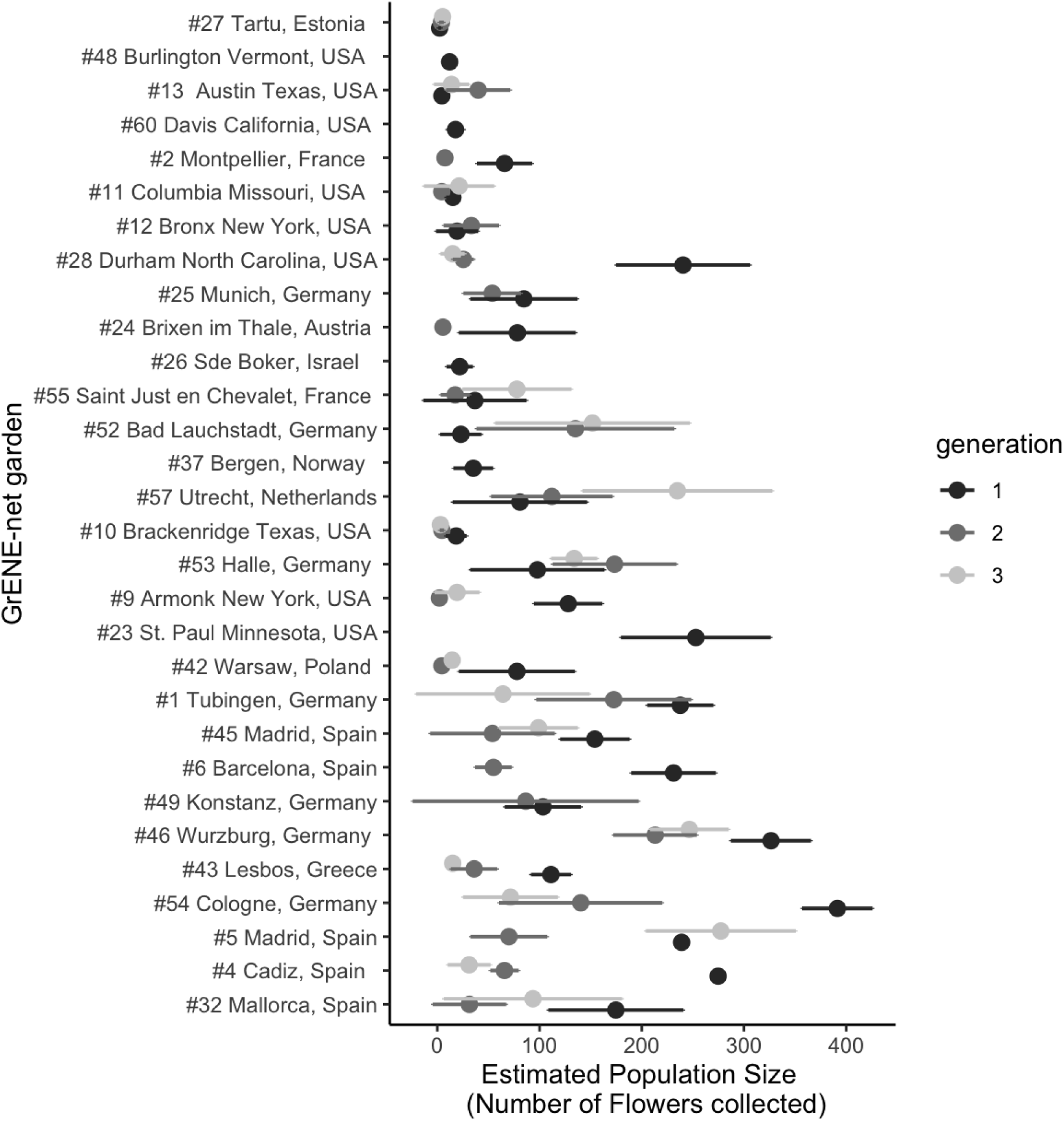
Population size across experimental gardens. Population size estimated from census-size-based model (**Text <u>S2</u>**), with experiments ordered by the degree of repeatability of evolution (low up, high down) based on **Fig. 1**. Intervals indicate 95%CI.

**Figure S9.**
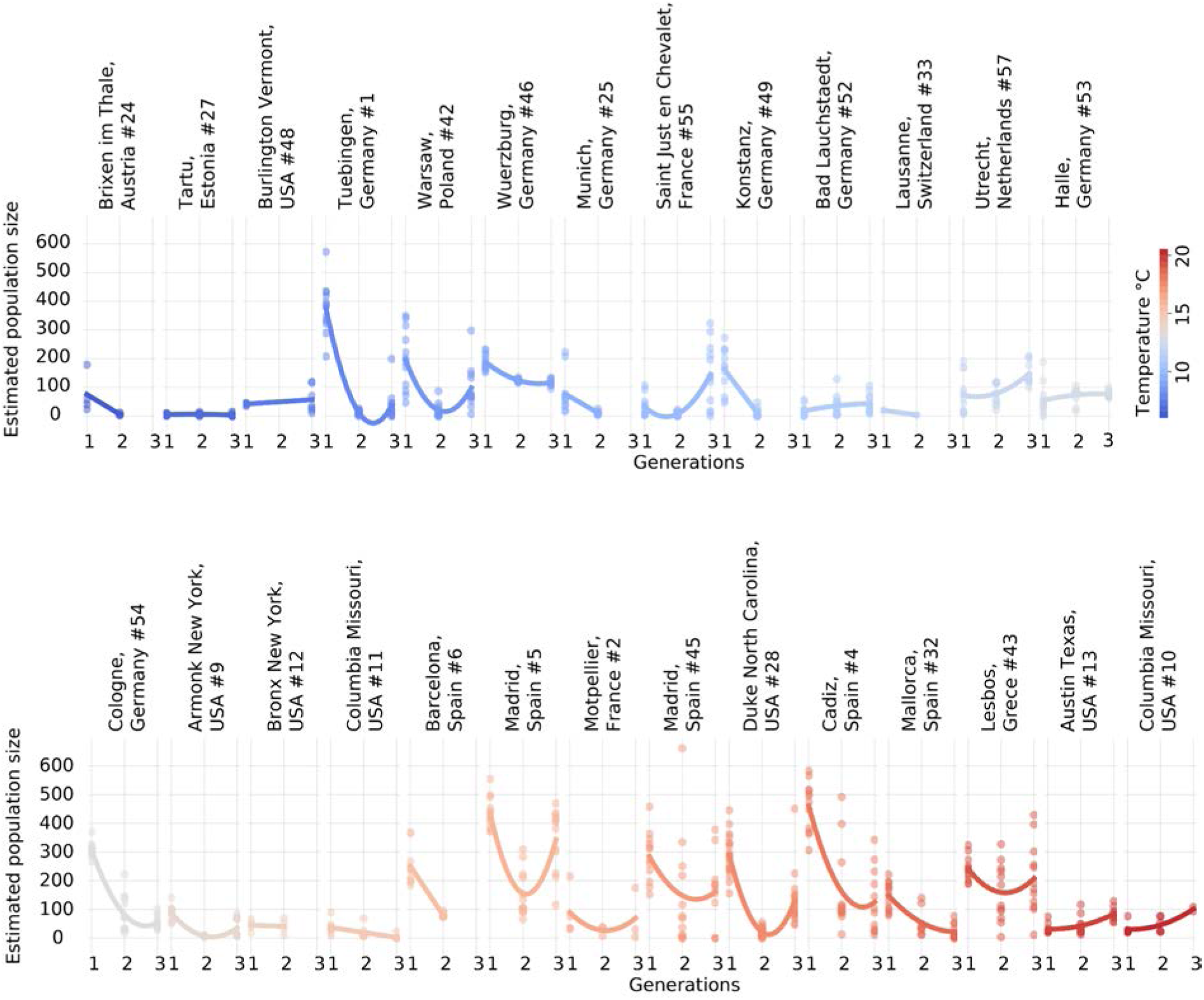
Evolutionary rescue by garden temperature. Population size trajectories fitted with a polynomial (y ∼ a + bx + cx^2^) across experimental gardens sorted by annual temperature.

**Figure S10.**
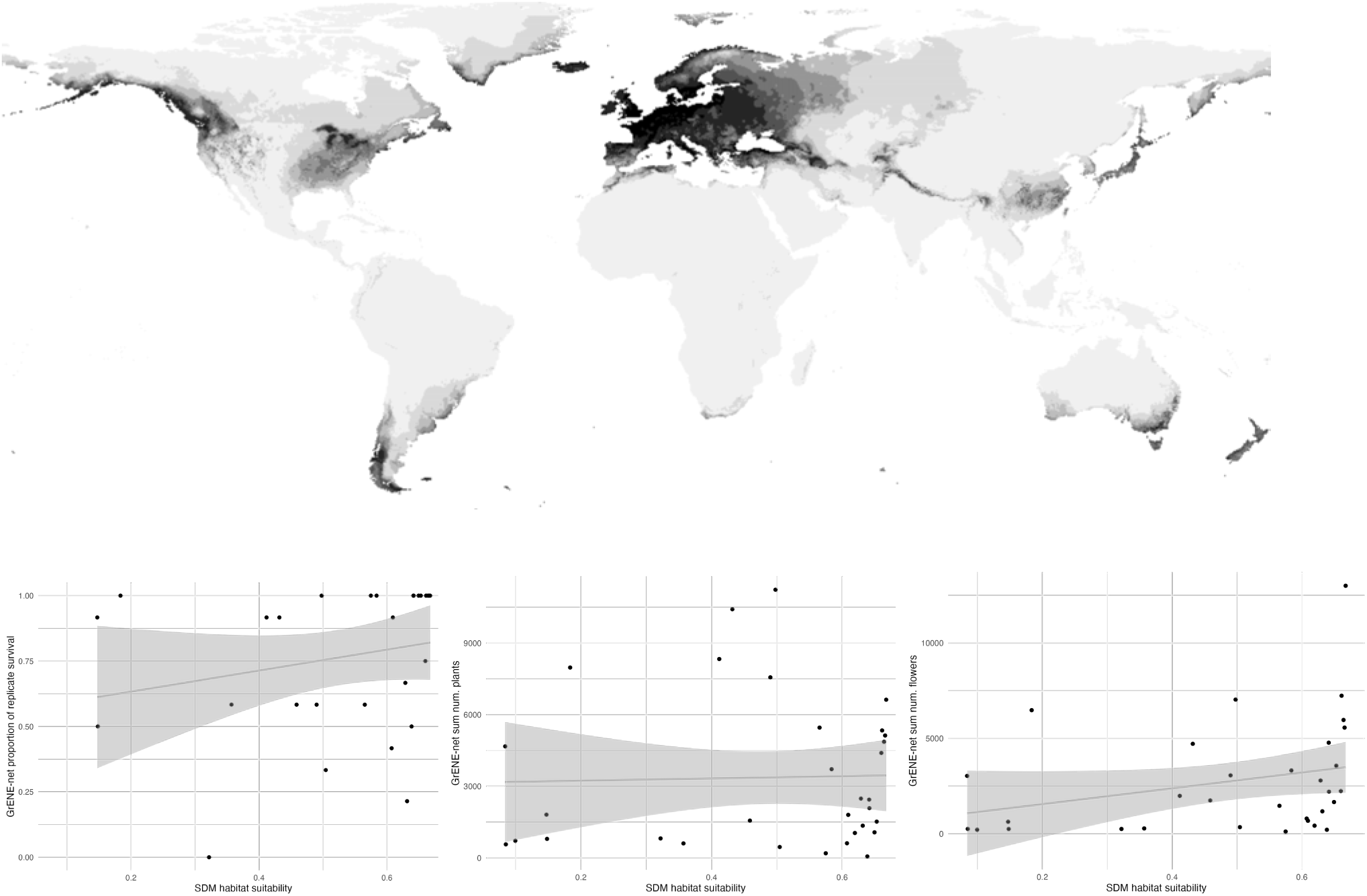
Species distribution model of *Arabidopsis* and population size metrics. (**upper**) Species Distribution Model or Environmental Niche Model fitted using MaxEnt and Worldclim v.2 (https://github.com/moiexpositoalonsolab/arabidopsisrange). Population size and flower distributions in comparison to species distribution model. Relationship between species distribution model (SDM) habitat suitability and population size and reproduction metrics. (**left**) Proportion of replicate survival in up to 5 years out of 12 starting replicates. (**middle**) The total number of flowers collected across all replicates within one location over years. (**right**) The sum of number of plants estimated from population census within one location over years. fig-compare-population-survival-with-SDM.R

### POOL SEQUENCING & GENOMIC QC

**Figure S11.**
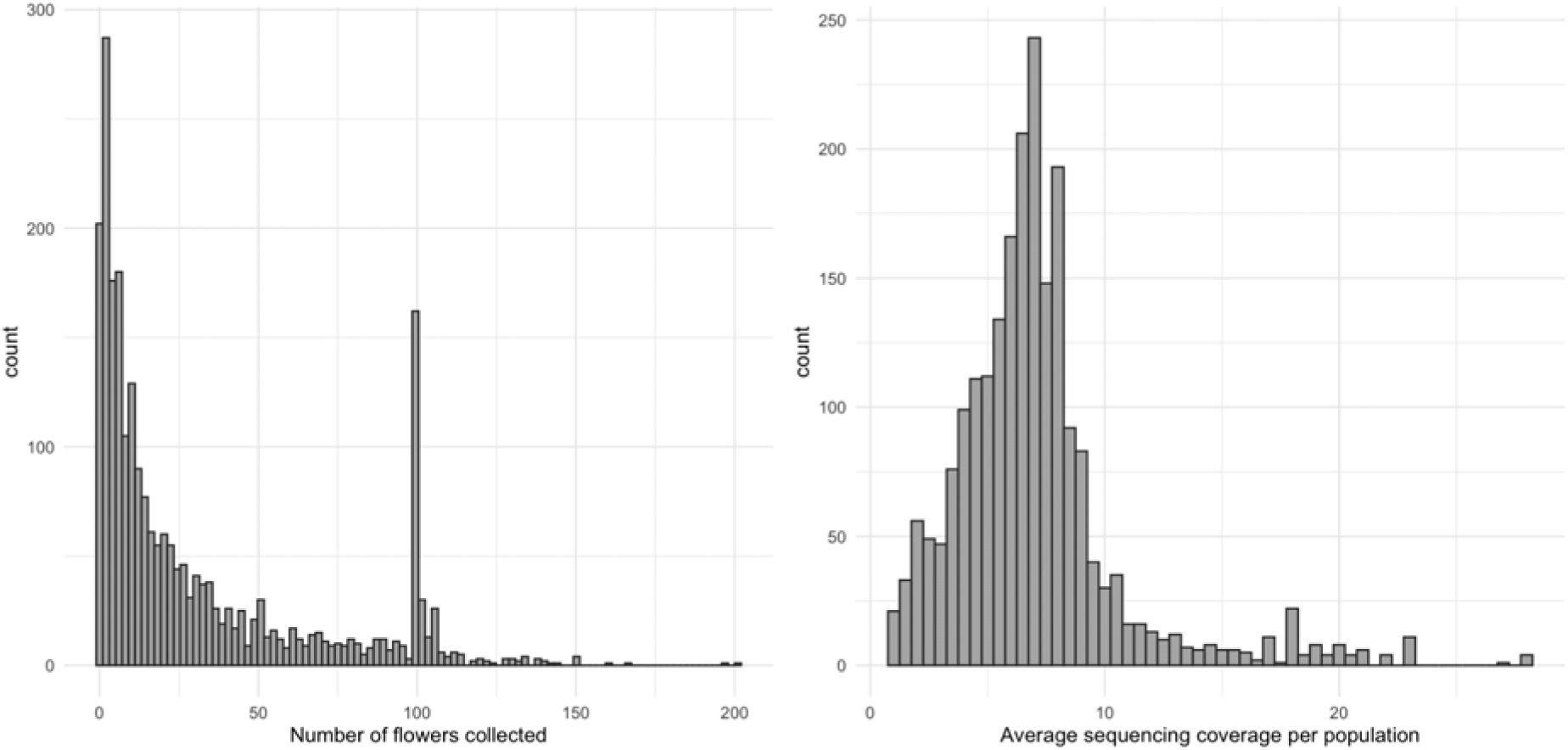
Distribution of flowers collected and sequencing coverage. Distribution of number of flowers collected per Pool-seq sample (**left**) and distribution of genome-wide sequencing depth per Pool-seq sample (**right**)

**Figure S12.**
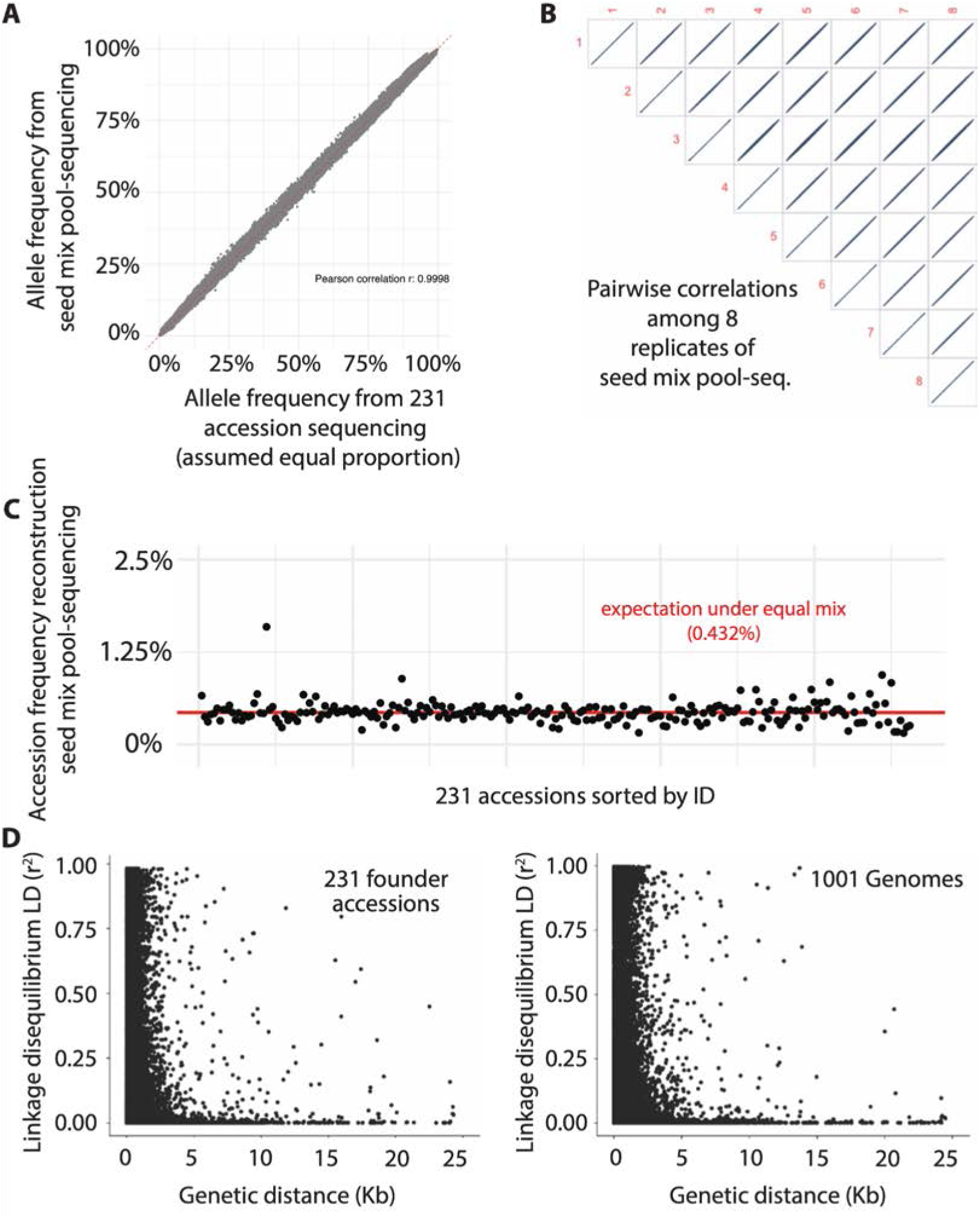
Allele and accession frequencies of the founder population. (A) Inference of allele frequencies using hapFIRE from deep Pool-seq sequencing of seed mixture of 231 accessions (y-axis) compared to the frequency expected from the 231 accessions VCF (x-axis). (B) Correlation of allele frequencies between 8 independent subsamples of seed deep Pool-sequencing. (C) The reconstructed founder accession frequencies. The red line indicates the equal mix (1/231 ∼ 0.4%) of all the founder accessions. (**D**) Linkage disequilibrium of the 231 founder accessions compared to the 1001 Genomes dataset.

**Figure S13.**
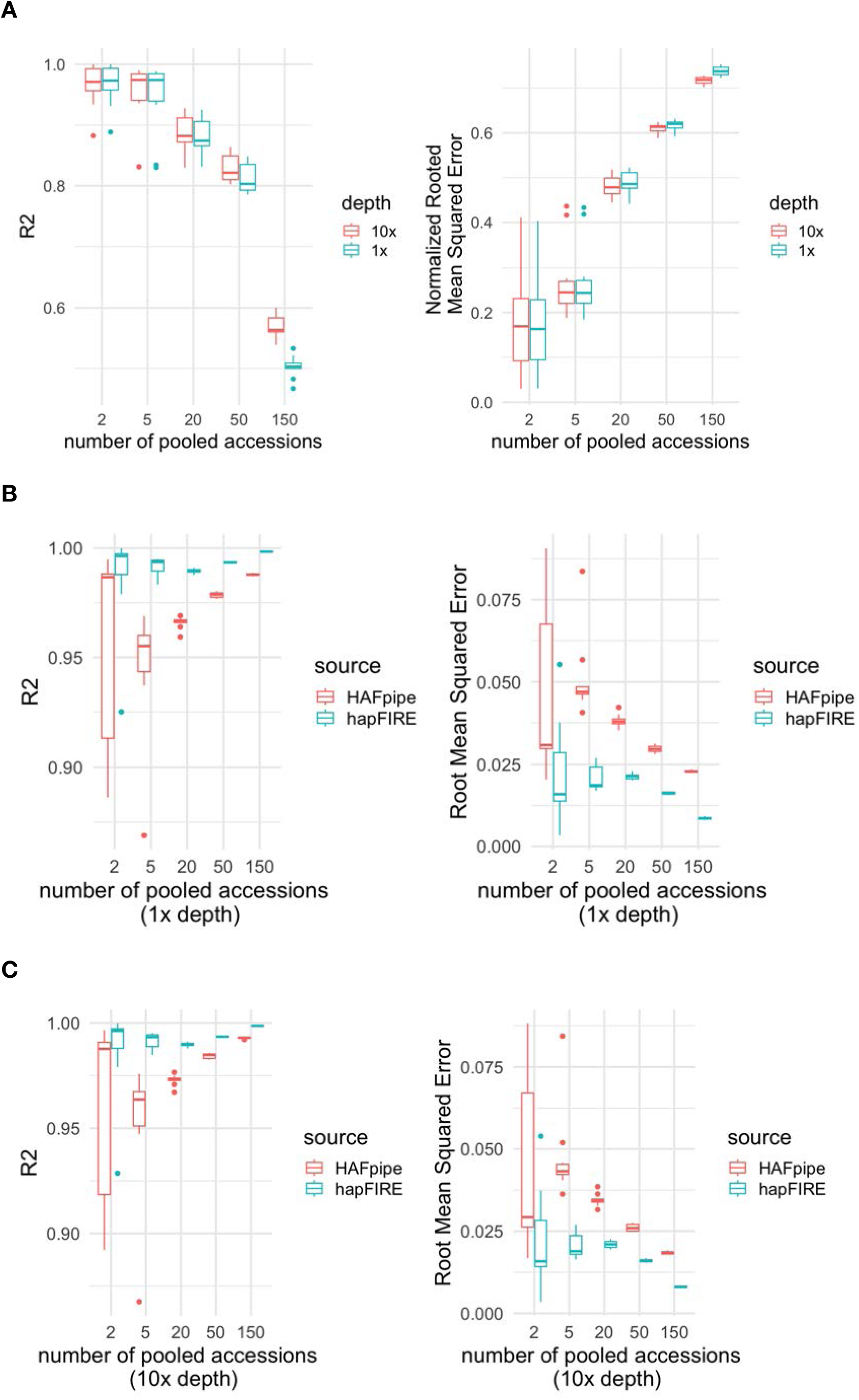
Accuracy of HapFIRE of accession and allele frequency from Pool-seq. (**A**) We simulated the pooled whole genome DNA sequencing with equal proportion of pooled accessions ranging from 2 to 150, and sequencing depth of 1x and 10x. We report *R^2^* and normalized root mean square error (RMSE/sd) of ***accession*** relative frequency. (**B**) Conducting the same simulations using 1x sequencing depth we study the accuracy of ***allele*** frequency *R^2^* and RMSE (corresponds to the % error in estimating allele frequency change). We compare our new HapFIRE with the original HAF-pipe wrapper (see **Text <u>S4</u>**). (**C**) Same as (B) with 10x sequencing depth.

**Figure S14.**
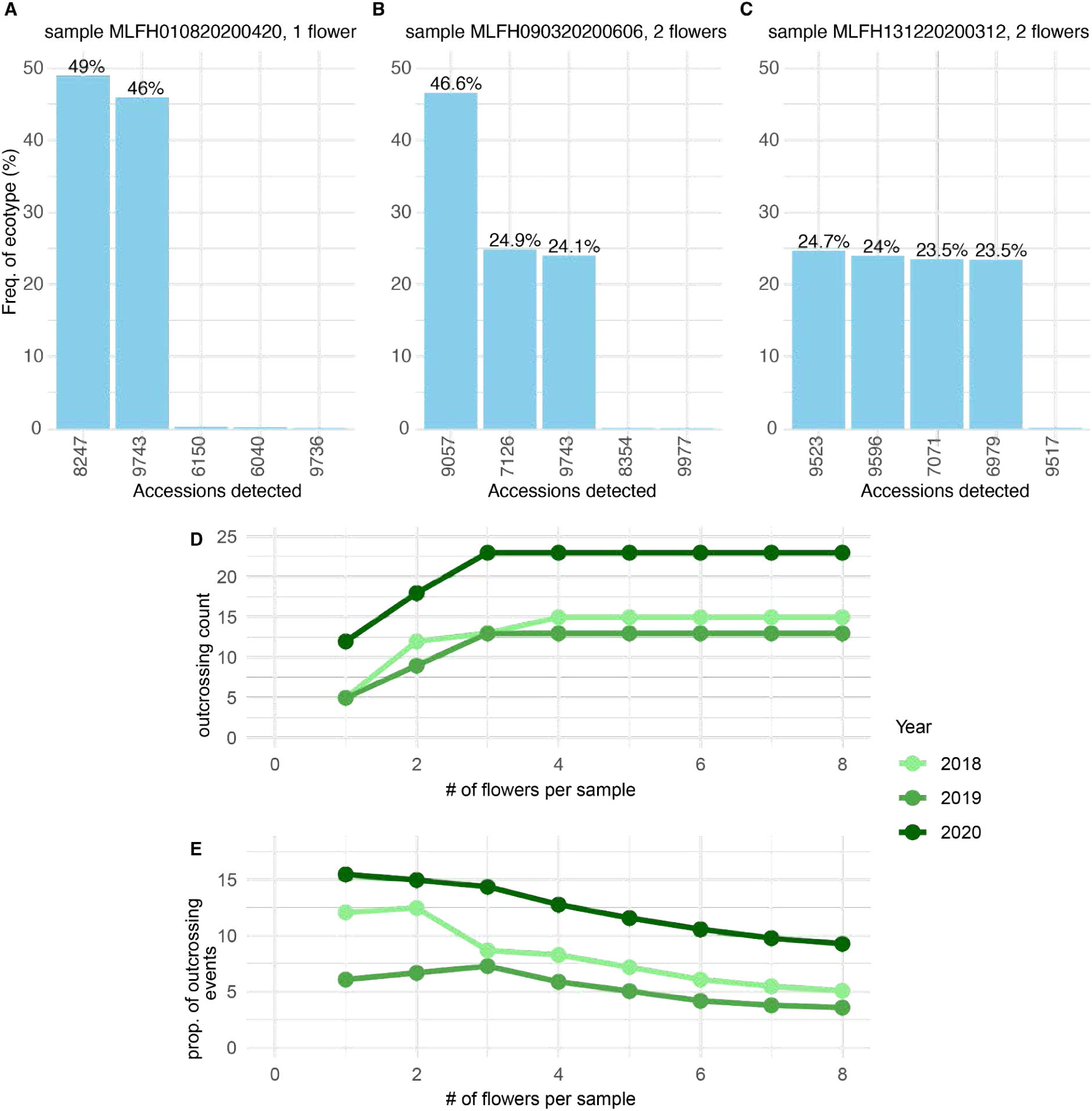
Approximate outcrossing rate based on accession fraction reconstruction. (**A-C**) Example sequencing of pooled samples of one or two flowers and reconstructed relative frequency of 231 accessions. (**A**) Sample illustrating a case of F1 with the expected 50/50%. (**B**) Sample where we detected an inbred line and an F1, with expected 50/25/25%. (**C**) Sample illustrating a cross of two F1 x F1 plants, with expected 25/25/25/25% (**C**). (**D**) Using this rationale, we used all samples with up to 8 flowers to identify outcrossing events (**Text <u>S5</u>**). (**E**) Approximate outcrossing rate based on samples with only one sample (*n*=8085) ranged 6-16%.

**Figure S15.**
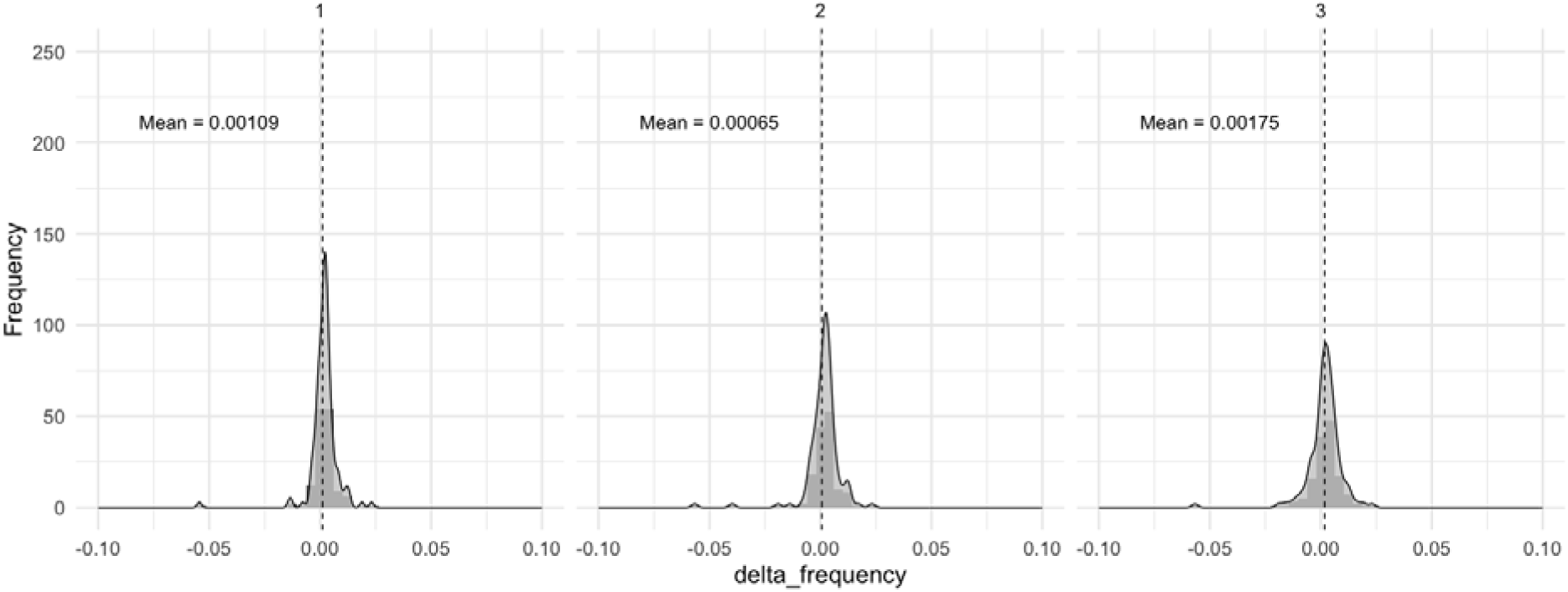
The distribution of mean genome-wide allele frequency changes under neutral simulation The figures from left to right are the distributions of mean allele frequency change, under random uniform sampling of the founder accessions over 3 discrete generations. Changes in LD pruned SNP frequencies are: founder to generation 1 population, founder to generation 2, and founder to generation 3.

**Figure S16.**
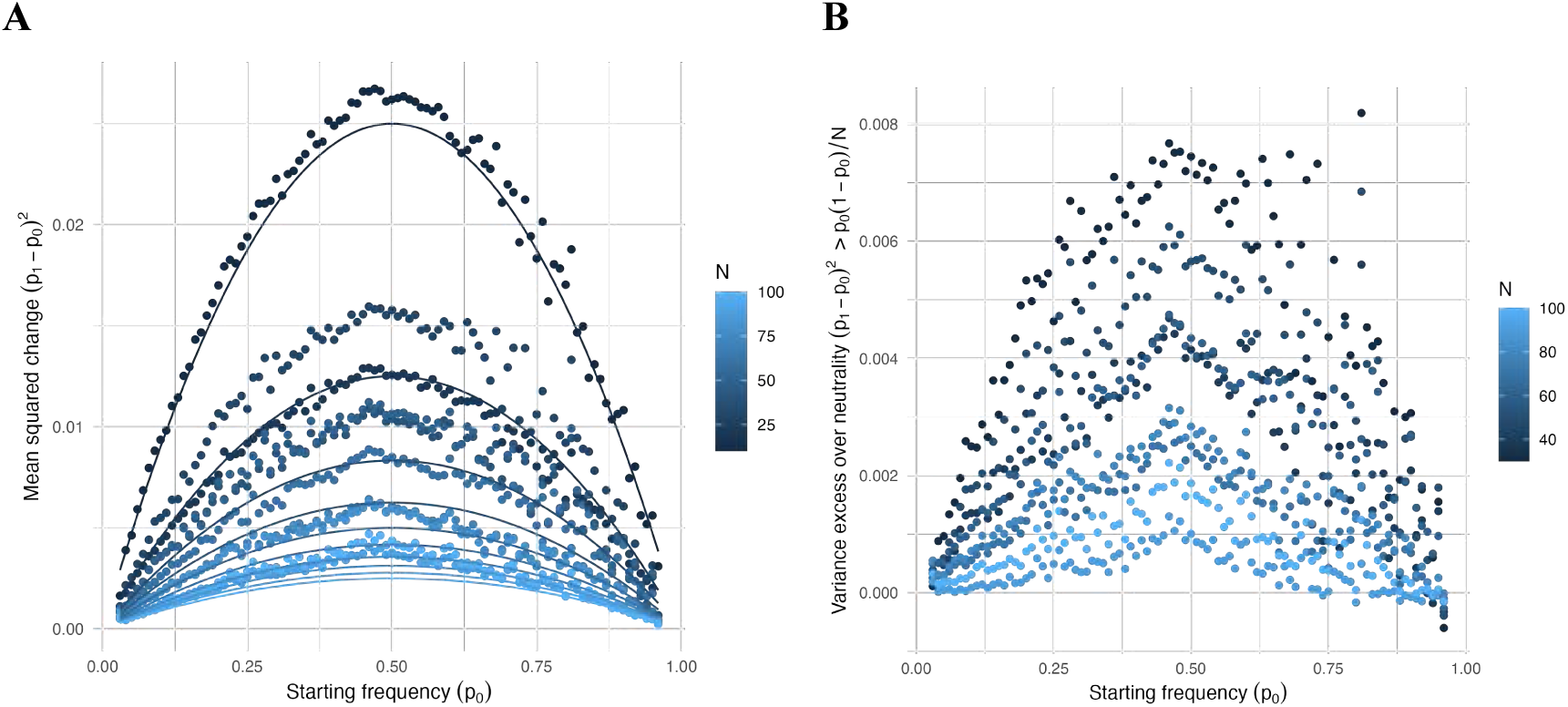
Expected frequency change under drift between two generations. Under drift we expect that the mean square change in frequency (i.e. variance in allele frequency) to be dependent on the starting frequency and population size following: *(p_1_-p_0_)^2^* = *p_0_ (1-p_0_) / N;* in a selfing plant. (**A**) Taking all the data across GrENE-net founder-to-generation 1 (n=11,663,490) we took the empirical average of *(p_1_-p_0_)^2^* for alleles rounded ±0.5% in population samples rounded ±5 flowers (dots), and overlaid theoretical expectation of *p_0_ (1-p_0_) / N* (lines). (**B**) Ratio excess of variance of empirical mean square / expectation calculated from (A). Ratio mean = 1.5. Ratio median[IQR] = 1.53 [1.10 - 1.80].

**Figure S17.**
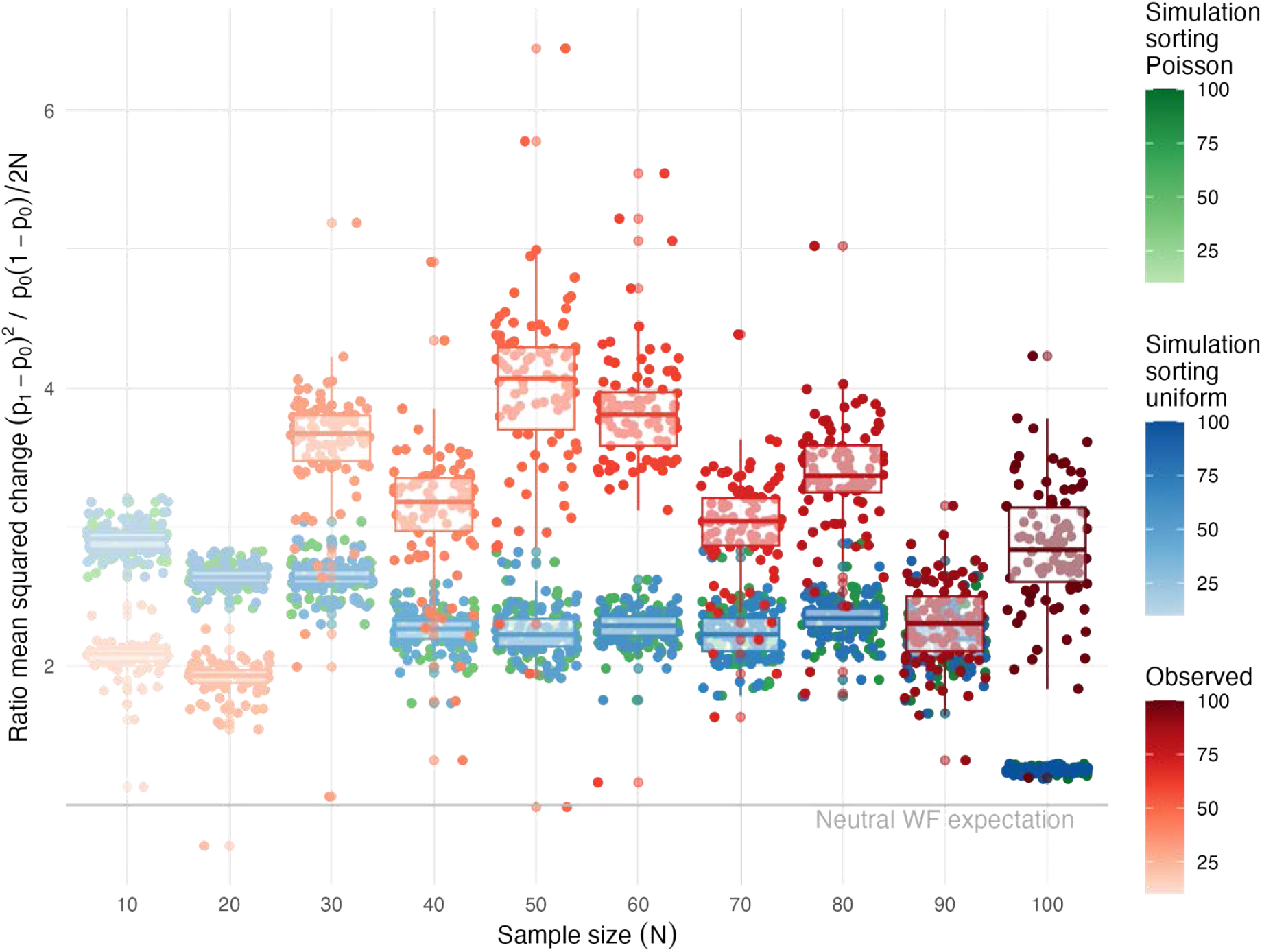
Expected frequency change under drift between two generations. The figure shows the observed variance in allele frequency change (red) compared to neutral Wright Fisher expectation, neutral simulation assuming uniform sampling of the accessions (blue) and neutral simulation assuming random sampling from a Poisson distribution (green).

### FST and LD pattern

**Figure S18.**
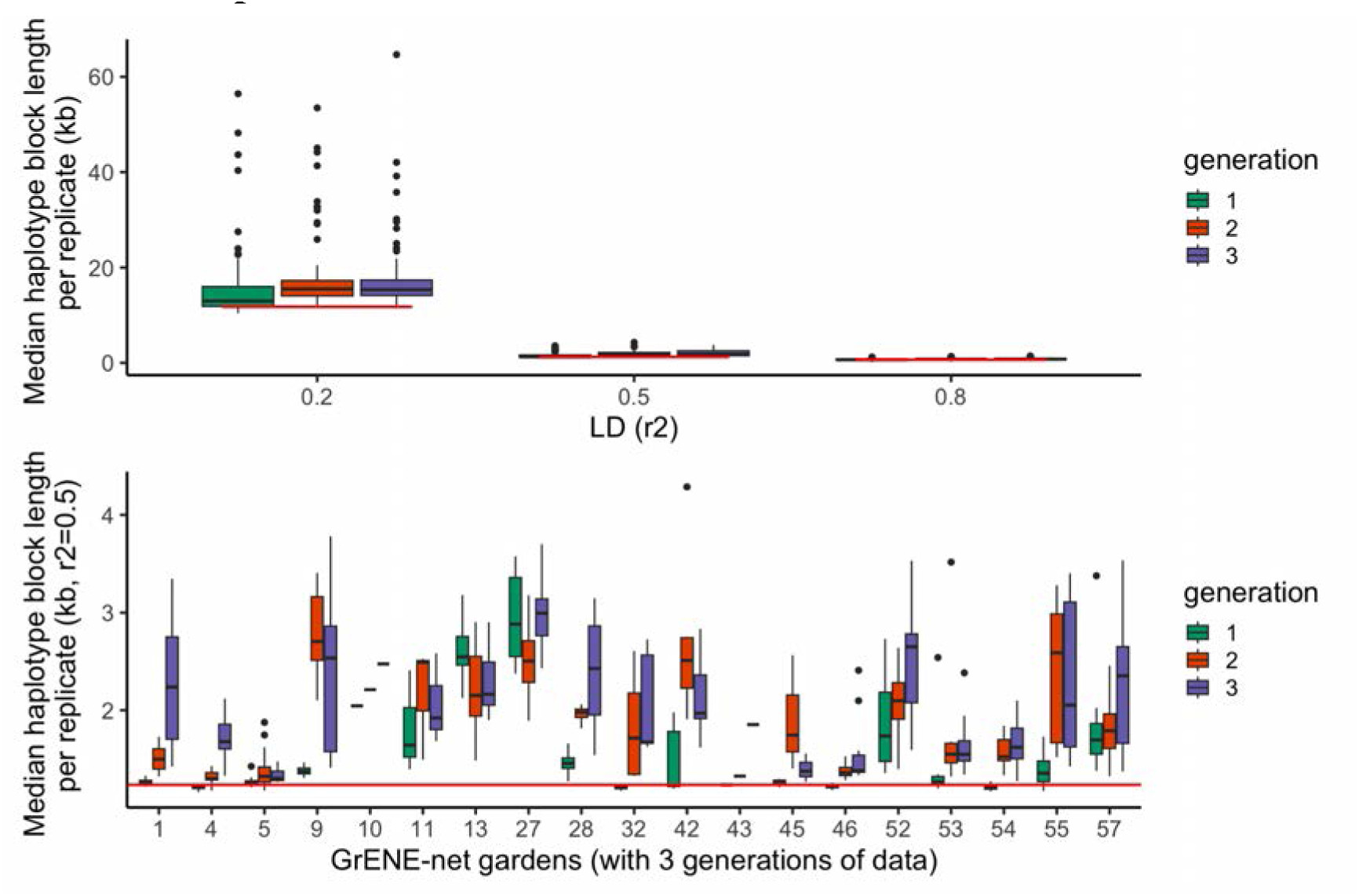
Genome-wide LD patterns across three generations Genome-wde LD pattern in three generations in GrENE-net assuming no recombination. The upper panel shows the median LD block length at different r2 cutoffs in three generations. The lower panel shows the LD block size (r2=0.5) in experimental gardens with three generations of data in three generations. The red line indicates the median block length in the founder population at different r2 thresholds.

**Figure S19.**
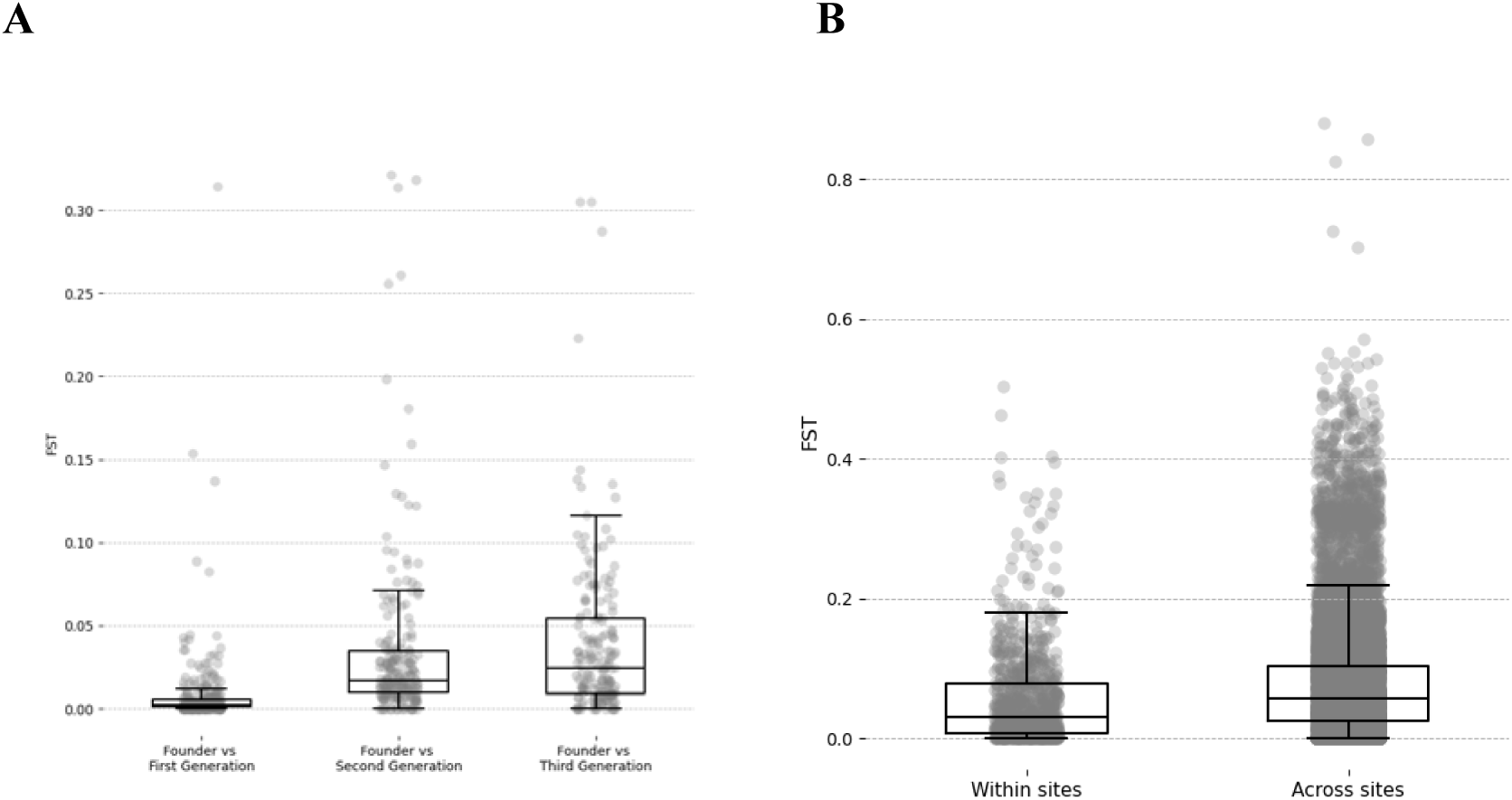
F_ST_ across generations. (A)There is an increase in F_ST_ values between the founder population and samples of the first, second and third generation (B) F_ST_ between samples within sites is significantly lower than F_ST_ between samples across sites. (Mann-Whitney U Test *P*=2×10^−90^, *n* = 50,403)

### ALLELE/ACCESSIONS RELATIVE FREQUENCY VISUALIZATIONS

**Figure S20.**
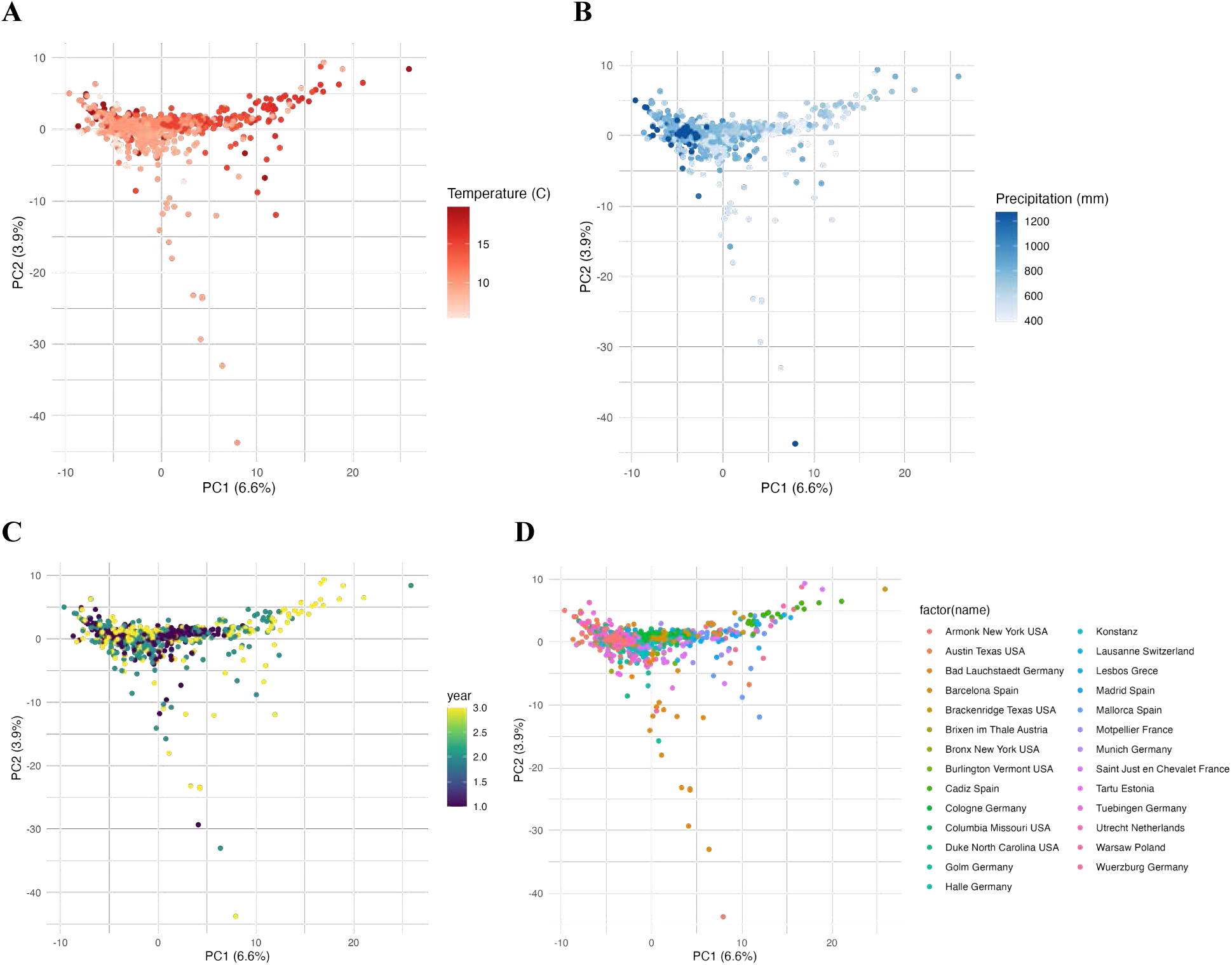
Principal Component Analysis of allele frequencies Principal Component Analysis (PCA) of genome-wide allele frequencies of LD-pruned SNPs (*n* = 13,985) colored by different variables: (**A**) Annual temperature of the experimental garden, (**B**) annual precipitation of the experimental gardens, (**C**) year of experimental sample, (**D**) experimental garden name.

**Figure S21.**
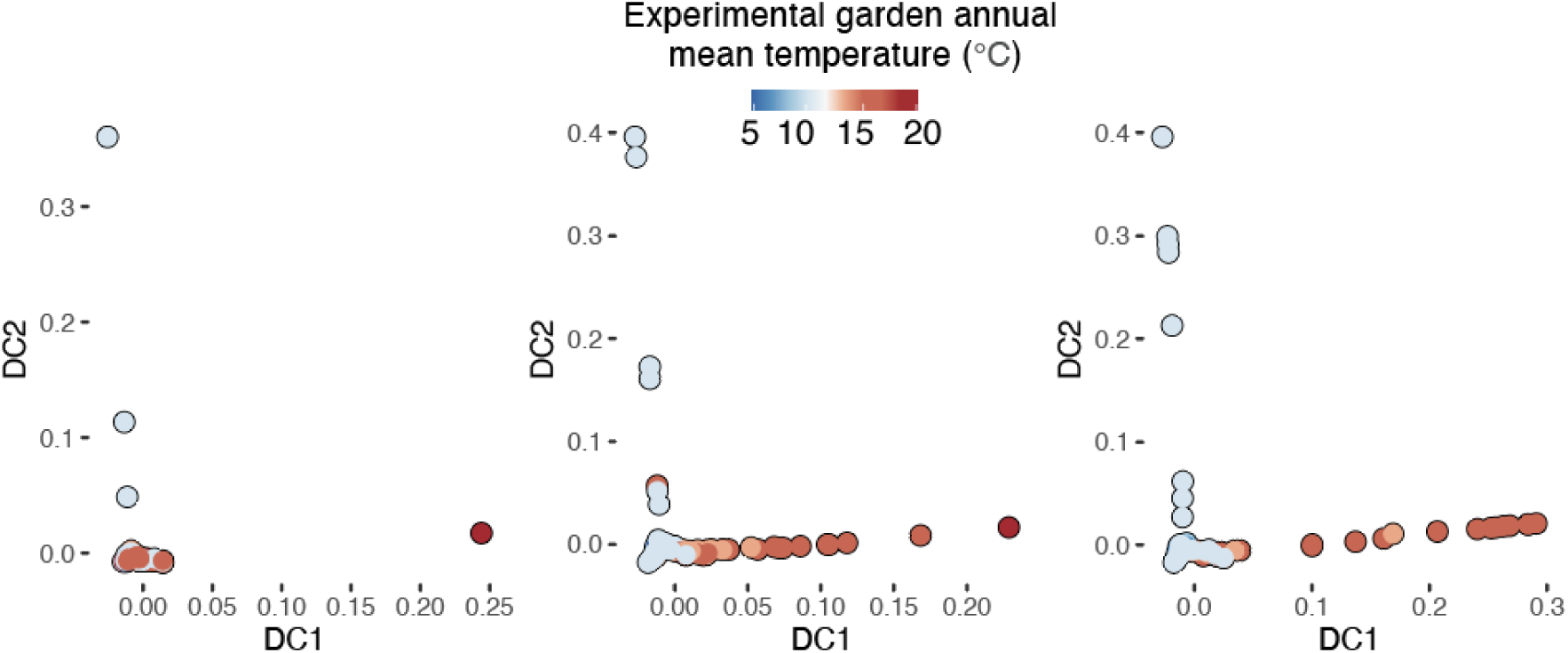
Diffusion map of allele frequencies per generation. Diffusion map of genome-wide allele frequencies of LD-pruned SNPs separated by generations and colored by the annual mean temperature of the experimental garden.

**Figure S22.**
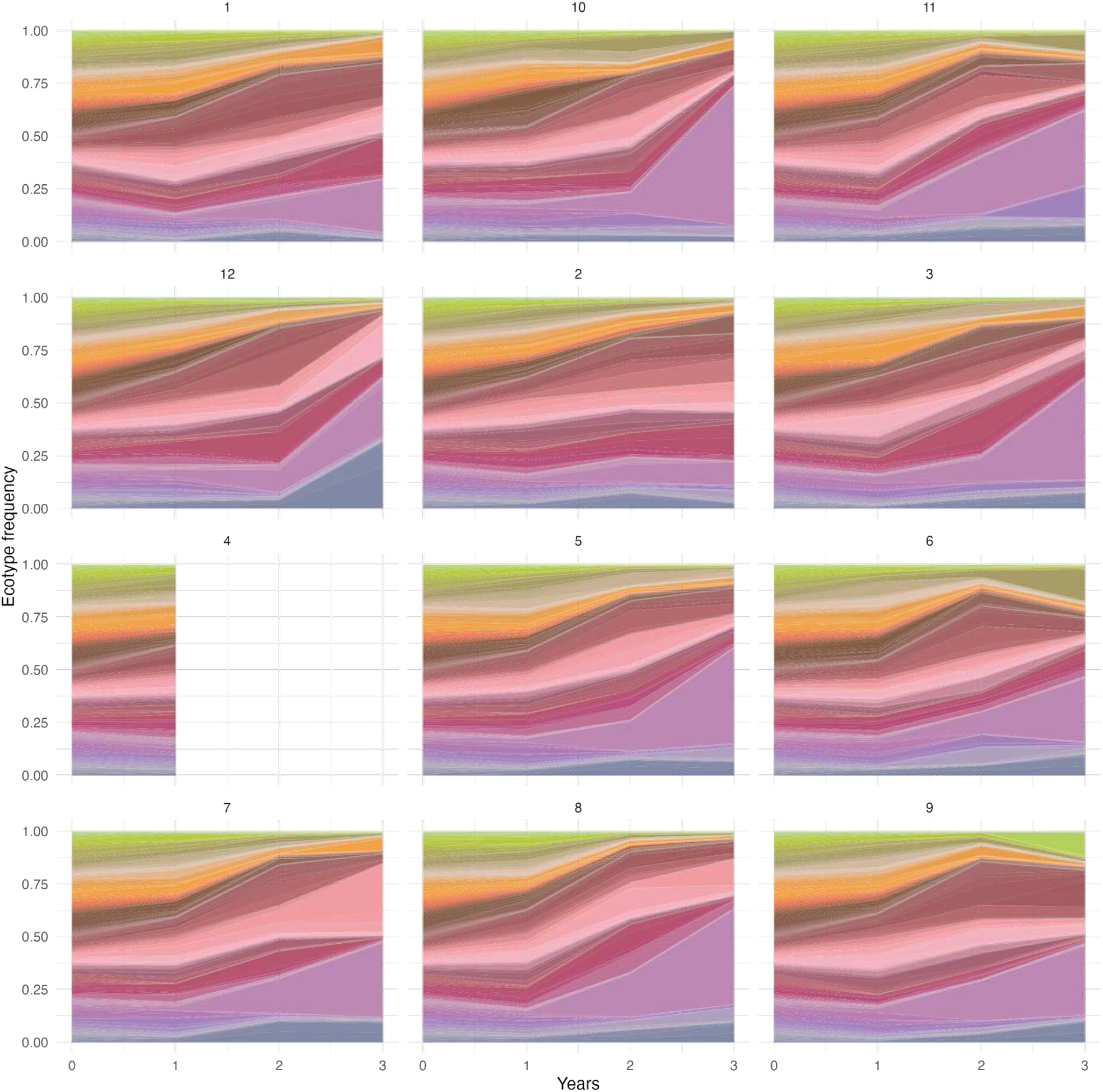
Muller plots of accessions across replicates in Cadiz, Spain. Example Muller plot of accessions trajectories in one site (#4, Cadiz, Spain) showcasing repeatability across replicates. Note how replicate 4 was lost.

**Figure S23.**
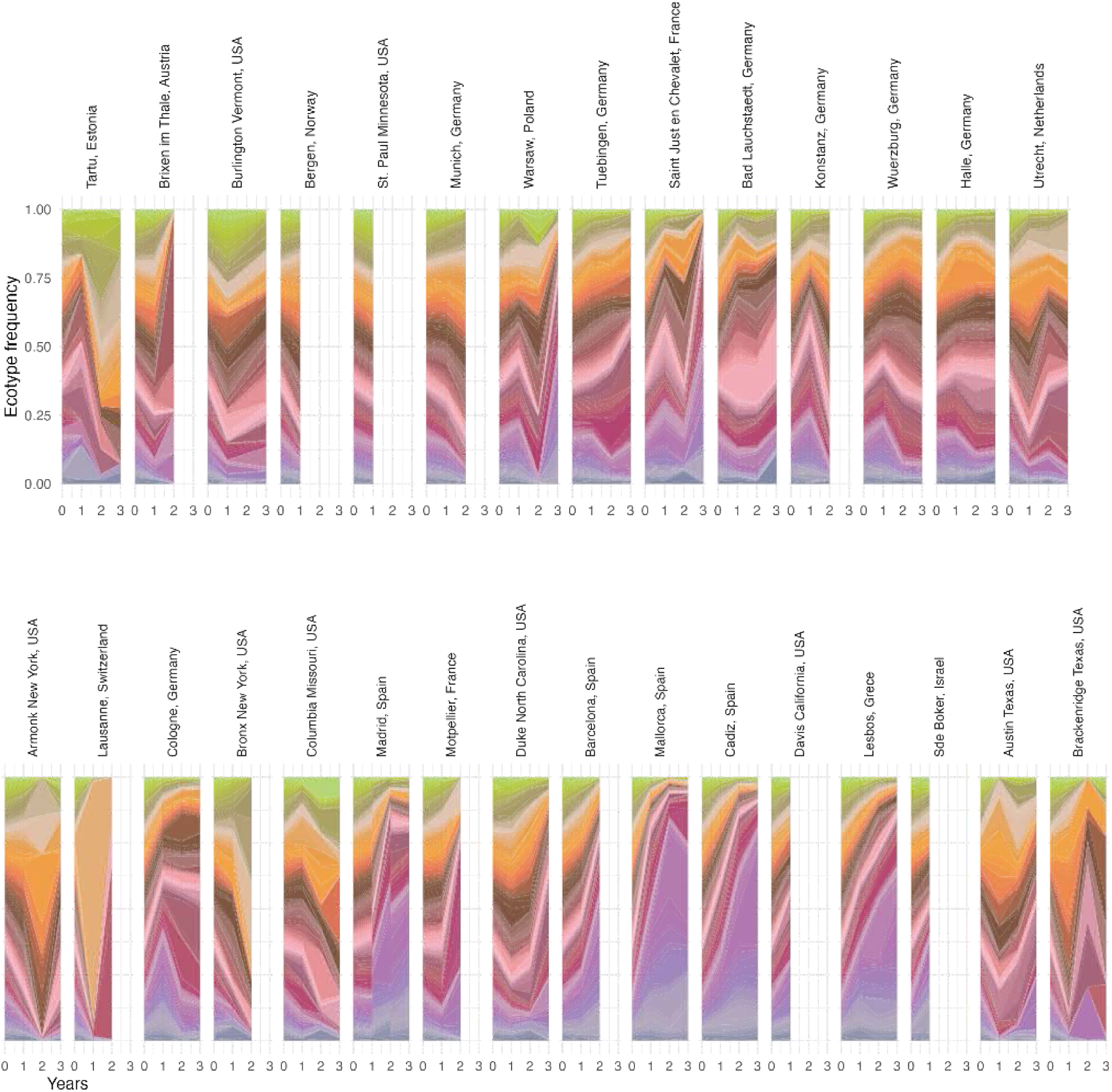
Muller plots of average accession trends across all sites Muller plot of average accession relative frequency for each site location sorting sites by annual temperature rank from cold (left) to warm (right). Colors within muller plots represent the relative proportion of accessions (*n* = 231) sorted in the y axis and color from accessions originated in cold (upper, green hues), to warm (lower, purple hues).

**Figure S24.**
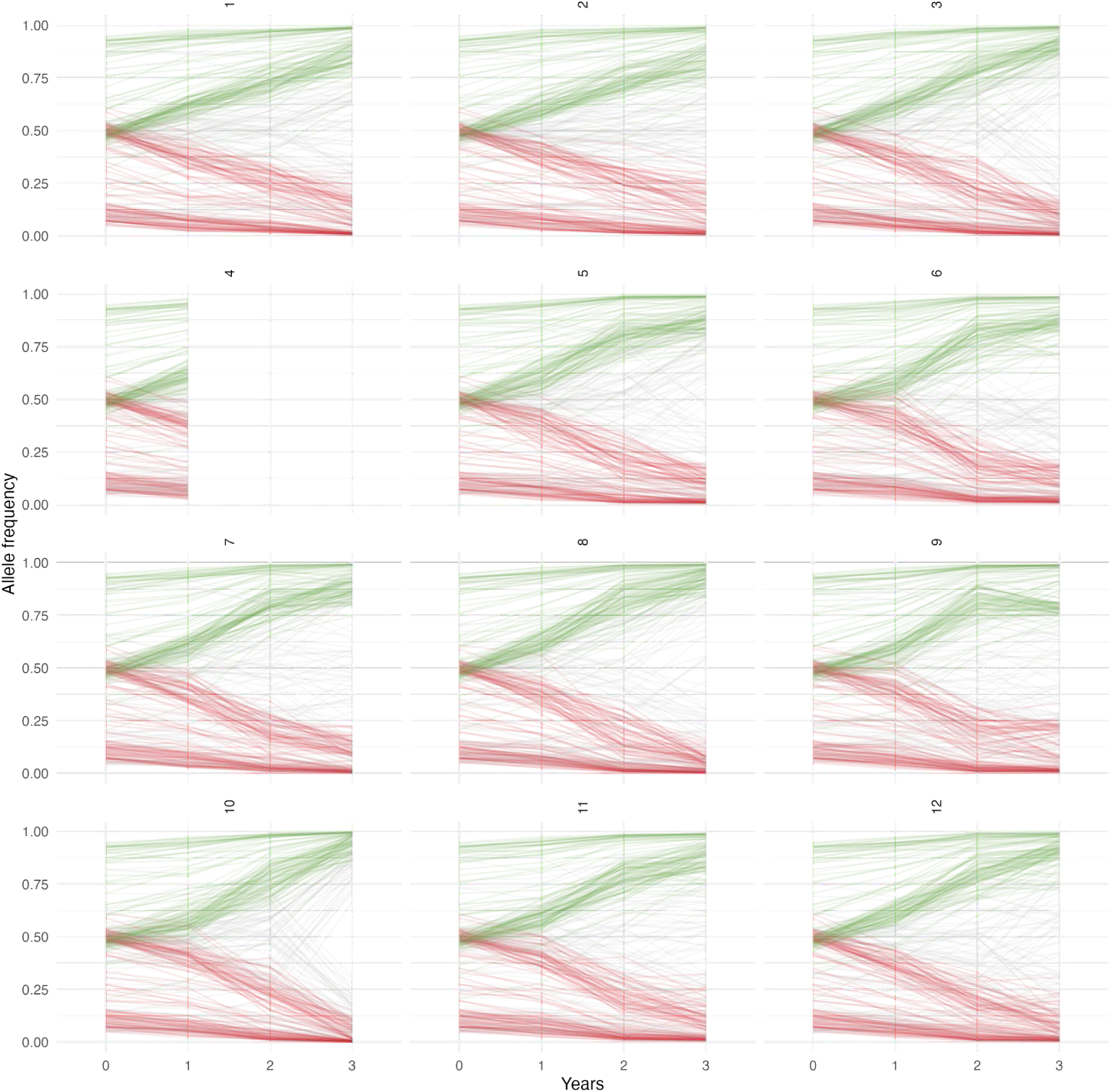
Allele trends in site #4 (Cadiz, Spain). Example allele frequency trajectories in one site (#4, Cadiz, Spain) showcasing repeatability across independent experimental population replicates. Note

**Figure S25.**
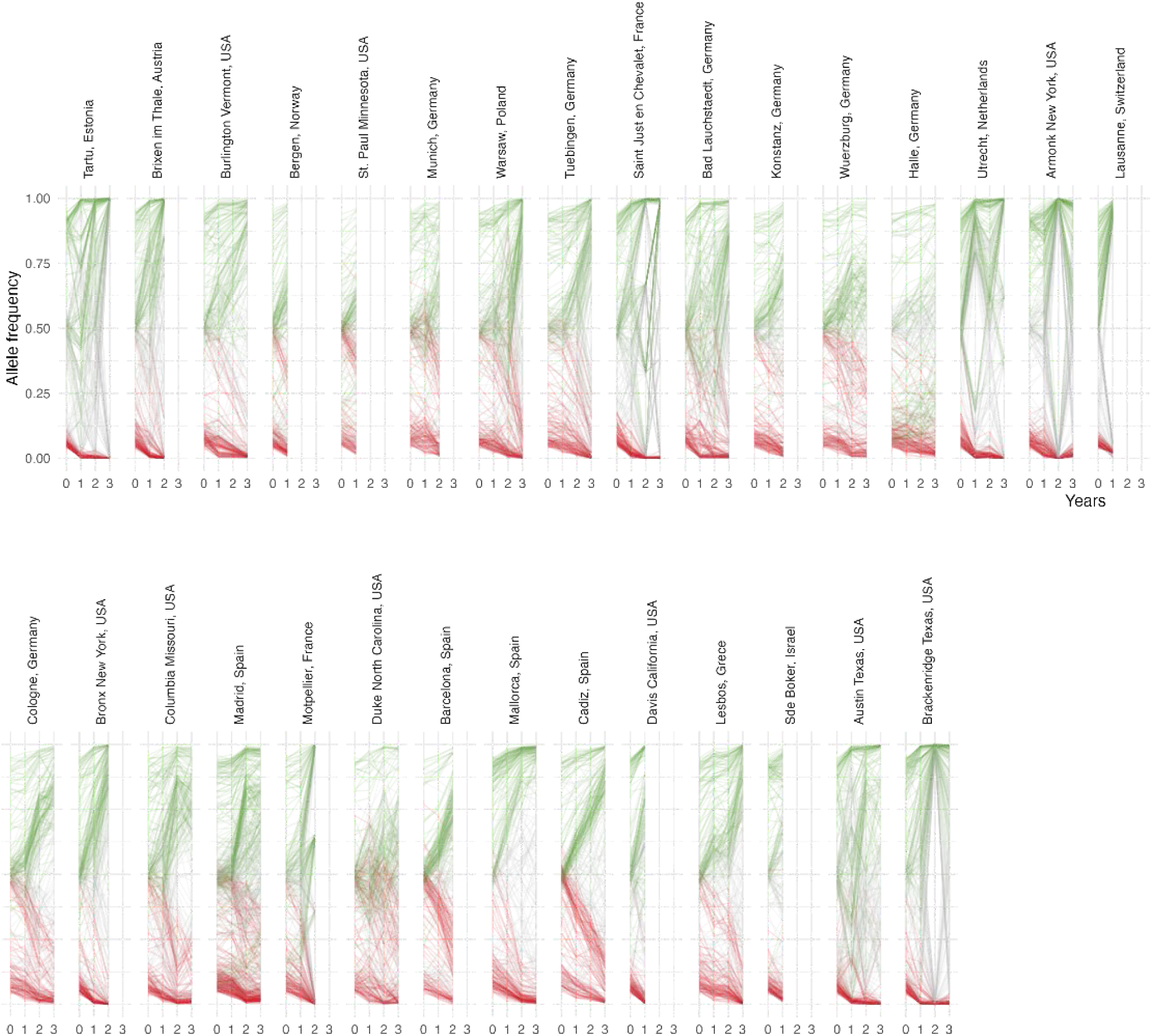
Allele trends across all sites. Displays of allele frequency changes across generations for all experimental gardens in one replicate, where the replicate chosen to be one that has the longest number of years in each site. The top 100 top LD-pruned alleles increasing (green) and decreasing (red) the most in frequency in a GLM are plotted. Sites are sorted based on annual temperature from cold (left) to warm (right).

### REPEATABILITY

**Figure S26.**
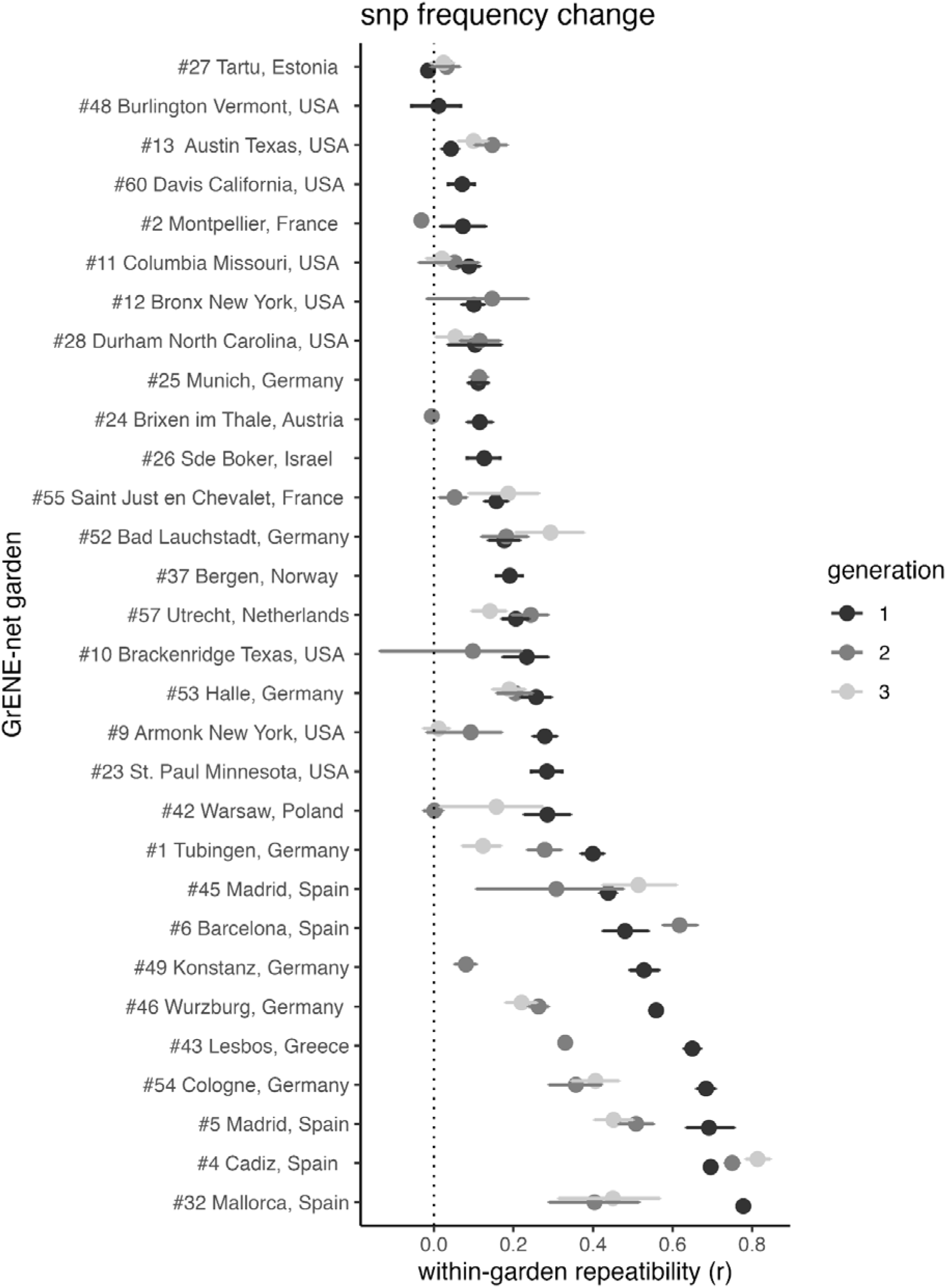
SNP-based evolutionary repeatability across experimental gardens and generations. Average pairwise correlation among 12 replicates within each experimental garden and year of SNP’s frequency change. Intervals represent 95% CI.

**Figure S27.**
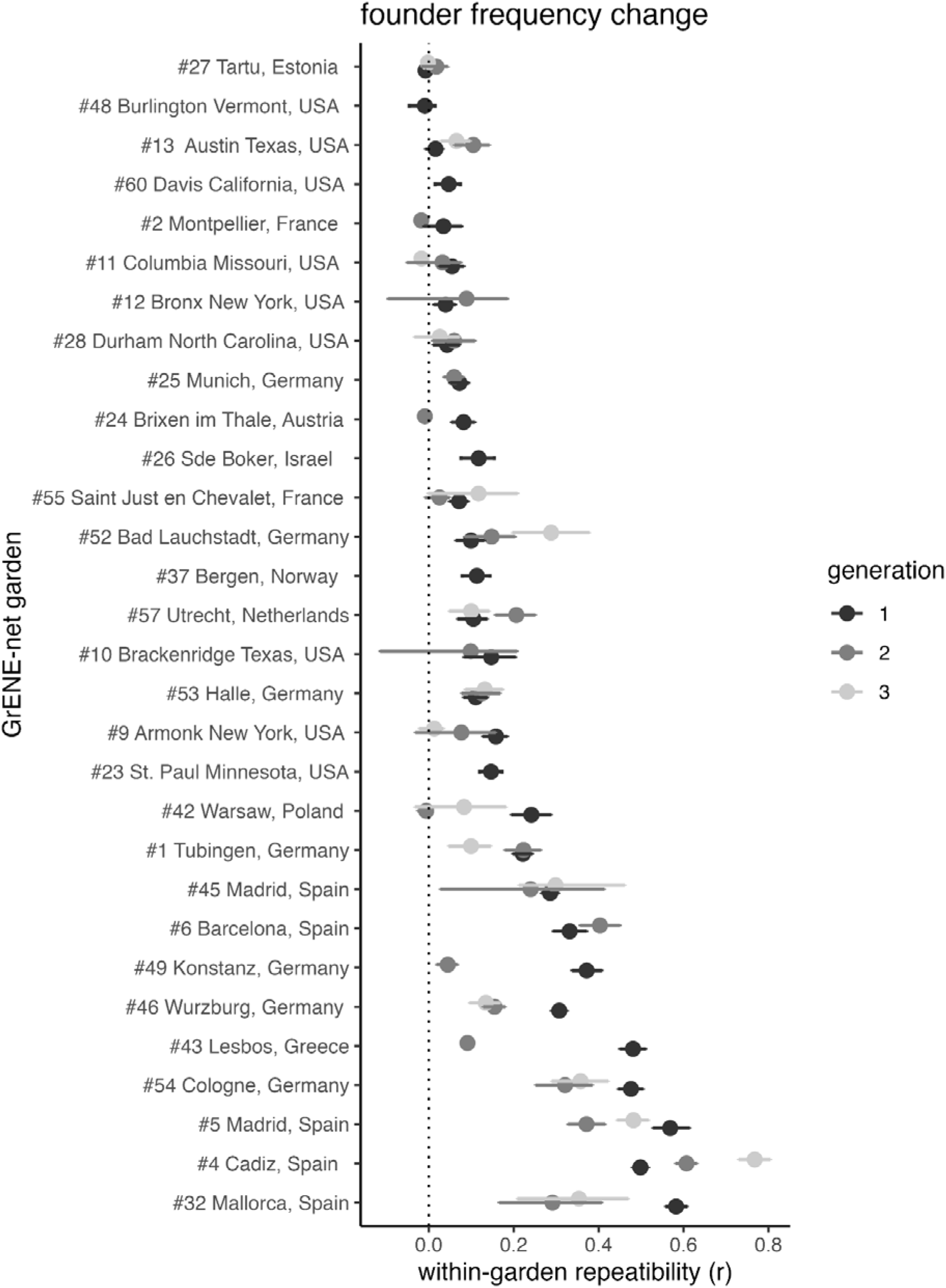
Accession-based evolutionary repeatability across experimental gardens and years. Average pairwise correlation among 12 replicates within each experimental garden and year of accessions’ change in frequency. Intervals represent 95% CI.

**Figure S28.**
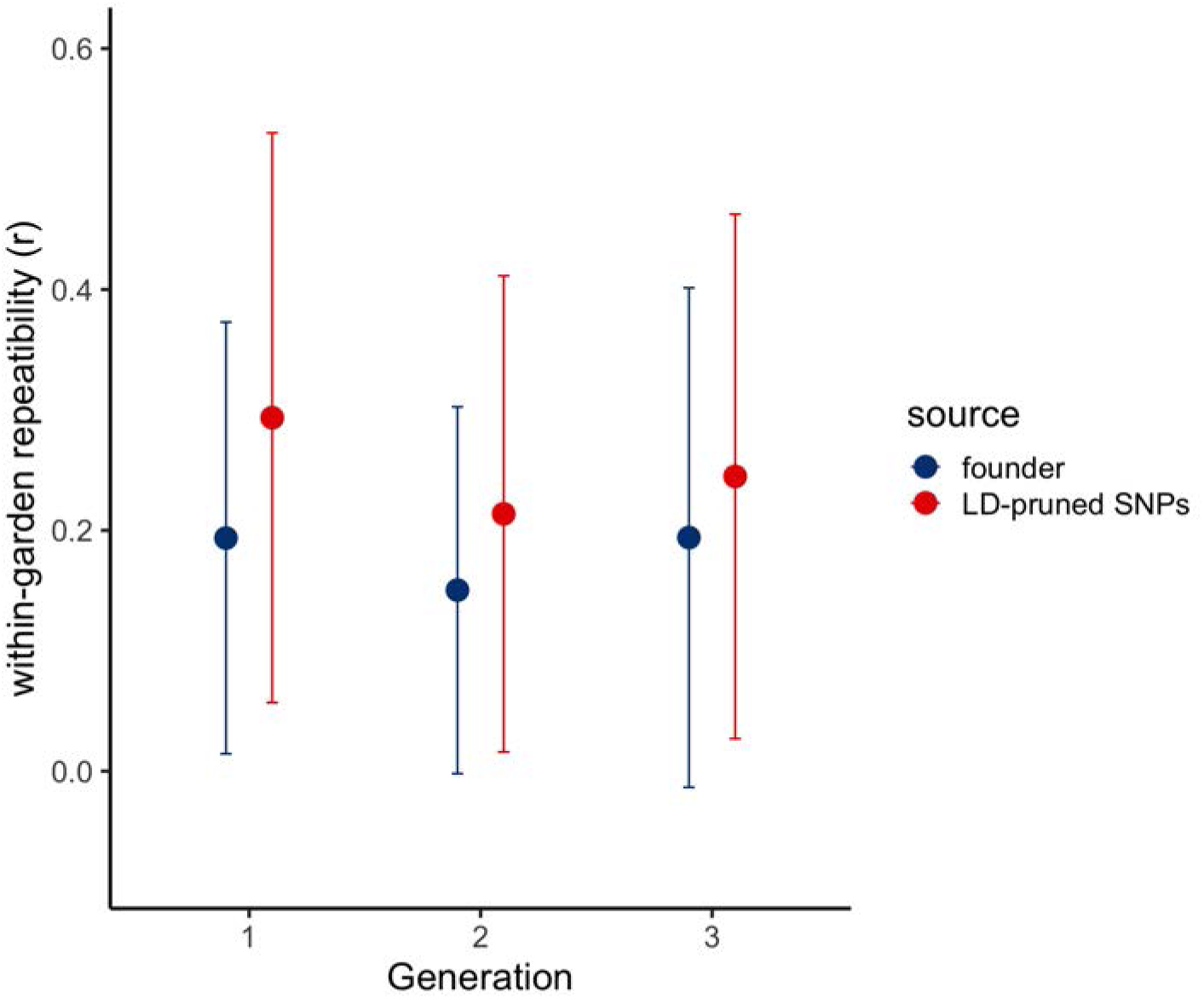
Evolutionary repeatability of all experimental gardens across generations. The dark blue indicates repeatability calculated from founder frequency changes, and red color indicates repeatability calculated from LD-pruned SNP frequency changes

**Figure S29.**
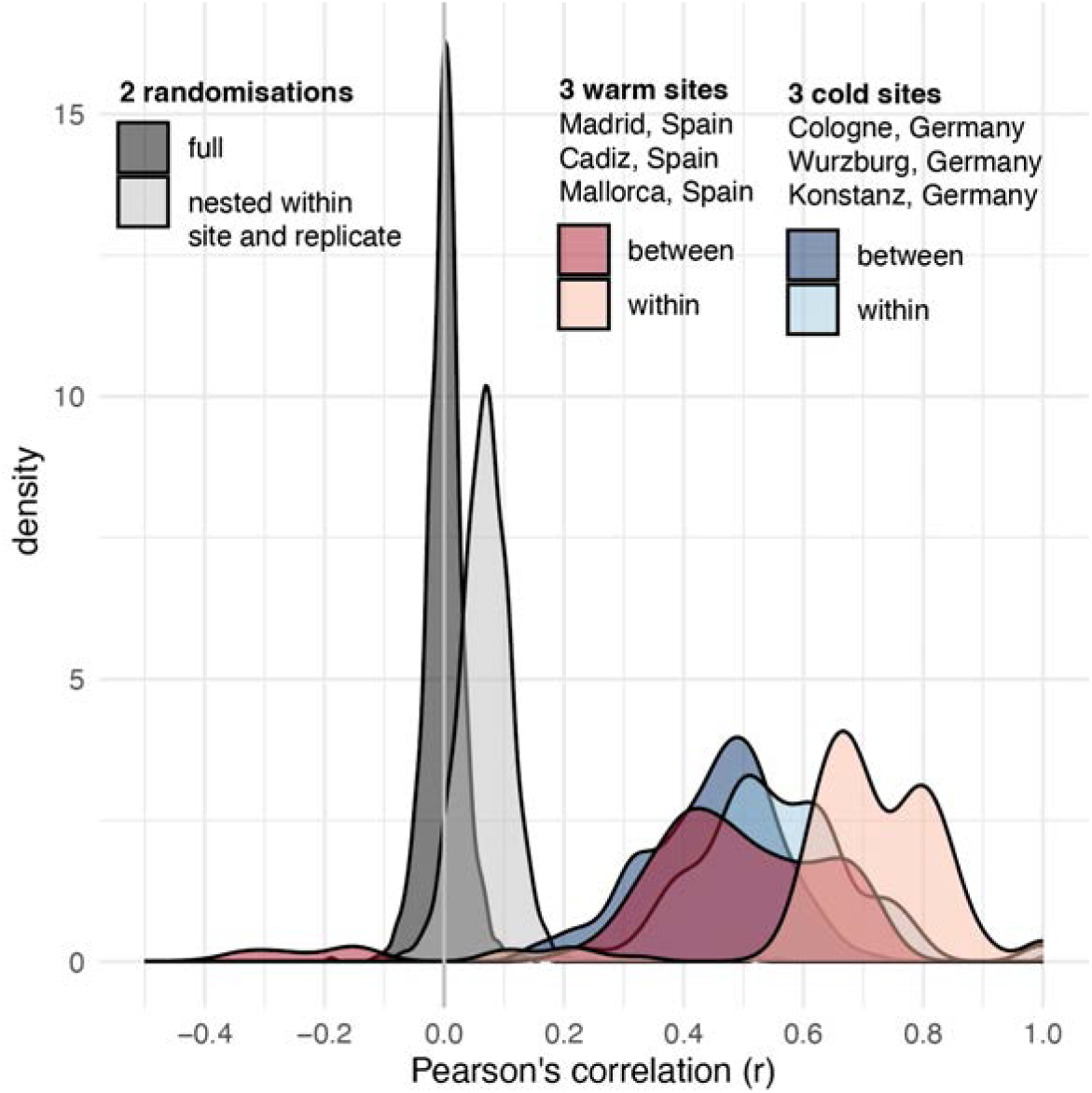
Evolution parallelism across different experimental gardens. This figure illustrates the distribution of Pearson’s correlation coefficients (r) for accession frequency changes between replicates within the same site (within-site comparisons) and between replicates from different sites with similar climates (between-site comparisons). The analysis was conducted using an example set of three warm location site gardens (Madrid, Cádiz, and Mallorca, Spain) and three cold location site gardens (Cologne, Würzburg, and Konstanz, Germany).

**Figure S30.**
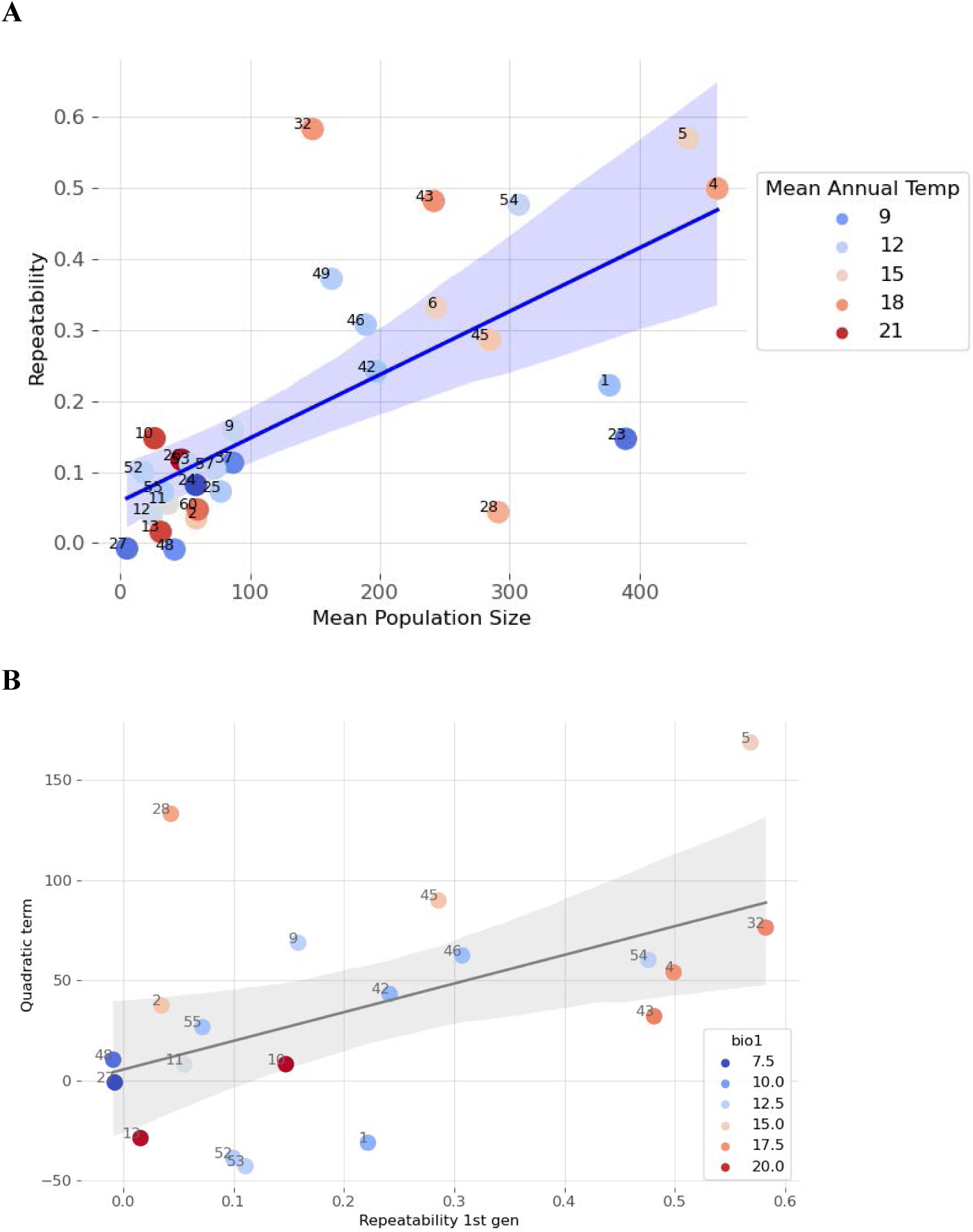
Relationship of evolutionary repeatability and population rescue. (A) There is a positive and significant relationship between mean population size and evolutionary repeatability in generation 1, as measured by pairwise correlations among accession changes within sites (*R^2^* = 0.476, *P* < 2.46×10^−5^). (B) There is a positive and significant relationship between the quadratic term estimated based on population size dynamics and evolutionary repeatability, as measured by pairwise correlations among accession changes within sites (*R^2^* = 0.270, *P* < 1.86×10^−2^,).

**Figure S31.**
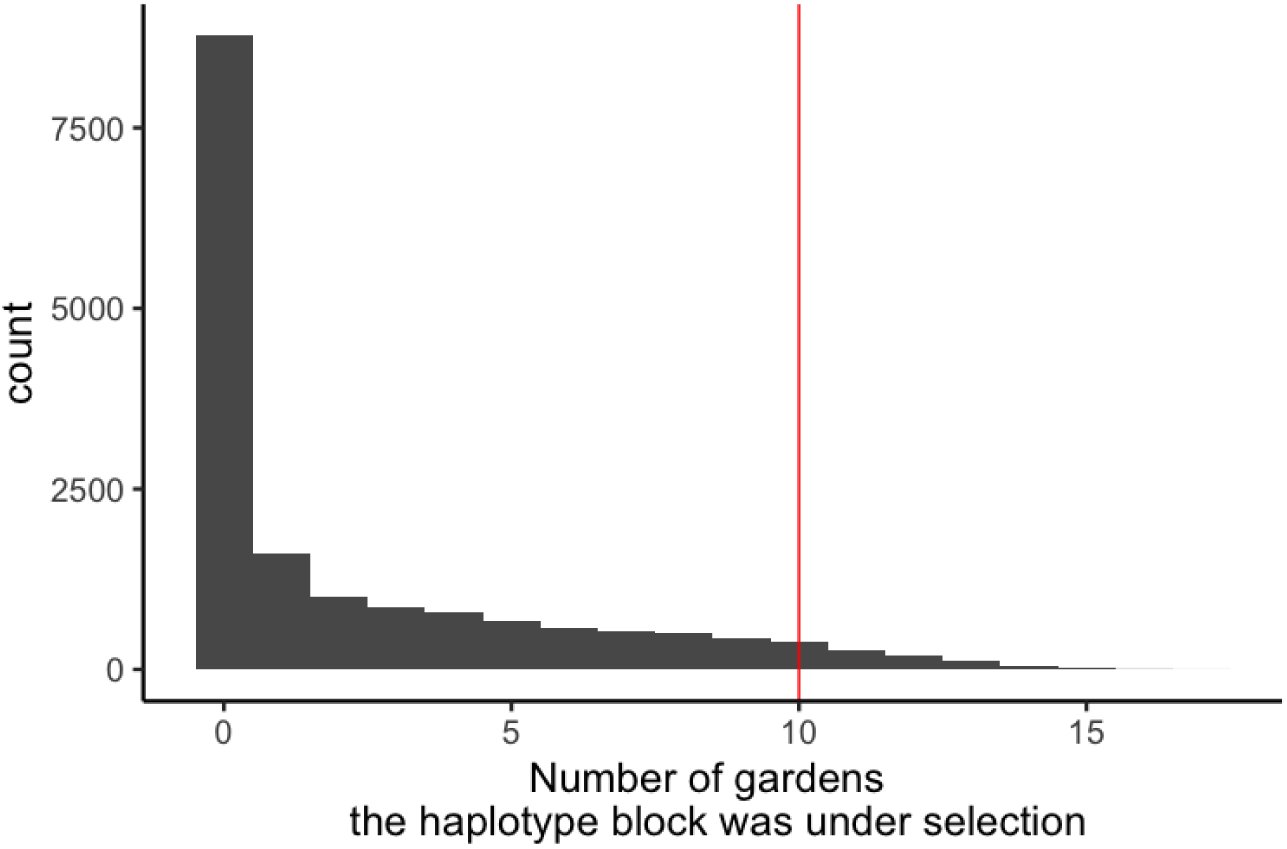
Overlap of natural selection signal across gardens. After collapsing the SNP-based selection signals per garden into 16,917 LD blocks, we plotted the distribution of LD blocks that were significant in one or more gardens. By conducting a permutation analysis, we established that it would be very unlikely to identify a block in 10 or more gardens only by chance (*P* = 10^−6^).

### STABILIZING SELECTION

**Figure S32.**
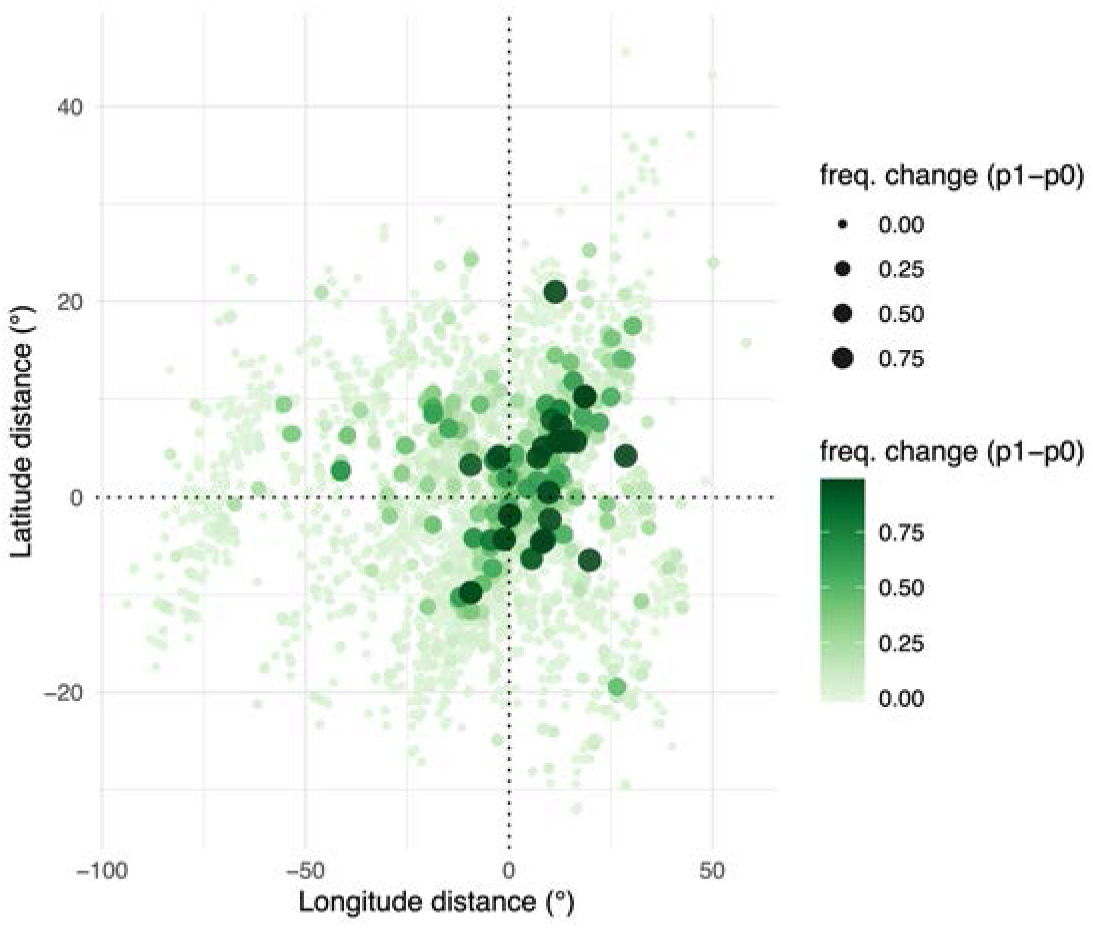
Local adaptation based on geographic distance of accessions to gardens. Climate distance of accession-to-garden is measured in degrees latitude or longitude across all Eurasian experimental gardens (*n* = 22) and all pairwise accession-garden-year combinations (*n* = 129,822).

**Figure S33.**
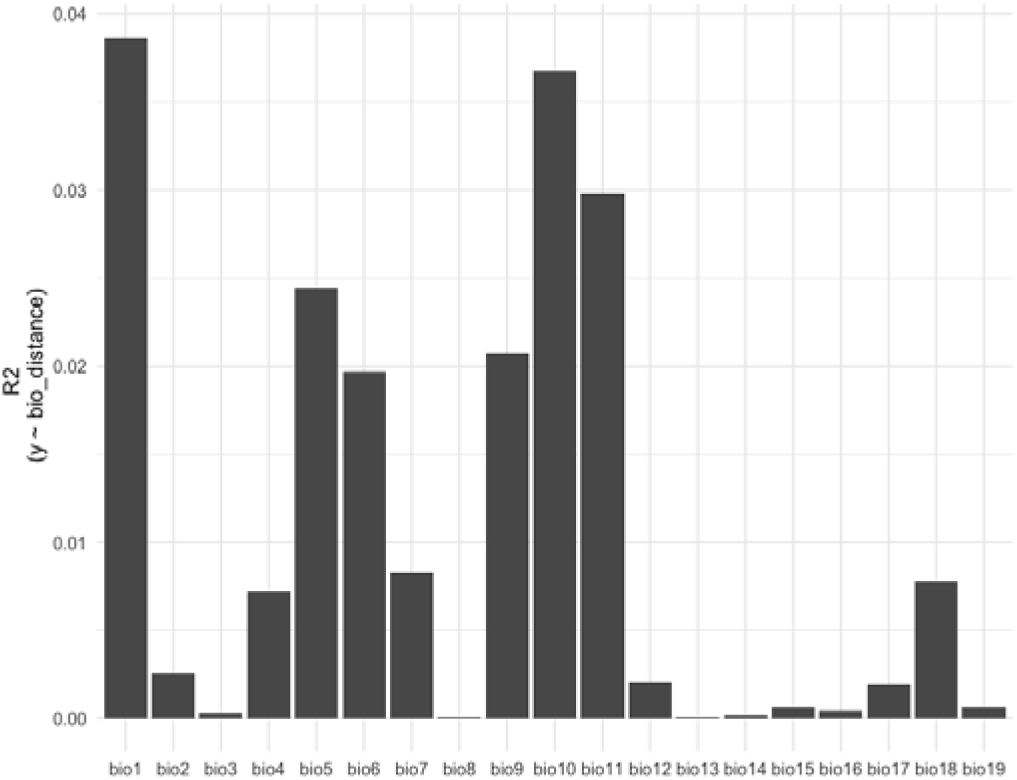
Stabilizing selection per BIOCLIM variables. The variance explained by the quadratic term of different BIOCLIM variables in the stabilizing selection model.

**Figure S34.**
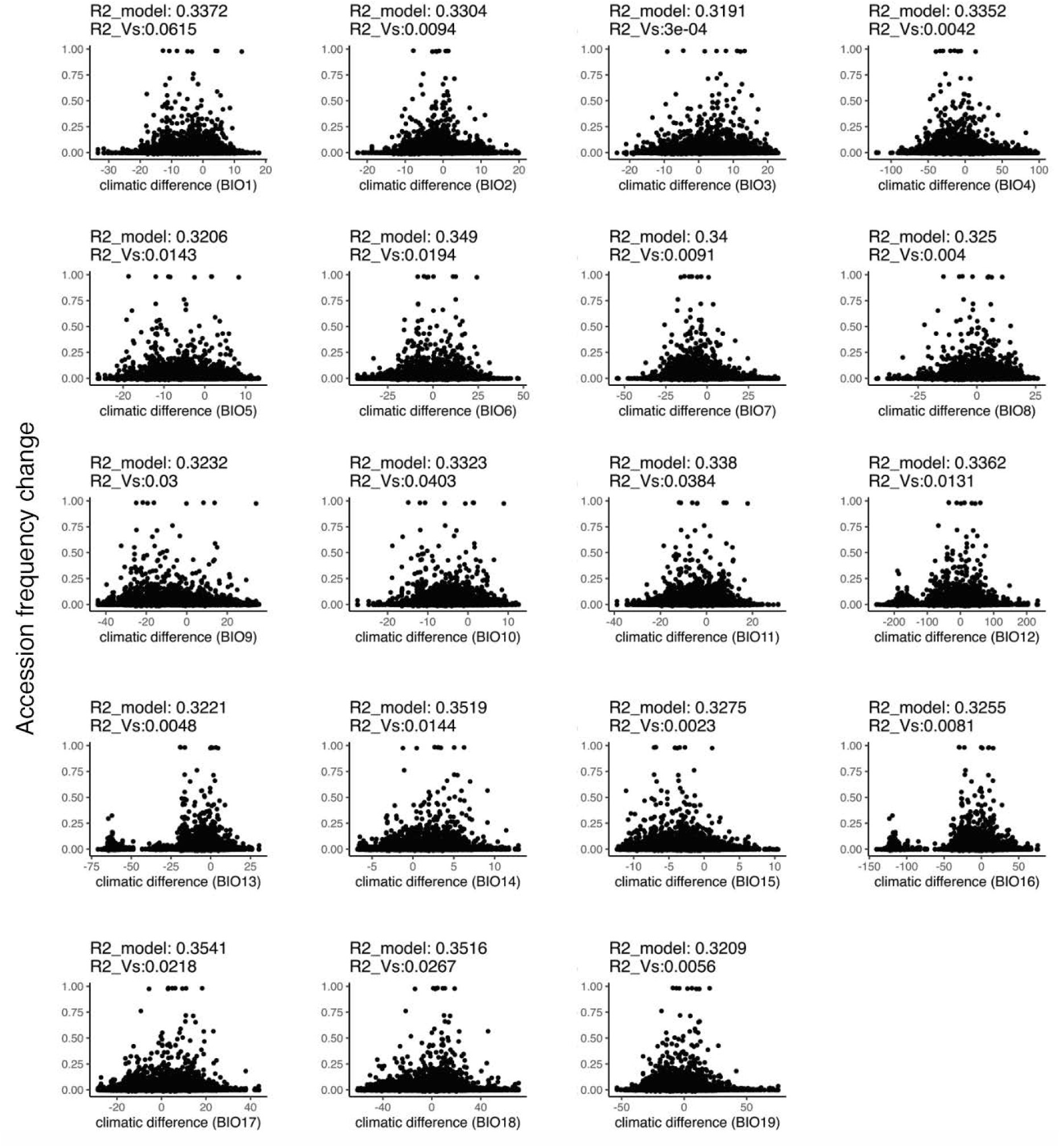
Gaussian stabilizing selection per BIOCLIM variables. The Gaussian stabilizing selection models for each BIOCLIM variable where y-axis represents accession frequency change in year 1, and x-axis represents climate distance (accession - garden). The value of *R^2^* model indicates the total variance explained by the model, and the value of *R^2^*of *Vs* indicates the proportion of variance explained by a simple (*z-z_opt_*)*^2^* / *Vs*.

**Figure S35.**
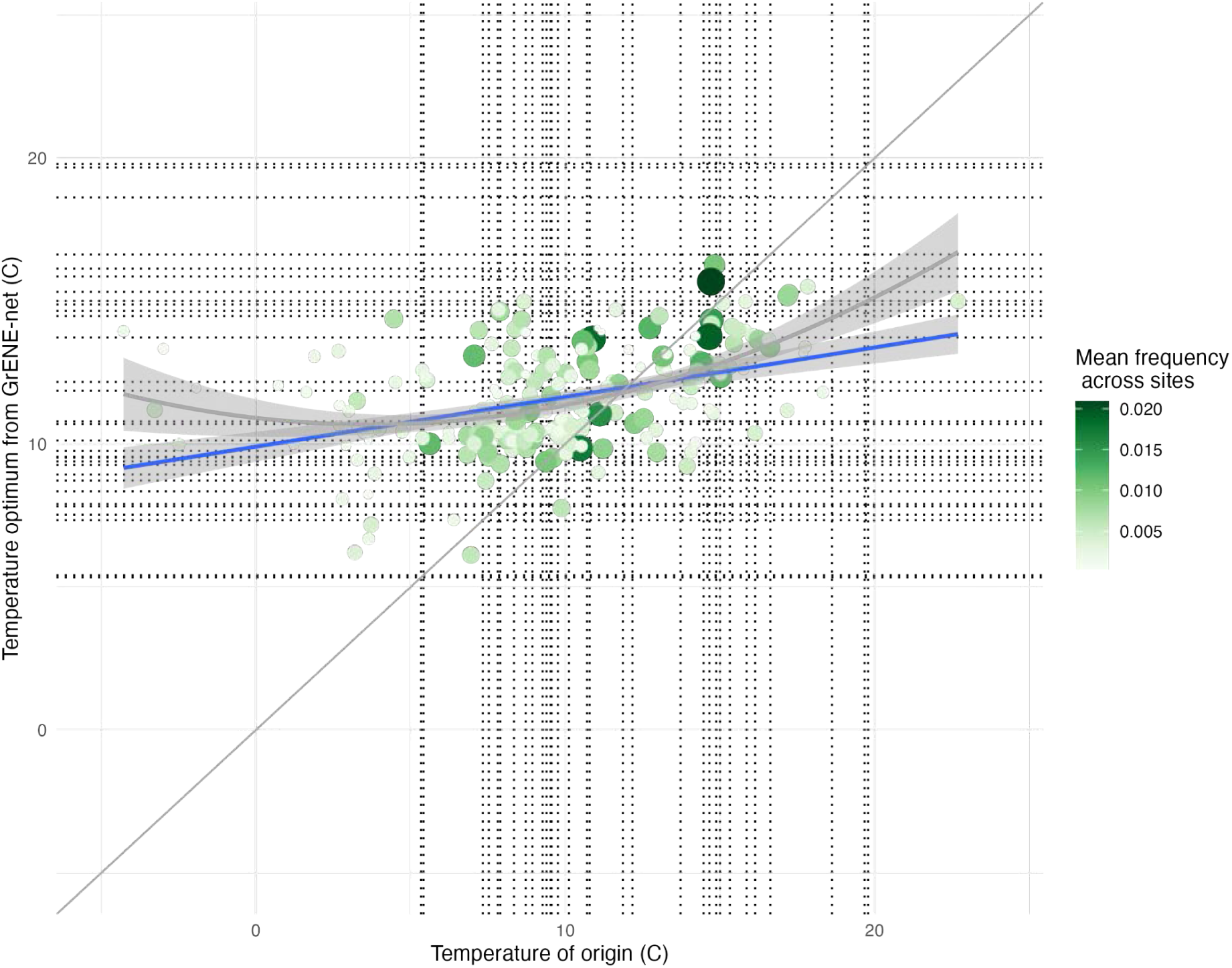
Accessions’ climate of origin vs inferred climate optimum. Accession climate of origin (x-axis) is correlated with a calculation of accessions’ climate optimum as inferred from their relative success across experimental gardens based on a weighted average of the experimental climate, *e_i_*, and accessions relative frequency *p_i_* to estimate a weighted environment average: *e_i_ × p_i_ /Σp_i_*. Vertical dotted lines mark the temperatures where field experiments are conducted (*n* = 30).

**Figure S36.**
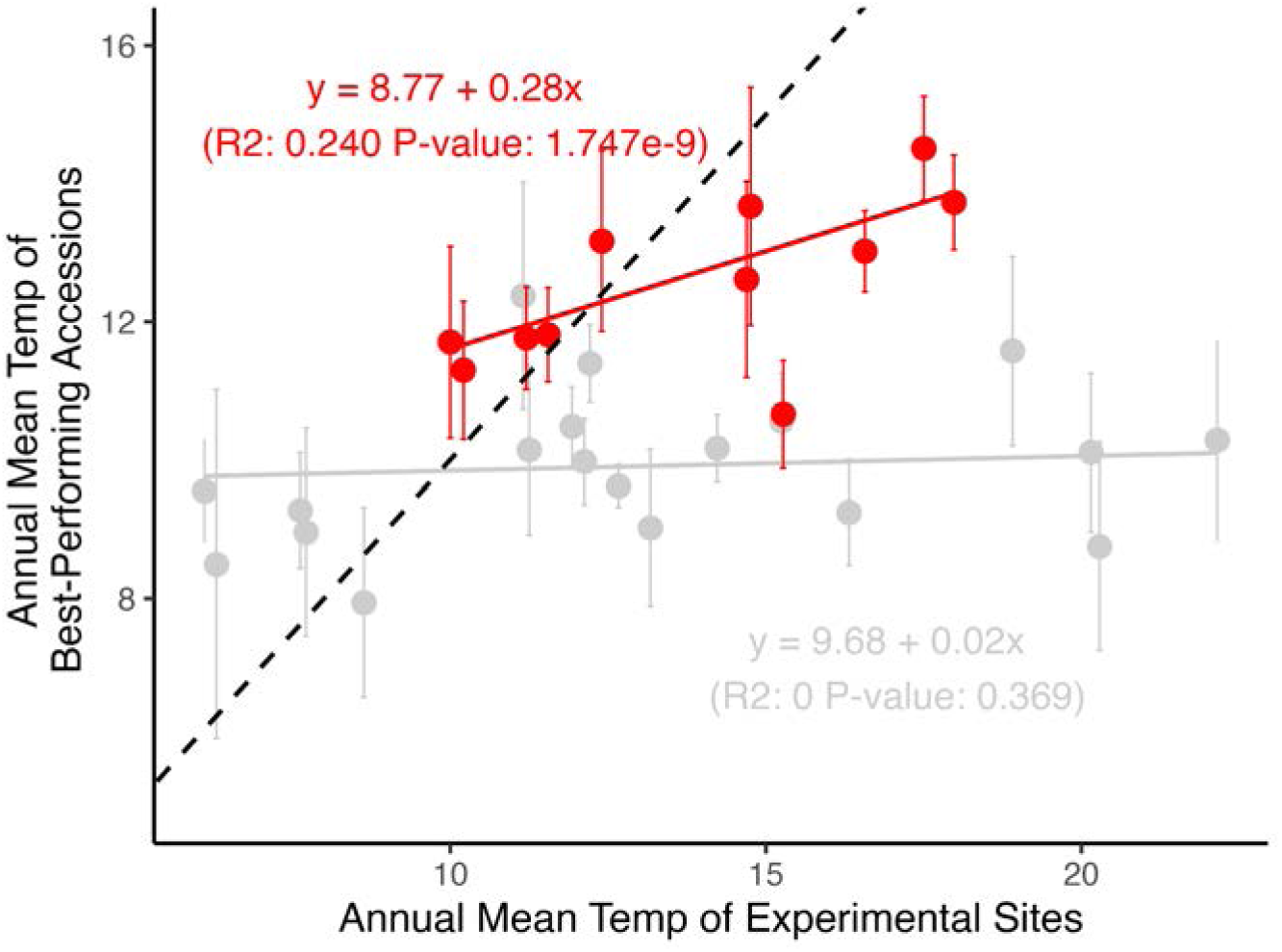
Top accessions’s climate of origin vs garden temperature. The red color indicates experimental gardens with SNP-based repeatability (Pearson’s correlation) *r >* 0.2, and grey color indicates experimental gardens with SNP-based repeatability *r <* 0.2. The dotted line indicates the y=x line.

**Figure S37.**
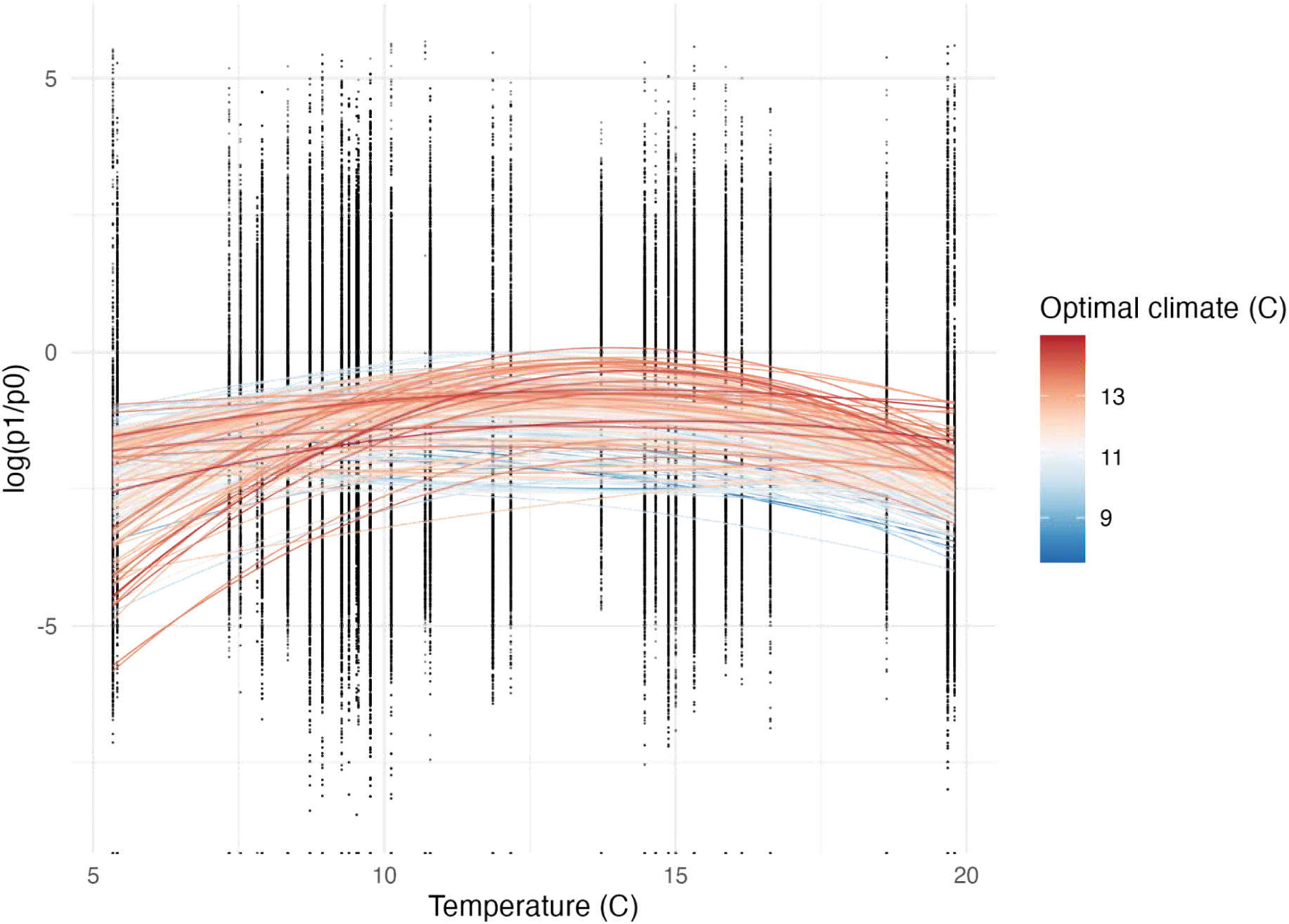
Accession relative frequency changes across experimental environments. To visualize a model-free local adaptation signal, we visualize accession relative frequency change in the first generation of experiments [log (p_t+1_ / p_t_)] (y-axis) is visualized against the accessions’ climate of origin (annual temperature at collection location) (x-axis). Dots indicate accession relative frequency observation (*n* = 30 experimental locations). Lines indicate a quadratic regression fit by accession. Line colors indicate the accession climate temperature of origin which helps visualize that warm originating accessions tend to increase in frequency in warmer experimental locations, and vice versa.

**Figure S38.**
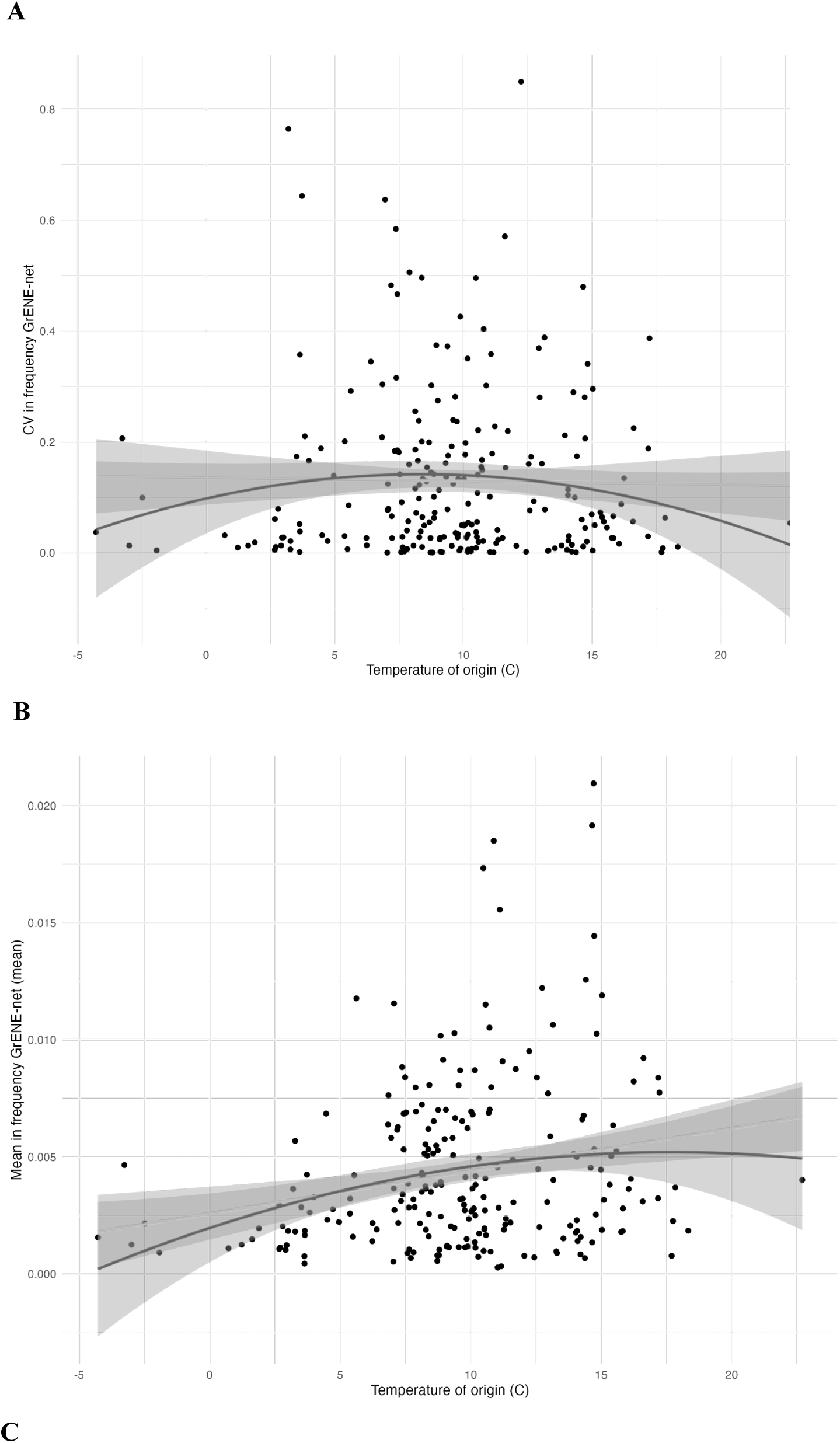
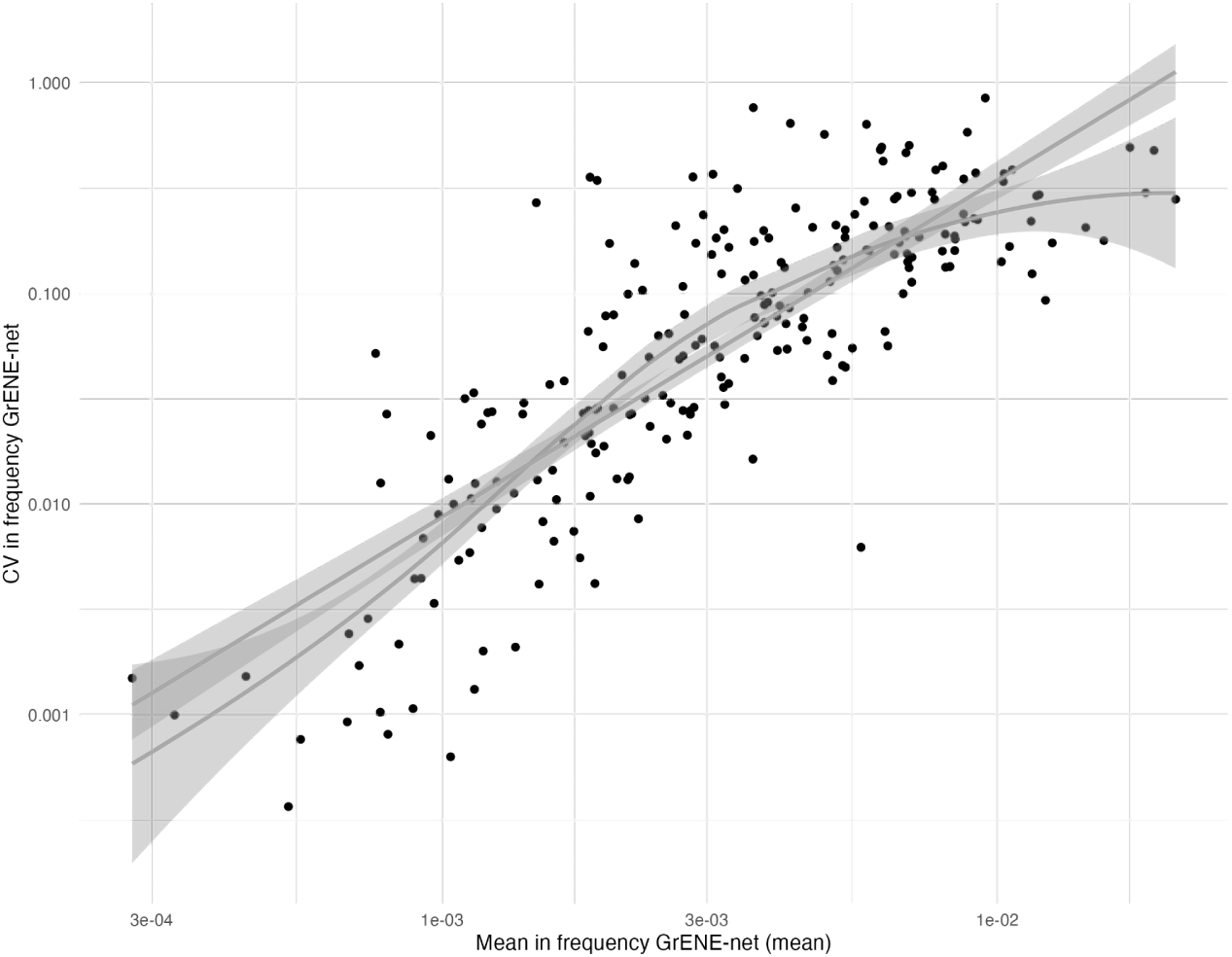
Summary statistics of accession relative frequency across experiments. To understand general patterns of accession relative frequency across experimental gardens. (**A**) Coefficient of Variation (CV) of accession relative frequency across all sites regressed against the accessions’ climate of origin. (**B**) Mean accession relative frequency across all sites positively correlates with the accessions’ temperature of origin, potentially indicating warm adapted accessions generally are more successful on average across experiments. (**C**) Positive relationship between the mean frequency change of an accession and the coefficient of variance, which indicates the most successful accessions on average are also more variable across environments.

**Figure S39.**
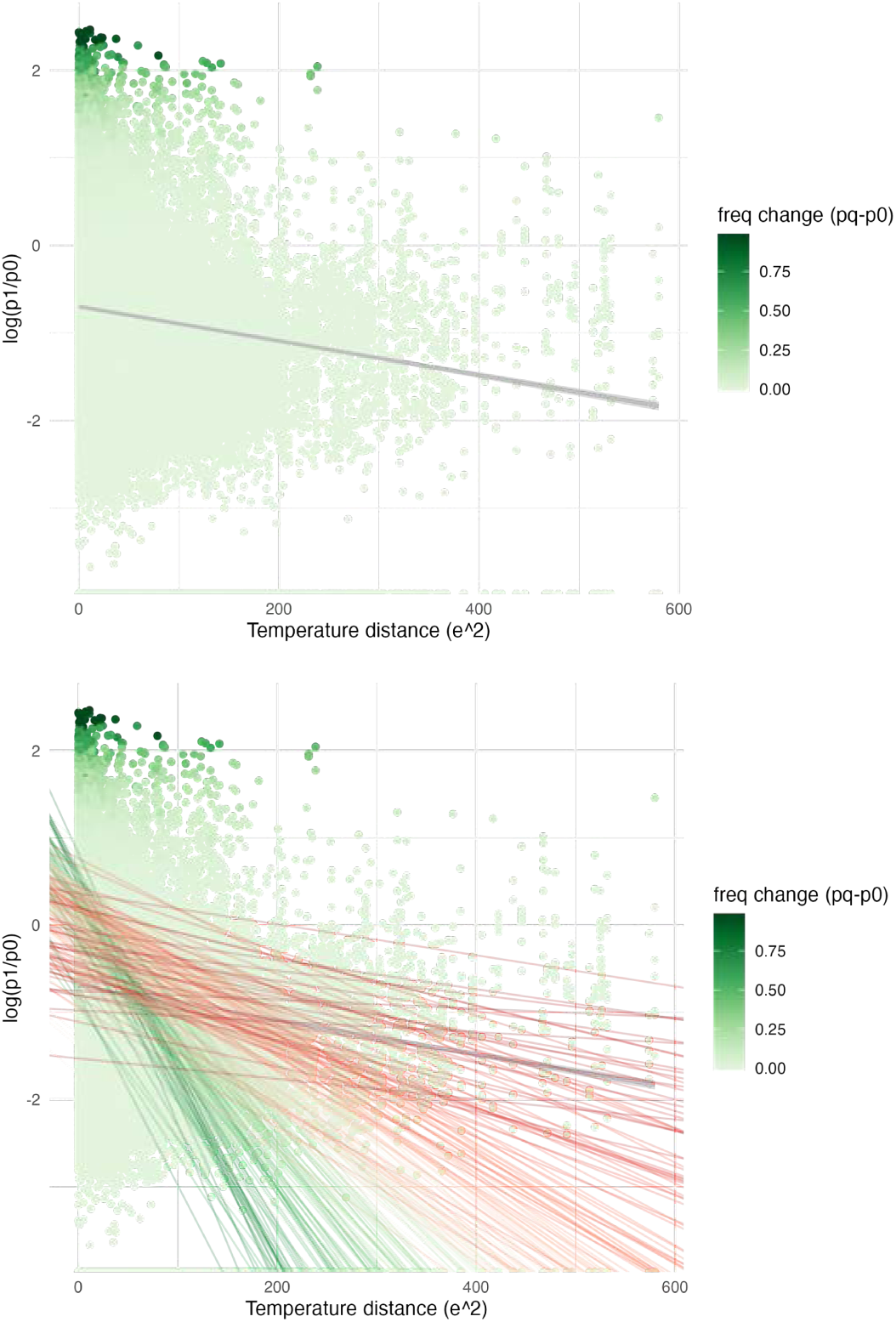
Stabilizing selection fitting with temperature by accession. (**A**) Stabilizing natural selection model fitting *log(p_t+1_ / p_t_)* = *log(W_max_ / ŵ) - V_s_^−1^ (z_origin_ - z_garden_)^2^* with a single *V_s_*regression. (**B**) Example of fitting a random slope MCMCglmm that can vary across accessions indicating accessions with shallow (red) to steep (dark green) slopes.

**Figure S40.**
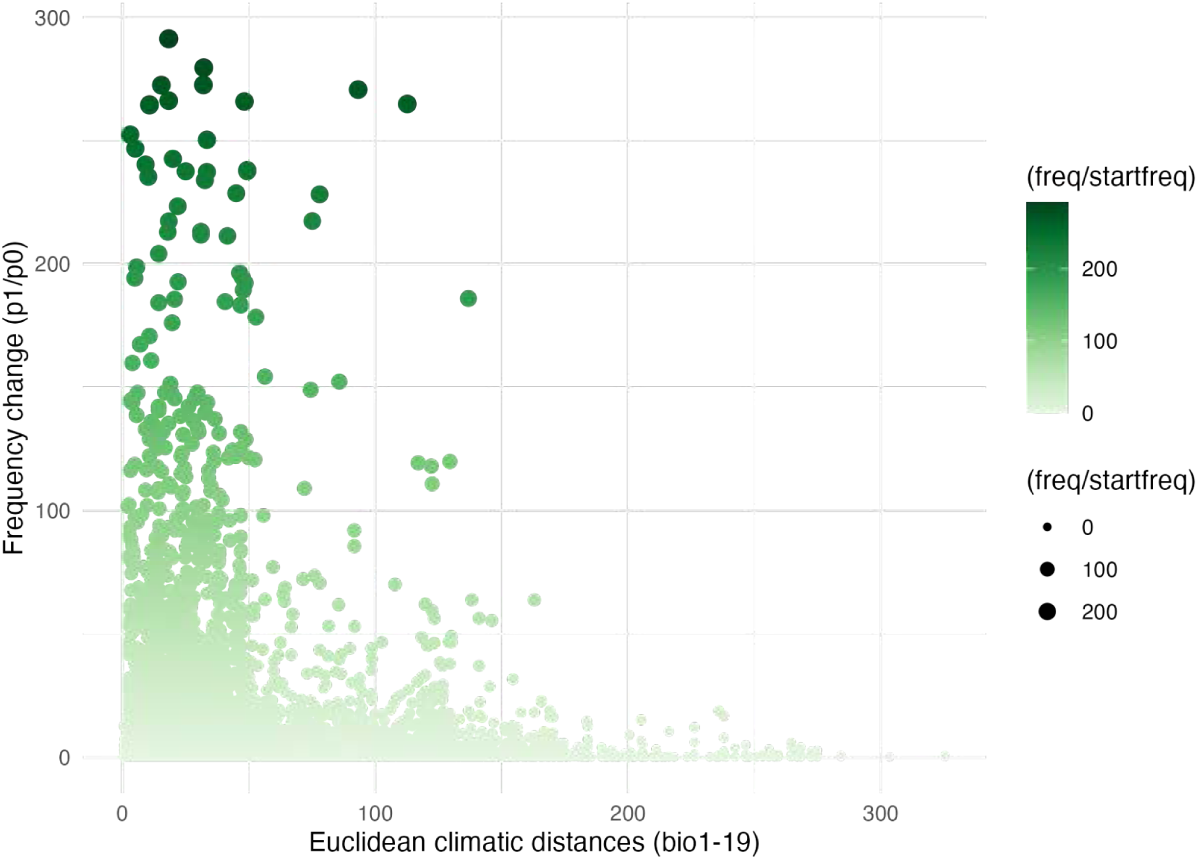
Stabilizing selection signal on euclidean distance of 19 climatic variables Stabilizing selection signal using accession-to-garden Euclidean distances of 19 variables rather than a single variable showcases the resilience of the stabilizing selection under a multivariate climate space.

**Figure S41.**
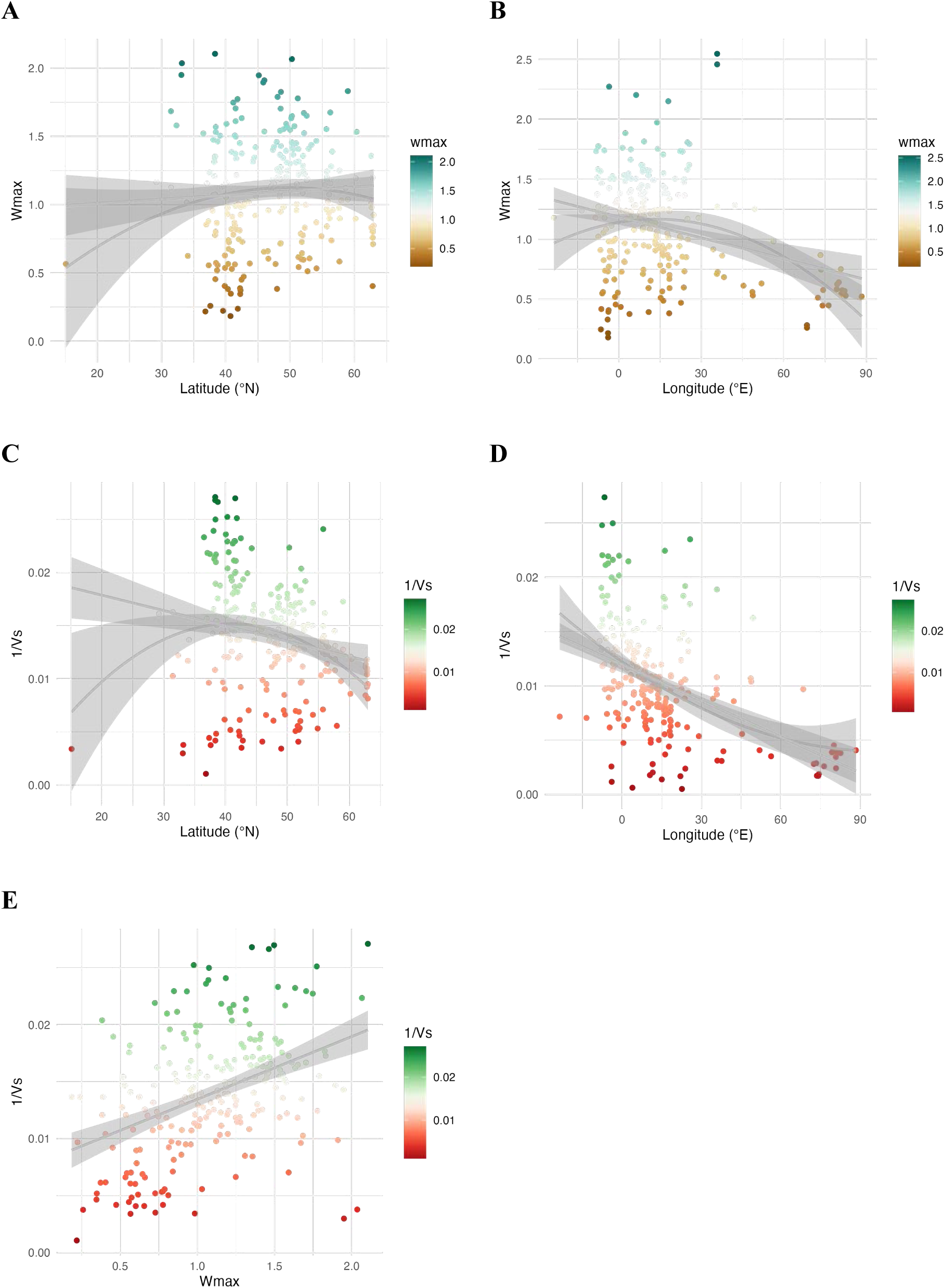
Stabilizing selection parameters per accession across geography. (**A**) Maximum relative fitness peak at optimum *W_max_* by accession across a latitudinal and (**B**) longitudinal gradient. (**C**) Fitness decline with distance parameter, *V ^−1^,* by accession along a latitudinal and (**D**) longitude gradient. (**E**) Relationship between *W_max_* and *V_s_^−1^*.

**Figure S42.**
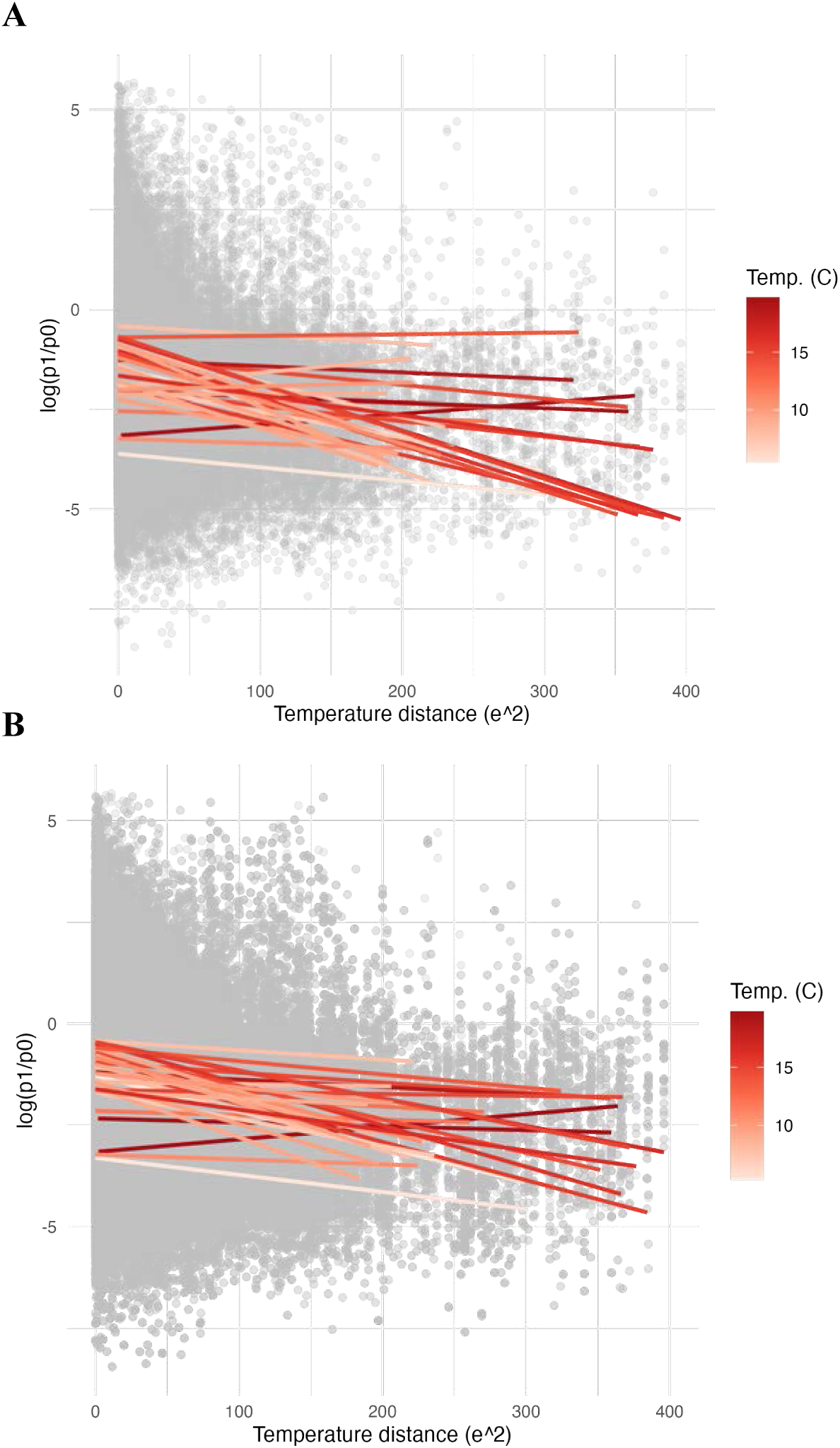
Stabilizing selection per site for terminal year and all years Fitting a regression for temperature distance to all sites truncated to e^2^<400 (to avoid non-linearities caused by south and north range distances). Slopes *Vs^−1^*are fitted per experimental (*n* = 30) where the color of the slope indicates the temperature of the experimental garden (red). (**A**) Terminal year. (**B**) All years combined.

### PHENOTYPES

**Figure S43.**
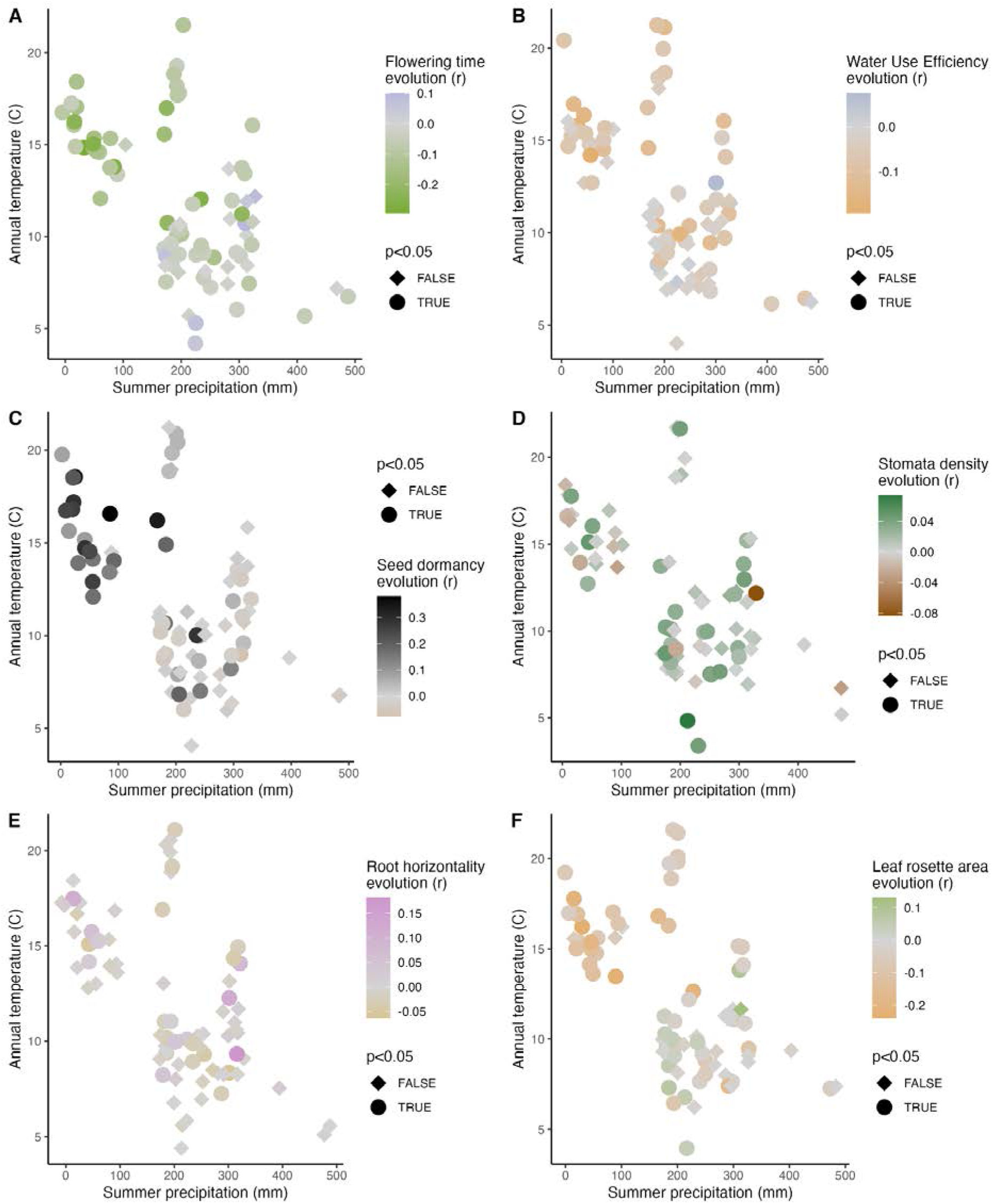
Phenotype-to-accession relative frequency correlation across environments. Using only sites from the Eurasian range (*n* = 22) and accessions with overlap with a homogenized dataset (*n* = 225), this figure shows the marginal phenotypic correlation between phenotypes measured in standard growth chamber conditions on the same founding accessions or imputed from a large consolidation effort, and accession relative frequency changes over one generation. (**A**) Flowering time measured at 10°C in growth chambers (*h^2^_kinship_ =* 0.93 [95%CI 0.898-0.975]). (**B**) Water use efficiency proxy (δ^13^C) (*h^2^_kinship_ =* 0.808 [95%CI 0.537-0.933]). (**C**) Seed dormancy (DSDS50, *h^2^_kinship_ =* 0.987 [95%CI 0.934-0.998]). (**D**) Stomata density (*h^2^_kinship_ =* 0.384 [95%CI 0.0579-0.773]). (**E**) Root horizontality (*h^2^_kinship_ =* 0.833 [95%CI 0.468-0.98]). (**F**) Leaf rosette area (*h^2^_kinship_ =* 0.941[95%CI 0.677-1]). Each panel displays the correlation (*r*) between phenotypic traits and accession relative frequency changes, plotted against annual temperature (°C) and annual precipitation (mm). Significant correlations (*P* < 0.05) are indicated, highlighting the relationship between phenotypic traits and evolutionary responses across different environmental conditions. NB: jitter was added to data points of similar climates a jitter of 2C temperature and 10 mm precipitation was added for visualization purposes.

**Figure S44.**
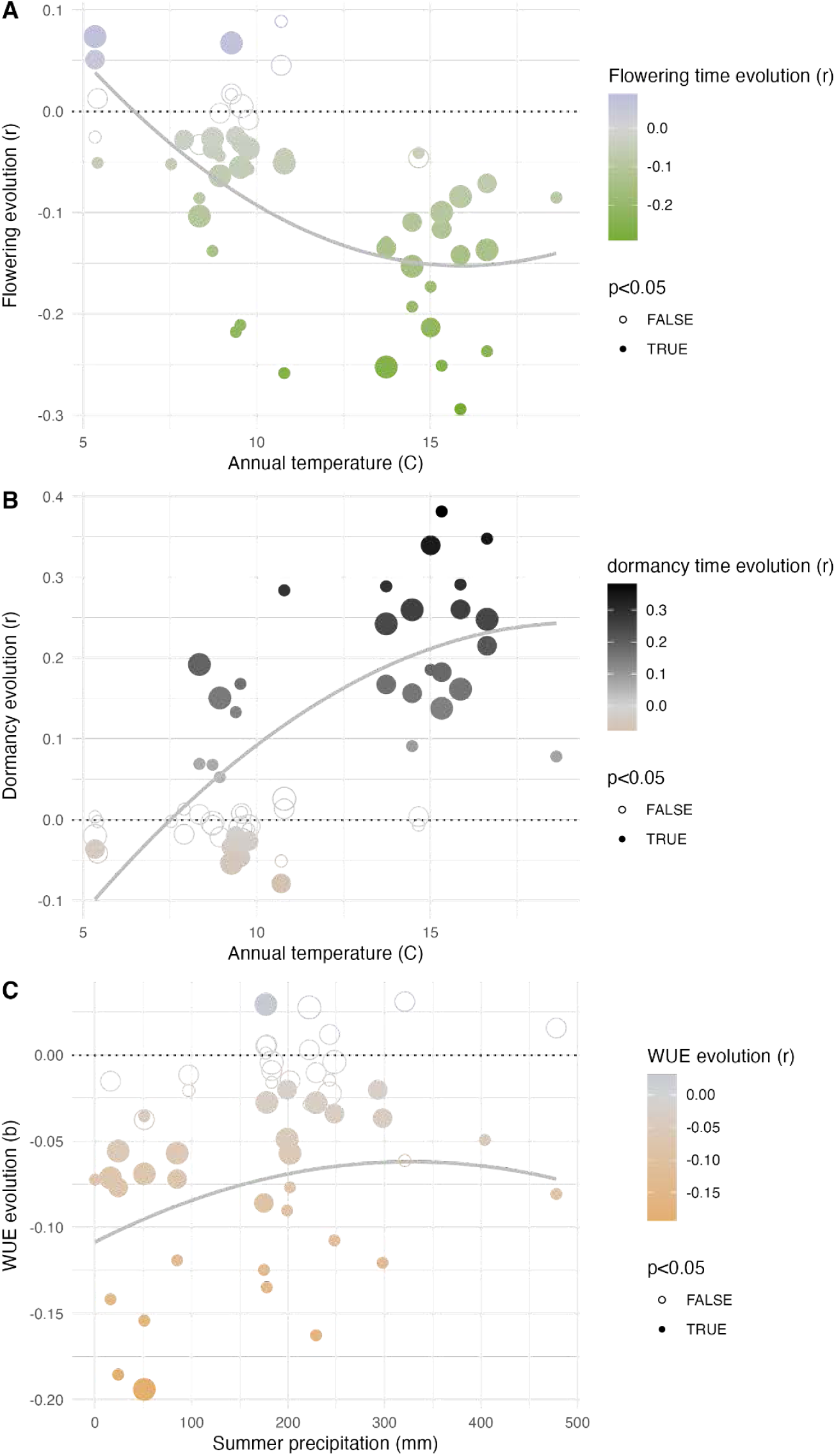
Phenotype evolution across experimental environmental gradients Correlation of accessions’ frequency with their flowering time (**A**), dormancy (DSDS50) (**B**) and water use efficiency (d^13^C) (**C**). Filled circles indicate significant correlations (*P<*0.05) and circle sizes indicate correlations with accession relative frequency between founders and different years (year 1 < year 2 < year 3). Lines indicate the best polynomial regression (y ∼ a + bx +cx^2^) using only the significant correlations.

### GENOME WIDE ASSOCIATIONS

**Figure S45.**
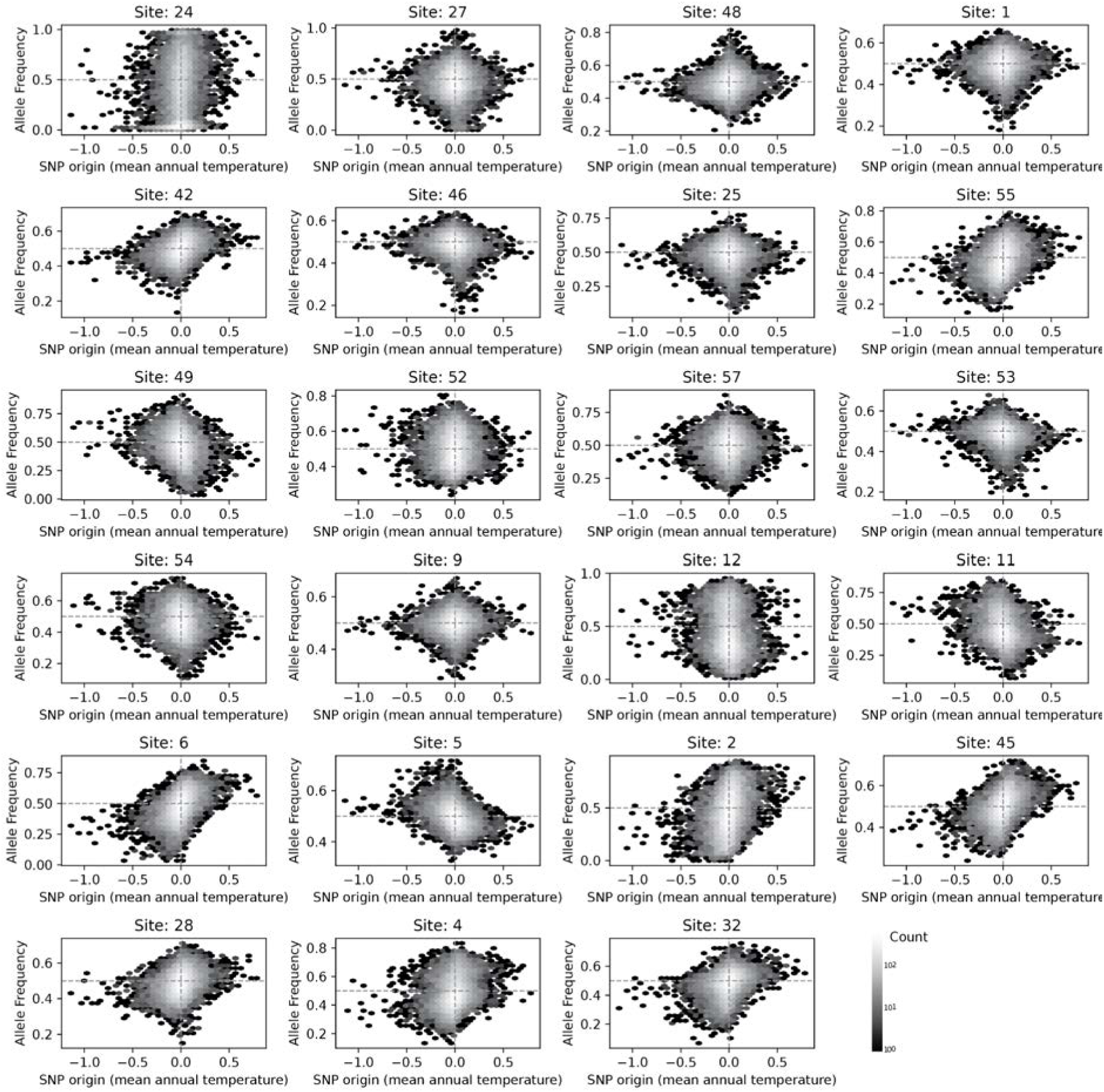
Change in allele frequency vs climatic origin for all sites. Temperature of origin of SNPs computed from those accessions carrying the allele and their location of origin compared to the allele frequency change over time at each common garden.

**Figure S46.**
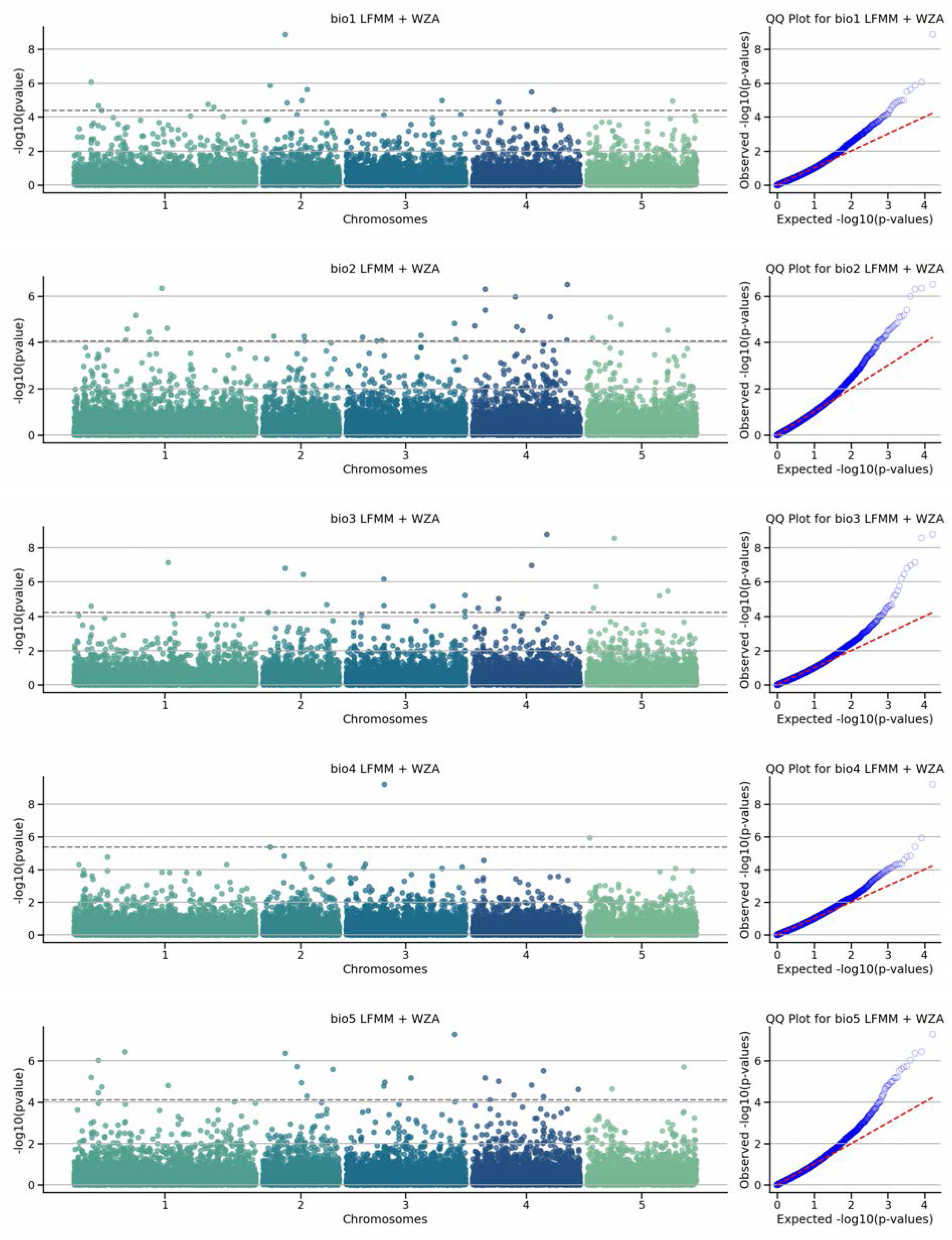
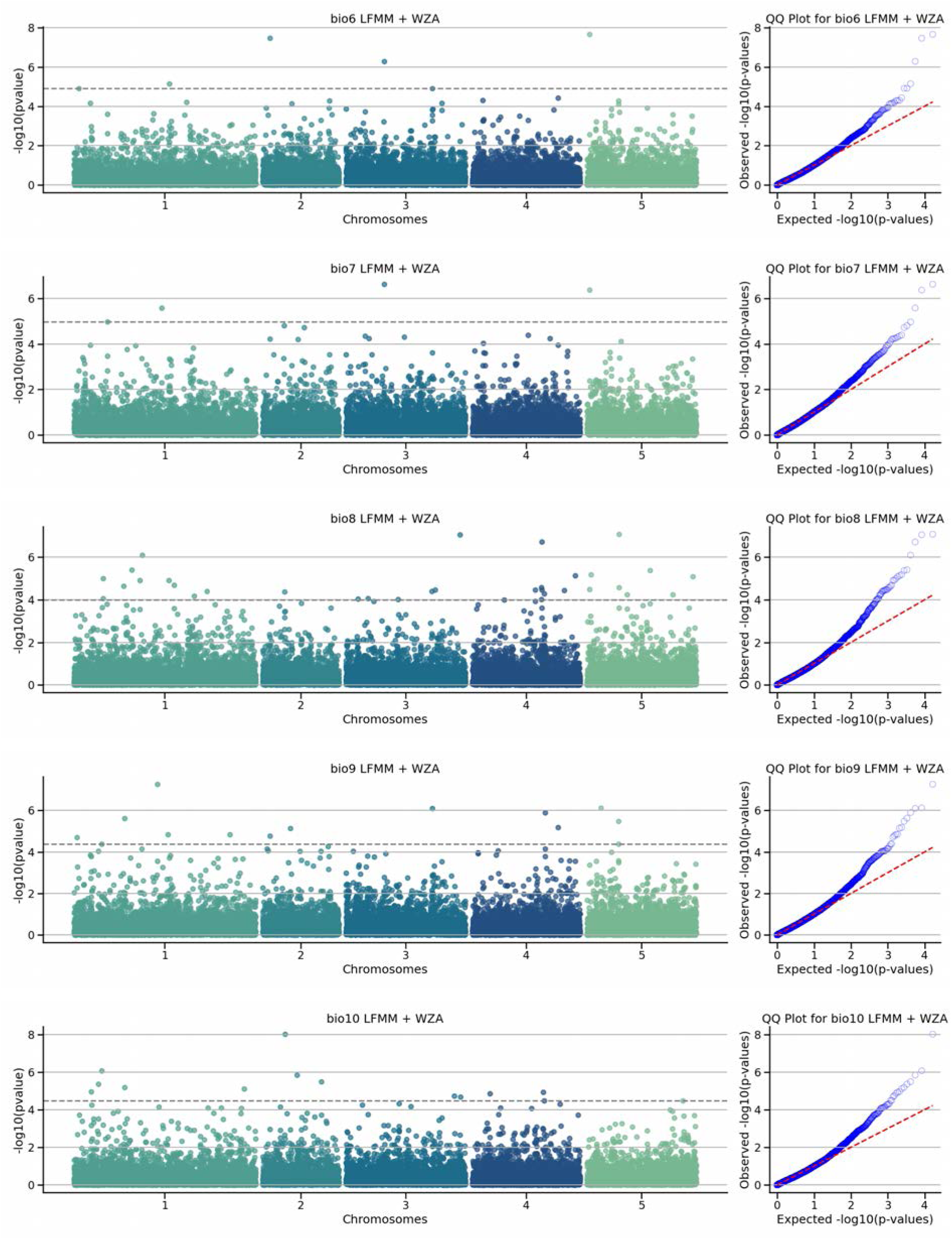
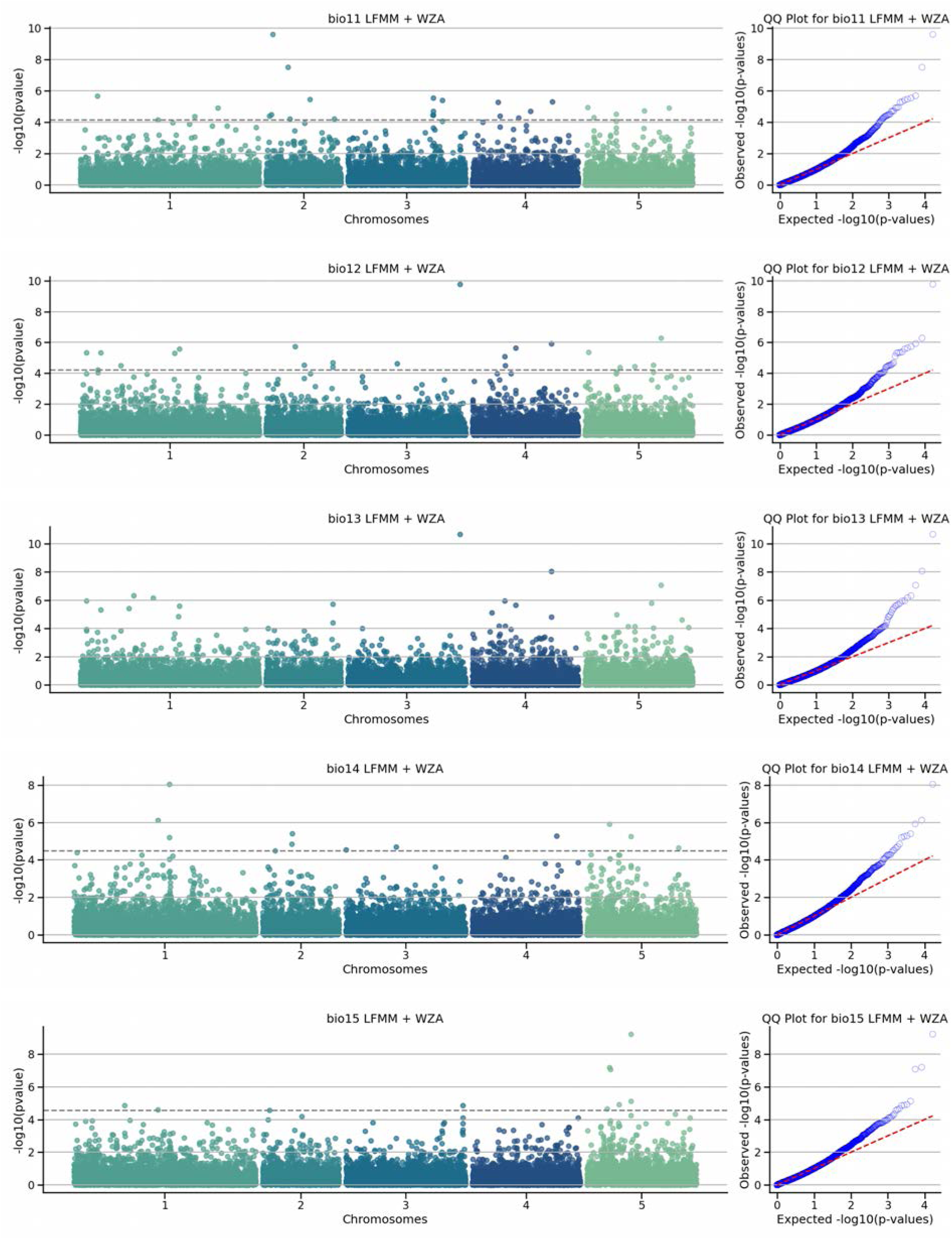
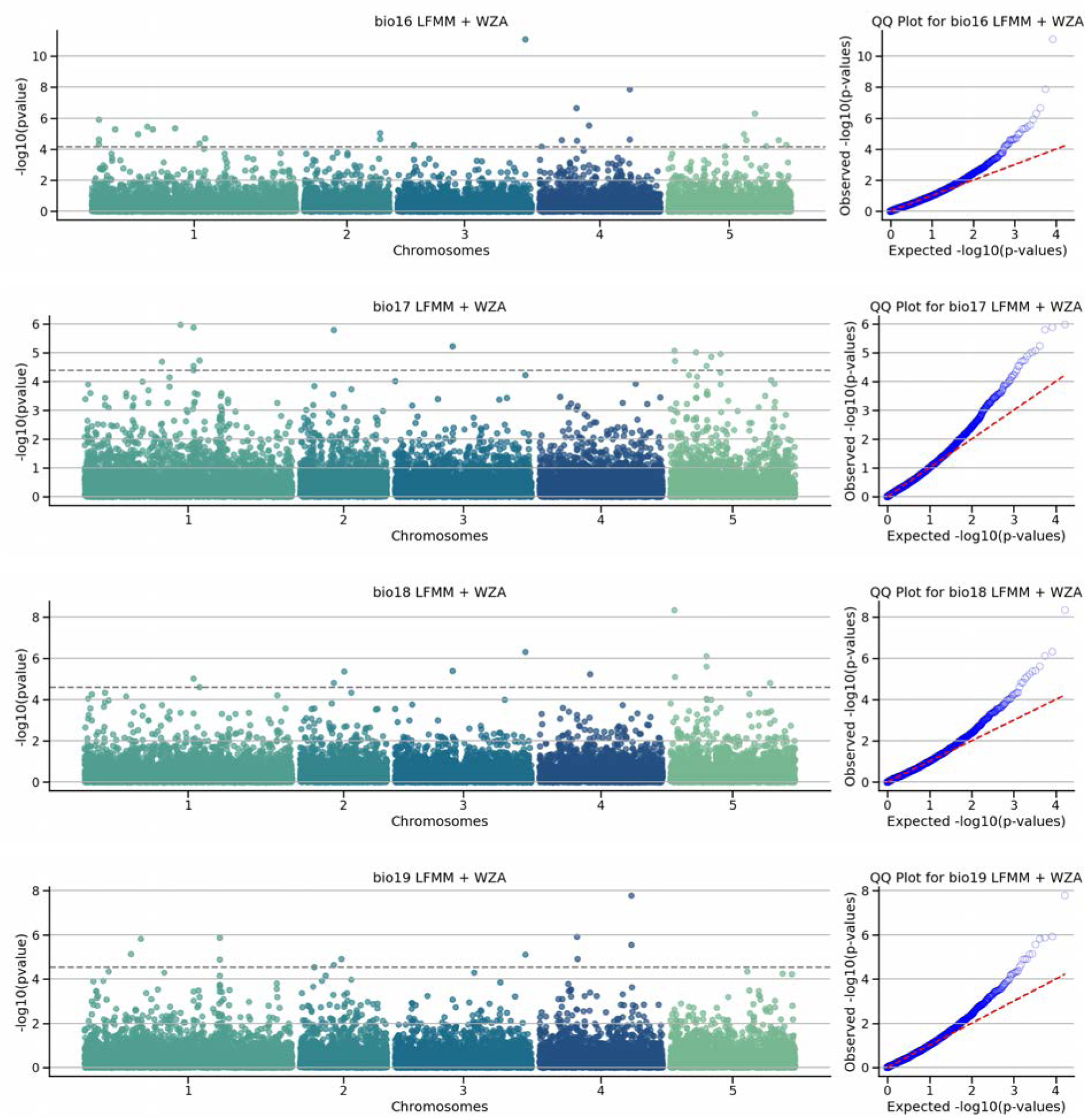
Manhattan and QQ plots of GEA using LFMM and WZA After performing GEA models with LFMM, the results were aggregated into haplotype blocks using the weighted Z-score (WZA). In the manhattan plot each dot represents a haplotype block. The dashed horizontal line indicates the significance threshold after correction for multiple testing using the Benjamini-Hochberg (BH).

**Figure S47.**
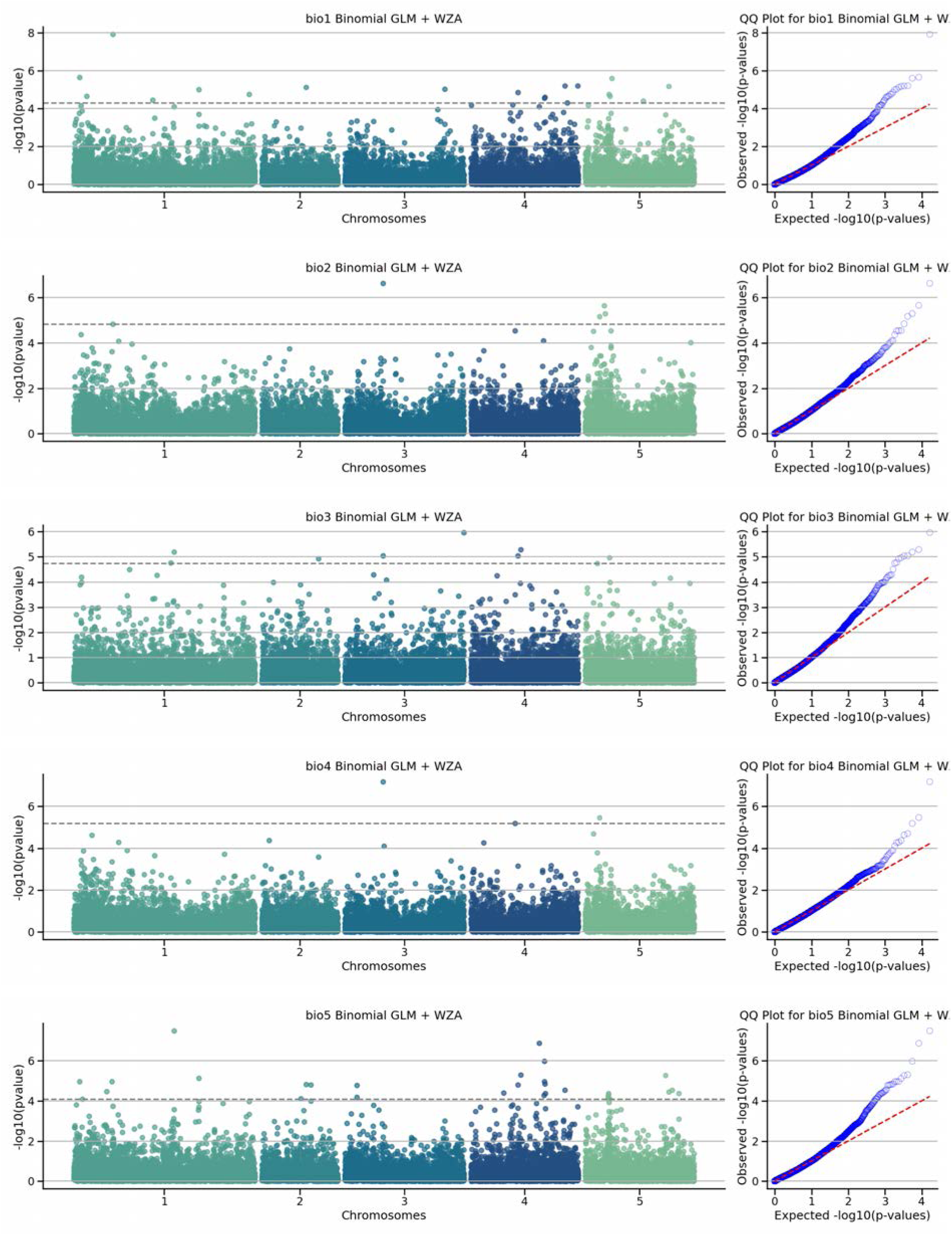
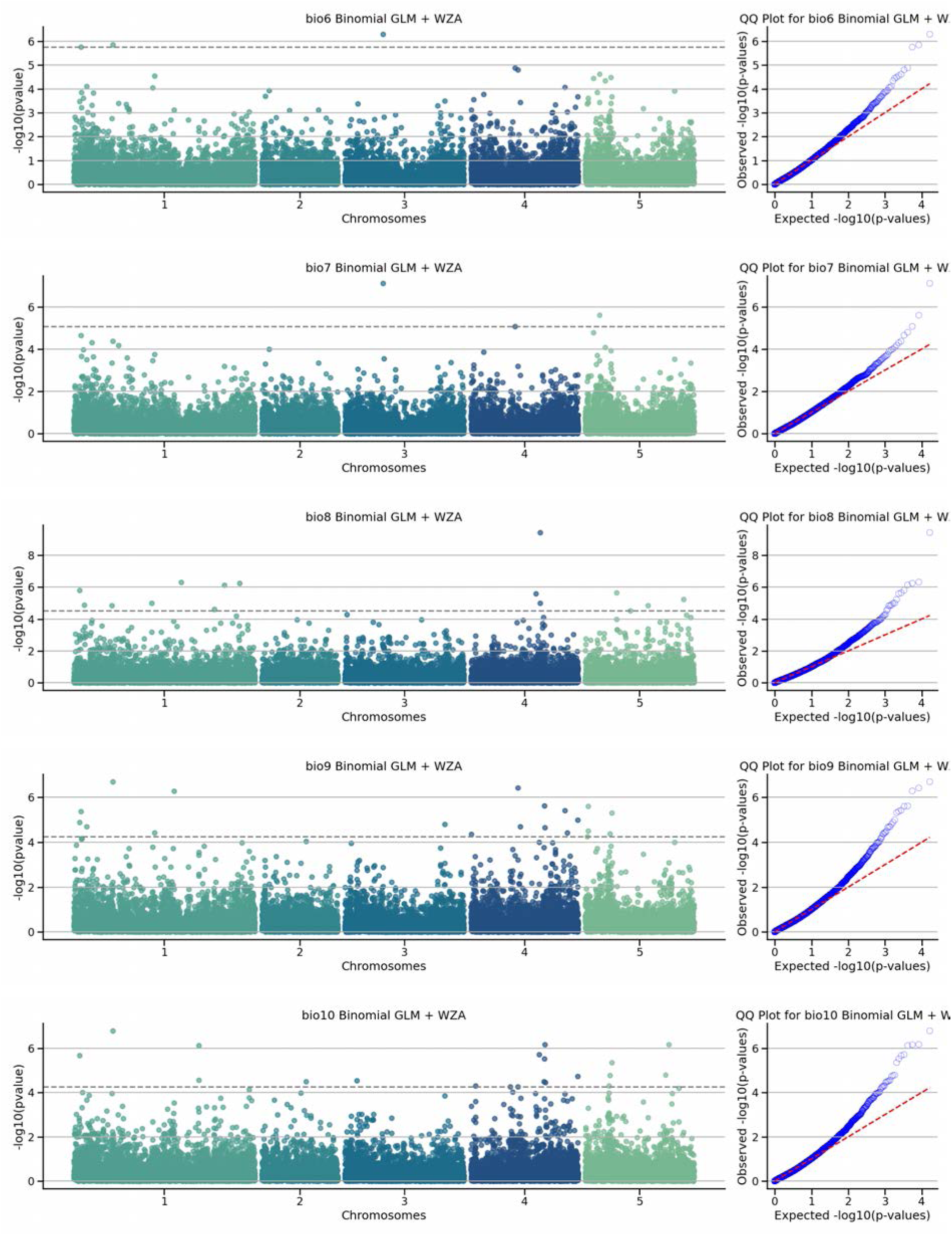
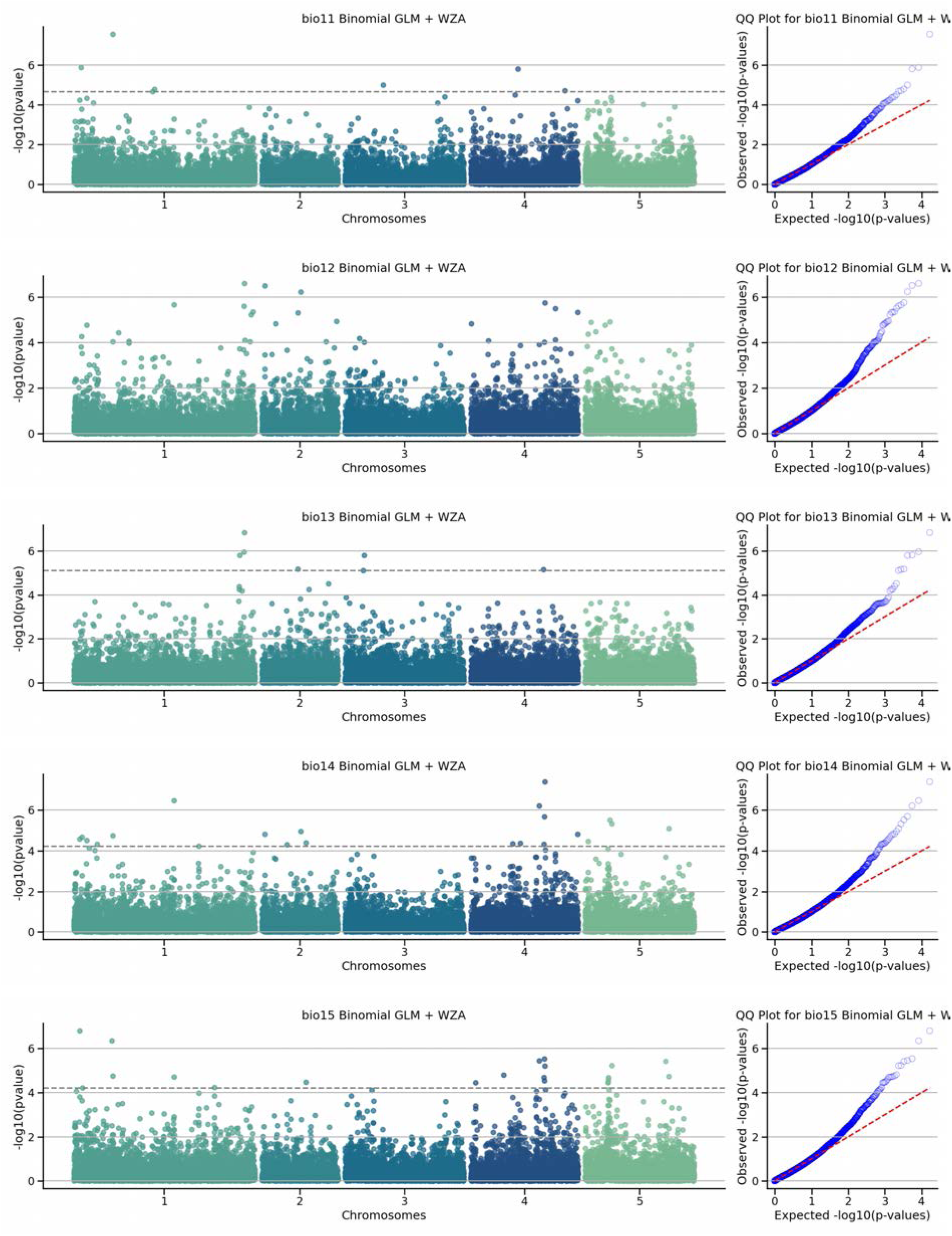
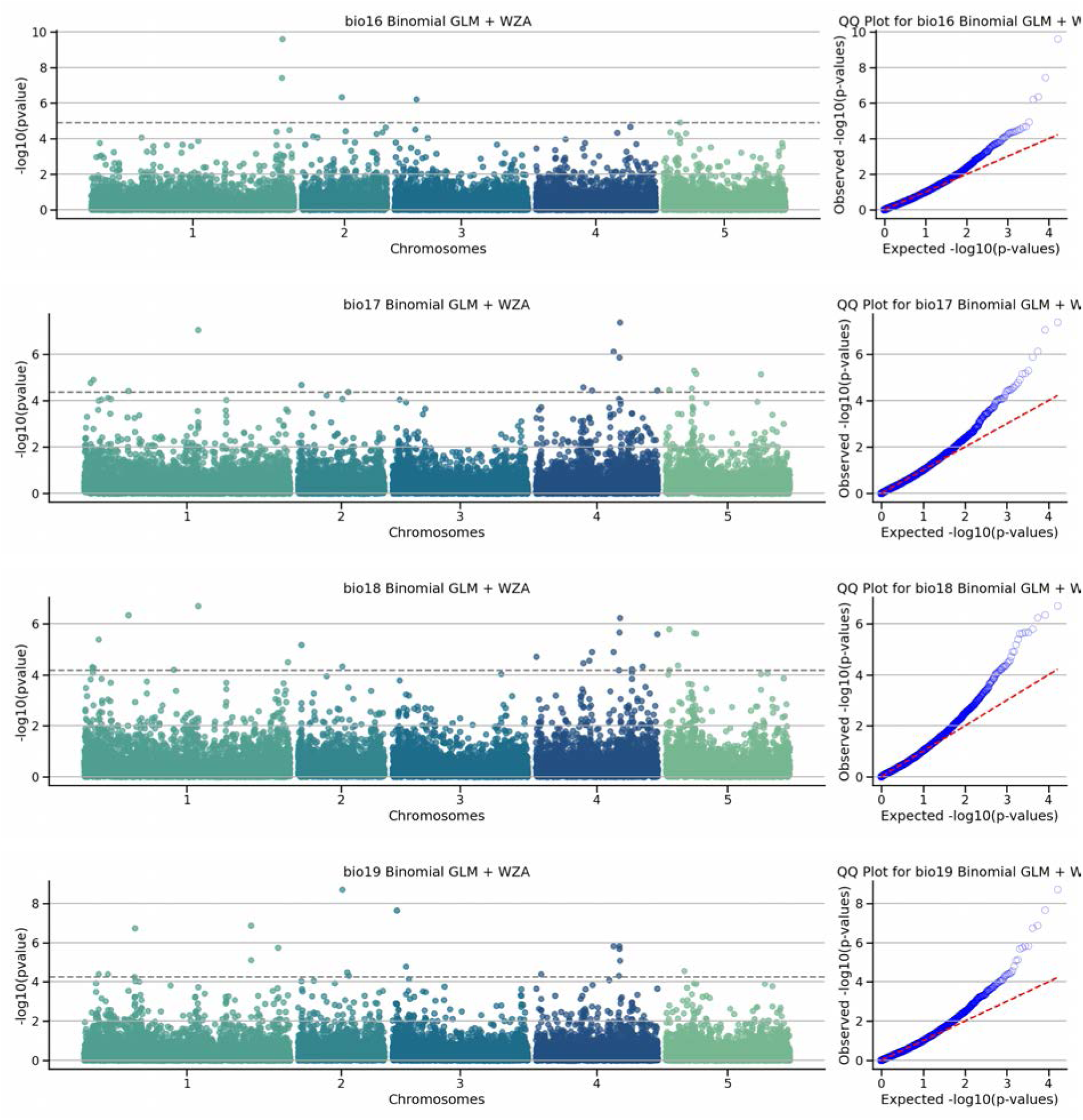
Manhattan and QQ plots of GEA using Binomial GLM and WZA After performing GEA models with a Binomial GLM, the results were aggregated into haplotype block using the weighted Z-score (WZA). In the manhattan plot each dot represents a haplotype block. The dashed horizontal line indicates the significance threshold after correction for multiple testing using the Benjamini-Hochberg (BH).

**Figure S48.**
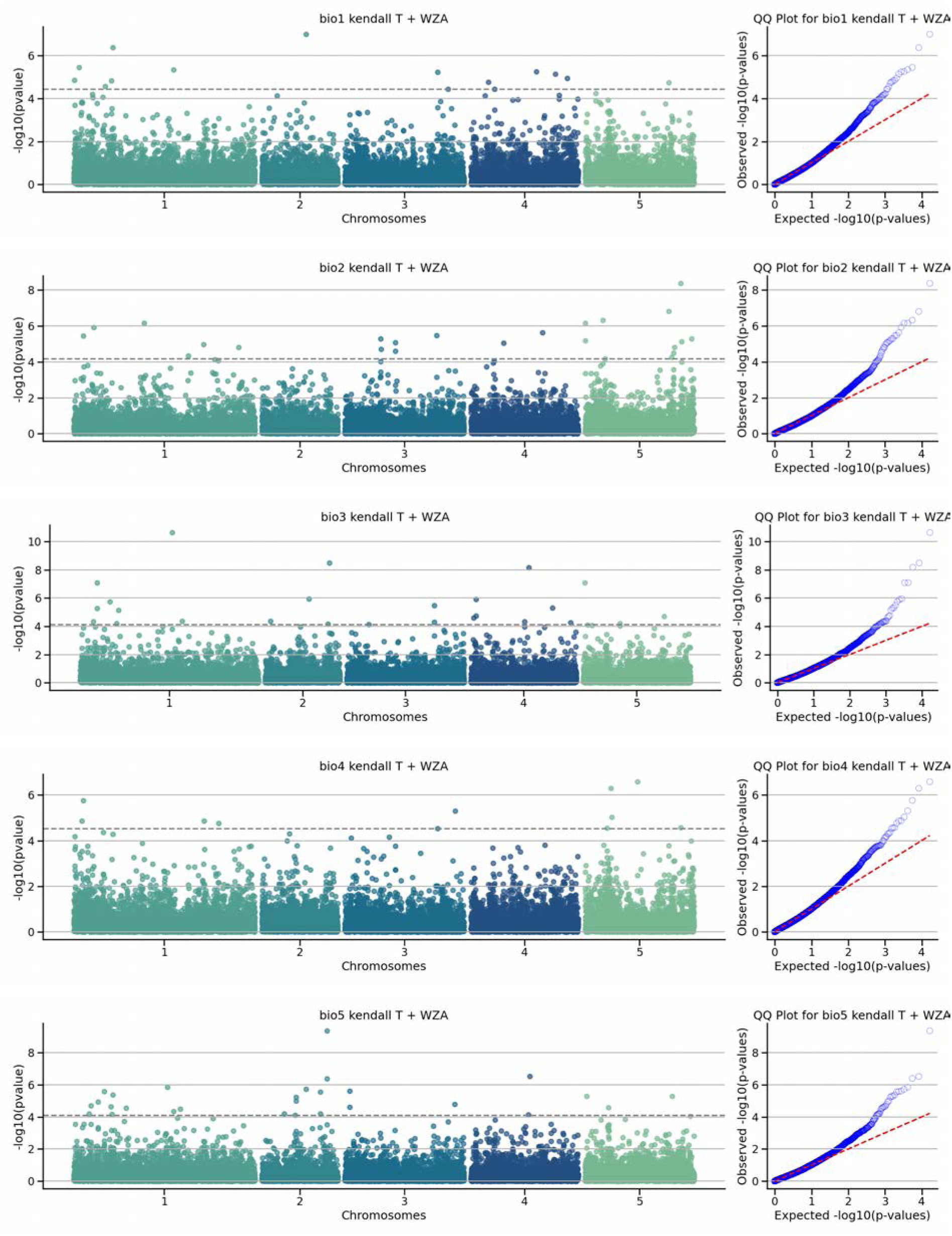
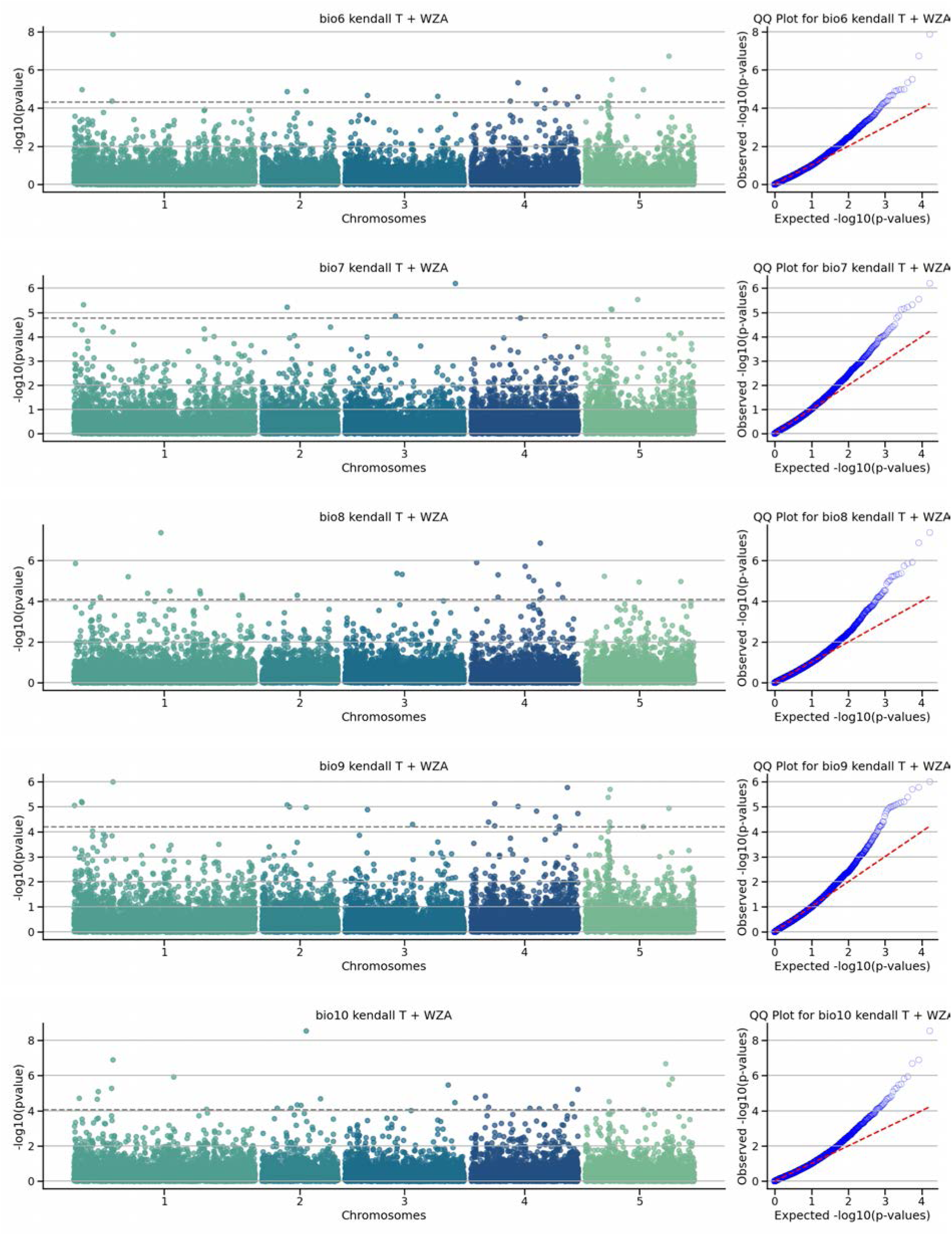
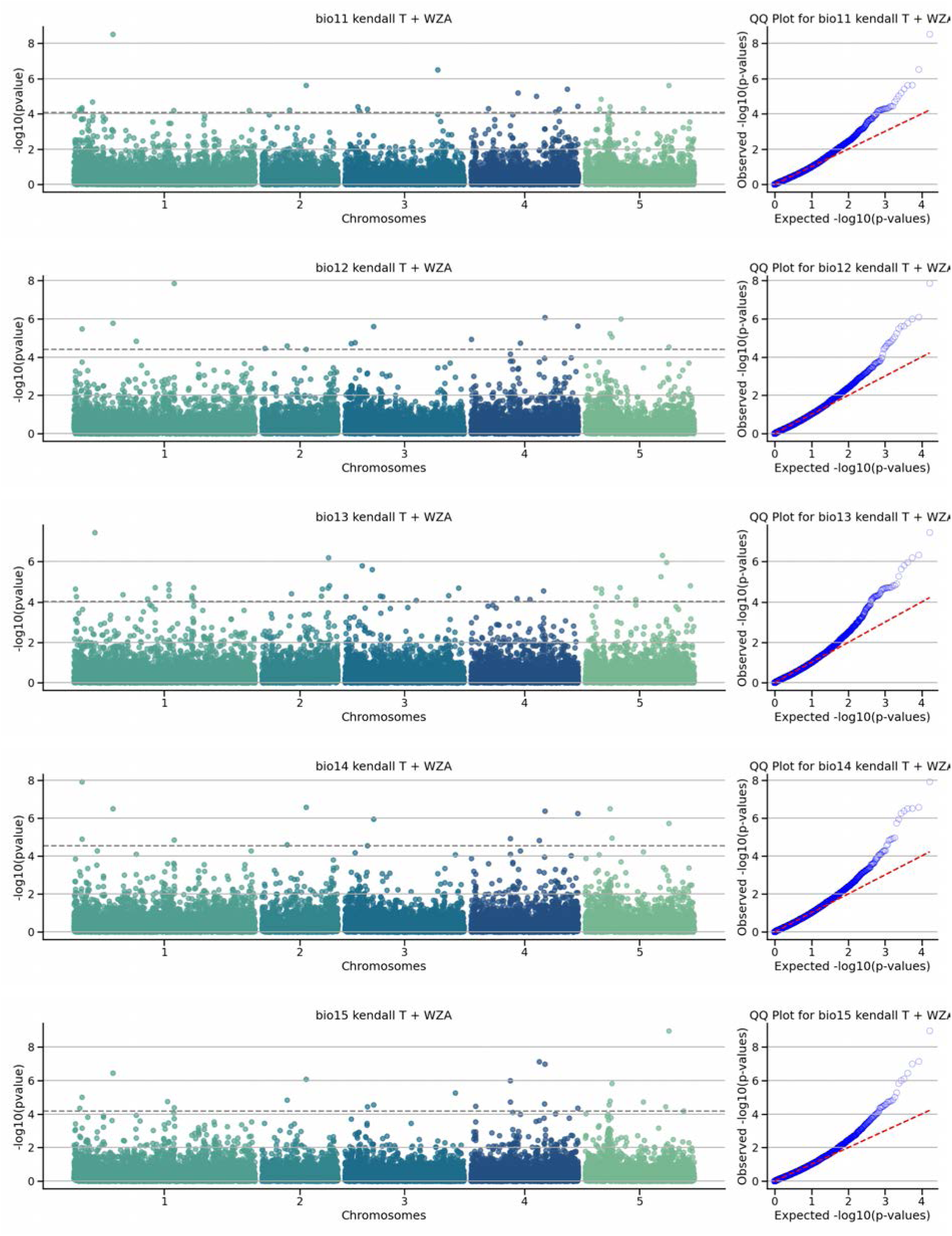
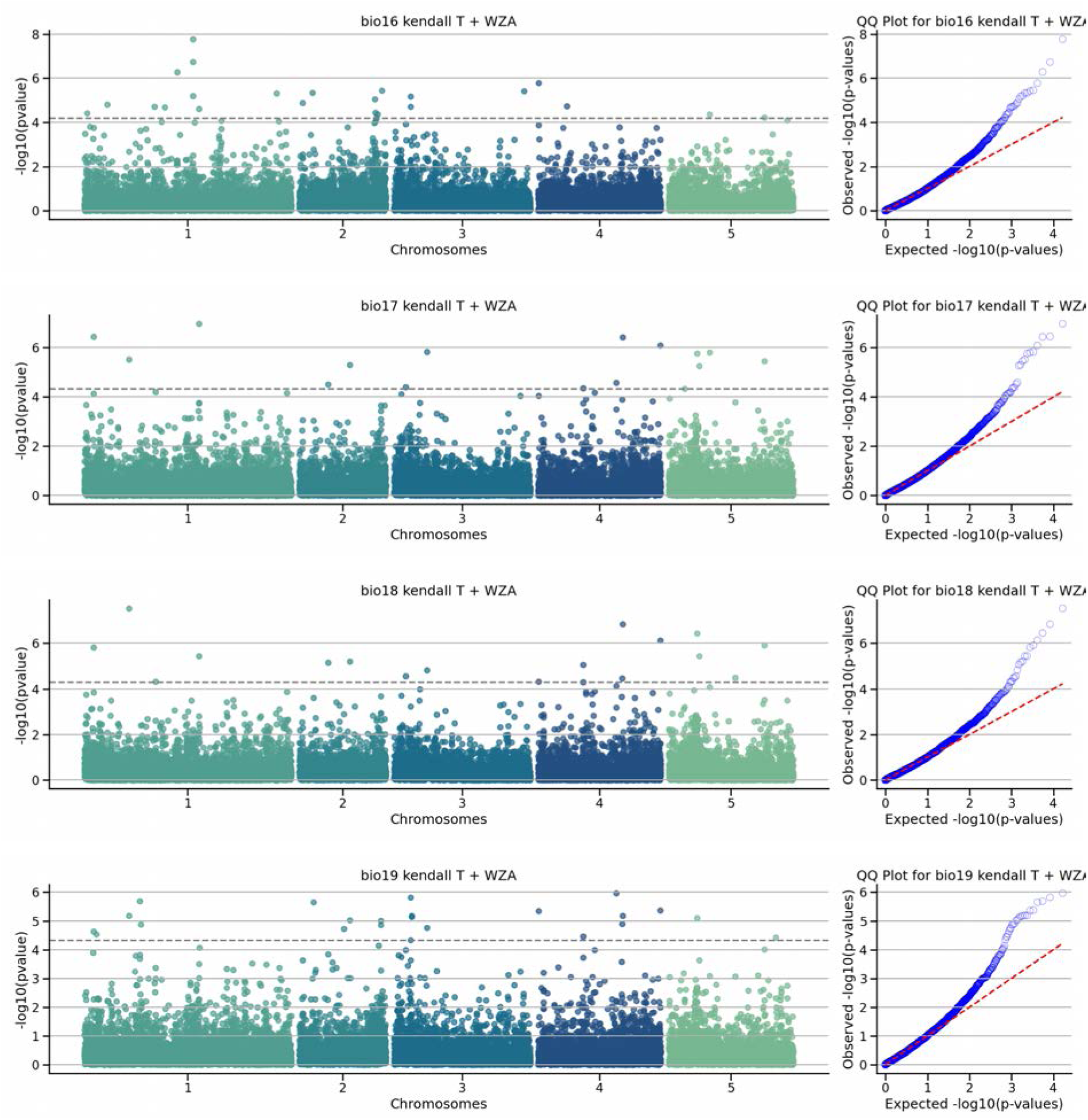
Manhattan and QQ plots of GEA usingKendall-τ correlation and WZA After performing GEA models with a Kendall-τ ranked correlation, the results were aggregated into haplotype blocks using the weighted Z-score (WZA). In the manhattan plot each dot represents a haplotype block. The dashed horizontal line indicates the significance threshold after correction for multiple testing using the Benjamini-Hochberg (BH).

**Figure S49.**
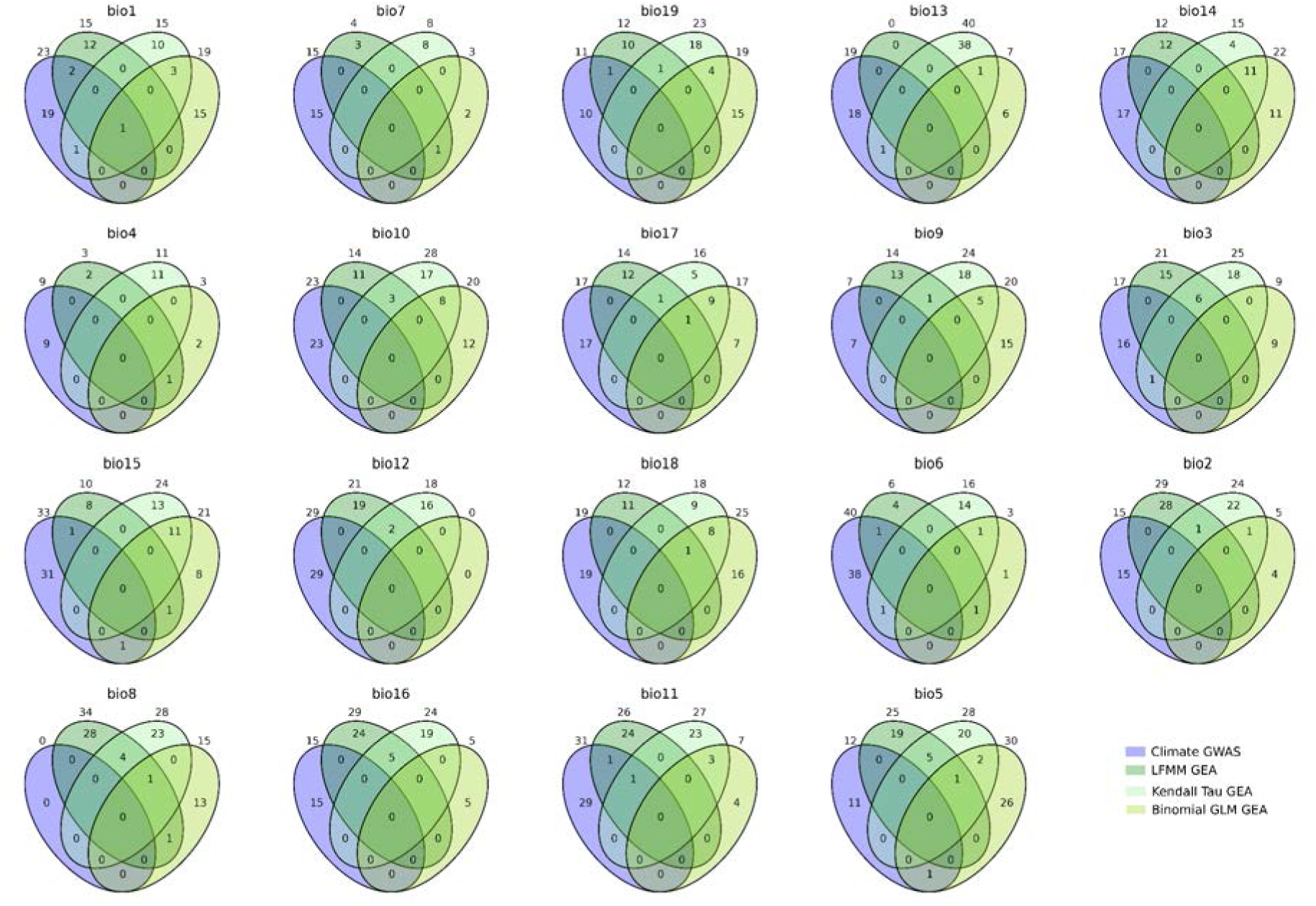
Venn diagrams of haplotype blocks identified as significant in eGEA models and classic climate GWAS. Extent of overlap on the haplotpye blocks identified in a classic climate GWAS and the GEA models. Table with the identified haplotype blocks and the genes present on them on **<u>S14</u>**

### GENE EXAMPLES

**Figure S50.**
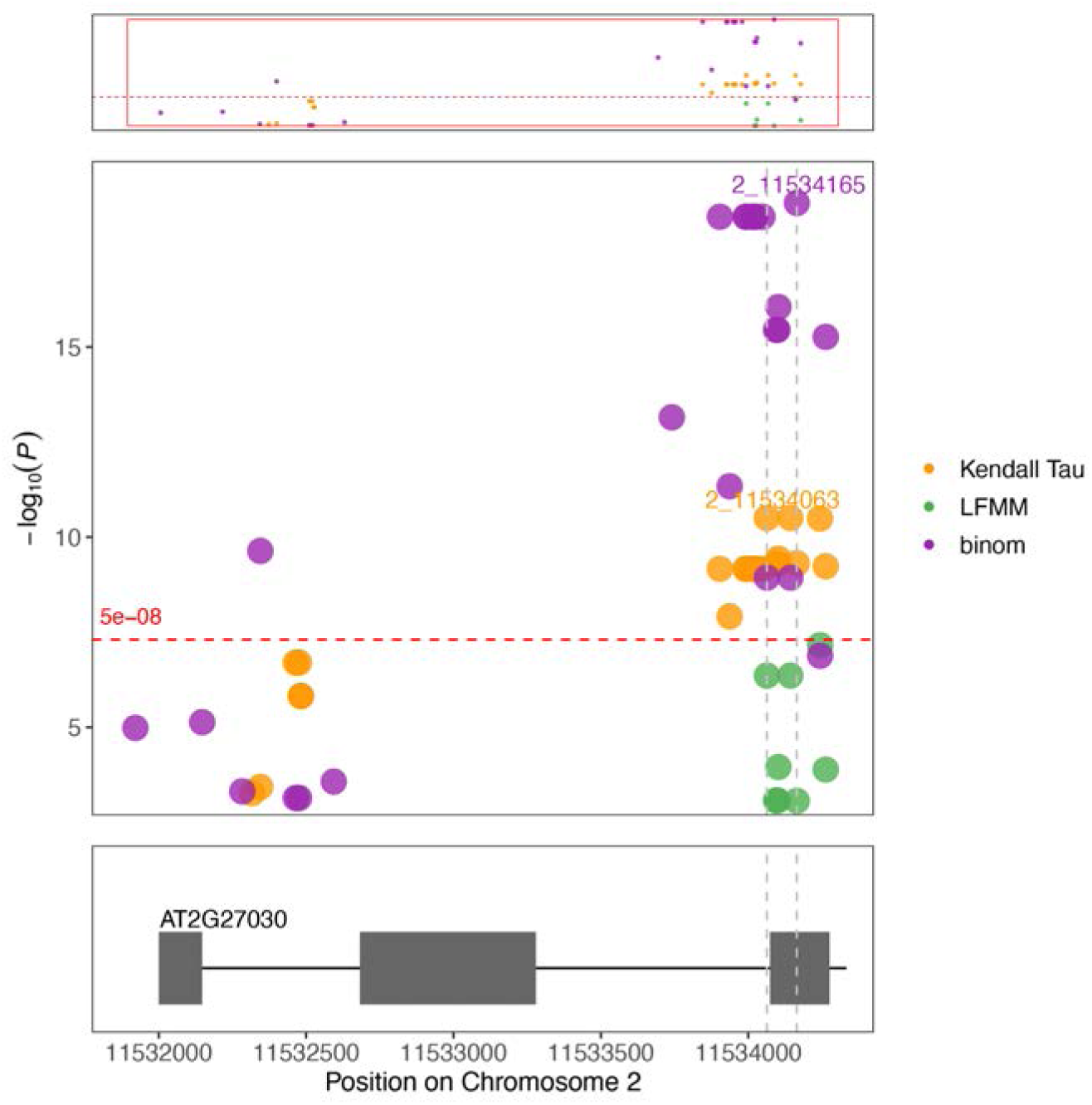
Manhattan plot of GEA with temperature in chromosome 2 Zoom in on chromosome 2 peaks of GrENE-net’s experimental evolution Genome-Environment Association (eGEA) using LFMM identifies *CAM5* gene (AT2G27030).

**Figure S51.**
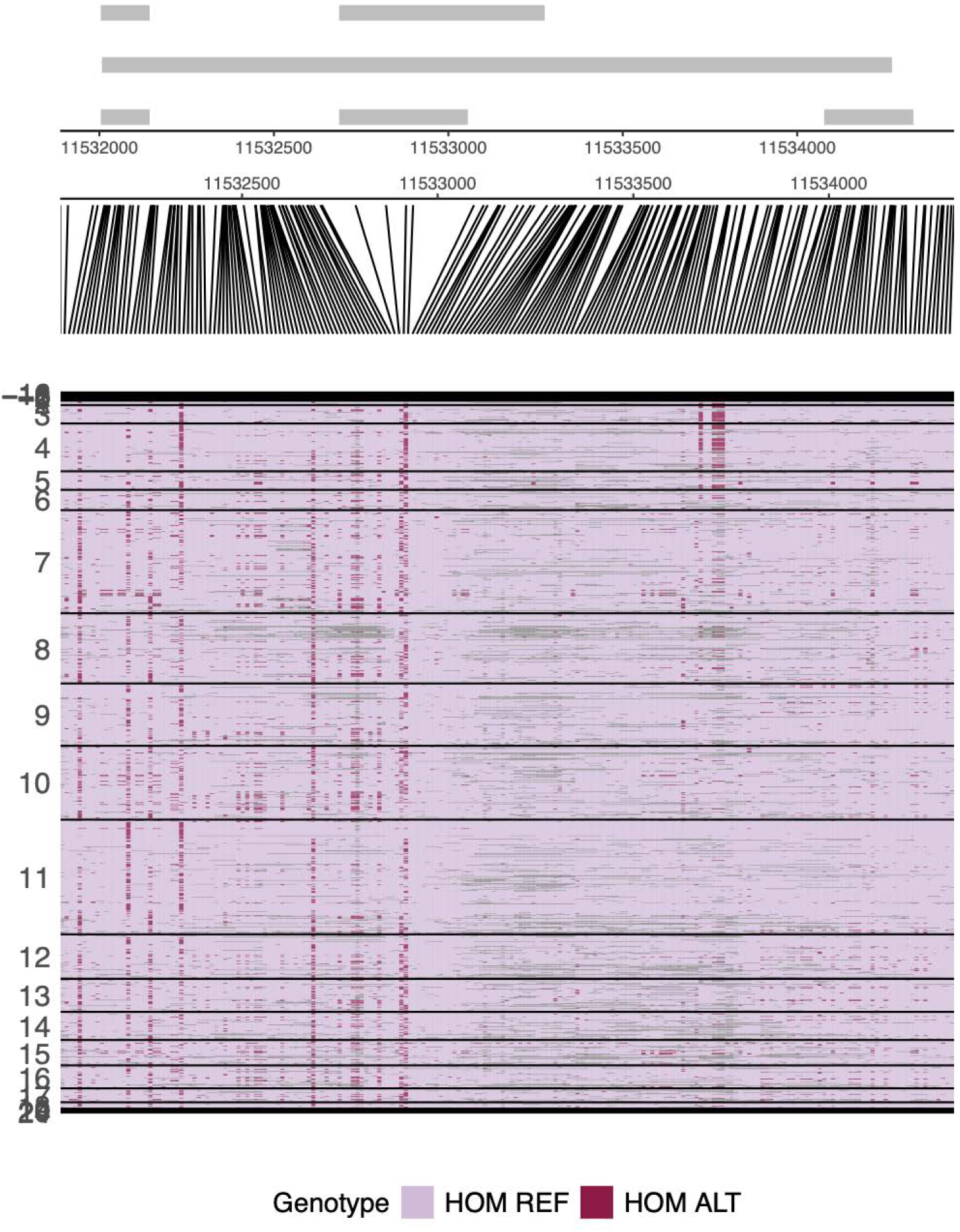
VCF visualization of *CAM5* Visualization of variant call format (VCF) table of 231 founder accessions showing reference or alternative genotypes, and grouped and sorted by annual temperature of origin (y-axis). TAIR10 gene models of *CAM* 5 are shown with exons in grey boxes.

**Figure S52.**
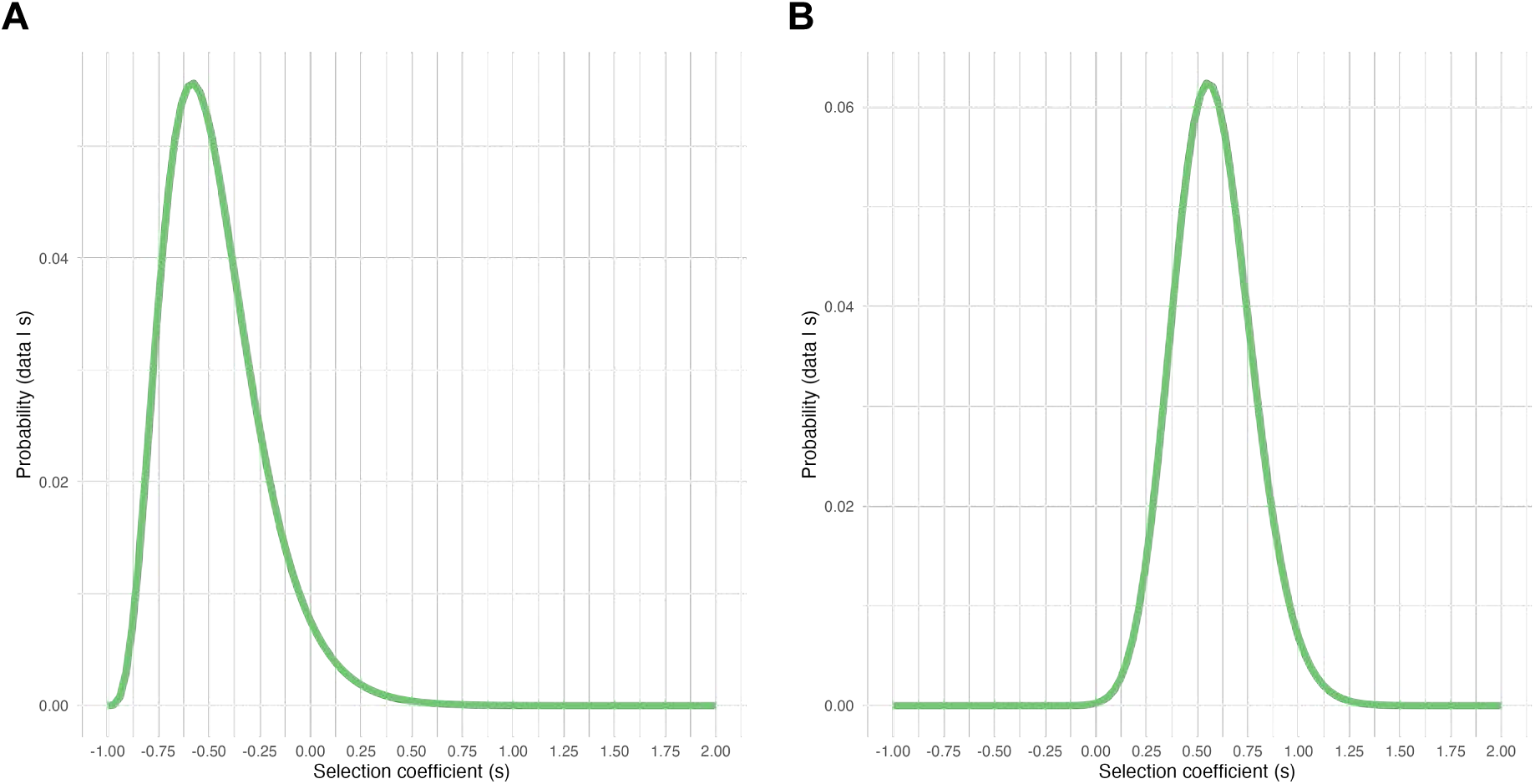
*CAM5* (AT2G27030) top SNP selection coefficient Selection coefficients estimated using information on the number of flower sampled and a Binomial likelihood in (**A**) Brixen im Thale, Austria as example of a cold site and (**B**) Cadiz, Spain, as example warm site.

**Figure S53.**
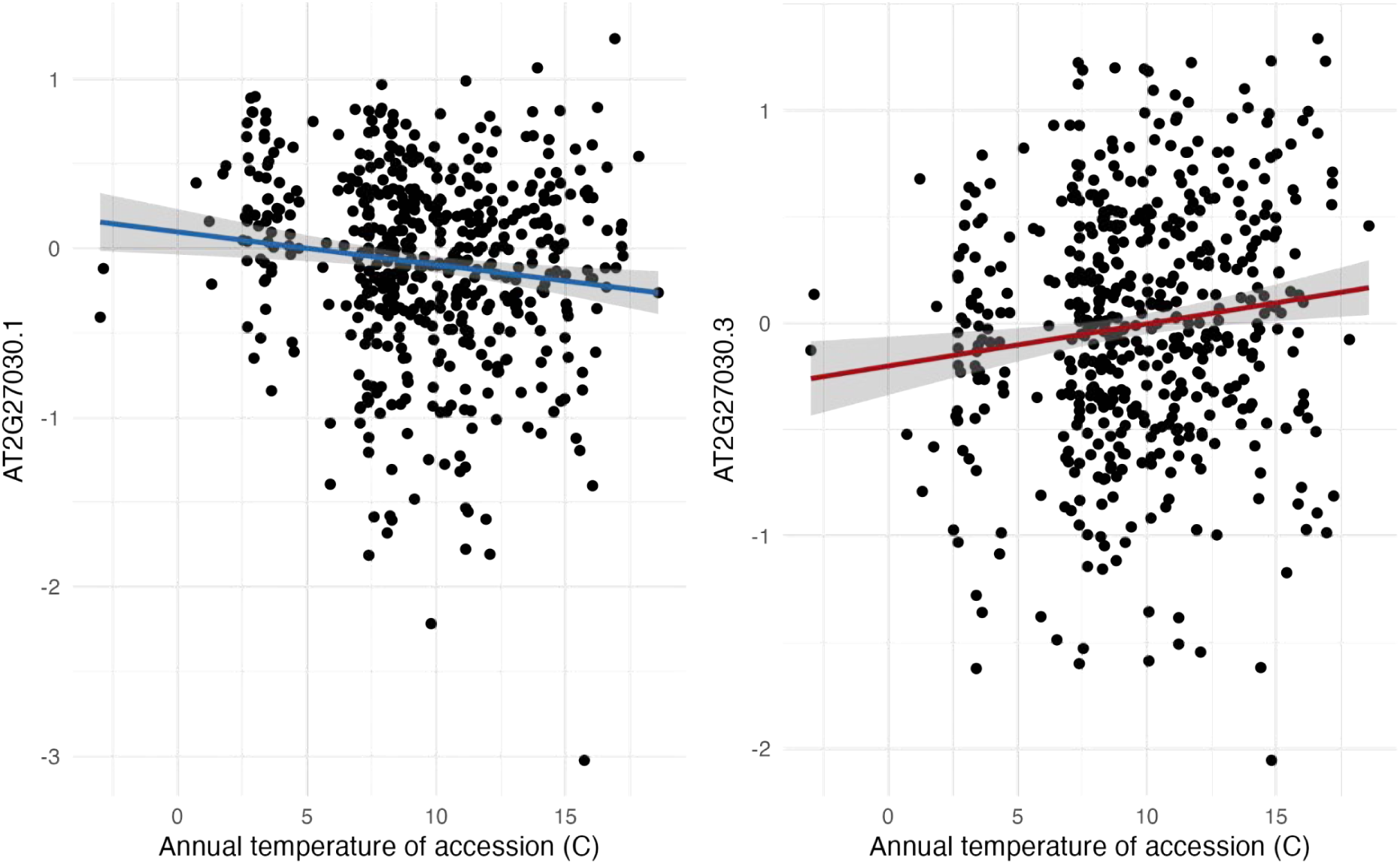
*CAM5* (AT2G27030) transcript expression across the 1001 transcriptomes. Normalized transcript expression (log2 with batch correction) from the 1001 Transcriptomees (*54*, *128*) of the splice variant 1 and 3 of *CAM5* is correlated with annual temperature of accessions at origin (*n* = 521, Spearman’s *r* = −0.124, +0.125, *P =* 0.0045, 0.0041; respectively).

**Figure S54.**
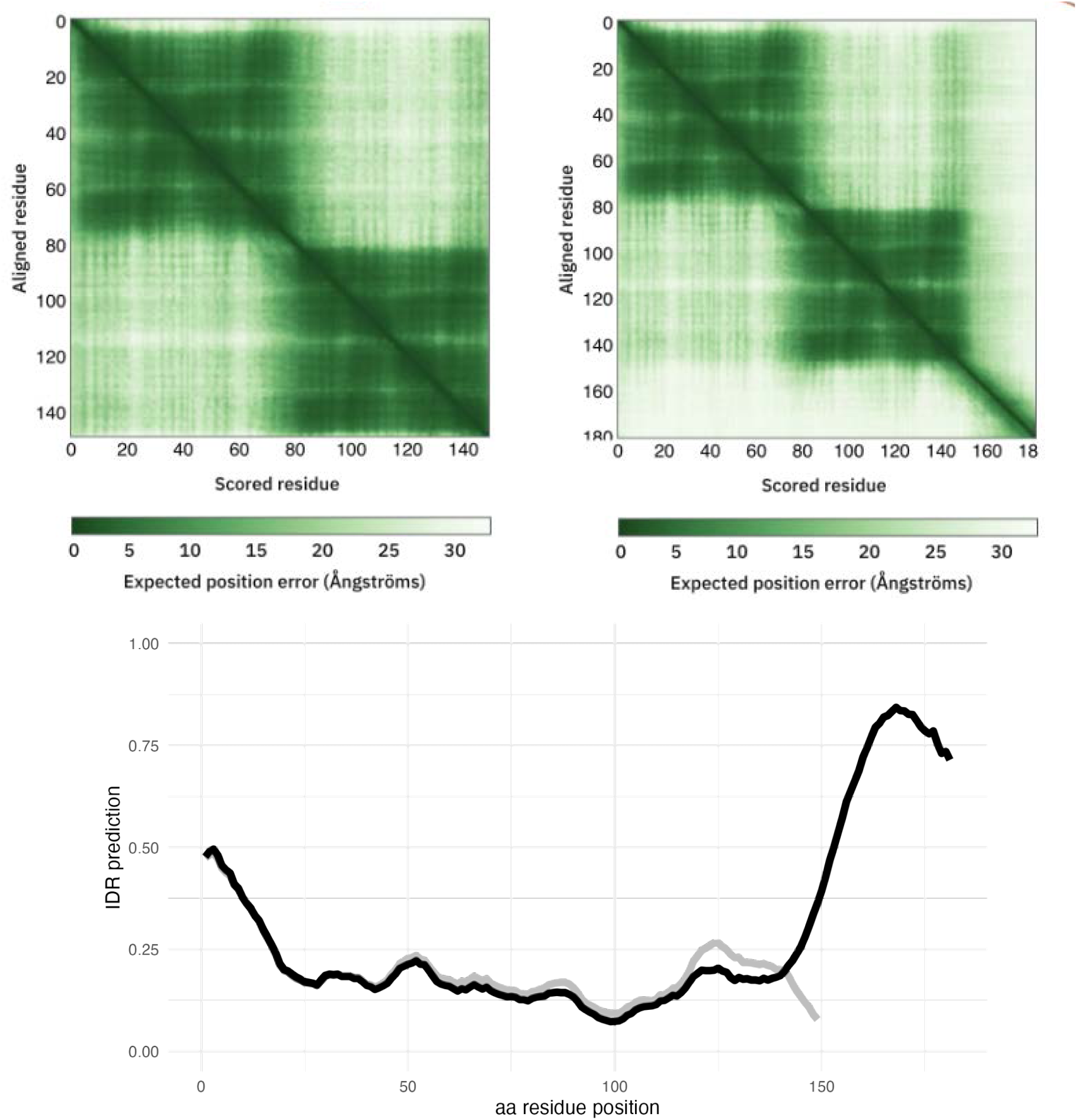
CAM5 AlphaFold2 protein structure. AlphaFold2 predictions of CAM5 proteins of the two splice variants. Splice variant AT2G27030.1 (https://alphafold.ebi.ac.uk/entry/Q682T9) (left) and splice variant AT2G27030.3 (https://alphafold.ebi.ac.uk/entry/F4IVN6) (right). IDR prediction on variant AT2G27030.1 (grey) and AT2G27030.3 (black) using *metapredict* v3.0.

**Figure S55.**
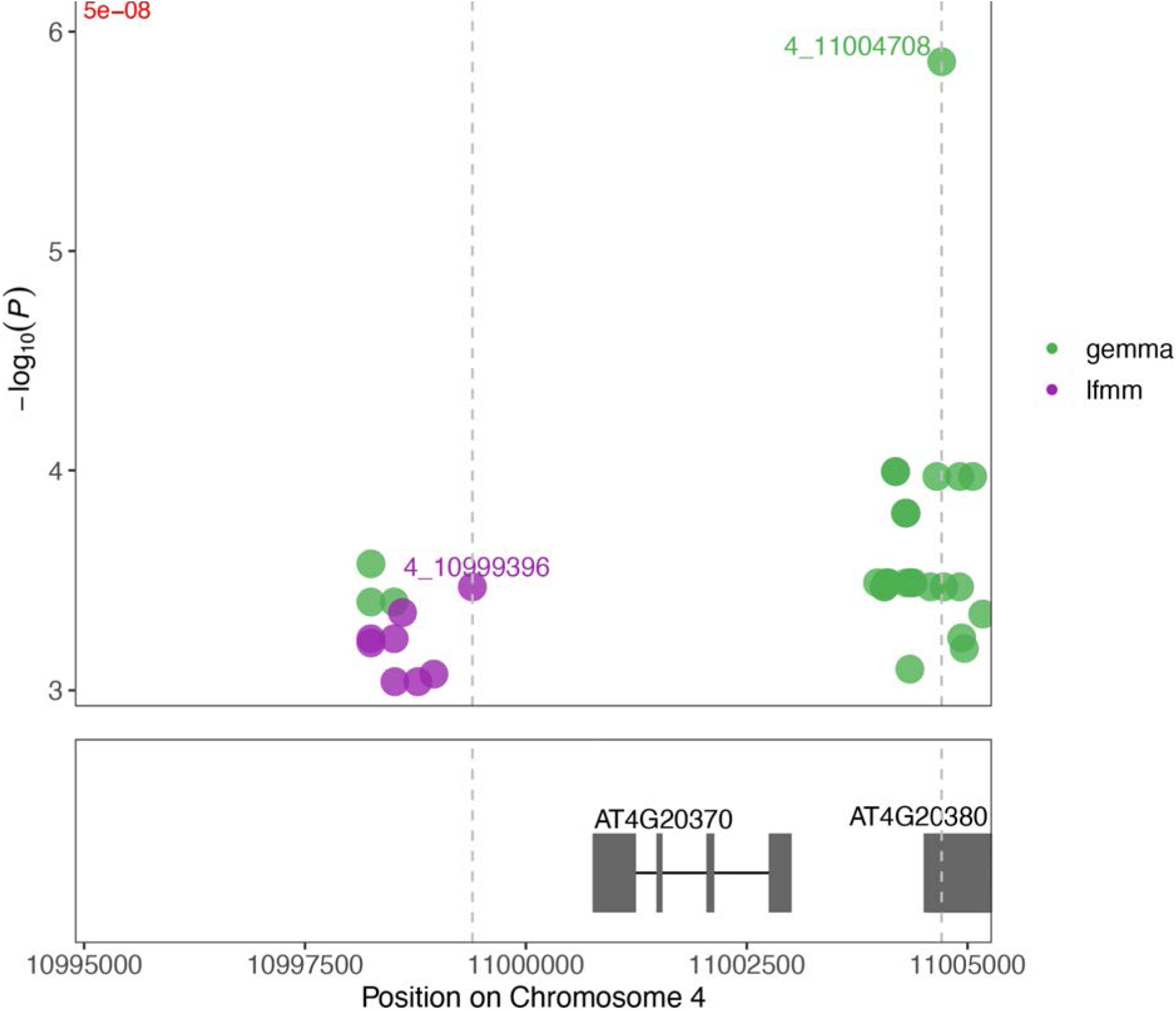
Manhattan plot of GEA with temperature on chromosome 4 Zoom in on chromosome 4 peaks of GrENE-net’s Genome-Environment Association (GEA) using LFMM identifies SNPs near *TSF* gene (AT4G20370). Two nearby SNPs are identified in the LFMM GEA of GrENE-net experimental evolution (purple) as well as LMM GEMMA GEA of the founder accessions data pinpointing associations of climate of origin of accessions (green). This region was already identified in addition to the nearby gene *LSD1* (AT4G20380,*CHILLING SENSITIVE 4 [ CHS4] or LESION SIMULATING DISEASE 1 [LSD1]*).

**Figure S56.**
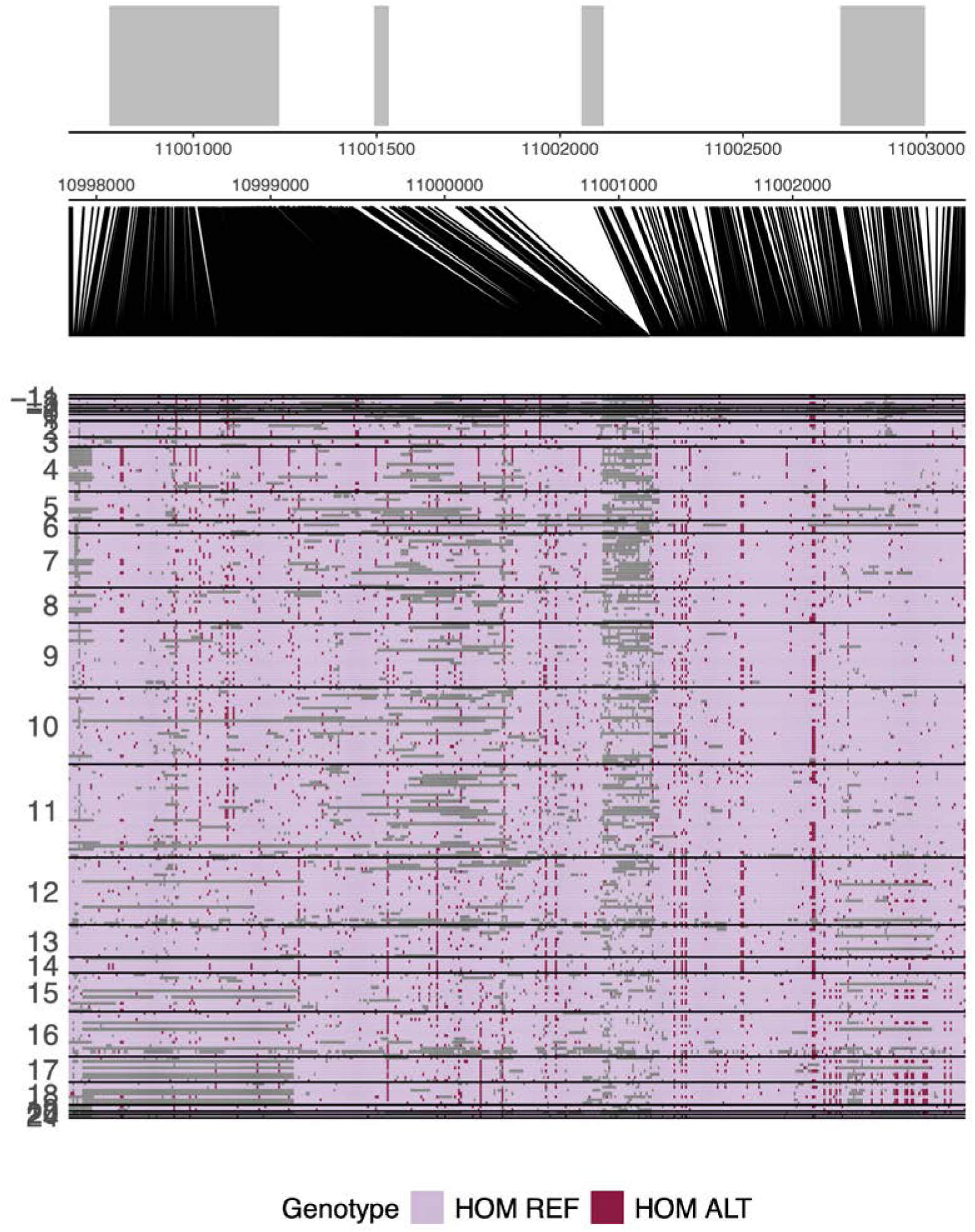
VCF visualization of *TSF* (AT4G20370) Visualization of variant call format (VCF) table of 231 founder accessions showing reference or alternative genotypes, and grouped and sorted by annual temperature of origin (y-axis). TAIR10 gene models of *TSF* are shown with exons in grey boxes.

**Figure S57.**
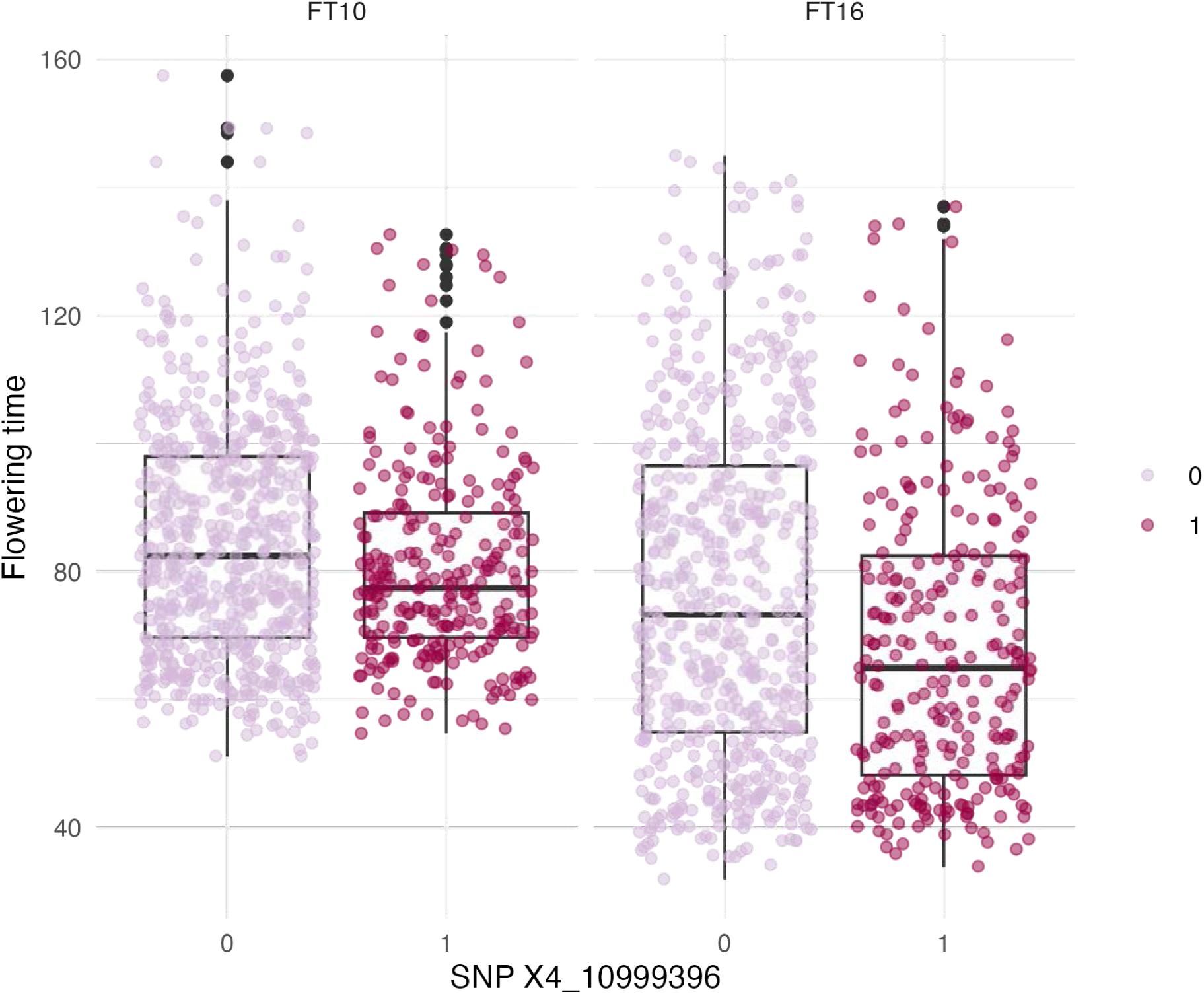
SNP near *TSF* gene identified in GEA and flowering time. Phenotype values of flowering time at 10°C and 16°C across the 1001 Genomes of *Arabidopsis* between the alternative and reference alleles. For FT10, the mean difference in flowering time was3.79 days (Wilcoxon test, *P* = 0.005), and for FT16, the mean difference was 8.64 days (Wilcoxon test, *P* = 1.2×10^−5^, *n* = 220). (For the effect of all alleles identified as significant near the TSF on flowering time see **Table <u>S18</u>**)

**Figure S58.**
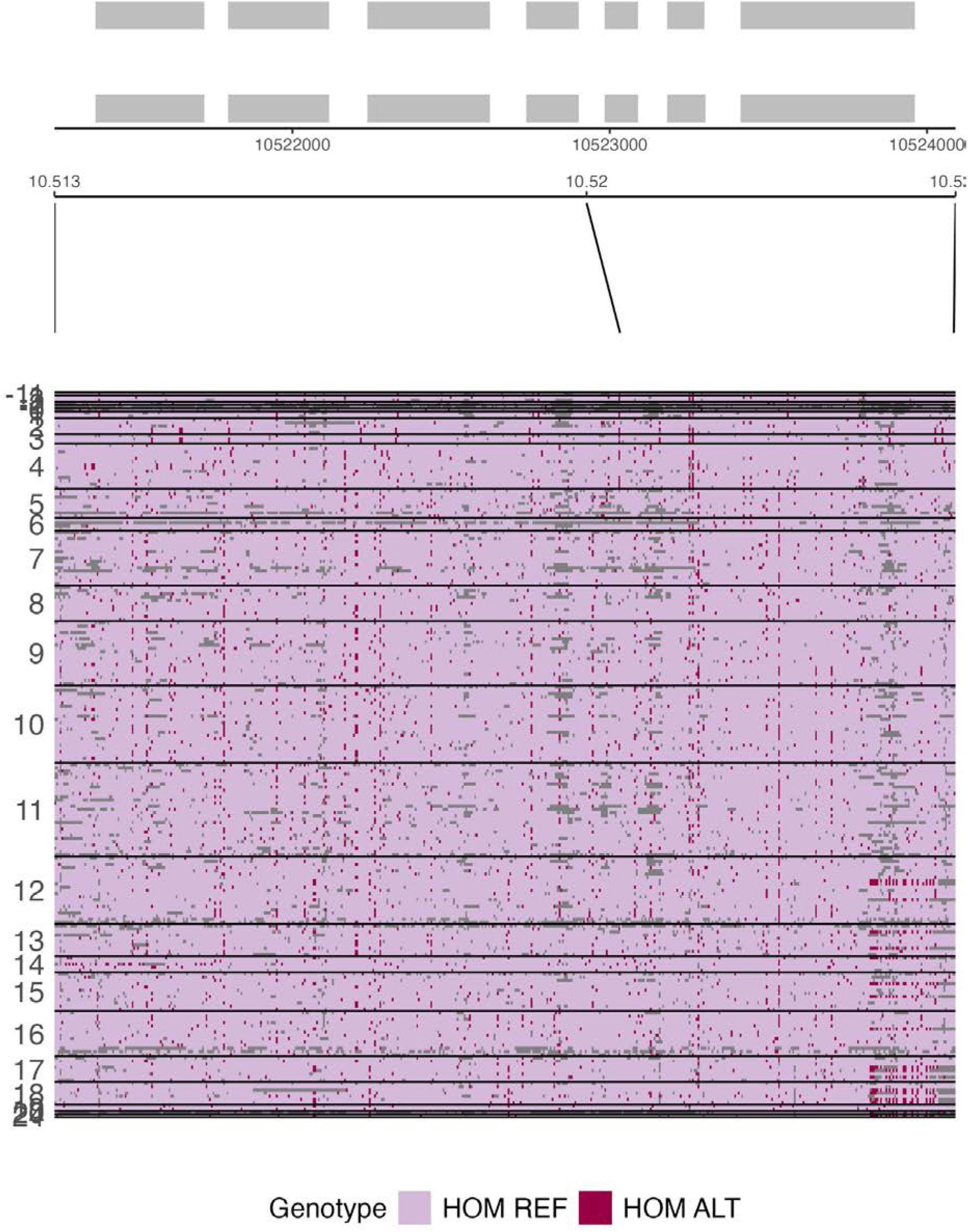
VCF visualization of *CYP707A1* (AT4G19230) Visualization of variant call format (VCF) table of 231 founder accessions showing reference or alternative genotypes, and grouped and sorted by annual temperature of origin (y-axis). TAIR10 gene models of *CYP606A1* are shown with exons in grey boxes.

**Figure S59.**
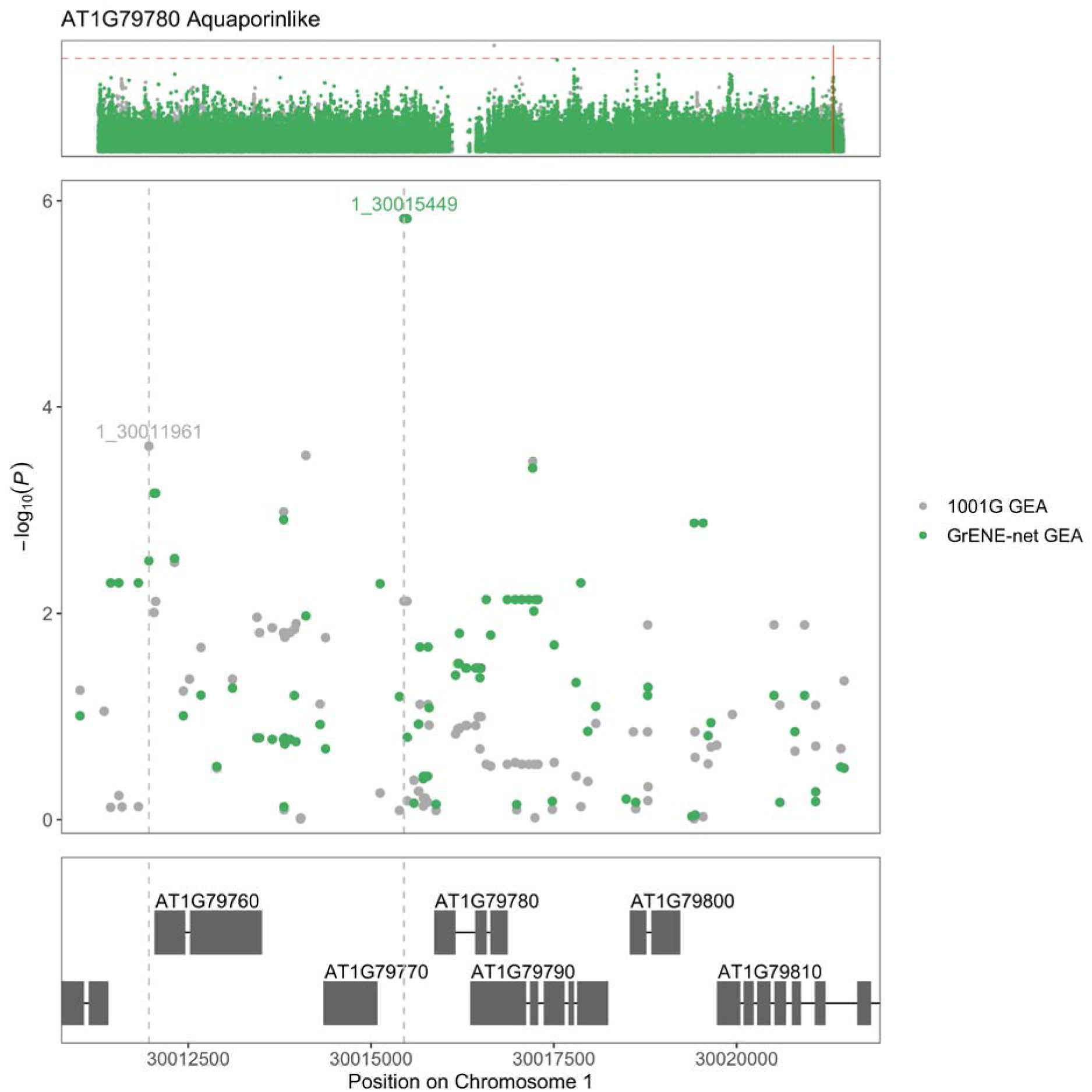
Manhattan plot of GEA with summer precipitation on Chromosome 1. Zoom in Chromosome 1 GrENE-net GEA LFMM conducted with summer precipitation (BIO18) about 424 bp away from Aquaporin-like protein gene start site (AT1G52180). This SNP is also downstream of an unknown CASP-like protein (*DUF1677*) (AT1G79770).

**Figure S60.**
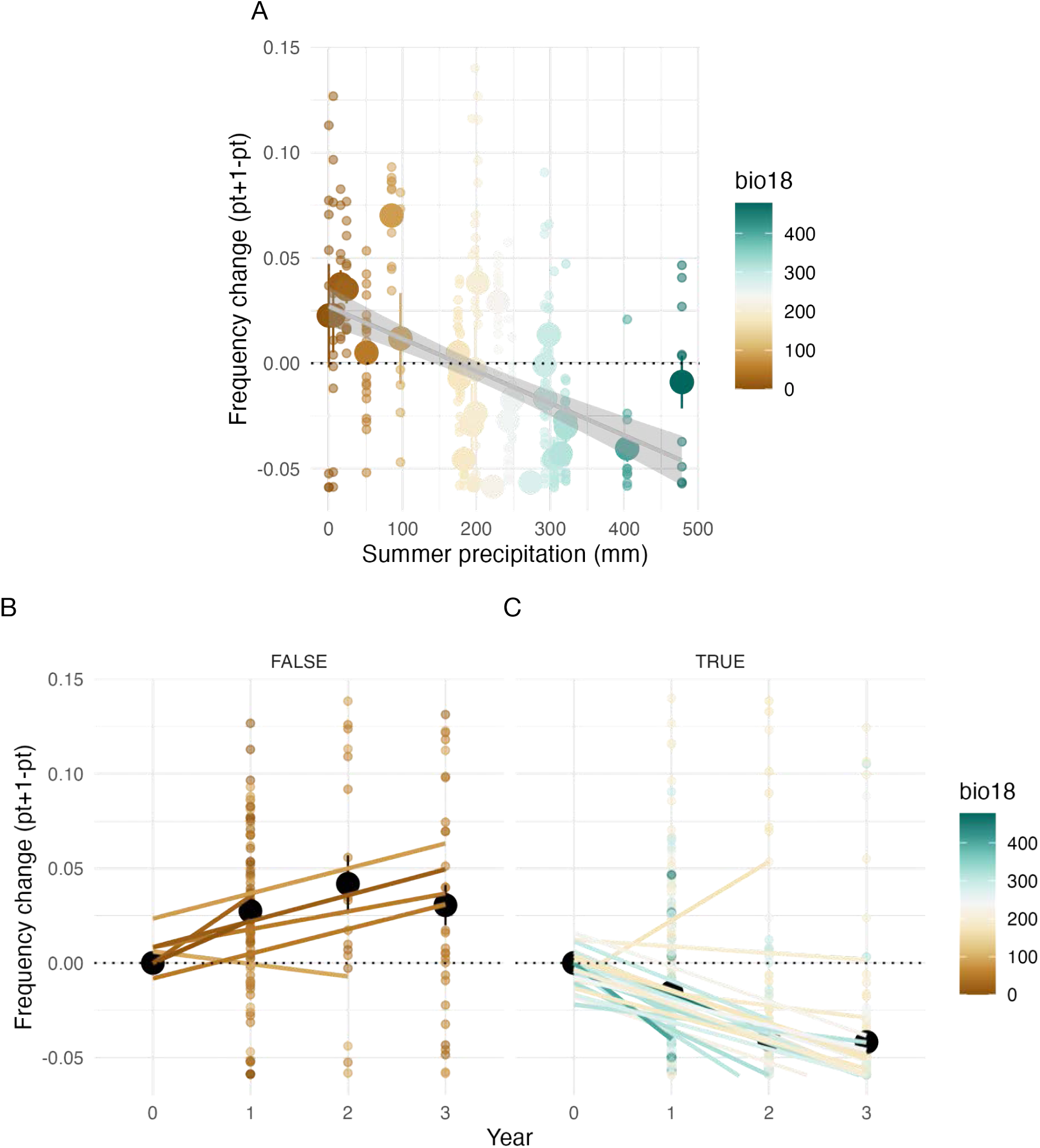
SNPs in Aquaporin-like protein (AT1G52180) change in frequency. (**A**) Correlation of allele frequency at top Aquaporin-like protein (AT1G52180) (bio18, summer precipitation [mm]). Temporal trends of top allele over years stratified by summer precipitation below 120 mm (**B**) or above (**C**).

### CROSS VALIDATION PREDICTABILITY

**Figure S61.**
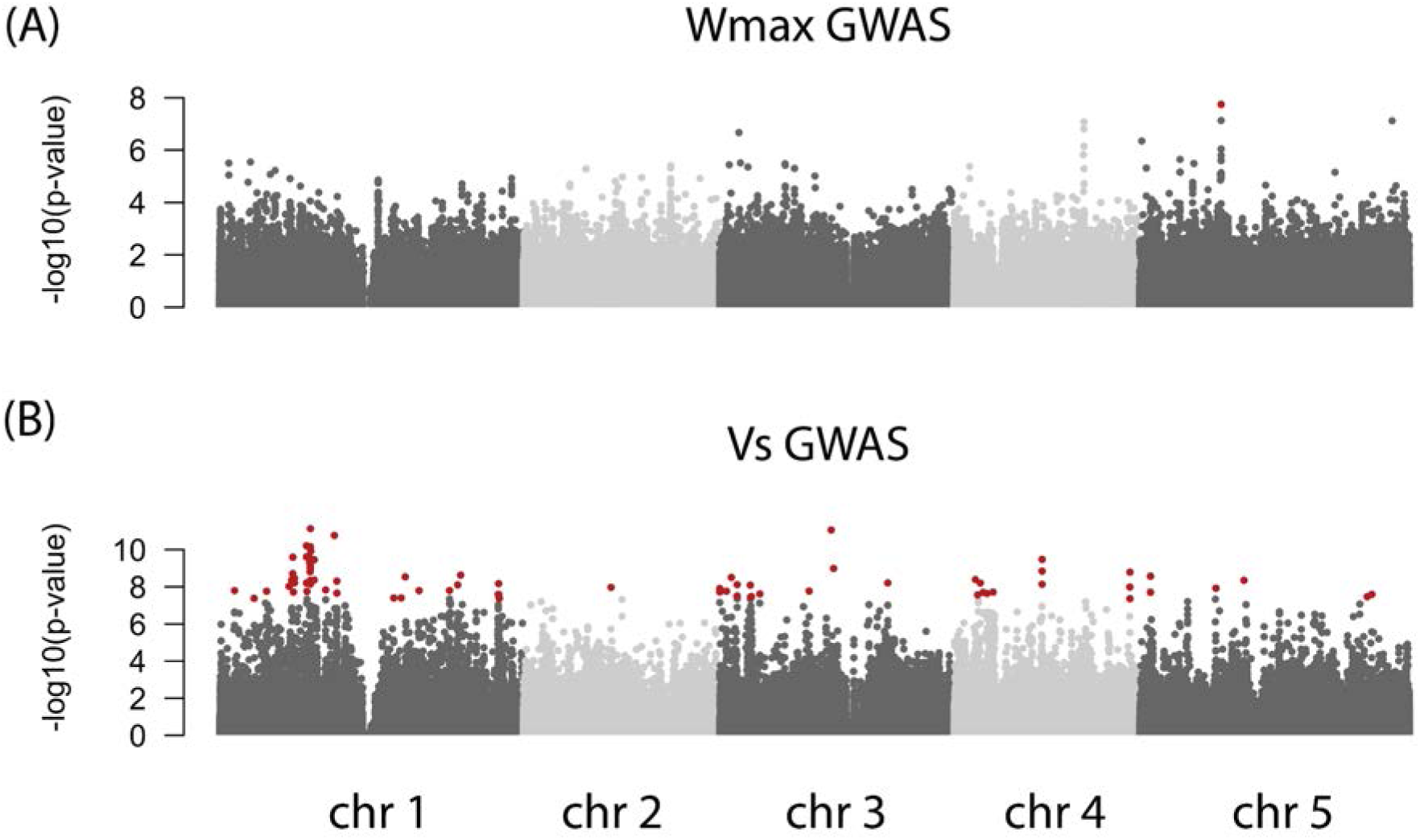
Genetic architecture of Gaussian stabilizing selection model parameters (**A**) Manhattan plot of estimated *W_max_* of 231 founder accessions. (**B**) Manhattan plot of estimated *V_s_* of 231 founder accessions. The red dots indicate significant SNPs after Bonferroni *P*-value adjustment. Variance explained was 91% (s.e 7.2%) for *Wmax* and 58.9% (s.e 16.5%) for *Vs*.

**Figure S62.**
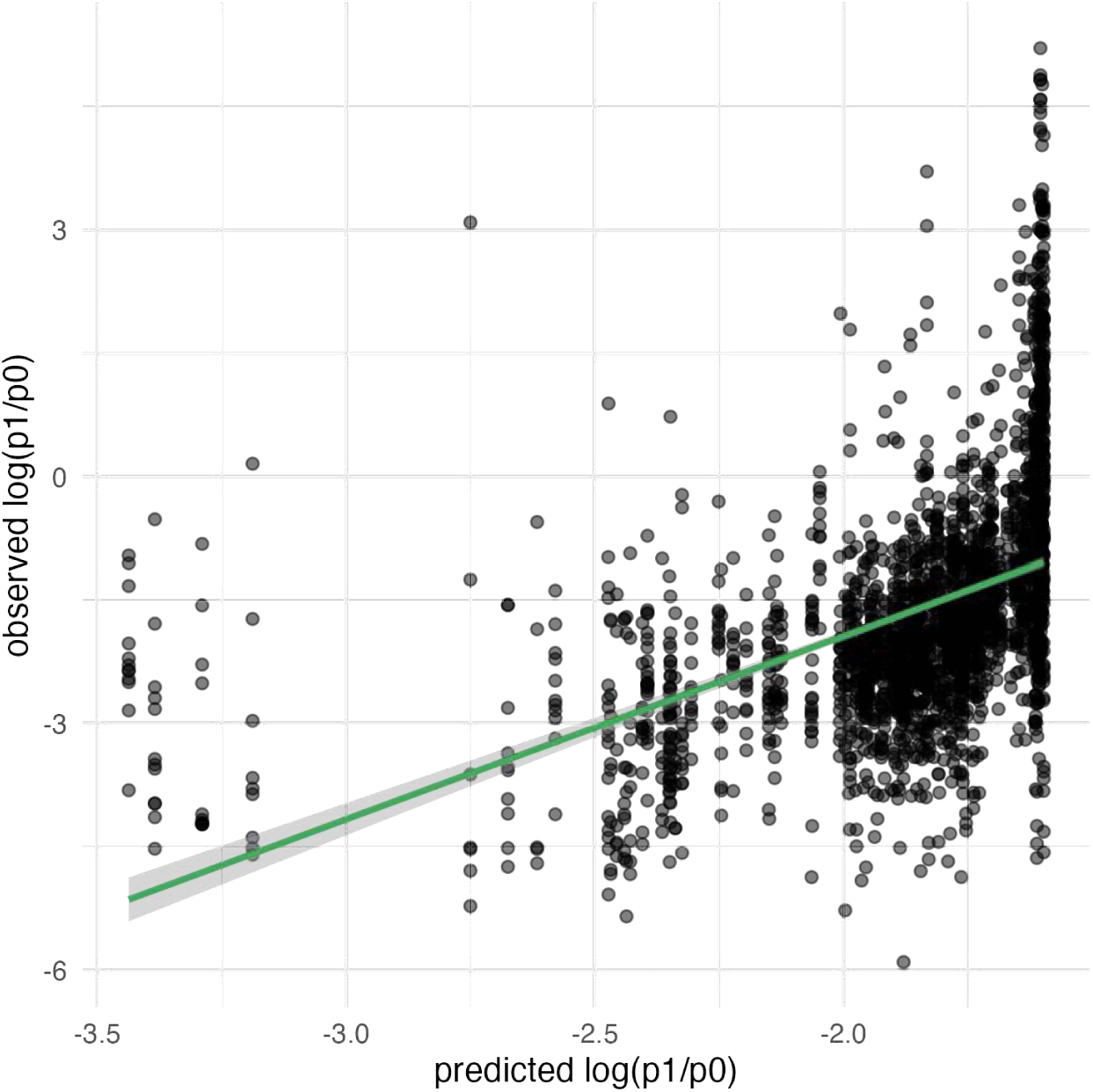
Predictability of terminal accession relative frequency in Cadiz, Spain. Fit of a stabilizing selection regression using *n*=29 sites and using site 4 (Cadiz, Spain) as testing site to visualize the predictability of accession relative frequency change.

**Figure S63.**
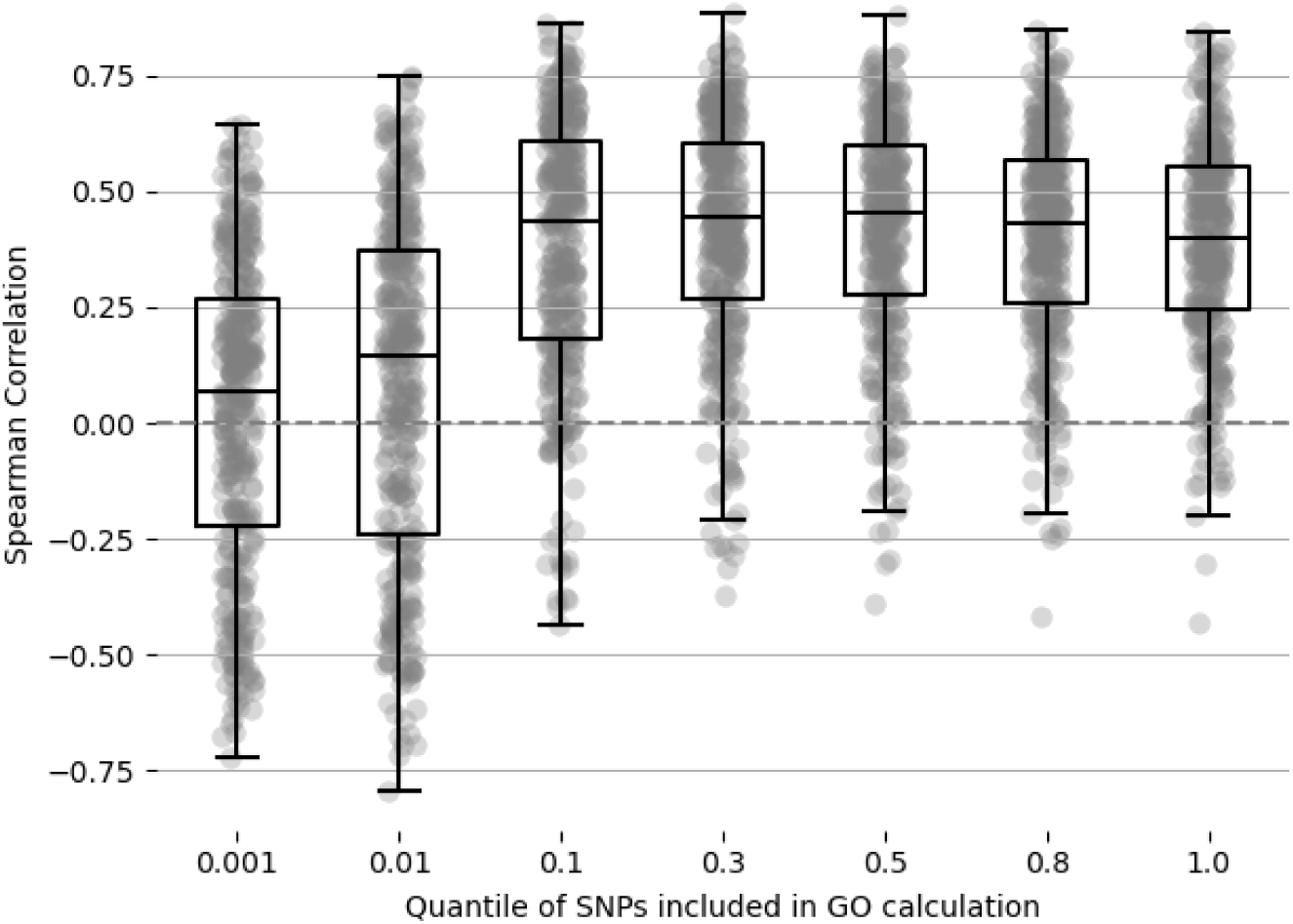
Predictive accuracy of accessions’ frequency change from GO models. Increasing the number of SNPs included in the calculation of Genomic Offset enhances the Spearman correlation with actual relative fitness of accessions. Each boxplot summarizes the correlations within specific SNP quantiles of *P-*values. Approximately the number of SNPs used in all leave-one-out cross-validation iterations were: 1,046 SNPs (quantile 0.001), 10k (quantile 0.01), 105k (quantile 0.1), 315k (quantile 0.3), 524k (quantile 0.5), 839k (quantile 0.8), and 1,049k (quantile 1.0).

**Figure S64.**
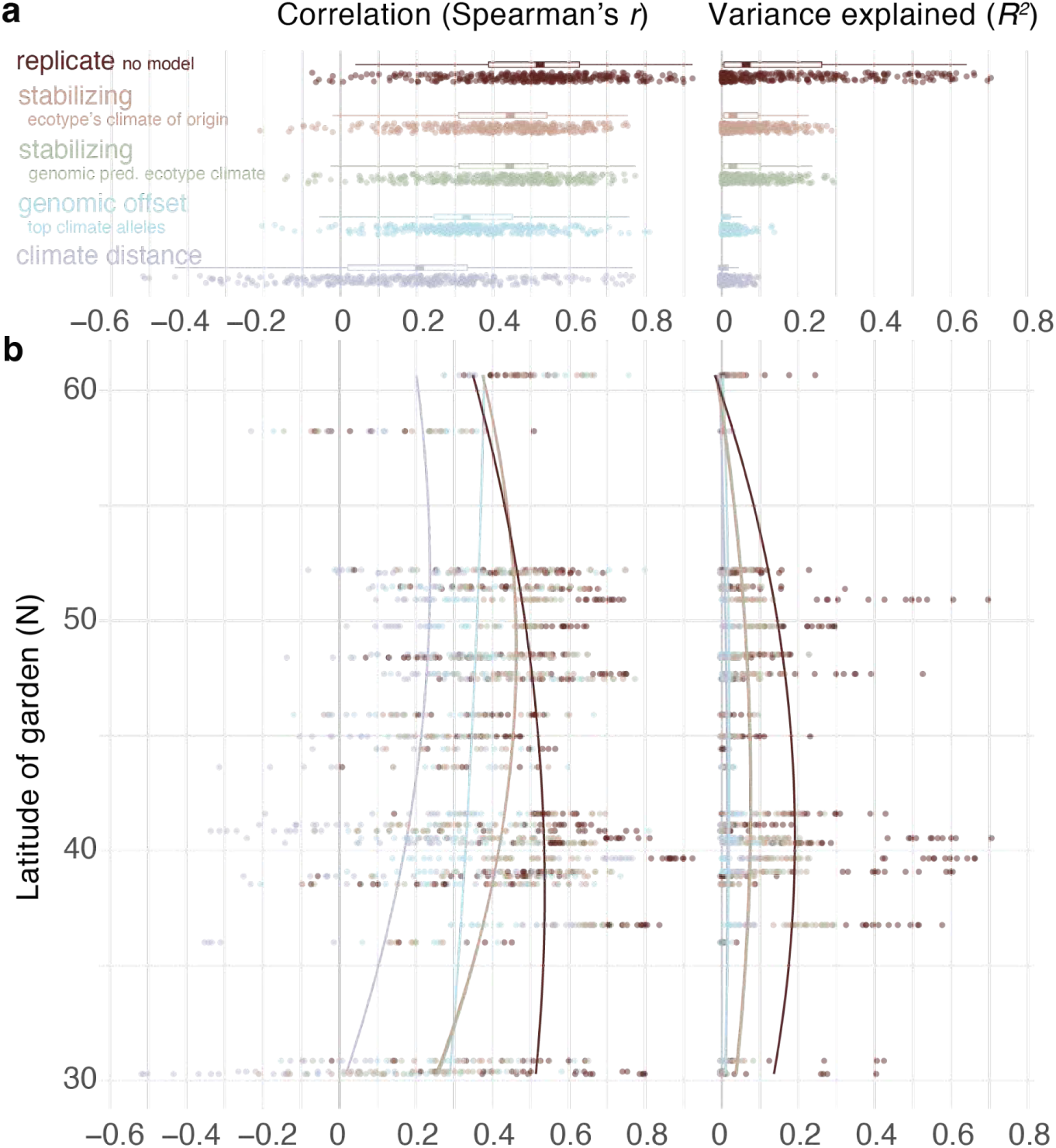
Predictability of accession relative frequency across model types. (**a**) Leave-one-out cross-validation of accession relative frequency change trajectories in one generation across all sites using different predictive models (*n=*325 predictions for each site and replicate). Stabilizing selection model of annual temperature using *V_s_ ^−1^* and *W_max_* per accession (n=231) to predict fitness declined by increased climate distance to origin (brown) or inferred from genomic-climate association (green), genomic offset quantifying missmatch using top SNPs identified in genome-wide climate (blue). As maximum reference, we use the predictability between replicates as the maximum possible predictability (dark brown), and as minimum baseline, we use a naive climate distance without any underlying evolutionary model (lila). (**b**) Leave-one-out predictability of each of the experimental gardens across a latitudinal gradient (n=5-12 surviving replicates per experiment). Curves are polynomial GLMs across all sites by latitude.

**Figure S65.**
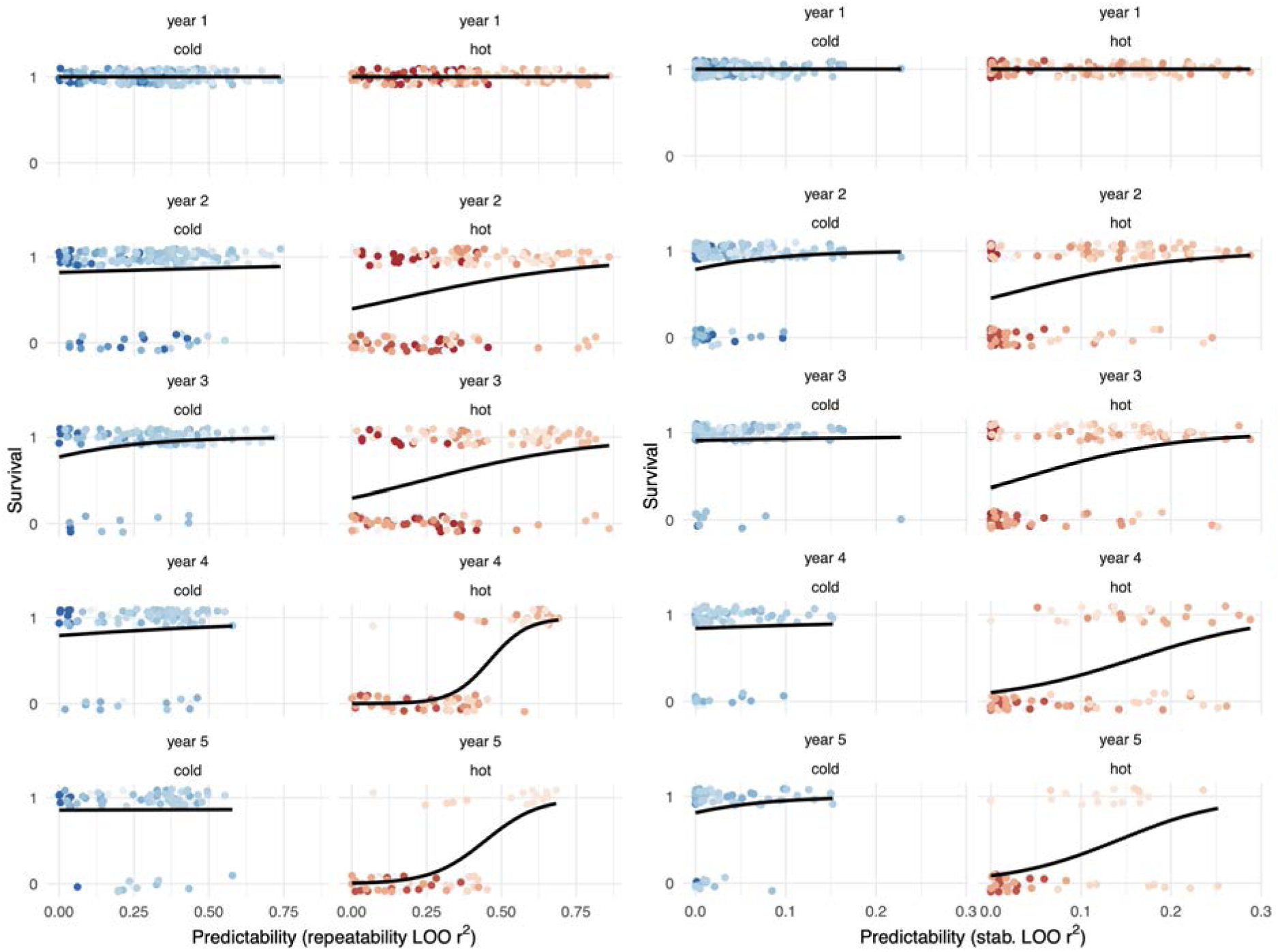
Evolutionary predictability and population survival over time. (**Left**) Logistic regressions of population survival across experiments classified as cold vs hot (∼10C threshold) over years. Color scale corresponds to annual temperature (BIO1). The predictor variable was predictability of evolutionary trajectories in a leave-one-out (LOO) cross-validation approach using other replicates within a garden as predictors. (**Right**) Same but using predictability from a genomic offset-like approach using stabilizing parameters, extended from main **Fig. 5**..

**Figure S66.**
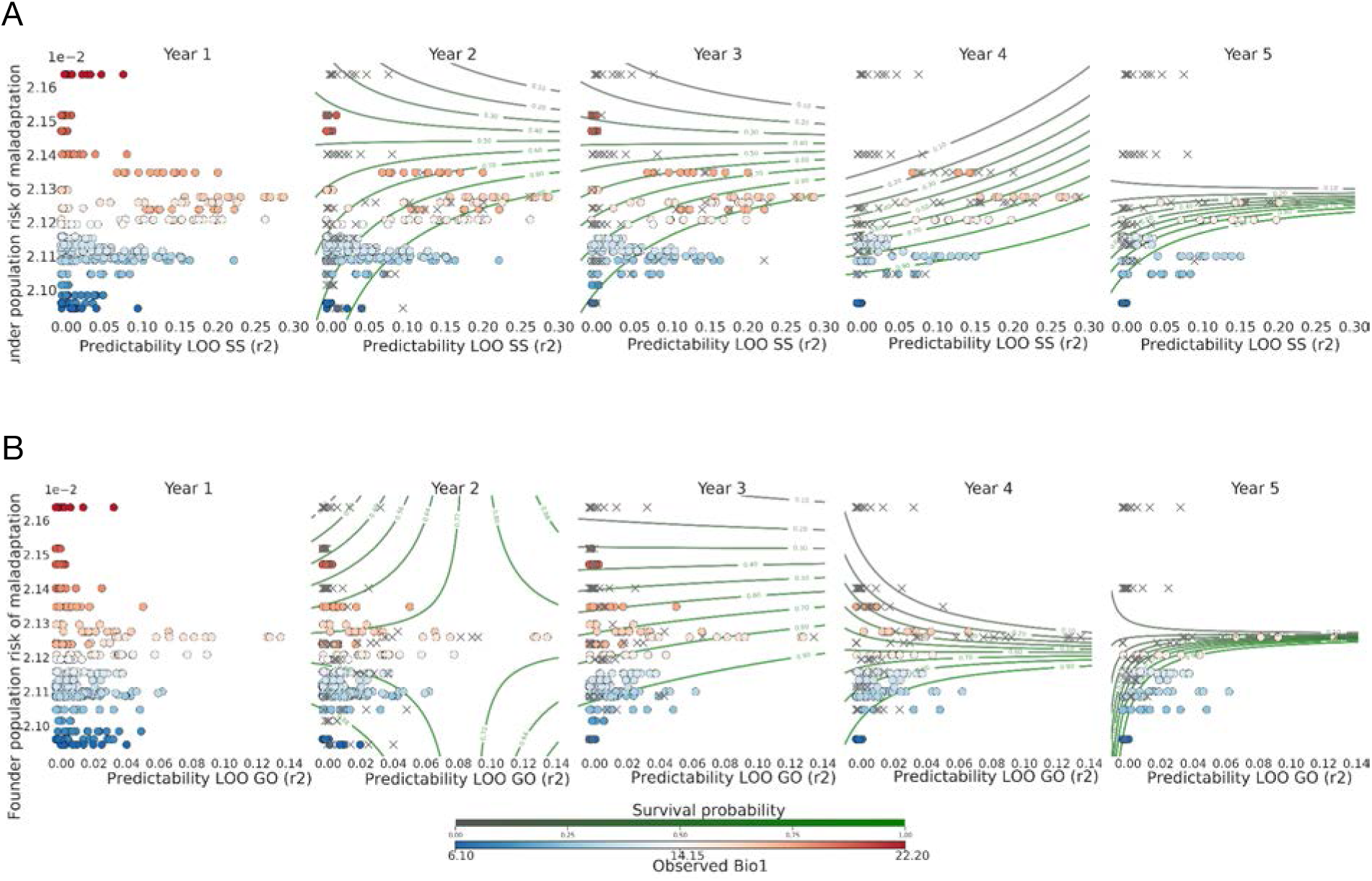
Survival probability surface GO predictions and evolutionary predictability. Isoclines of predicted survival probability as a function of first year predictability (Predictability LOO Stabilizing Selection model (*r^2^*) for panel A and Predictability LOO GO (*r^2^*) for panel B), maladaptation risk (GO) of the founder population and their interaction for five years. The contour lines represent the predicted probability based on a logistic regression model including green lines indicating higher predicted survival and grey lines indicating lower survival. Observed data points are shown as circles (colored by the mean annual temperature of the experimental site), and crosses indicate extinction of the plot. After Year 2, the variable GO became a significant predictor of survival (P < 0.001 for all subsequent years).

